# Rewiring of gene expression in circulating white blood cells is associated with pregnancy outcome in heifers (Bos taurus)

**DOI:** 10.1101/2020.03.18.997379

**Authors:** Sarah E. Moorey, Bailey N. Walker, Michelle F. Elmore, Joshua B. Elmore, Soren P. Rodning, Fernando H. Biase

## Abstract

Infertility is a challenging phenomenon in cattle that reduces the sustainability of beef production worldwide. Here, we tested the hypothesis that gene expression profiles of protein-coding genes expressed in peripheral white blood cells (PWBCs), and circulating micro RNAs in plasma, are associated with female fertility, measured by pregnancy outcome. We drew blood samples from 17 heifers on the day of artificial insemination and analyzed transcript abundance for 10496 genes in PWBCs and 290 circulating micro RNAs. The females were later classified as pregnant to artificial insemination, pregnant to natural breeding or not pregnant. We identified 1860 genes producing significant differential coexpression (eFDR<0.002) based on pregnancy outcome. Additionally, 237 micro RNAs and 2274 genes in PWBCs presented differential coexpression based on pregnancy outcome. Furthermore, using a machine learning prediction algorithm we detected a subset of genes whose abundance could be used for blind categorization of pregnancy outcome. Our results provide strong evidence that transcript abundance in circulating white blood cells is associated with fertility in heifers.

## INTRODUCTION

Cattle production systems provide approximately 28% ^1^ of the global protein supply. Improving cattle production efficiency is essential for farmers to attain sustainable production and support the growing demand for animal protein ^1^. Infertility is a critical problem in agriculturally important large animals. For instance, the prevalence of infertility ranges from 64% to 95%, averaging 15% in beef heifers ^2-14^. First breeding success greatly influences the lifetime efficiency of beef replacement heifers. Heifers that calve early in their first calving season experience increased productivity and longevity than their later calving herd mates ^15-18^. Furthermore, the genetic correlation between yearling pregnancy rate and lifetime pregnancy rate is high (0.92-0.97) ^19,20^. Therefore, the ability to identify heifers that experience optimal fertility during the first breeding is essential to the sustainability of beef cattle production systems.The genetic architecture of fertility in beef heifers is very complex ^21-28^. In line with this complexity, female reproductive traits have low heritability, with traits related to female reproductive fitness in cattle, such as heifer pregnancy and first service conception, range from 0.07 to 0.20 ^2,21,29-33^ and 0.03 to 0.18 ^2,21,29^, respectively. The examination of the genetic components of fertility in beef heifers have yielded several genes potentially associated with fertility traits ^21-28^; but the effect of these markers are minimal, and there is no clear redundancy in genetic markers identified across breeds or subspecies.

Beyond the genomic profiling, the analysis of multiple layers of an individual’s molecular blueprint is likely key for understanding the underlying biology of complex traits ^34^. In line with this rationale, expression-trait association studies have emerged as a means to better understand complex traits ^35,36^. Specifically related to fertility, there is evidence that circulating micro RNA (miRNA) profiles are indicative of reproductive function ^37^. Furthermore, we have previously identified differences in the transcriptome of peripheral white blood cells (PWBC) among heifers of different fertility potential ^38^.

In the present study, our objective was to investigate gene transcripts in PWBCs and circulating micro RNA (miRNA) profiles in heifers enrolled in fixed time artificial insemination (AI). We utilized a multi-dimensional approach to investigate differences in coexpression of protein-coding genes in PWBCs, and coexpression between circulating miRNA and mRNAs in PWBCs.

## RESULTS

### Overview of the experiment and data generated

We generated transcriptome data for mRNA from PWBC and small RNA from plasma collected at the time of AI from heifers that became pregnant to AI (AI-preg, n=6), pregnant later in the breeding season from natural breeding with bulls NB (NB, NB-preg, n=6), or failed to become pregnant (non-pregnant, n=5, Fig. 1A). We produced, on average, 25.8 and 7.1 million reads from mRNA and small RNA, respectively. Following the filtering for lowly expressed genes, 10496 protein coding genes in PWBC and 290 miRNAs in plasma samples were used for differential gene expression, coexpression and prediction analyses (Fig. 1A). Notably, three of the five non-pregnant samples clustered separately from other subjects based on the expression levels of protein-coding genes (Fig. 1B). We note that all heifers were phenotypically similar; and no marked distinction was observed among these three heifers relative to the remainder of the group on the day of AI. Contrariwise, there was no separation of samples based on the miRNA expression levels (Fig. 1C).

**Fig. 1.**
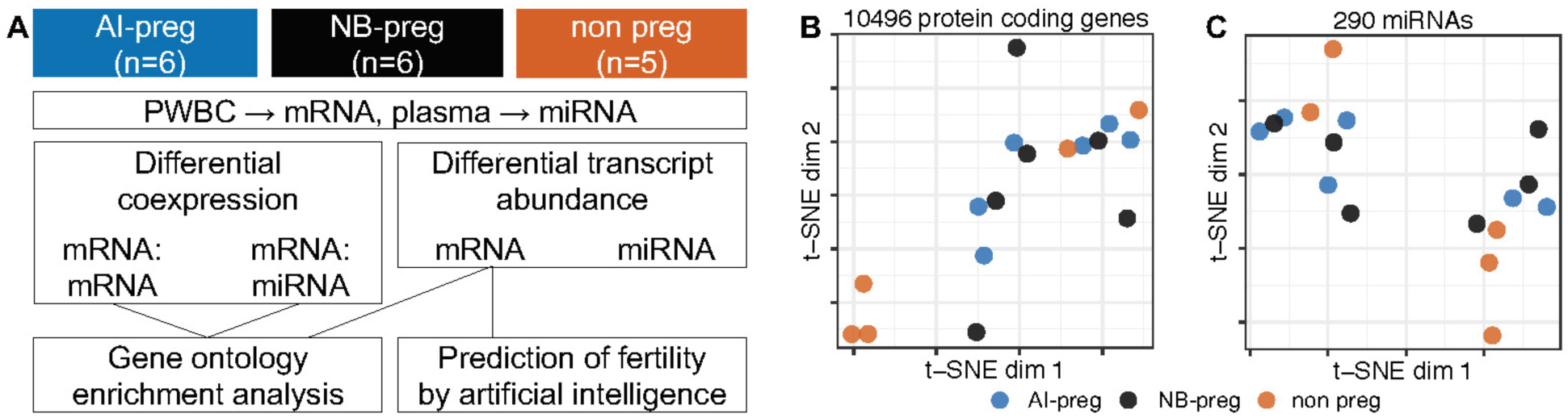
Overview of the experimental design and data. (A) Schematics of the experimental design, classification scheme based on fertility, data produced and overview of the analyses. (B,C) Dimensionality reduction of the transcriptome data obtained from the mRNA abundance of protein-coding genes in PWBCs and plasma extracellular miRNA.

### Differential mRNA:mRNA coexpression associated with pregnancy outcome

First, we asked whether there was a general coexpression profile of the protein-coding genes expressed in PWBCs amongst all 17 samples. We identified that 1633 genes that formed 2017 pairs with highly significant correlated expression levels (|r|>0.98, FDR<0.02, Fig. 2A, Supplementary Table S1). Notably, ∼99% of those significant correlations were positive (Fig. 2A). Several genes showing coexpression were associated with ‘translation’, ‘positive regulation of gene expression’, ‘B cell differentiation’, and ‘negative regulation of extrinsic apoptotic signaling pathway’ amongst other biological processes (FDR<0.1, Supplementary Table S2). This result indicates the occurrence of gene regulatory networks in PWBCs that are biologically meaningful for the immune system.

**Fig. 2.**
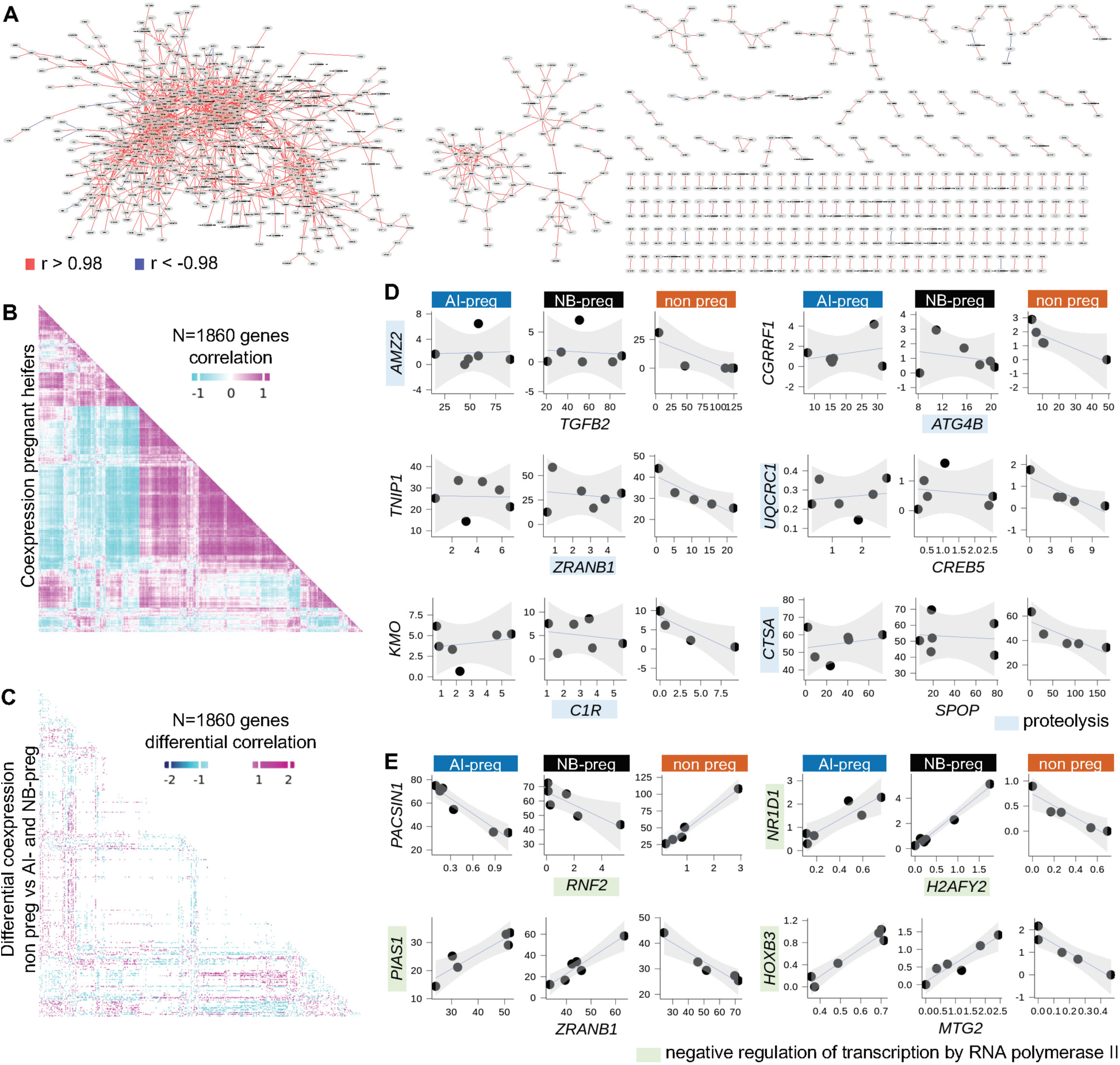
mRNA:mRNA coexpression in PWBCs. (A) Network of genes with highly correlated mRNA abundance. (B) Coexpression of 1860 protein-coding genes in pregnant heifers (AI-preg and NB-preg). (C) Profile of the differential coexpression in heifers classified as non-pregnant relative to the counterparts that became pregnant. (D) Scatterplot of selected genes associated with “proteolysis” that were coexpressed in heifers classified as non-pregnant relative to the counterparts that became pregnant. (E) Scatterplot of selected genes associated with “negative regulation of transcription by RNA polymerase II” that showed inverted coexpression in heifers classified as non-pregnant relative to the counterparts that became pregnant. All scatterplots present transcript abundance in Log2(counts per million).

Next, we investigated if the pair-wise gene coexpression differs between animals that became pregnant relative to those that did not become pregnant. We identified 1860 genes with significant differential correlation based on the pregnancy outcome (eFDR<0.002, Fig. 2B,C, Supplementary Table S3, Supplementary Fig. S1). Of these 1860 genes, 963 genes formed 716 negative, 1053 genes formed 708 positive, four genes lost two positive, and 40 genes formed 23 inverted coexpression connections in non-pregnant heifers relative to both AI-preg and NB-preg heifers. Among the 963 genes that formed new negative coexpression connections in non-pregnant heifers, there was enrichment of the biological processes “proteolysis” (n=38 genes) and “translation” (n=52 genes, FDR<0.1, see Fig. 2D for selected genes and Supplementary Table S4 for list of genes). Five of the 40 genes that formed inverted coexpression connection in not-preg heifers relative to the pregnant counterparts were associated with “negative regulation of transcription by RNA polymerase II” (FDR<0.1, see Fig. 2E for selected genes and Supplementary Table S5 for a list of genes). These results indicate that physiological factors associated with fertility or infertility act on the transcription protein-coding genes in PWBCs.

### Differential miRNA:mRNA coexpression associated with pregnancy outcome

The generation of data from small RNA (from plasma) and mRNA (from PWBCs) from the same subjects (n=17) permitted us to ask whether circulating miRNAs have correlated expression with protein-coding genes expressed in the PWBCs. We identified 141 pairs of miRNA:mRNA with |r|>0.85 (eFDR<0.0001, Supplementary Fig. S2) that were formed by 106 protein-coding genes and 33 miRNAs (Supplementary Table S6). Among the 106 protein-coding genes existing in high correlation with miRNA, we identified three significantly enriched biological processes, namely: “peptidyl-threonine phosphorylation”, “cilium assembly” and “regulation of gene expression” (Fig. 3A, Supplementary Table S7). Of notice, 30 of the 141 miRNA:mRNA pairs that presented negative correlation between the expression levels have been previously identified as interacting pairs in the miRWalk database (Fig. 3B). Additionally, bta-mir-744 and bta-mir-320a emerged as the two most connected miRNAs (13 and 6 connections, respectively). It is evident that circulating miRNAs have regulatory roles over transcripts of protein-coding genes expressed in the PWBCs.

**Fig. 3.**
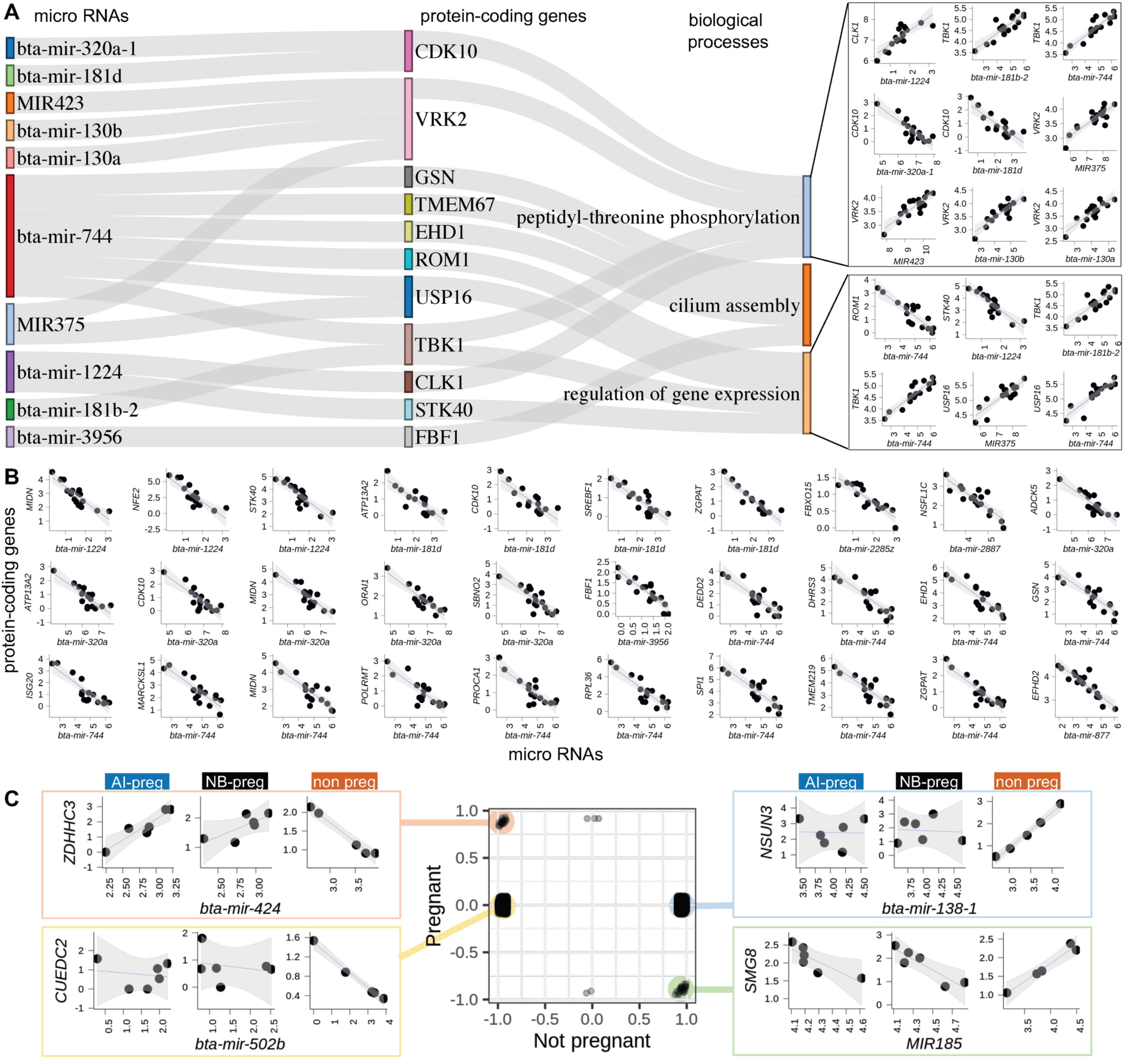
miRNA from plasma coexpressed with mRNA in PWBCs. (A) miRNA:mRNA pairs significantly coexpressed whose genes are significantly enriched in specific biological processes. (B) miRNA:mRNA pairs significantly coexpressed with evidence of interaction from miRBase. (C) The center chart presents pair-wise gene Pearson’s correlation for the corresponding group. The scatterplots to each side show selected miRNA:mRNA pairs representing differential coexpression between miRNA:mRNA in heifers classified as non-pregnant relative to the counterparts that became pregnant. All scatterplots presenting transcript abundance are in Log2(counts per million).

We also interrogated miRNA:mRNA pairs for differential coexpression associated with pregnancy outcome. We identified 2330 and 2738 miRNA:mRNA pairs with gain of negative and positive correlated expression, respectively, (|r|>0.9, eFDR≤0.01, Supplementary Fig. S3) in non-pregnant heifers that were not identified compared to pregnant heifers (both AI-preg and NB-preg, Fig. 3C). In addition, there were 51, two and three connections showing inverted, loss of negative and loss of positive correlations in non-pregnant heifers relative to the pregnant (both AI-preg and NB-preg) counterparts (Fig. 3C, see Supplementary Table S8 for genes). Among the 1220 protein-coding genes associated with gain of negative correlation with miRNAs, there were biological processes significantly enriched, namely: “immunoglobulin mediated immune response”, “negative regulation of DNA-binding transcription factor activity”, “lymphocyte chemotaxis” and “mitochondrial respiratory chain complex I assembly” (FDR<0.1, Supplementary Table S9). Among the 1288 genes associated with gain of positive correlation with miRNAs, the following biological processes were significantly enriched: “regulation of DNA replication” and “cellular response to BMP stimulus” (FDR<0.1, Supplementary Table S10). These results indicate that several miRNA and mRNA pairs have their connections altered in relation to the heifer’s fertility potential.

### Differential gene expression associated with pregnancy outcome

Differential gene expression analysis revealed 67 (55 up- and 12 down-regulated) and 81 (63 up- and 18 down-regulated) protein-coding genes with altered expression profiles between PWBC of AI-preg versus non-pregnant, and NB-preg versus non-pregnant heifers, respectively (eFDR≤0.05, Fig. 4A, Supplementary Tables S11 and S12, Supplementary Fig. S4). Notably, 26 genes were common between both pregnant groups versus the non-pregnant group (Fig. 4B). Gene ontology and pathway analyses of differentially expressed genes between AI-preg and Non-pregnant heifers revealed significant enrichment for the terms “signal transduction” (*ARAP3, LSP1, NFAM1, OR12D2, RARA*; FDR<0.05). When differentially expressed genes between NB-preg and non-pregnant heifers were analyzed, significant enrichment was determined for the terms “signal transduction” (*ARAP3, HRH2, LSP1, TIAM2, TNFRSF17, TNFRSF1A*; FDR=0.062) and the KEGG Pathway “cytokine-cytokine receptor interaction” (*IFNGR2, IL5RA, LTB, TNFRSF17, TNFRSF1A*; FDR=0.0045). Plasma miRNA profiles were very similar among samples from all pregnancy outcomes with the exception of miR-11995 that was more abundant in AI-preg versus non-pregnant heifers. These results indicate that quantitative differences exist in transcript abundance of protein-coding genes expressed in PWBC relative to the pregnancy outcome.

**Fig. 4.**
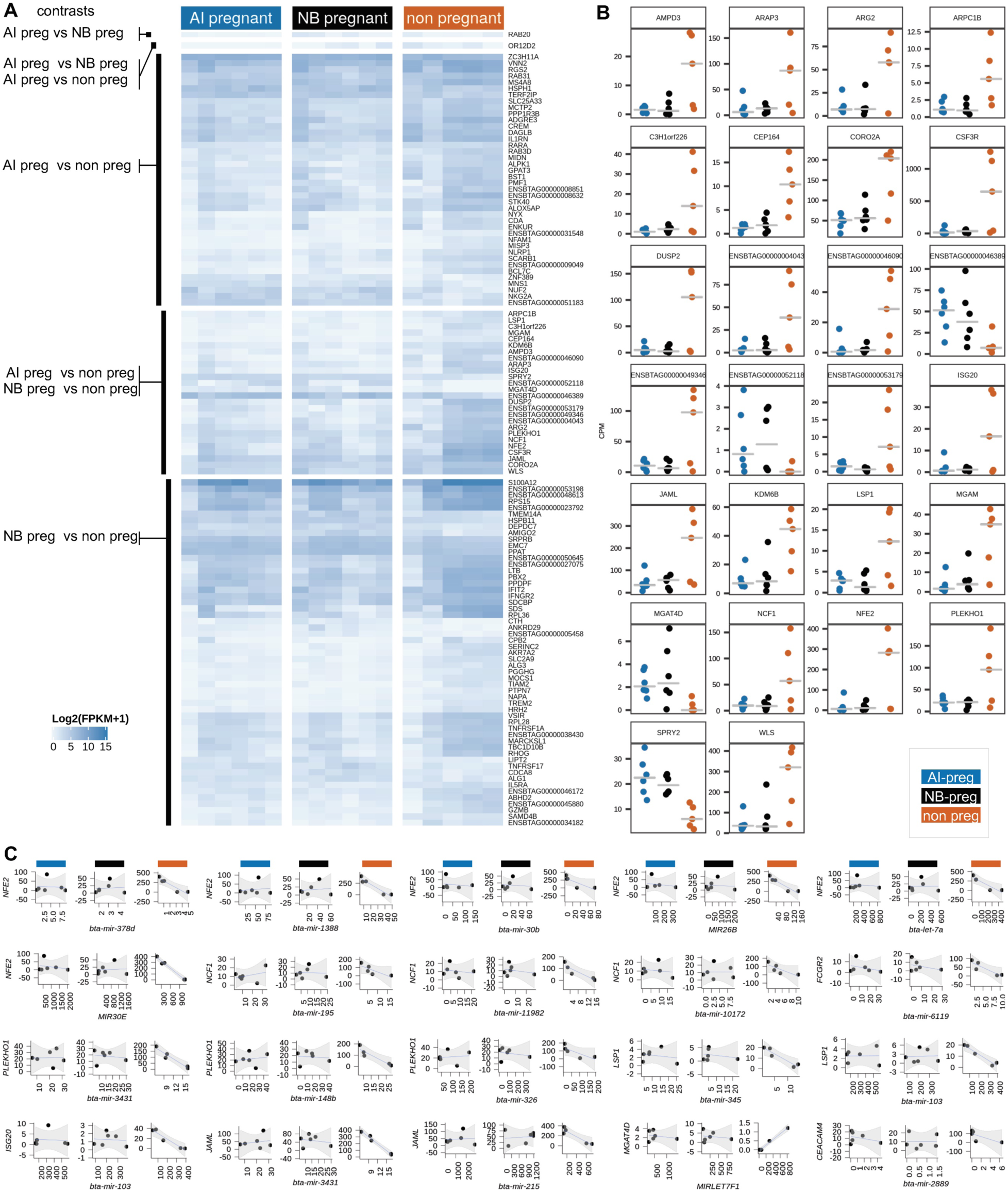
Differential coexpression among heifers of different reproductive outcome. (A) Heatmap depicting the expression levels of genes differentially expressed according to specific contrasts. (B) Transcript abundance in counts per million of the genes differentially expressed on both contrasts: AI-preg vs non-pregnant and NB-preg vs non-pregnant. (C) miRNA:mRNA pairs significantly coexpressed whose protein-coding genes are differentially expressed in PWBCs based on pregnancy outcome. All scatterplots present transcript abundance in Log2(counts per million).

Next, we asked if differential gene expression could be associated with differential miRNA:mRNA coexpression. An intersection of the results identified 12 differentially expressed genes (10 annotated genes) in non-pregnant heifers relative to the AI-preg and NB-preg groups that also gained coexpression with 27 miRNAs (Fig. 4C). Notably, all genes that had higher expression in non-pregnant heifers (*CEACAM4, FCGR2, ISG20, JAML, LSP1, NCF1, NFE2, PLEKHO1*) showed a gain in negative coexpression with specific miRNAs. By contrast *MGAT4D* was more lowly expressed in non-pregnant heifers, and gained a positive coexpression with a miRNA in non-pregnant heifers.

### Assessment of transcript levels as predictors of pregnancy outcome

Motivated by the observations that several genes have their abundance altered in relation to the fertility potential pregnancy (Fig. 4A), we asked whether transcript abundances could serve as predictive features across datasets. We previously generated gene expression values for a similar dataset (GSE103628 ^38^), which also consisted of transcriptome data from PWBCs collected from heifers at the time of AI, and were classified according to their pregnancy outcome as AI-preg (n=6) or non-pregnant (n=6). This dataset was particularly fitting for this interrogation because the sampling occurred at the same site in the year prior. We combined the expression values of the previously produced dataset (year one) with those of the current dataset (year two) and contrasted the transcript abundance between heifers that were categorized as AI-preg versus non-pregnant accounting for the different years as a fixed effect in the model. This analysis identified 198 genes whose abundance were significantly different between the two experimental groups (AI-preg, non-pregnant, FDR<0.03, Fig. 5A, Supplementary Table S13).

**Fig. 5.**
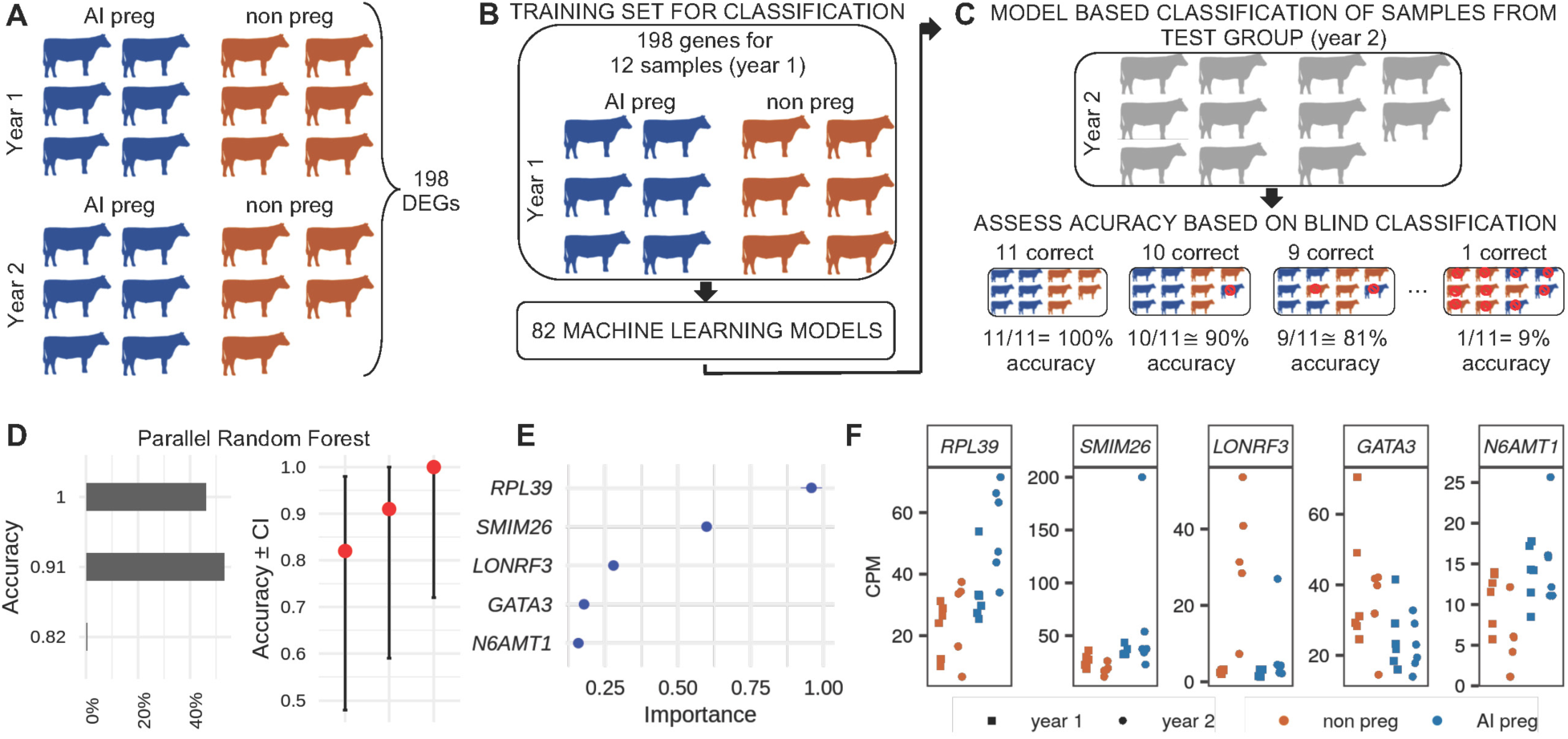
Prediction of pregnancy outcome based on mRNA levels and machine learning algorithms. (A) Representation of the contrast between gene expression levels of heifers classified as AI-preg and non-pregnant from two years, and the number of differentially expressed genes (DEGs). (B) Depiction of the training set used as input for the modeling of the transcriptome dataset by multiple machine learning algorithms. (C) Depiction of the data used as the test set for prediction of pregnancy outcome, and the representation of the calculation of accuracy. (D) Accuracy of blind pregnancy prediction based on the algorithm “parallel random forest” in 2000 trials. (E) Variable importance for the top five genes. (F) Transcript levels for the top five genes most informative for the machine learning algorithm to classify pregnancy outcome.

We used the transcript abundance values for these 198 genes obtained from samples collected in year one as input (training set) for machine learning algorithms to derive models of prediction to discriminate pregnancy outcome (AI-preg or non-pregnant, Fig. 5B). Next, we used the transcript abundance values for these 198 genes obtained from samples collected in year two as input to the algorithms for a blind classification of pregnancy outcome (AI-preg or non-pregnant, Fig. 5C). The accuracy of classifications was assessed based on the known pregnancy outcome, and parallelized random forest was the algorithm that produced the greatest proportion of correct classification of all 11 heifers tested (926 out of 2000 predictions, 46.3%), followed by 1062 out of 2000 predictions with correct classification for 10 out of 11 heifers (53.1%, Fig. 5D). Notably, the top five genes that showed the largest degree of importance for the classification were *RPL39, SMIM26, LONRF3, GATA3, N6AMT1* (Fig. 5E), of which, *RPL39, SMIM26, N6AMT1* were up-regulated and *LONRF3* and *GATA3* were down-regulated in AI-preg heifers relative to the non-pregnant group (Fig. 5F).

## DISCUSSION

Here, we report a multilayered transcriptome analysis of transcripts from blood sampled on the day of AI. The main objective of our study was to understand gene regulatory networks ^39^ occurring in PWBCs, and rewiring that can occur based on the heifers’ fertility potential. We focused on the day of artificial insemination because it is the last day prior to the fertilization, when the heifer has the first opportunity to become pregnant. We interrogated mRNA and miRNA abundance levels obtained from the same individuals using the coexpression framework, which is a valuable systems biology approach to gain insights into the biology of complex traits ^39^. Furthermore, using machine learning ^40^, we detected consistent altered gene transcript abundance associated with infertility across independent datasets. Collectively, our results provide important aspects of regulatory networks in PWBCs that shed light on the physiological status of an animal and its preparedness for becoming pregnant.

We must note that the empirical results reported in this study must be considered in light of some limitations. Here, we evaluated transcripts present in a subset of cells (white blood cells) in one tissue (blood). It is important to acknowledge that the data generated does not capture the totality of the complexity involving infertility, nor the multitude of factors influencing infertility. Furthermore, this is a snapshot of one day in the heifer’s life which we are using to make inferences on the first breeding season. Thus, the results presented here cannot be extrapolated to the consecutive breeding seasons in the heifer’s life. Finally, the sample size does not capture all the variability existent within cattle. Thus, we do not generalize the results beyond the scope of this research.

Similar to work reported in humans ^41^, our gene network analysis focusing on mRNAs detected several protein-coding genes with strong coexpression in PWBCs. Interestingly, nearly all of the correlated expression inferred as statistically significant across all heifers (|r|>0.98, FDR<0.02) were positive. As expected ^42^, in silico functional gene co-expression analysis indicated groups of genes that are linked to specific biological processes. A contrast of coexpressing pairs of genes between samples collected from non-pregnant heifers and the two groups of heifers that became pregnant displayed a large number of connections present in non-pregnant heifers that were not identified in either AI-preg or NB-preg heifers. Our strict criteria (|r|>0.99, eFDR<0.002) utilized to identify these altered connections indicate functional regulatory relationships between selected genes.

Surprisingly, a great majority of the altered connections between gene pairs present were gained in non-pregnant heifers, while there was minor loss of significant connections in non-pregnant heifers relative to the AI-preg and NB-preg groups. The enrichment for specific biological processes of genes that formed new negative connections was clear evidence of the biological relevance of these new connections. Notably, among genes annotated with proteolytic activity, *ADAM19* ^43^, *ASRGL1* ^44^, *ATG4B* ^45^, *DPEP2* ^46^, *IMMP2L* ^47^, *PSMB9* ^48^, *SERPINE2* ^49^, *UCHL3* ^50^, *USP42* ^51^ have been implicated in female infertility. A more drastic change in patterns of coexpression was observed with the inversion of gene connections (significant pairwise coexpression in one group with significant coexpression with opposite direction in another group). These gene circuits changing from activation to repression are explained by current models of incoherent feed-forward loops ^52^. Cumulatively, the large occurrence of new connections and the inverted connections created in PWBCs exclusively in non-pregnant heifers indicate that gene regulation is altered as an adaptative response to a stimulus.

Our profiling of small RNAs from plasma identified 263 annotated miRNAs, out of which 131 and 122 were previously identified in sera and circulating exosomes, respectively, among other cattle organs ^53^. Only 38 annotated miRNAs were profiled in peripheral blood mononucleated cells in humans ^54^, which demonstrates that our plasma samples were depleted of circulating white blood cells.

Immuno-related cells composing the PWBC population can uptake circulating small RNAs ^55^. Although our data did not allow us to trace the origin of the tissue that produced the miRNAs, we detected 33 circulating miRNAs whose abundance are associated with the abundance of 106 protein-coding genes. As reported in humans ^56^, we quantified positively and negatively correlated abundances between miRNA and protein-coding genes. Negative correlations mostly occur due to the targeting of mRNA by a specific miRNA. Indeed, we found several pairs of miRNA:mRNA negatively correlated that had been identified as interacting pairs that fit this model. By comparison, the positive coexpression between miRNA:mRNA pairs can be explained by a miRNA targeting an intermediate gene, which in turn is negatively associated with a third gene, thereby creating a positive correlation between a miRNA:mRNA pair (see Fig. 3 in reference ^57^). In our results, we only identify examples of miRNA:mRNA pairs that follow the intermediate action of miRNAs when relaxing the stringent criteria for inference of significant coexpression. Regardless of the model of interaction, it was notable that many protein-coding genes that were coexpressed with miRNAs were enriched for specific biological processes, including “regulation of gene expression”. These results add further evidence to the growing body of literature suggesting that circulating miRNAs act as mediators of cellular function ^58,59^.

We also tested whether circulating miRNA and PWBC mRNA pairs have differential coexpression relative to the pregnancy outcome. Following the same trend observed for miRNA:mRNA coexpression in PWBCs, there were 1000-fold more miRNA:mRNA connections that were present in heifers that did not become pregnant compared to the loss in coexpression. Results from gene enrichment analysis support the notion that genes forming new miRNA:mRNA coexpression connections in non-pregnant heifers function in the immune system, although our data do not permit inferences on whether particular immune processes were more or less functional in non-pregnant heifers.

Critical aspects of female fertility are tied to an appropriate regulation of the immune system. Maternal immune tolerance to pregnancy starts with the exposure to proteins in the semen ^60^, and is carried throughout placentation and pregnancy establishment ^61^. Notably, 24 out of 26 differentially expressed genes between pregnant (AI-preg and NB-preg) and non-pregnant heifers showed greater abundance in non-pregnant heifers relative to the other two groups. (AI-preg and NB-preg). While we did not detect differential abundance of miRNAs relative to pregnancy outcome, the gain of coexpression between miRNAs and mRNAs from differentially expressed genes functionally linked with immunology is in line with regulatory roles that certain miRNAs exert on the immune system ^62^. Specifically, bta-miR-30b ^63^, bta-miR-148b, bta-miR-195 ^64^, bta-miR-326 ^65^, bta-miR-378d ^66^, bta-miR-1388 ^63^, bta-miR-2889 ^66^, bta-miR-11982 ^67^ have been reported to be differentially expressed in pathogen infected cells or tissues relative to control counterparts. This result lead us to hypothesize that the heifers in the non-pregnant group have a heightened activity of specific cells or functions in the immune system relative to the heifers that became pregnant. It remains unclear, however, if this possible higher activity was stimulated by extrinsic factors or is within the naturally-occurring variation ^68^.

The results of differential gene expression and differential coexpression fostered a hypothesis that transcript abundance in the PWBCs can be used as predictors of pregnancy outcome. We tested this hypothesis by applying machine learning algorithms on two groups of heifers that were bred in 2015 (year one) and 2016 (year two). Parallel random forest emerged as the algorithm with over 90% efficiency of classification nearly all trials executed, which confirms the potential of accurate classification of samples using RNA-seq data under the case-control framework ^69,70^. The results show that while not one single gene emerges as a potential biomarker, the accumulated information of transcript abundance from multiple genes can be powerful for the identification of fertility potential in cattle.

Although we detected altered coexpression with high confidence, and in some instances, with significant enrichment of genes in specific gene ontology terms, the consequence of the differential coexpression on those biological functions remains undetermined. In addition, it remains unclear whether the altered patterns of gene expression profiles are connected to immune functions that are related to infertility ^71^. The data analyzed here does not allow us to infer causality ^72^, however our results converge to a strong indication that differences in gene expression profiles are associated ^72^ with fertility in cattle.

In conclusion, analyses of mRNA from PWBCs and circulating extracellular miRNA by multiple approaches (differential gene expression, differential gene coexpression and machine learning classification) provided complementary evidence that gene expression profiles in the PWBCs are associated with the fertility potential in beef heifers. The overall difference detected in transcript abundance for specific genes is reproducible across samples and show promise for the development of molecular features indicating fertility potential in beef heifers. The analysis of transcriptome data is critical for the understanding of economically important complex traits^73^; and our findings show that transcript levels may be relevant for selection of animals more likely to contribute to a sustainable beef production system.

## METHODS

All procedures with animals were approved by Institutional Animal and Care and Use Committee at Auburn University. All methods related to animal handling were performed in accordance with the guidelines stipulated by the Institutional Animal and Care and Use Committee at Auburn University and the Guide for the Care and Use of Laboratory Animals.

### Heifer classification according to pregnancy outcome

Crossbred heifers (n=32, Angus-Simmental, *Bos taurus taurus*) were developed as replacement heifers ^74,75^ at the Black Belt Research and Extension Center (Auburn University). All animals were exposed to the same vaccination protocol which included an injection of ULTRABAC® 7 and BOVI-SHIELD GOLD® FP™ 5 L5 at approximately 6 months of age followed by a second injection of each product at approximately 9 months of age. A final injection of BOVI-SHIELD GOLD® FP® 5 VL5 HB was administered to all heifers at approximately 12 months of age. Thirty-four days prior to AI, heifers were evaluated for reproductive tract score (scale of 1-5 ^76^) and body condition score (BCS; scale of 1-9 ^77^) for the assertion that morphological and physiological conditions were suitable for breeding. One animal was eliminated from the breeding herd at this time due to a reproductive tract score of 2. The remaining animals presented body condition score 6 and reproductive tract score 3 (n=9), 4 (n=14) or 5 (n=8).

Thirty one animals, 14 months of age on average, underwent estrous synchronization for fixed-time artificial insemination utilizing the 7-Day Co-Synch + CIDR protocol ^78^. An experienced veterinarian inseminated all heifers with semen from one sire of proven fertility (Deer Valley All In, Select Sires). Fourteen days after insemination, heifers were exposed to two fertile bulls for natural breeding for 60 days. An experienced veterinarian performed pregnancy evaluation by transrectal palpation on days 69 and 120 post artificial insemination. Presence or absence of a conceptus, alongside morphological features indicating fetal age were used to classify heifers as pregnant to AI (AI-pregnant), pregnant to natural breeding (NB-pregnant), or non-pregnant. At the end of the breeding season, 16, nine and six heifers were classified as AI-pregnant, NB-pregnant or non-pregnant, respectively.

### Collection, processing and preservation of biological material

Immediately after AI, 10ml of blood was drawn from the jugular vein into vacutainers containing 18mg K2 EDTA (Becton, Dickinson and Company, Franklin, NJ). The tubes were inverted to prevent blood coagulation and immersed in ice for approximately 4 hours. Plasma and buffy coat were separated by centrifugation for 10 minutes at 2000×g at 4°C. Plasma was removed (∼1.5ml) and centrifuged 10 minutes at 500×g at 4°C for further removal of cells. The plasma devoid of cells was then transferred to a new microcentrifuge tube and preserved at - 80°C. The buffy coat was removed and deposited into 14ml of red blood cell lysis solution (0.15 M ammonium chloride, 10mM potassium bicarbonate, 0.1mM EDTA, Cold Spring Harbor Protocols) for 10 minutes at room temperature (24-25°C). After a centrifugation, 800xg for 10 minutes, the solution was discarded and the pellet containing PWBCs was re-suspended in 200 μl of RNAlater® (Lifetechnologies™, Carlsbad, CA). PWBC samples were stored at -80°C.

### RNA extraction, library preparation, and RNA sequencing

Total RNA was isolated from PWBCs of 18 heifers (AI-pregnant (n=6), NB-pregnant (n=6), and non-pregnant (n=6)) using TRIzol™ reagent (Invitrogen, Carlsbad, CA) ^79,80^ following the manufacturer’s protocol. RNA yield was quantified using the Qubit™ RNA Broad Range Assay Kit (Eurogene, OR) on a Qubit^®^ Fluorometer, and integrity was assessed on an Agilent 2100 Bioanalyzer (Agilent, Santa Clara, CA) using an Agilent RNA 6000 Nano kit (Agilent, Santa Clara, CA). RNA was submitted for sequencing if RNA integrity number was >7. Approximately 500 μg of total RNA was used as input for the TruSeq Stranded mRNA Library Prep kit (Illumina, Inc., San Diego, CA); and library preparation was carried out following manufacturer’s instructions. Libraries were quantified with the Qubit™ dsDNA High Sensitivity Assay Kit (Eurogene, OR); and quality was evaluated using the High Sensitivity DNA chip (Agilent, Santa Clara, CA) on an Agilent 2100 Bioanalyzer. Libraries were sequenced in a HiSeq2500 system at the Genomic Services Laboratory at HudsonAlpha, Huntsville, AL to generate 125nt long, pair-end reads.

Total RNA (circulating extracellular RNA) was isolated from 1ml of plasma by adding 9 ml of TRIzol™ reagent and following manufacturer’s protocols through organic phase separation. The aqueous phase containing the RNA was then collected and purified with the RNA Clean & Concentrator-5 kit (ZymoResearch, Irvine, CA), according to manufacturer’s instructions. Total RNA yield was quantified in test samples using the Qubit™ RNA Broad Range Assay Kit averaging 21.4ng, whereas samples prepared for sequencing were not quantified and all of the eluted RNA was directly used for library preparation.

Total RNA obtained from plasma was desiccated to a volume of 2.5μl for library preparation with the TruSeq Small RNA Library Prep kit (Illumina, Inc., San Diego, CA) following manufacturer’s instructions, with the minor alteration that reagent volumes were halved in all reactions until reverse transcription was performed, as described elsewhere ^81^. After PCR amplification, libraries were loaded onto a 10% polyacrylamide gel ^82^ and electrophoresis was carried out for three hours at 100 volts alongside a 25/100 bp mixed DNA ladder (Bioneer, Alameda, CA); and the custom DNA ladder included in the TruSeq Small RNA Library Prep kit. The gel was soaked in GelRed® Nucleic Acid Gel Stain (Biotium, Inc., Fremont, CA) for DNA staining. DNA bands in the range of 145 -160 base pairs were isolated from the gel by the crush-and-soak method ^83^ with the following modifications. Gel fragments were placed into a 0.5 ml tube containing small holes at the bottom, mounted into a 2ml tube. The tubes were centrifuged at 14,000xg for two minutes at room temperature. The crushed gel was soaked in 200μl of water at 4°C overnight, followed by removal of the solution containing the library to a new tube.

Concentration of each library was determined by Qubit™ dsDNA High Sensitivity Assay Kit, and the volume corresponding to 1.5ng of each library was pooled for further purification with the DNA Clean & Concentrator™-5 (ZymoResearch, Irvine, CA) following the manufacturer’s instructions. The pool was quantified with Qubit™ dsDNA High Sensitivity Assay Kit, and the DNA profile was evaluated using the High Sensitivity DNA chip on an Agilent 2100 Bioanalyzer. Libraries were sequenced in a NextSeq system at the Genomic Services Laboratory at HudsonAlpha, Huntsville, AL to generate 75nt long, single end reads.

Raw sequences and metadata are deposited in the Gene Expression Omnibus database (GSE146041)

### RNA sequencing data processing

Sequences were submitted to a custom build bioinformatics pipeline^84-86^. Reads were aligned to the bovine genome (ARS-UCD1.2 obtained from Ensembl ^87^) using HISAT2 ^88^. Sequences aligning to multiple places on the genome, or with 5 or more mismatches, were filtered out. The sequences were then marked for duplicates, and non-duplicated pairs of reads (mRNA) or single reads (miRNA) were used for transcript quantification at the gene level. The fragments were counted against the Ensembl gene annotation ^89^ (version 1.87) using featureCounts ^90^. RNAseq data from one animal (non-pregnant) were removed from analysis due to low sequencing yield (613,765 reads).

### Filtering of lowly expressed genes

All analytical procedures were carried out in R software ^91^. Supplementary code contains all scripts used for analytical procedures described below and the creation of the plots. This code and all original data can be obtained on figshare ^92^.

Counts per million (CPM), and fragments per kilobase per million reads (FPKM) were calculated using “edgeR” ^93^. In order to remove quantification uncertainty associated to lowly expressed genes and erroneous identification of differentially expressed genes ^94,95^, we retained genes with at least 2 CPM for PWBC mRNA or at least 1 CPM for plasma miRNA in five or more samples for downstream analyses. Further analytical procedures were carried out with genes annotated as “protein-coding” and “miRNA” for the PWBC RNA-sequencing and plasma small RNA-sequencing, respectively.

### Pair-wise gene coexpression and differential coexpression for protein-coding genes

First, we calculated pair-wise Pearson’s correlation coefficient (*r*) ^96,97^ for all 17 animals sampled. Significance of the correlations was assessed using the “TestCor” package based on the empirical statistical test and the large-scale correlation test with normal approximation^98^ for the calculation of the false discovery rate (FDR). Further biological interpretation of results was performed on pairs of genes if abs(r)>0.98, which corresponded to a nominal P≤1.2×10^−16^ and FDR<0.02.

Next, we calculated within group pair-wise Pearson’s correlation coefficient (*r*) for each group: AI-pregnant, NB-pregnant, and non-pregnant. We interpreted a differential correlation as biologically meaningful based on the following scenarios, assuming that groups 1 and 2 can be assigned arbitrarily: inverted correlation, if |*r*_*gruop* 1_ − *r*_*gruop* 2_| > 1.95; gained correlation in group 1 and lost correlation in group 2, if |*r*_*group* 1_| > 0.99 and −0.1 < *r*_*group* 2_ < 0.1.

Significance was estimated by the eFDR approach ^85,99,100^. We resampled (5000 randomizations) the labels of pregnancy outcome and calculated *r* for each pair of genes in the scrambled data set (*r*_*scramble*_). The eFDR was obtained based on the proportion of the *r*_*scramble*_ greater than a specified *r* (i.e.: for r=0.99, eFDR<0.002, Supplementary Fig. S1), relative to the total number of correlations calculated using the formulas described elsewhere ^85,99,100^.

### mRNA-miRNA coexpression and differential coexpression

We calculated Pearson’s correlation (*r*) ^96,97^ values between miRNA and mRNA genes for all 17 samples. Significance was estimated by the eFDR approach ^85,99,100^ by shuffling (5000 randomizations) the animal identification on mRNA dataset, thus breaking the common source of both RNA types (Supplementary Fig. S2).

Pearson’s correlation (r) values were calculated for mRNA-miRNA transcript pairs within groups: AI-pregnant, NB-pregnant, and non-pregnant heifers. We interpreted a differential correlation as biologically meaningful based on the following scenarios |*r*_*group* 1_| > 0.99 and −0.1 < *r*_*group* 2_ < 0.1. Significance was assessed by the eFDR approach (Supplementary Fig. S3). We further interrogated the data for experimentally validated miRNA-mRNA interactions obtained from miRWalk database (version 3)^101,102^.

### Differential expression of mRNA and miRNA

Differences of transcript levels between groups based on the pregnancy outcome were determined from fragment counts using “edgeR” ^93^ and “DESeq2” ^103^. For each comparison, a gene was initially inferred as differentially expressed if the nominal P value was ≤0.03 in both “edgeR” and “DESeq2” analyses. This nominal P value corresponded to eFDR≤0.05 (Supplementary Fig. S4), as calculated according to the procedure and formulas outlined elsewhere ^85,99,100^ using 10000 randomizations of sample reshuffling. Genes inferred as differentially expressed based on P values were further selected through “leave-one-out” cross validation ^104,105^. Only genes whose significance were maintained after all rounds of sampling were further considered for biological interpretation.

### Testing for enrichment of gene ontology terms and KEGG pathways

Differentially expressed genes and genes that displayed differential coexpression among heifers of AI-pregnant, NB-pregnant, and non-pregnant classifications were analyzed for enrichment of Gene Ontology ^106^ categories and KEGG Pathways ^107^. In all tests, the background gene list consisted of all genes detected as expressed in our dataset ^108^. The GO annotations and transcript lengths were obtained from BioMart ^109^ and KEGG annotation was obtained from the “org.Bt.eg.db” package. We used “GoSeq” package ^110^ to estimate significance of category enrichment using the sampling method ^110^. We utilized FDR for multiple testing under dependency ^111^, and inferred statistical significance of GO terms and KEGG pathways if FDR < 0.10.

### Assessment of predictive potential of gene expression levels by machine learning algorithms

We focused the mRNA sequencing data from AI-pregnant and non-pregnant heifers from this report and data we collected from the same site on the previous year and published elsewhere ^38,86^. The raw sequences from the previous dataset were realigned, and data were treated as described above.

To remove the genes not informative to the machine learning algorithm, we identified differentially expressed genes between the two groups (AI-pregnant and non-pregnant). We used the same procedures and standards aforementioned with the exception that we added year as a fixed effect into the model. There were 198 genes differentially expressed between these two groups (AI-preg and non-pregnant). We performed a variance stabilizing transformation on the counts for these 198 genes, and the transformed data were used as input.

For the machine learning approach, we used the expression data for the 198 DEGs from year one (2015) as a training set and the data for the same group of genes from year two (2016) as a test set. Initially we processed the training set for the development of models and assessed the prediction accuracy for 86 classification algorithms (Supplementary Fig. S5). Accuracy was determined by the number of samples correctly classified divided by the number of samples tested (n=11). Six models that showed prediction accuracy ≥0.9 (classified correctly at least 10 samples, accuracy P value ≤0.01, Supplementary Fig. S5), were further subjected to 2000 predictions with different initial ‘seed’ numbers.

## Author contributions

S.E.M. performed sample collection, library preparations, contributed to the data analysis, prepared the first draft and edited the final draft. B.N.W. contributed to sample collection, assisted on the laboratory assays and editing of the final draft. M.F.E. is responsible for the acquisition and curation of the animal database. J.B.E and S.P.R. were responsible for the health and reproductive management of the animals. F.B. conceived designed and supervised the experiments. All authors reviewed the manuscript.

## Competing interests

The author(s) declare no competing interests.

## Data availability

All raw sequencing files and raw read counts produced in this work are deposited in the National Center for Biotechnology Information under the Gene Expression Omnibus database, accession: GSE146041. All the intermediate data and results produced from raw files can be reproduced based on the codes available on Supplementary Code.

## Notes

https://doi.org/10.6084/m9.figshare.11985666.v2

### Competing Interest Statement

The authors have declared no competing interest.

