## Supplementary code for "Rewiring of gene expression in circulating white blood cells is associated with pregnancy outcome in heifers (Bos taurus)"

">

**June 2020**


-
-
- Script to count the features according to Ensembl annotation
  - mRNA
  - small RNA
- Script to create the matrix with the counts for all samples
  - mRNA
  - small RNA
- Data Overview
  - Figures 1b,c
- Differential mRNA:mRNA coexpression associated with pregnancy outcome
  - Supplementary Table 1, data used for Figure 1a
  - Supplementary Table 2
  - Supplementary Fig. 1a,b
  - Figures 1b,c
  - Supplementary Table 4
  - Figure 1d
  - Supplementary Table 5
- Differential miRNA:mRNA coexpression associated with pregnancy outcome
  - Supplementary table 6
  - Supplementary Fig. 2
  - Supplementary Table 7
  - Figure 3a
  - Figure 3b
  - Supplementary Fig. 3
  - Supplementary Table 8
  - Figure 3c
  - Supplementary Table 9
  - Supplementary Table 10
- Differential gene expression associated with pregnancy outcome
  - AI Pregnant vs non pregnant
    - Results from edgeR
    - Results from edgeR both years
    - Results from DESeq
    - Results from DESeq from both years
  - Supplementary Fig. 4
  - Supplementary table 11
- DEG Analysis for mRNA
  - AI Pregnant vs Pregnant to Natural Service
    - Results from edgeR
    - Results from DESeq
- DEG Analysis for mRNA
  - Pregnant to Natural Service vs non pregnant
    - Results from edgeR
    - Results from DESeq
  - Supplementary table 12
  - Figure 4a
  - Figure 4b
  - Figure 4c
- DEG Analysis for miRNA
  - Pregnant to Artificial Insemination vs non pregnant
    - Results from edgeR
    - Results from DESeq
  - AI Pregnant vs Pregnant to Natural Service
    - Results from edgeR
    - Results from DESeq
  - Pregnant to Natural Service vs non pregnant

- Results from edgeR
- Results from DESeq
- Assessment of transcript levels as predictors of pregnancy outcome
  - Supplementary Fig. 5
  - Figure 5d
  - Figure 5e
  - Figure 5f


This file was created to permit reproducibility of the findings described in the paper. Please direct questions to Fernando Biase: ***fbiase*** at ***vt.edu***
The raw data is deposited on GEO repository under the following access GSE146041 (https://www.ncbi.nlm.nih.gov/geo/query/acc.cgi?acc=GSE146041). For the reproduction of this code please download the R binary file containing all the objects corresponding to the data used in this work: resource_data_2019_06_28.RData (https://github.com/Biase-Lab/heifer_transcriptome/blob/master/resources/resource_data_2019_06_28.RData)

```
library("DESeq2", lib.loc="/usr/lib/R/site-library")
library('edgeR', lib.loc="/usr/lib/R/site-library")
library("statmod", lib.loc="/usr/lib/R/site-library")
library("ggplot2", lib.loc="/usr/lib/R/site-library")
library("ggrepel", lib.loc="/usr/lib/R/site-library")
library("biomaRt", lib.loc="/usr/lib/R/site-library")
library("goseq", lib.loc="/usr/lib/R/site-library")
library("WGCNA", lib.loc="/usr/lib/R/site-library")
library ("reshape2", lib.loc="/usr/lib/R/site-library")
library ("bigmemory", lib.loc="/usr/lib/R/site-library")
library("doParallel", lib.loc="/usr/lib/R/site-library")
library("gtools", lib.loc="/usr/lib/R/site-library")
library('org.Bt.eg.db', lib.loc="/usr/lib/R/site-library")
library('DT', lib.loc="/usr/lib/R/site-library")
library("Rtsne", lib.loc="/usr/lib/R/site-library")
library("ggpubr", lib.loc="/usr/lib/R/site-library")
library('grid', lib.loc="/usr/lib/R/site-library")
library('gridExtra', lib.loc="/usr/lib/R/site-library")
library("flashClust", lib.loc="/usr/lib/R/site-library")
library("parallelDist", lib.loc="/usr/lib/R/site-library")
library("ComplexHeatmap", lib.loc="/usr/lib/R/site-library")
library("circlize", lib.loc="/usr/lib/R/site-library")
library("caret", lib.loc="/usr/lib/R/site-library")
library("caretEnsemble", lib.loc="/usr/lib/R/site-library")
library("foreach", lib.loc="/usr/lib/R/site-library")
library("TestCor", lib.loc="/usr/lib/R/site-library")
library("scales", lib.loc="/usr/lib/R/site-library")
library("stringi", lib.loc="/usr/lib/R/site-library")
library("networkD3" , lib.loc="/usr/lib/R/site-library")

#source("multiplot.R") #from Cookbook for R Multiple graphs on one page
#(http://www.cookbook-r.com/Graphs/Multiple_graphs_on_one_page_%28ggplot2%29/)
```

### Script to count the features according to Ensembl annotation

#### mRNA

Code used to produce the counts for mRNA data for each animal. Note that this was executed outside R as a bash command on the terminal.

```
#!/bin/bash
for folder in /data/auburn/heifer_pregnancy/proj_2018/align_genome/SL*; do
#input files
gff_file=/home/fernando/genomes/Bos_taurus.ARS-UCD1.2.95.gtf
alignment=$folder/Aligned.out.merged.filtered.sorted.bam
#output files
sample=$(echo $folder | awk '{n=split($0,a,"/");print a[7]}')
output=/data/auburn/heifer_pregnancy/proj_2018/count_mRNA/$sample.count
/home/fernando/bioinfo/subread-1.6.2-Linux-x86_64/bin/featureCounts -s2 -a $gff_file -o $output -F 'GTF' -t 'exo
n' -g 'gene_id' --ignoreDup -p -T 10  $alignment
done;
```

#### small RNA

Code used to produce the counts for small RNA data for each animal. Note that this was executed outside R as a bash command on the terminal.

```
#!/bin/bash
for folder in /data/auburn/heifer_pregnancy/proj_2018/alignment_small_rna_a/heifer*; do
#input files
gff_file=/home/fernando/genomes/Bos_taurus.ARS-UCD1.2.95.gtf
alignment=$folder/aligned.sorted.bam
#output files
sample=$(echo $folder | awk '{n=split($0,a,"/");print a[7]}')
output=/data/auburn/heifer_pregnancy/proj_2018/count_small_rna_ensembl/$sample.count
/home/fernando/bioinfo/subread-1.6.2-Linux-x86_64/bin/featureCounts -a $gff_file -o $output -F 'GTF' -t 'exon' -g
'gene_id' --ignoreDup  -T 10  $alignment
done;
```

### Script to create the matrix with the counts for all samples

#### mRNA

Code used to produce the matrix containing all the counts for mRNA data.

```
setwd("/data/auburn/heifer_pregnancy/proj_2018/count_mRNA")
files<-list.files(".", recursive=T, pattern=".count" )
files<-files[grep("summary", files, invert = TRUE)]
files<-files[grep("featurecount", files, invert = TRUE)]
#create the count data
count<-data.frame()
count_pwbc<-data.frame(matrix(nrow=27570))
for (n in 1:length(files)) {
  count<-read.delim(files[n], header =TRUE, sep = "\t",  stringsAsFactors = FALSE,comment.char = "#")
  count<-count[,c(1,7)]
  count_pwbc<-cbind(count_pwbc,count)
}

dim(count_pwbc) #27570
rownames(count_pwbc)<-count_pwbc[,2]
count_pwbc<-count_pwbc[,c(3,5,7,9,11,13,15,17,19,21,23,25,27,29,31,33,35,37)]
colnames(count_pwbc)<-substr(colnames(count_pwbc), 55,62)
dim(count_pwbc) #27570
head(count_pwbc)

#write.table(count_pwbc,file="/home/fernando/auburn/heifer_pregnancy/proj_2018/analysis/resources/2019_05_02_unfi
ltered_count_pwbc.txt", sep = "\t",append = FALSE, quote = FALSE, row.names = TRUE)
```

Code used to produce the matrix containing all the counts for mRNA data produced in 2017. Original data can be obtained from GSE103628 (https://www.ncbi.nlm.nih.gov/geo/query/acc.cgi?acc=GSE103628)

```
setwd("/data/auburn/heifer_pregnancy/proj_2018/count_mRNA_proj2017")
files<-list.files(".", recursive=T, pattern=".count" )
files<-files[grep("summary", files, invert = TRUE)]
files<-files[grep("featurecount", files, invert = TRUE)]
#create the count data
count<-data.frame()
count_pwbc_2017<-data.frame(matrix(nrow=27570))
for (n in 1:length(files)) {
  count<-read.delim(files[n], header =TRUE, sep = "\t",  stringsAsFactors = FALSE,comment.char = "#")
  count<-count[,c(1,7)]
  count_pwbc_2017<-cbind(count_pwbc_2017,count)
}

dim(count_pwbc_2017) #27570
rownames(count_pwbc_2017)<-count_pwbc_2017[,2]
count_pwbc_2017<-count_pwbc_2017[,c(3,5,7,9,11,13,15,17,19,21,23,25,27,29,31,33,35,37,39,41,43,45,47,49)]
colnames(count_pwbc_2017)<-substr(colnames(count_pwbc_2017),74,81)
dim(count_pwbc_2017) #27570
head(count_pwbc_2017)

#write.table(count_pwbc_2017,file="/home/fernando/auburn/heifer_pregnancy/proj_2018/analysis/resources/2019_05_02
_unfiltered_count_pwbc_2017.txt", sep = "\t",append = FALSE, quote = FALSE, row.names = TRUE)
```


```
setwd("/data/auburn/heifer_pregnancy/proj_2018/count_mRNA_proj2017")
files<-list.files(".", recursive=T, pattern=".count" )
files<-files[grep("summary", files, invert = TRUE)]
files<-files[grep("featurecount", files, invert = TRUE)]
#create the count data
count<-data.frame()
count_pwbc_2017<-data.frame(matrix(nrow=27570))
for (n in 1:length(files)) {
  count<-read.delim(files[n], header =TRUE, sep = "\t",  stringsAsFactors = FALSE,comment.char = "#")
  count<-count[,c(1,7)]
  count_pwbc_2017<-cbind(count_pwbc_2017,count)
}

dim(count_pwbc_2017) #27570
rownames(count_pwbc_2017)<-count_pwbc_2017[,2]
count_pwbc_2017<-count_pwbc_2017[,c(3,5,7,9,11,13,15,17,19,21,23,25,27,29,31,33,35,37,39,41,43,45,47,49)]
colnames(count_pwbc_2017)<-substr(colnames(count_pwbc_2017),74,81)
dim(count_pwbc_2017) #27570
head(count_pwbc_2017)

#write.table(count_pwbc_2017,file="/home/fernando/auburn/heifer_pregnancy/proj_2018/analysis/resources/2019_05_02
_unfiltered_count_pwbc_2017.txt", sep = "\t",append = FALSE, quote = FALSE, row.names = TRUE)
```


### small RNA

Code used to produce the matrix containing all the counts for miRNA data.

```
setwd("/data/auburn/heifer_pregnancy/proj_2018/count_small_rna_ensembl")
files<-list.files(".", recursive=T, pattern=".count" )
files<-files[grep("summary", files, invert = TRUE)]
files<-files[grep("featurecount", files, invert = TRUE)]
#create the count data
count<-data.frame()
count_miRNA_plasma<-data.frame(matrix(nrow=27570))
for (n in 1:length(files)) {
  count<-read.delim(files[n], header =TRUE, sep = "\t",  stringsAsFactors = FALSE,comment.char = "#")
  count<-count[,c(1,7)]
  count_miRNA_plasma<-cbind(count_miRNA_plasma,count)
}

dim(count_miRNA_plasma) #27570
rownames(count_miRNA_plasma)<-count_miRNA_plasma[,2]
count_miRNA_plasma<-count_miRNA_plasma[,c(3,5,7,9,11,13,15,17,19,21,23,25,27,29,31,33,35,37)]
colnames(count_miRNA_plasma)<-substr(colnames(count_miRNA_plasma), 64,73)
dim(count_miRNA_plasma) #27570
head(count_miRNA_plasma)

#write.table(count_miRNA_plasma,file="/home/fernando/auburn/heifer_pregnancy/proj_2018/analysis/resources/2019_05
_02_unfiltered_count_miRNA_plasma.txt", sep = "\t",append = FALSE, quote = FALSE, row.names = TRUE)
```

Code to generate objects with annotation. Executed and the files saved on February 05 2019.

```
cow<-useMart("ensembl", dataset = "btaurus_gene_ensembl")
annotation.ensembl.symbol<-getBM(attributes = c('ensembl_gene_id', 'external_gene_name','description','hgnc_symbo
l','gene_biotype','transcript_length'),  values = "*", mart = cow)
annotation.ensembl.symbol<-getBM(attributes = c('ensembl_gene_id','external_gene_name','description','hgnc_symbo
l','gene_biotype','transcript_length'),  values = "*", mart = cow)
annotation.ensembl.symbol<-annotation.ensembl.symbol[order(annotation.ensembl.symbol$ensembl_gene_id, -annotatio
n.ensembl.symbol$transcript_length),]
annotation.ensembl.symbol<-annotation.ensembl.symbol[!duplicated(annotation.ensembl.symbol$ensembl_gene_id),]
annotation.GO.biomart<-getBM(attributes = c('ensembl_gene_id', 'external_gene_name','go_id','name_1006','namespac
e_1003'),  values = "*", mart = cow)
```

Set the working directories according to your path structure.

```
rm(list=ls())
sourcefile_path<-"/mnt/storage/auburn/heifer_pregnancy/proj_2018/analysis/resources"
result_path<-"/mnt/storage/auburn/heifer_pregnancy/proj_2018/analysis/results"
```

Create the .RData with all the data needed

```
count_miRNA_plasma<-read.delim(paste(sourcefile_path, "2019_05_02_unfiltered_count_miRNA_plasma.txt.bz2", sep="/"
), stringsAsFactors=FALSE)
count_pwbc<-read.delim(paste(sourcefile_path, "2019_05_02_unfiltered_count_pwbc.txt.bz2", sep="/"), stringsAsFact
ors=FALSE)
count_pwbc_2017<-read.delim(paste(sourcefile_path, "2019_05_02_unfiltered_count_pwbc_2017.txt.bz2", sep="/"), str
ingsAsFactors=FALSE)

gene.length<-read.delim(paste(sourcefile_path, "gene_length.txt.bz2", sep="/"), stringsAsFactors=FALSE)
annotation.ensembl.symbol<-read.delim(paste(sourcefile_path, "2019_02_05_anontation_cow.txt.bz2", sep="/"), strin
gsAsFactors=FALSE)
annotation.GO.biomart<-read.delim(paste(sourcefile_path, "2019_02_05_anontation_GO_cow.txt.bz2", sep="/"), string
sAsFactors=FALSE)

bta_miRWalk_CDS<-read.delim(paste(sourcefile_path, "bta_miRWalk_CDS.txt.bz2", sep="/"), stringsAsFactors=FALSE)
bta_miRWalk_3UTR<-read.delim(paste(sourcefile_path, "bta_miRWalk_3UTR.txt.bz2", sep="/"), stringsAsFactors=FALSE)
bta_miRWalk_5UTR<-read.delim(paste(sourcefile_path, "bta_miRWalk_5UTR.txt.bz2", sep="/"), stringsAsFactors=FALSE)
gc()
#save(count_miRNA_plasma, count_pwbc, count_pwbc_2017, gene.length, annotation.ensembl.symbol, annotation.GO.biom
art,bta_miRWalk_CDS,bta_miRWalk_3UTR,bta_miRWalk_5UTR, file=paste(sourcefile_path, "resource_data_2019_06_28.RDat
a", sep="/"), compress="bzip2")
```

Load the files needed for analyses.

```
load( file=paste(sourcefile_path,"resource_data_2019_06_28.RData", sep="/"), verbose =TRUE)
```

```
## Loading objects:
##   count_miRNA_plasma
##   count_pwbc
##   count_pwbc_2017
##   gene.length
##   annotation.ensembl.symbol
##   annotation.GO.biomart
##   bta_miRWalk_CDS
##   bta_miRWalk_3UTR
##   bta_miRWalk_5UTR
```

```
source(paste(sourcefile_path,"multiplot.R", sep="/"))
```

Filtering of the mRNA dataset.

```
#Filter to remove the library with extremely low read counts
count_pwbc<-count_pwbc[,-3]
#Filter to remove lowly expressed genes
keep<-rowSums( cpm(count_pwbc) >= 2 ) >= 5
count_pwbc<-count_pwbc[keep,]

#Filter to retain only protein coding genes
count_pwbc<-merge(count_pwbc,annotation.ensembl.symbol, by.x="row.names", by.y="ensembl_gene_id", all=FALSE)
count_pwbc<-count_pwbc[ which(count_pwbc$gene_biotype=='protein_coding'), ]
rownames(count_pwbc)<-count_pwbc$Row.names
count_pwbc<-count_pwbc[,c(10,4,16,18,8,5,9,15,11,2,6,12,3,13,14,7,17)]
colnames(count_pwbc) <- c("AI_92", "AI_99", "AI_114", "AI_117", "AI_119", "AI_120", "NB_93", "NB_103", "NB_108",
"NB_111", "NB_116", "NB_121", "NP_94", "NP_101", "NP_107", "NP_113", "NP_115")
```

```
#Filter 2017 data to remove lowly expressed genes
count_pwbc_2017<-count_pwbc_2017[,c(13,14,15,16,17,18,19,20,21,22,23,24)]
keep<-rowSums( cpm(count_pwbc_2017) >= 2 ) >= 6
count_pwbc_2017<-count_pwbc_2017[keep,]
dim(count_pwbc_2017) #10602
```

```
## [1] 10602    12
```

```
#Filter 2017 data to retain only protein coding genes
count_pwbc_2017<-merge(count_pwbc_2017,annotation.ensembl.symbol, by.x="row.names", by.y="ensembl_gene_id", all=F
ALSE)
count_pwbc_2017<-count_pwbc_2017[ which(count_pwbc_2017$gene_biotype=='protein_coding'), ]
rownames(count_pwbc_2017)<-count_pwbc_2017[,1]
count_pwbc_2017<-count_pwbc_2017[,c(2:13)]
colnames(count_pwbc_2017) <- c("AI_63", "NP_59", "NP_61", "NP_53", "AI_62", "AI_49", "AI_42", "AI_55", "NP_39",
"AI_36","NP_64", "NP_58")
```

```
merged_datasets<-merge(count_pwbc,count_pwbc_2017, by="row.names", all=FALSE)
rownames(merged_datasets)<-merged_datasets[,1]
merged_datasets<-merged_datasets[,2:30]
```

Calculate counts per million reads and fragments per kilobase per million reads.

```
gene_length<-subset(gene.length, Geneid %in% rownames(count_pwbc))
gene_length<-gene_length[order(gene_length$Geneid),]
count_pwbc<-count_pwbc[order(rownames(count_pwbc)),]
table(rownames(count_pwbc)==gene_length$Geneid) #all True
```

```
##
##  TRUE
## 10496
```

```
cpm_pwbc<-cpm(count_pwbc,normalized.lib.sizes = TRUE,log = FALSE)
rpkm_pwbc<-rpkm(count_pwbc, gene.length = gene_length$Length, normalized.lib.sizes = TRUE,log = FALSE)

rpkm_AIpreg<-rpkm_pwbc[,c(1,2,3,4,5,6)]
#dim(rpkm_AIpreg) #10496 X 6
#head(rpkm_AIpreg)
rpkm_NBpreg<-rpkm_pwbc[,c(7,8,9,10,11,12)]
#dim(rpkm_NBpreg) #10496 X 6
#head(rpkm_NBpreg)
rpkm_NOTpreg<-rpkm_pwbc[,c(13,14,15,16,17)]
#dim(rpkm_NOTpreg) #10496 X 5
#head(rpkm_NOTpreg)

cpm_AIpreg<-cpm_pwbc[,c(1,2,3,4,5,6)]
cpm_NBpreg<-cpm_pwbc[,c(7,8,9,10,11,12)]
cpm_NOTpreg<-cpm_pwbc[,c(13,14,15,16,17)]
```

```
#for merged dataset
gene_length_merged<-subset(gene.length, Geneid %in% rownames(merged_datasets))
gene_length_merged<-gene_length_merged[order(gene_length_merged$Geneid),]
merged_datasets<-merged_datasets[order(rownames(merged_datasets)),]
table(rownames(merged_datasets)==gene_length_merged$Geneid) #all True
```

```
##
## TRUE
## 9960
```

```
cpm_merged_datasets<-cpm(merged_datasets,normalized.lib.sizes = TRUE,log = FALSE)
rpkm_merged_datasets<-rpkm(merged_datasets, gene.length = gene_length_merged$Length, normalized.lib.sizes = TRUE,
log = FALSE)

rpkm_AIpreg_merged<-rpkm_merged_datasets[,c(1,2,3,4,5,6,18,22,23,24,25,27)]
#dim(rpkm_AIpreg_merged) #9960 X 6
#head(rpkm_AIpreg_merged)

rpkm_NOTpreg_merged<-rpkm_merged_datasets[,c(13,14,15,16,17,19,20,21,26,28,29)]
#dim(rpkm_NOTpreg_merged) #9960 X 5
#head(rpkm_NOTpreg_merged)
```

Filtering of the miRNA dataset.

```
#Filter to remove the library with extremely low read counts
count_miRNA_plasma<-count_miRNA_plasma[,-17]
#Filter to remove data with low read counts
keep<-rowSums( cpm(count_miRNA_plasma) > 1 ) >= 5
count_miRNA_plasma<-count_miRNA_plasma[keep,]
dim(count_miRNA_plasma) #1086
```

```
## [1] 1086   17
```

```
#Fiter to retain only microRNAs
count_miRNA_plasma<-merge(count_miRNA_plasma,annotation.ensembl.symbol, by.x="row.names", by.y="ensembl_gene_id",
all=FALSE)
count_miRNA_plasma<-count_miRNA_plasma[ which(count_miRNA_plasma$gene_biotype=='miRNA'), ]
rownames(count_miRNA_plasma)<-count_miRNA_plasma$Row.names
count_miRNA_plasma<-count_miRNA_plasma[,c(15,18,8,11,12,13,16,3,5,6,10,14,17,2,4,7,9)]
names(count_miRNA_plasma) <- c("AI_92", "AI_99", "AI_114", "AI_117", "AI_119", "AI_120", "NB_93", "NB_103", "NB_1
08", "NB_111", "NB_116", "NB_121", "NP_94", "NP_101", "NP_107", "NP_113", "NP_115")
```

Calculate counts per million reads and fragments per kilobase per million reads.

```
gene_length_miRNA<-subset(gene.length, Geneid %in% rownames(count_miRNA_plasma))
gene_length_miRNA<-gene_length_miRNA[order(gene_length_miRNA$Geneid),]
count_miRNA_plasma<-count_miRNA_plasma[order(rownames(count_miRNA_plasma)),]
table(rownames(count_miRNA_plasma)==gene_length_miRNA$Geneid) #all TRUE
```

```
##
## TRUE
##  290
```

```
cpm_miRNA<-cpm(count_miRNA_plasma,normalized.lib.sizes = TRUE,log = FALSE)
rpkm_miRNA<-rpkm(count_miRNA_plasma, gene.length = gene_length_miRNA$Length, normalized.lib.sizes = TRUE,log = FA
LSE)

#write.table(cpm_miRNA,file="/mnt/storage/auburn/heifer_pregnancy/proj_2018/analysis/results/2020_02_17_filtered_
count_miRNA_plasma.txt", sep = "\t",append = FALSE, quote = FALSE, row.names = TRUE)

rpkm_AIpreg_miRNA<-rpkm_miRNA[,c(1,2,3,4,5,6)]
rpkm_NBpreg_miRNA<-rpkm_miRNA[,c(7,8,9,10,11,12)]
rpkm_NOTpreg_miRNA<-rpkm_miRNA[,c(13,14,15,16,17)]

cpm_AIpreg_miRNA<-cpm_miRNA[,c(1,2,3,4,5,6)]
cpm_NBpreg_miRNA<-cpm_miRNA[,c(7,8,9,10,11,12)]
cpm_NOTpreg_miRNA<-cpm_miRNA[,c(13,14,15,16,17)]
```

### Data Overview

### Figures 1b,c

```
data.for.tsne.mRNA<-t(log2(rpkm_pwbc+1))
data.for.tsne.miRNA<-t(log2(rpkm_miRNA+1))
set.seed(9434)
tsne_mRNA <- Rtsne(data.for.tsne.mRNA, dims = 2, theta = 0, perplexity=2, verbose=FALSE, max_iter = 500000)
tsne_miRNA <- Rtsne(data.for.tsne.miRNA, dims = 2, theta = 0, perplexity=2, verbose=FALSE, max_iter = 500000)
```

```
group<-factor(c("Preg_AI","Preg_AI","Preg_AI","Preg_AI","Preg_AI","Preg_AI","Preg_NB","Preg_NB","Preg_NB","Preg_N
B","Preg_NB","Preg_NB","Not_Preg","Not_Preg","Not_Preg","Not_Preg","Not_Preg"), levels=c("Preg_AI","Preg_NB", "No
t_Preg"))

TSNE_mRNA<-data.frame("DIM1"=tsne_mRNA$Y[,1], "DIM2"=tsne_mRNA$Y[,2], "group"=group )
TSNE_miRNA<-data.frame("DIM1"=tsne_miRNA$Y[,1], "DIM2"=tsne_miRNA$Y[,2], "group"=group )

plot1<-ggplot(data=TSNE_mRNA, aes(x=DIM1,y=DIM2) ) +
  geom_point(aes(colour = factor(group)), size=3 , alpha =0.8 ) +
  scale_color_manual(values= c("#0072B2", "#000000","#D55E00"), labels=c("AI-preg","NB-preg", "non preg") ,name
 =NULL) +
  scale_y_continuous(name="t-SNE dim 2")+
  scale_x_continuous(name="t-SNE dim 1")+
  ggtitle("10496 protein coding genes")+
  theme_bw(base_size = 11)+
  theme(aspect.ratio =1,
        axis.text=element_blank(),
        legend.position = "none",
        plot.title = element_text(lineheight=.8, hjust=0.5, vjust=-1,size= 11),
        plot.margin = unit(c(0,0.2,0,0),"cm")
  )

plot2<-ggplot(data=TSNE_miRNA, aes(x=DIM1,y=DIM2) ) +
  geom_point(aes(colour = factor(group)), size=3 , alpha =0.8 ) +
  scale_color_manual(values= c("#0072B2", "#000000","#D55E00"), labels=c("AI-preg","NB-preg", "not-preg") ,name
 =NULL) +
  scale_y_continuous(name="t-SNE dim 2")+
  scale_x_continuous(name="t-SNE dim 1")+
  ggtitle("290 miRNAs")+
  theme_bw(base_size = 11)+
  theme(aspect.ratio =1,
        axis.text=element_blank(),
        legend.position = "none",
        plot.title = element_text(lineheight=.8, hjust=0.5,vjust=-1, size= 11),
        plot.margin = unit(c(0,0,0,0.2),"cm")
  )
leg <- get_legend(
  ggplot(data=TSNE_miRNA, aes(x=DIM1,y=DIM2) ) +
  geom_point(aes(colour = factor(group)), size=3 , alpha =0.8 ) +
  scale_color_manual(values= c("#0072B2", "#000000","#D55E00"), labels=c("AI-preg","NB-preg", "non preg") ,name
 =NULL) +
  theme_bw(base_size = 12)+
  theme(legend.position = "top",legend.direction="horizontal"))


#font_size 12 for figure
#pdf(paste(result_path,"/2019_06_29_tsne_plots.pdf", sep=""), width=4.5, height=3.2)
#grid.arrange(plot1,plot2, as_ggplot(leg), layout_matrix=rbind(c(1:2),3), heights=unit(c(3,2), c("in", "mm")))
#dev.off()
grid.arrange(plot1,plot2, as_ggplot(leg), layout_matrix=rbind(c(1:2),3), heights=unit(c(3,2), c("in", "mm")))
```

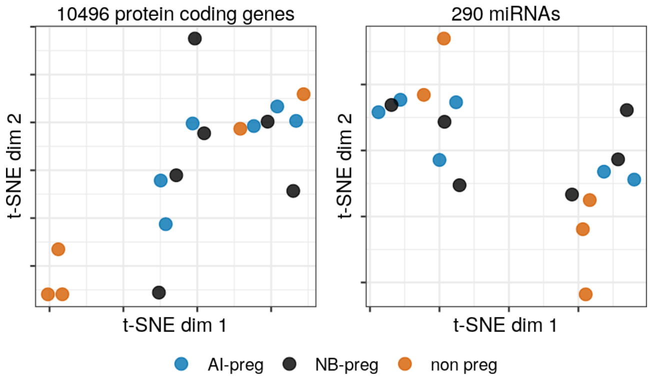


```
rm(plot1,plot2, leg, TSNE_miRNA, TSNE_mRNA, data.for.tsne.mRNA, data.for.tsne.miRNA, tsne_mRNA, tsne_miRNA)
```

### Differential mRNA:mRNA coexpression associated with pregnancy outcome

#### Supplementary Table 1, data used for Figure 1a

Create correlation matrices for all animals.

```
correlation_all_animals<-corAndPvalue(t(log2(rpkm_pwbc+1)),method="pearson", use="pairwise.complete.obs")
correlation_all_animals_a<-melt(correlation_all_animals$cor)
correlation_all_animals_a$pvalue<-melt(correlation_all_animals$p)$value
rm(correlation_all_animals)
correlation_all_animals_a<-correlation_all_animals_a[!(correlation_all_animals_a$Var1 == correlation_all_animals_
a$Var2),]
correlation_all_animals_a<-correlation_all_animals_a[!duplicated(t(apply(correlation_all_animals_a[,c(1:2)], 1, s
ort))),]

correlation_sig_fdr0.02<-ApplyFdrCor(t(log2(rpkm_pwbc+1)), alpha=0.02, stat_test="empirical", method="LCTnorm", v
ect=TRUE)

correlation_all_animals_a$eFDR0.02<-correlation_sig_fdr0.02

correlation_all_animals_b<-correlation_all_animals_a[abs(correlation_all_animals_a$value) > 0.98,]

correlation_all_animals_b$gene_1<-annotation.ensembl.symbol$external_gene_name[match( correlation_all_animals_b$V
ar1, annotation.ensembl.symbol$ensembl_gene_id )]
correlation_all_animals_b$gene_2<-annotation.ensembl.symbol$external_gene_name[match( correlation_all_animals_b$V
ar2, annotation.ensembl.symbol$ensembl_gene_id )]
correlation_all_animals_b$Var1<-as.character(correlation_all_animals_b$Var1)
correlation_all_animals_b$Var2<-as.character(correlation_all_animals_b$Var2)

correlation_all_animals_b$gene_1<-ifelse(correlation_all_animals_b$gene_1=="", correlation_all_animals_b$Var1,cor
relation_all_animals_b$gene_1)
correlation_all_animals_b$gene_2<-ifelse(correlation_all_animals_b$gene_2=="", correlation_all_animals_b$Var2,cor
relation_all_animals_b$gene_2)
#write.table(correlation_all_animals_b, file= paste(result_path,"/2020_01_16_gene_coexpression_all_animals_b.tx
t", sep=""), append = FALSE, quote = FALSE, sep = "\t" ,row.names = FALSE)
knitr::kable(head(correlation_all_animals_b), caption = "First five rows of Supplementary table 1",format = 'pand
oc')
```

First five rows of Supplementary table 1

| | Var1 | Var2 | value | pvalue | eFDR0.02 | gene_1 | gene_2 |
|---|---|---|---|---|---|---|---|
| 149546 | ENSBTAG00000006914 | ENSBTAG00000000032 | 0.9847809 | 0 | TRUE | PRR3 | ABI3 |
| 151225 | ENSBTAG00000011444 | ENSBTAG00000000032 | 0.9808916 | 0 | TRUE | HINT2 | ABI3 |
| 236218 | ENSBTAG00000014262 | ENSBTAG00000000064 | 0.9861767 | 0 | TRUE | BZW2 | FEN1 |
| 276256 | ENSBTAG00000008954 | ENSBTAG00000000077 | 0.9835958 | 0 | TRUE | PSMB9 | ADSL |
| 277968 | ENSBTAG00000013623 | ENSBTAG00000000077 | 0.9842698 | 0 | TRUE | TIMM13 | ADSL |
| 307043 | ENSBTAG00000007084 | ENSBTAG00000000080 | 0.9821463 | 0 | TRUE | MAP3K14 | GRWD1 |

#### Supplementary Table 2

Test biological processes for enrichment with genes that are highly correlated in PWBCs in all heifers.

```
all_genes<-data.frame( gene=rownames(count_pwbc), stringsAsFactors=FALSE )
rownames(all_genes)<-all_genes$gene

N_expressed_genes<-length(all_genes$gene)

annotation.genelength.biomart_vector<-gene_length[,2]
names(annotation.genelength.biomart_vector)<-gene_length[,1]

annotation.GO.BP.biomart<-annotation.GO.biomart[annotation.GO.biomart$namespace_1003=="biological_process", c(1,3
)]
annotation.GO.BP.biomart<-annotation.GO.BP.biomart[annotation.GO.BP.biomart$ensembl_gene_id %in% rownames(all_gen
es),]
annotation.GO.MF.biomart<-annotation.GO.biomart[annotation.GO.biomart$namespace_1003=="molecular_function", c(1,3
)]
annotation.GO.MF.biomart<-annotation.GO.MF.biomart[annotation.GO.MF.biomart$ensembl_gene_id %in% rownames(all_gen
es),]
```

```
test.genes<-data.frame(a=unique(c(as.character(correlation_all_animals_b$Var1),as.character(correlation_all_anima
ls_b$Var2))), stringsAsFactors=FALSE)
all_genes_numeric<-as.integer(all_genes$gene %in%test.genes$a)
names(all_genes_numeric)<-all_genes$gene

N_DEGs<-length(test.genes$a)

set.seed(9830)
pwf<-nullp(all_genes_numeric, bias.data=annotation.genelength.biomart_vector, plot.fit=FALSE )
GO_BP_Cats_mRNA_all_heifers<-goseq(pwf,gene2cat=annotation.GO.BP.biomart, method ="Sampling", repcnt = 7000, use_
genes_without_cat=FALSE)
GO_BP_Cats_mRNA_all_heifers<-GO_BP_Cats_mRNA_all_heifers[GO_BP_Cats_mRNA_all_heifers$numDEInCat>4,]
GO_BP_Cats_mRNA_all_heifers$BY_FDR<-p.adjust(GO_BP_Cats_mRNA_all_heifers$over_represented_pvalue, method ="fdr")
GO_BP_Cats_mRNA_all_heifers<-GO_BP_Cats_mRNA_all_heifers[with(GO_BP_Cats_mRNA_all_heifers, order(BY_FDR,over_repr
esented_pvalue, -numDEInCat)), ]
#head(GO_BP_Cats_mRNA_all_heifers, n=20)

GO_BP_Cats_mRNA_all_heifers$fold_enrichment<-(GO_BP_Cats_mRNA_all_heifers$numDEInCat/N_DEGs)/(GO_BP_Cats_mRNA_all
_heifers$numInCat/N_expressed_genes)
annotation.GO.BP.biomart_testgenes<-annotation.GO.BP.biomart[annotation.GO.BP.biomart$ensembl_gene_id %in% test.g
enes$a, ]
GO_BP_Cats_mRNA_all_heifers<-merge(GO_BP_Cats_mRNA_all_heifers,annotation.GO.BP.biomart_testgenes, by.x="categor
y", by.y="go_id", all.x=TRUE, all.y=FALSE)
GO_BP_Cats_mRNA_all_heifers<-merge(GO_BP_Cats_mRNA_all_heifers, annotation.ensembl.symbol, by.x="ensembl_gene_id"
, by.y="ensembl_gene_id", all=FALSE, all.x=TRUE, all.y=FALSE)
GO_BP_Cats_mRNA_all_heifers<-GO_BP_Cats_mRNA_all_heifers[with(GO_BP_Cats_mRNA_all_heifers, order(BY_FDR,term)), ]
GO_BP_Cats_mRNA_all_heifers<-GO_BP_Cats_mRNA_all_heifers[GO_BP_Cats_mRNA_all_heifers$BY_FDR<0.2,]

#write.table(GO_BP_Cats_mRNA_all_heifers, file= paste(result_path,"2019_11_10_GO_BP_Cats_mRNA_cor_above98_all_hei
fers.txt", sep="/"), append = FALSE, quote = FALSE, sep = "\t" ,row.names = FALSE)
rm( correlation_all_animals_a)
```

```
knitr::kable(head(GO_BP_Cats_mRNA_all_heifers), caption = "First five rows of Supplementary Table 2")
```

First five rows of Supplementary Table 2

| | ensembl_gene_id | category | over_represented_pvalue | under_represented_pvalue | numDEInCat | numInCat | term |
|---|---|---|---|---|---|---|---|
| 36 | ENSBTAG00000000359 | GO:0002183 (GO:0002183) | 0.0001428 | 1 | 8 | 13 | cytoplasm translatio initiation |
| 170 | ENSBTAG00000001988 | GO:0002183 (GO:0002183) | 0.0001428 | 1 | 8 | 13 | cytoplasm translatio initiation |
| 212 | ENSBTAG00000002734 | GO:0002183 (GO:0002183) | 0.0001428 | 1 | 8 | 13 | cytoplasm translatio initiation |
| 369 | ENSBTAG00000004861 | GO:0002183 (GO:0002183) | 0.0001428 | 1 | 8 | 13 | cytoplasm translatio initiation |

| | ensembl_gene_id | category | over_represented_pvalue | under_represented_pvalue | numDEInCat | numInCat | term |
|---|---|---|---|---|---|---|---|
| 553 | ENSBTAG00000006543 | GO:0002183 (GO:0002183) | 0.0001428 | 1 | 8 | 13 | cytoplasm translation initiation |
| 885 | ENSBTAG00000010790 | GO:0002183 (GO:0002183) | 0.0001428 | 1 | 8 | 13 | cytoplasm translation initiation |

Create correlation matrices for AI pregnant, NB pregnant and non pregnant animals.

```
correlation_AIpreg<-corAndPvalue(t(log2(rpkm_AIpreg+1)),method="pearson", use="pairwise.complete.obs")
correlation_AIpreg_a<-melt(correlation_AIpreg$cor)
correlation_AIpreg_a$pvalue<-melt(correlation_AIpreg$p)$value
rm(correlation_AIpreg)

correlation_preg_NB<-corAndPvalue(t(log2(rpkm_NBpreg+1)),method="pearson", use="pairwise.complete.obs")
correlation_preg_NB_a<-melt(correlation_preg_NB$cor)
correlation_preg_NB_a$pvalue<-melt(correlation_preg_NB$p)$value
rm(correlation_preg_NB)

correlation_NOTpreg<-corAndPvalue(t(log2(rpkm_NOTpreg+1)),method="pearson", use="pairwise.complete.obs")
correlation_NOTpreg_a<-melt(correlation_NOTpreg$cor)
correlation_NOTpreg_a$pvalue<-melt(correlation_NOTpreg$p)$value
rm(correlation_NOTpreg)
```

Merge the correlation matrices

```
pearson_correlation<-cbind(correlation_AIpreg_a, correlation_preg_NB_a, correlation_NOTpreg_a)
pearson_correlation<-pearson_correlation[,c(1:4,7,8,11,12)]
colnames(pearson_correlation)<-c('gene_1', 'gene_2', 'cor_AI', 'pvalue_AI', 'cor_NB', 'pvalue_NB','cor_NP', 'pval
ue_NP')
#head(pearson_correlation)

pearson_correlation<-pearson_correlation[!(pearson_correlation$gene_1 == pearson_correlation$gene_2),]
dim(pearson_correlation) # 110155520
```

```
## [1] 110155520         8
```

```
pearson_correlation<-pearson_correlation[!duplicated(t(apply(pearson_correlation[,c(1:2)], 1, sort))),]
dim(pearson_correlation) # 55077760
```

```
## [1] 55077760        8
```

### Supplementary Fig. 1a,b

Calculate the empirical false discovery rate for co-expression and differential coexpression.

```
permutation_matrix<-matrix(,nrow=200100,ncol=17)
#head(permutation_matrix)
#dim(permutation_matrix)

for (i in seq(1:200100)){
  sampling <- sample(1:17,17, replace = FALSE)
  if ( ! identical(sampling , c(1:17))){
    permutation_matrix[i,]<-sample(1:17,17, replace = FALSE)
  }}

permutation_matrix<-permutation_matrix[!duplicated(permutation_matrix),]
dim(permutation_matrix)
head(permutation_matrix)

permutation_matrix<-permutation_matrix[sample(1:dim(permutation_matrix)[1], 5000),]
rand<-dim(permutation_matrix)[1]

#diff.coexp<-seq(0.005, 0.05, 0.005)
diff.coexp<-seq(1.6, 2, 0.05)

results <- filebacked.big.matrix(length(diff.coexp),rand, type="double", init=0, separated=FALSE,
                                 backingfile="incidence_matrix.bin",
                                 descriptor="incidence_matrix.desc")
mdesc_result<- describe(results)

cl <- makeCluster(10)
registerDoParallel(cl)

results[,]<-foreach(i = diff.coexp, .combine='rbind',.packages=c("reshape2","bigmemory"), .inorder=TRUE, .verbose
=TRUE ) %:%
            foreach(j = 1:rand, .combine='cbind',.packages=c("reshape2","bigmemory"), .inorder=FALSE, .verbose=TR
UE ) %dopar% {

   rpkm_pwbc_scramble<-log2(rpkm_pwbc[,permutation_matrix[j,]] +1)
   cor_1<-(cor(t(rpkm_pwbc_scramble[,1:6])))
   cor_2<-(cor(t(rpkm_pwbc_scramble[,7:11])))
   length(which(abs(cor_1-cor_2) > i))/2
  }
stopCluster(cl)

total.rand <-  5000 * 55077760

qvalue<-data.frame(diff_correlation = diff.coexp,
                   e.pvalue= (rowSums(results[,]+1))/(total.rand+1),
                   e.pvalue.round= round((rowSums(results[,]+1))/(total.rand+1) ,6))

write.table(qvalue, file= paste(result_path,"/2019_06_26_gene_coexpression_inverted_empiricalFDR.txt", sep=""), a
ppend = FALSE, quote = FALSE, sep = "\t" ,row.names = FALSE)

rm(results)
system("rm incidence_matrix.bin")
system("rm incidence_matrix.desc")

coexp<-seq(0.9, 1, 0.01)

results <- filebacked.big.matrix(length(coexp),rand, type="double", init=0, separated=FALSE,
                                 backingfile="incidence_matrix.bin",
                                 descriptor="incidence_matrix.desc")
mdesc_result<- describe(results)

cl <- makeCluster(10)
registerDoParallel(cl)

results[,]<-foreach(i = coexp, .combine='rbind',.packages=c("reshape2","bigmemory"), .inorder=TRUE, .verbose=TRUE
) %:%
            foreach(j = 1:rand, .combine='cbind',.packages=c("reshape2","bigmemory"), .inorder=FALSE, .verbose=TR
UE ) %dopar% {

   rpkm_pwbc_scramble<-log2(rpkm_pwbc[,permutation_matrix[j,]] +1)
   cor_1<-(cor(t(rpkm_pwbc_scramble[,1:6])))
   length(which(abs(cor_1) > i))/2
  }
stopCluster(cl)

total.rand <-  5000 * 55077760

qvalue<-data.frame(correlation = coexp,
                   e.pvalue= (rowSums(results[,]+1))/(total.rand+1),
                   e.pvalue.round= round((rowSums(results[,]+1))/(total.rand+1) ,6))

#write.table(qvalue, file= paste(result_path,"/2019_06_27_gene_coexpression_empiricalFDR.txt", sep=""), append =
```

```
 FALSE, quote = FALSE, sep = "\t" ,row.names = FALSE)

rm(results)
system("rm incidence_matrix.bin")
system("rm incidence_matrix.desc")
rm(permutation_matrix)
```

```
eFDR_coexpression<-read.delim(paste(result_path,"/2019_06_27_gene_coexpression_empiricalFDR.txt", sep=""), row.na
mes=NULL,header =TRUE, stringsAsFactors =FALSE)
eFDR_coexpression_inverted<-read.delim(paste(result_path,"/2019_06_26_gene_coexpression_inverted_empiricalFDR.tx
t", sep=""), row.names=NULL,header =TRUE, stringsAsFactors =FALSE)
```

```
font_size<-9

plot1<-ggplot()+
  geom_point(data=eFDR_coexpression, aes(x=correlation , y=e.pvalue.round),color="black", size=3, shape=16)+
  geom_line(data=eFDR_coexpression, aes(x=correlation , y=e.pvalue.round),color="black", size=0.1,linetype=3)+
  scale_y_continuous(name="empirical FDR", limits = c(0, 0.1), breaks=seq(0,0.1, 0.005))+
  scale_x_continuous(name="correlation (r)", limits = c(0.9, 1), breaks=seq(0.9,1, 0.01))+
  ggtitle("Gene coexpression within a group of heifers \n of same reproductive outcome")+
  theme_bw()+
  theme(panel.grid= element_blank(),
        panel.background = element_blank(),
        panel.grid.minor = element_blank(),
        panel.grid.major = element_line(color="lightgray"),
        plot.background = element_blank(),
        axis.title=element_text(color="black", size=font_size),
        axis.text=element_text(color="black", size=font_size),
        panel.spacing = unit(c(0.4,0.4,0.4,0.4),"cm"),
        plot.margin = unit(c(0.5,0.5,0.5,0.5),"cm"),
        legend.position="none",
        plot.title = element_text(lineheight=.8, hjust=0.5, size= font_size))

plot2<-ggplot()+
  geom_point(data=eFDR_coexpression_inverted, aes(x=diff_correlation , y=e.pvalue.round),color="black", size=3, s
hape=16)+
  geom_line(data=eFDR_coexpression_inverted, aes(x=diff_correlation , y=e.pvalue.round),color="black", size=0.1,l
inetype=3)+
  scale_y_continuous(name="empirical FDR", limits = c(0, 0.01), breaks=seq(0,0.01, 0.001))+
  scale_x_continuous(name="absolute difference between correlation values", limits = c(1.5, 2), breaks=seq(1.5, 2
, 0.05))+
  ggtitle("Inverted gene coexpression \n between groups of heifers")+
  theme_bw()+
  theme(panel.grid= element_blank(),
        panel.background = element_blank(),
        panel.grid.minor = element_blank(),
        panel.grid.major = element_line(color="lightgray"),
        plot.background = element_blank(),
        axis.title=element_text(color="black", size=font_size),
        axis.text=element_text(color="black", size=font_size),
        panel.spacing = unit(c(0.4,0.4,0.4,0.4),"cm"),
        plot.margin = unit(c(0.5,0.5,0.5,0.5),"cm"),
        legend.position="none",
        plot.title = element_text(lineheight=.8, hjust=0.5 , size= font_size ))

#font_size 12 for figure
#pdf(paste(result_path,"2019_11_10_eFDR_plots.pdf", sep="/"), width=10, height=5)
#grid.arrange(plot1,plot2, ncol = 2)
#dev.off()

grid.arrange(plot1,plot2, ncol = 2)
```

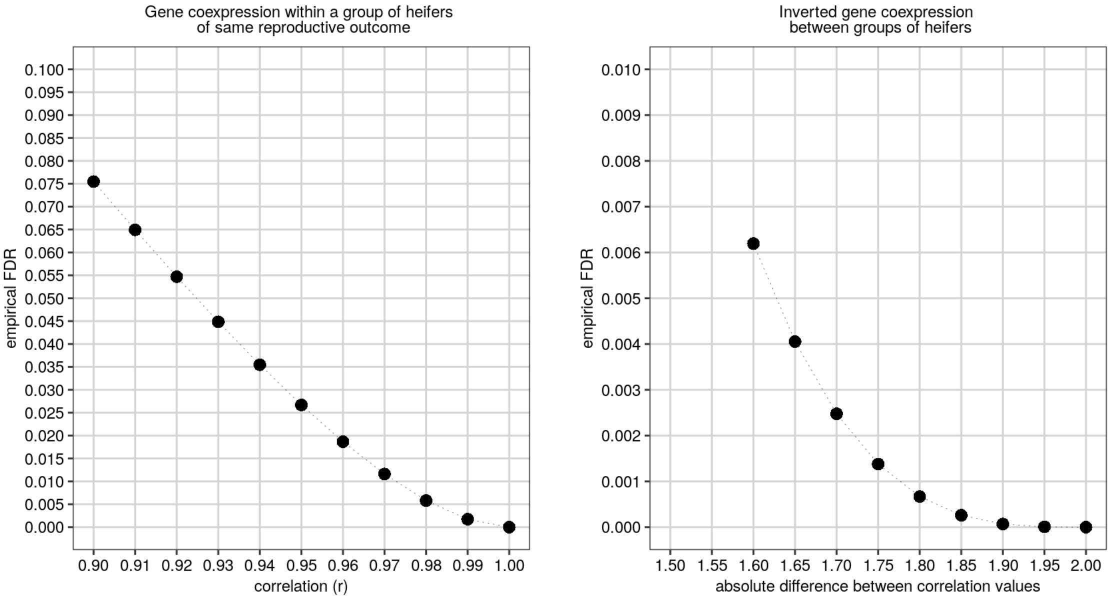


Perform calculations to determine differential coexpression.

```
pearson_correlation$subtraction_cor_AI_cor_NP<-pearson_correlation$cor_AI - pearson_correlation$cor_NP
pearson_correlation$subtraction_cor_NB_cor_NP<-pearson_correlation$cor_NB - pearson_correlation$cor_NP
pearson_correlation$subtraction_cor_AI_cor_NB<-pearson_correlation$cor_AI - pearson_correlation$cor_NB


pearson_correlation$result_cor_AI_cor_NP<-ifelse(abs(pearson_correlation$subtraction_cor_AI_cor_NP) > 1.95 , "inv
erted",
                                   ifelse(abs(pearson_correlation$cor_AI) >0.99 & (pearson_correlation$cor_NP > -
0.1 & pearson_correlation$cor_NP < 0.1) , "loss",
                                          ifelse( (pearson_correlation$cor_AI > -0.1 & pearson_correlation$cor_AI
< 0.1)  &  abs(pearson_correlation$cor_NP) >0.99, "gain" , "nothing")))

pearson_correlation$result_cor_NB_cor_NP<-ifelse(abs(pearson_correlation$subtraction_cor_NB_cor_NP) > 1.95 , "inv
erted",
                                   ifelse(abs(pearson_correlation$cor_NB) >0.99 & (pearson_correlation$cor_NP > -
0.1 & pearson_correlation$cor_NP < 0.1) , "loss",
                                          ifelse( (pearson_correlation$cor_NB > -0.1 & pearson_correlation$cor_NB
< 0.1)  &  abs(pearson_correlation$cor_NP) >0.99, "gain" , "nothing")))

pearson_correlation$result_cor_AI_cor_NB<-ifelse(abs(pearson_correlation$subtraction_cor_AI_cor_NB) > 1.95 , "inv
erted",
                                   ifelse(abs(pearson_correlation$cor_AI) >0.99 & (pearson_correlation$cor_NB > -
0.1 & pearson_correlation$cor_NB < 0.1) , "loss",
                                          ifelse( (pearson_correlation$cor_AI > -0.1 & pearson_correlation$cor_AI
< 0.1)  &  abs(pearson_correlation$cor_NB) >0.99, "gain" , "nothing")))


pearson_correlation$result_cor_AI_and_NB_cor_NP<-ifelse( (abs(pearson_correlation$subtraction_cor_AI_cor_NP) + ab
s(pearson_correlation$subtraction_cor_NB_cor_NP)) / 2 > 1.9 , "inverted",
                                   ifelse( (abs(pearson_correlation$cor_AI) >0.99  & abs(pearson_correlation$cor_
NB) >0.99)  & (pearson_correlation$cor_NP > -0.1 & pearson_correlation$cor_NP < 0.1) , "loss",
                                          ifelse( (pearson_correlation$cor_AI > -0.1 & pearson_correlation$cor_AI
< 0.1 & pearson_correlation$cor_NB > -0.1 & pearson_correlation$cor_NB < 0.1 )  &  abs(pearson_correlation$cor_N
P) > 0.99, "gain" , "nothing")))

pearson_correlation$result_cor_AI_and_NB_cor_NP_a<-ifelse( (abs(pearson_correlation$subtraction_cor_AI_cor_NP) +
 abs(pearson_correlation$subtraction_cor_NB_cor_NP)) / 2 > 1.9 , "inverted",
                                   ifelse( (pearson_correlation$cor_AI >0.99  & pearson_correlation$cor_NB >0.99)
& (pearson_correlation$cor_NP > -0.1 & pearson_correlation$cor_NP < 0.1) , "loss_positive",
                                   ifelse( (pearson_correlation$cor_AI < -0.99  & pearson_correlation$cor_NB < -
0.99)  & (pearson_correlation$cor_NP > -0.1 & pearson_correlation$cor_NP < 0.1) , "loss_negative",
                                   ifelse( (pearson_correlation$cor_AI > -0.1 & pearson_correlation$cor_AI < 0.1
 & pearson_correlation$cor_NB > -0.1 & pearson_correlation$cor_NB < 0.1 )  &  pearson_correlation$cor_NP > 0.99,
"gain_positive" ,
                                   ifelse( (pearson_correlation$cor_AI > -0.1 & pearson_correlation$cor_AI < 0.1
 & pearson_correlation$cor_NB > -0.1 & pearson_correlation$cor_NB < 0.1 )  &  pearson_correlation$cor_NP < -0.99,
"gain_negative" ,"nothing")))))

#write.table(pearson_correlation, file= paste(result_path,"/2020_01_28_differential_gene_coexpression.txt", sep
=""), append = FALSE, quote = FALSE, sep = "\t" ,row.names = FALSE)


table(pearson_correlation$result_cor_AI_and_NB_cor_NP)
```

```
##
##     gain inverted     loss  nothing
##     1424       23        2 55076311
```

```
table(pearson_correlation$result_cor_AI_and_NB_cor_NP_a)
```

```
##
## gain_negative gain_positive      inverted loss_positive       nothing
##           716           708            23             2      55076311
```

### Figures 1b,c

Subset the matrix of correlation based on the results from pearson_correlation$result_cor_AI_and_NB_cor_NP_a

```
correlation_AIpreg<-cor(t(log2(rpkm_AIpreg+1)),method="pearson", use="pairwise.complete.obs")
correlation_preg_NB<-cor(t(log2(rpkm_NBpreg+1)),method="pearson", use="pairwise.complete.obs")
correlation_NOTpreg<-cor(t(log2(rpkm_NOTpreg+1)),method="pearson", use="pairwise.complete.obs")

correlation_AI_NB_preg<-(correlation_AIpreg + correlation_preg_NB)/2

pearson_correlation_a<-pearson_correlation[!(pearson_correlation$result_cor_AI_and_NB_cor_NP_a == "nothing"), ]

#write.table(pearson_correlation_a, file= paste(result_path,"/2020_01_28_differential_gene_coexpression_only_sign
ificant.txt", sep=""), append = FALSE, quote = FALSE, sep = "\t" ,row.names = FALSE)


correlation_AI_NB_preg<-correlation_AI_NB_preg[row.names(correlation_AI_NB_preg) %in% unique(c(as.character(pears
on_correlation_a$gene_1), as.character(pearson_correlation_a$gene_2))),
                                              colnames(correlation_AI_NB_preg)  %in% unique(c(as.character(pearso
n_correlation_a$gene_1), as.character(pearson_correlation_a$gene_2)))]
correlation_NOTpreg<-correlation_NOTpreg[row.names(correlation_NOTpreg) %in% unique(c(as.character(pearson_correl
ation_a$gene_1), as.character(pearson_correlation_a$gene_2))),
                                        colnames(correlation_NOTpreg)  %in% unique(c(as.character(pearson_correla
tion_a$gene_1), as.character(pearson_correlation_a$gene_2)))]



#rm(correlation_AIpreg,correlation_preg_NB,correlation_NOTpreg)

dissimilarity<-as.matrix(parDist(x= correlation_AI_NB_preg, method = "euclidean", diag = TRUE, upper = FALSE))
rownames(dissimilarity)<-colnames(correlation_AI_NB_preg)
colnames(dissimilarity)<-colnames(correlation_AI_NB_preg)
geneTree <- flashClust(as.dist(dissimilarity), method = "complete")
rm(dissimilarity)

subtraction_matrix <- correlation_NOTpreg - correlation_AI_NB_preg

correlation_AI_NB_preg<-correlation_AI_NB_preg[geneTree$order,geneTree$order ]
subtraction_matrix<-subtraction_matrix[geneTree$order,geneTree$order ]


subtraction_matrix[(subtraction_matrix >-1.8 & subtraction_matrix < -1.09) |  (subtraction_matrix > -0.89 & subtr
action_matrix < 0.89) | (subtraction_matrix > 1.09 & subtraction_matrix < 1.8)] <- 0

subtraction_matrix[upper.tri(subtraction_matrix)]<-0

correlation_AI_NB_preg[upper.tri(correlation_AI_NB_preg)]<-0
```

```
ht<-Heatmap(correlation_AI_NB_preg,
        name="correlation",
        cluster_rows= FALSE,
        cluster_columns = FALSE,
        show_row_names = FALSE,
        show_column_names = FALSE,
        col = colorRamp2(c(-2, -1,0, 1, 2), c("midnightblue","cyan1", "white", "magenta", "mediumvioletred"), spa
ce = "LAB"),
        heatmap_legend_param = list(title = "correlation  ",title_position = "lefttop",legend_direction = "horizo
ntal", title_gp = gpar(col = "black", fontsize = 5),labels_gp = gpar(col = "black", fontsize = 4 ),grid_height=un
it(0.5, "mm"),legend_width=unit(1.5, "cm"))
)
#png(paste(result_path,"/2019_07_08_AI_NB_heatmap_a.png", sep=""),width = 570, height = 625,res = 300)
#draw(ht, heatmap_legend_side = "bottom")
#dev.off()
draw(ht, heatmap_legend_side = "bottom")
```

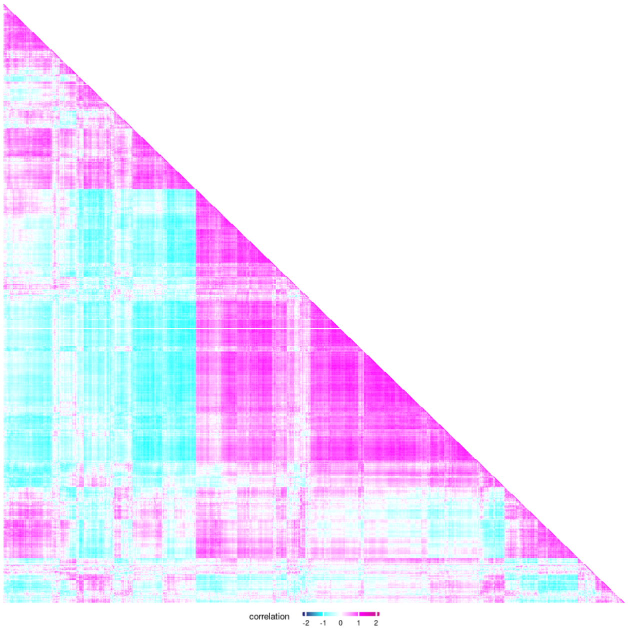


```
ht<-Heatmap(subtraction_matrix,
        name="correlation",
        cluster_rows= FALSE,
        cluster_columns = FALSE,
        show_row_names = FALSE,
        show_column_names = FALSE,
        col = colorRamp2(c(-2, -1,0, 1, 2), c("midnightblue","cyan1", "white", "magenta", "mediumvioletred"), spa
ce = "LAB"),
        heatmap_legend_param = list(title = "correlation  ",title_position = "lefttop",legend_direction = "horizo
ntal", title_gp = gpar(col = "black", fontsize = 5),labels_gp = gpar(col = "black", fontsize = 4 ),grid_height=un
it(0.5, "mm"),legend_width=unit(1.5, "cm"))
)

#png(paste(result_path,"/2019_07_08_non_preg_differential_coex_heatmap.png", sep=""),width = 570, height = 625,re
s = 300)
#draw(ht, heatmap_legend_side = "bottom")
#dev.off()
draw(ht, heatmap_legend_side = "bottom")
```

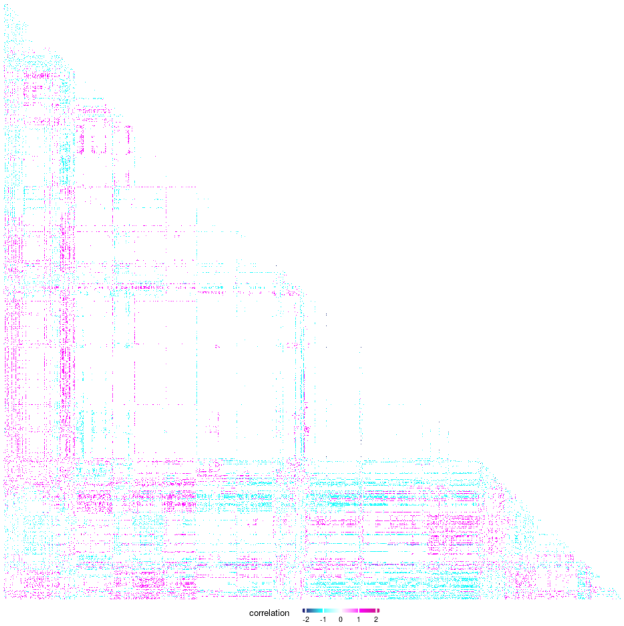


Test biological processes for enrichment with mRNAs that formed negative coexpression and calculate fold change enrichment. (AI and NB vs NP)

```
pearson_correlation_a<-pearson_correlation[pearson_correlation$result_cor_AI_and_NB_cor_NP_a == "gain_positive",]
test.genes<-data.frame(a=unique(c(as.character(pearson_correlation_a$gene_1),as.character(pearson_correlation_a$g
ene_2))), stringsAsFactors=FALSE)
all_genes_numeric<-as.integer(all_genes$gene %in%test.genes$a)
names(all_genes_numeric)<-all_genes$gene

N_DEGs<-length(test.genes$a)

set.seed(9830)
pwf<-nullp(all_genes_numeric, bias.data=annotation.genelength.biomart_vector, plot.fit=FALSE )
GO_BP_Cats_mRNA_AI_and_NB_and_NP<-goseq(pwf,gene2cat=annotation.GO.BP.biomart, method ="Sampling", repcnt = 7000,
use_genes_without_cat=FALSE)
GO_BP_Cats_mRNA_AI_and_NB_and_NP<-GO_BP_Cats_mRNA_AI_and_NB_and_NP[GO_BP_Cats_mRNA_AI_and_NB_and_NP$numDEInCat>4
,]
GO_BP_Cats_mRNA_AI_and_NB_and_NP$BY_FDR<-p.adjust(GO_BP_Cats_mRNA_AI_and_NB_and_NP$over_represented_pvalue, metho
d ="fdr")
GO_BP_Cats_mRNA_AI_and_NB_and_NP<-GO_BP_Cats_mRNA_AI_and_NB_and_NP[with(GO_BP_Cats_mRNA_AI_and_NB_and_NP, order(B
Y_FDR,over_represented_pvalue, -numDEInCat)), ]
head(GO_BP_Cats_mRNA_AI_and_NB_and_NP, n=20)

GO_BP_Cats_mRNA_AI_and_NB_and_NP$fold_enrichment<-(GO_BP_Cats_mRNA_AI_and_NB_and_NP$numDEInCat/N_DEGs)/(GO_BP_Cat
s_mRNA_AI_and_NB_and_NP$numInCat/N_expressed_genes)
annotation.GO.BP.biomart_testgenes<-annotation.GO.BP.biomart[annotation.GO.BP.biomart$ensembl_gene_id %in% test.g
enes$a, ]
GO_BP_Cats_mRNA_AI_and_NB_and_NP<-merge(GO_BP_Cats_mRNA_AI_and_NB_and_NP,annotation.GO.BP.biomart_testgenes, by.x
="category", by.y="go_id", all.x=TRUE, all.y=FALSE)
GO_BP_Cats_mRNA_AI_and_NB_and_NP<-merge(GO_BP_Cats_mRNA_AI_and_NB_and_NP, annotation.ensembl.symbol, by.x="ensemb
l_gene_id", by.y="ensembl_gene_id", all=FALSE, all.x=TRUE, all.y=FALSE)
GO_BP_Cats_mRNA_AI_and_NB_and_NP<-GO_BP_Cats_mRNA_AI_and_NB_and_NP[with(GO_BP_Cats_mRNA_AI_and_NB_and_NP, order(B
Y_FDR,term)), ]
GO_BP_Cats_mRNA_AI_and_NB_and_NP<-GO_BP_Cats_mRNA_AI_and_NB_and_NP[GO_BP_Cats_mRNA_AI_and_NB_and_NP$BY_FDR<0.2,]
```

### Supplementary Table 4

```
pearson_correlation_a<-pearson_correlation[pearson_correlation$result_cor_AI_and_NB_cor_NP_a == "gain_negative",]
test.genes<-data.frame(a=unique(c(as.character(pearson_correlation_a$gene_1),as.character(pearson_correlation_a$g
ene_2))), stringsAsFactors=FALSE)
all_genes_numeric<-as.integer(all_genes$gene %in%test.genes$a)
names(all_genes_numeric)<-all_genes$gene

N_DEGs<-length(test.genes$a)

set.seed(9830)
pwf<-nullp(all_genes_numeric, bias.data=annotation.genelength.biomart_vector, plot.fit=FALSE )
GO_BP_Cats_mRNA_AI_and_NB_and_NP<-goseq(pwf,gene2cat=annotation.GO.BP.biomart, method ="Sampling", repcnt = 7000,
use_genes_without_cat=FALSE)
GO_BP_Cats_mRNA_AI_and_NB_and_NP<-GO_BP_Cats_mRNA_AI_and_NB_and_NP[GO_BP_Cats_mRNA_AI_and_NB_and_NP$numDEInCat>4
,]
GO_BP_Cats_mRNA_AI_and_NB_and_NP$BY_FDR<-p.adjust(GO_BP_Cats_mRNA_AI_and_NB_and_NP$over_represented_pvalue, metho
d ="fdr")
GO_BP_Cats_mRNA_AI_and_NB_and_NP<-GO_BP_Cats_mRNA_AI_and_NB_and_NP[with(GO_BP_Cats_mRNA_AI_and_NB_and_NP, order(B
Y_FDR,over_represented_pvalue, -numDEInCat)), ]
#head(GO_BP_Cats_mRNA_AI_and_NB_and_NP, n=20)

GO_BP_Cats_mRNA_AI_and_NB_and_NP$fold_enrichment<-(GO_BP_Cats_mRNA_AI_and_NB_and_NP$numDEInCat/N_DEGs)/(GO_BP_Cat
s_mRNA_AI_and_NB_and_NP$numInCat/N_expressed_genes)
annotation.GO.BP.biomart_testgenes<-annotation.GO.BP.biomart[annotation.GO.BP.biomart$ensembl_gene_id %in% test.g
enes$a, ]
GO_BP_Cats_mRNA_AI_and_NB_and_NP<-merge(GO_BP_Cats_mRNA_AI_and_NB_and_NP,annotation.GO.BP.biomart_testgenes, by.x
="category", by.y="go_id", all.x=TRUE, all.y=FALSE)
GO_BP_Cats_mRNA_AI_and_NB_and_NP<-merge(GO_BP_Cats_mRNA_AI_and_NB_and_NP, annotation.ensembl.symbol, by.x="ensemb
l_gene_id", by.y="ensembl_gene_id", all=FALSE, all.x=TRUE, all.y=FALSE)
GO_BP_Cats_mRNA_AI_and_NB_and_NP<-GO_BP_Cats_mRNA_AI_and_NB_and_NP[with(GO_BP_Cats_mRNA_AI_and_NB_and_NP, order(B
Y_FDR,term)), ]
GO_BP_Cats_mRNA_AI_and_NB_and_NP<-GO_BP_Cats_mRNA_AI_and_NB_and_NP[GO_BP_Cats_mRNA_AI_and_NB_and_NP$BY_FDR<0.1,]
#write.table(GO_BP_Cats_mRNA_AI_and_NB_and_NP, paste(result_path, "2019_12_06_GO_BP_Cats_mRNA_diff_coexpression_g
ain_neg_AI_and_NB_and_NP.txt", sep="/"), quote=FALSE, sep="\t", row.names=FALSE)
```

```
knitr::kable(head(GO_BP_Cats_mRNA_AI_and_NB_and_NP), caption = "First five rows of Supplementary Table 4",format
 = 'pandoc')
```

First five rows of Supplementary Table 4

| ensembl_gene_id | category | over_represented_pvalue | under_represented_pvalue | numDEInCat | numInCat | term |
|---|---|---|---|---|---|---|

| | ensembl_gene_id | category | over_represented_pvalue | under_represented_pvalue | numDEInCat | numInCat | term |
|---|---|---|---|---|---|---|---|
| 10 | ENSBTAG00000000131 | GO:0006508 (GO:0006508) | 0.0005713 | 0.9998572 | 38 | 232 | proteolysis |
| 51 | ENSBTAG00000000725 | GO:0006508 (GO:0006508) | 0.0005713 | 0.9998572 | 38 | 232 | proteolysis |
| 83 | ENSBTAG00000001134 | GO:0006508 (GO:0006508) | 0.0005713 | 0.9998572 | 38 | 232 | proteolysis |
| 104 | ENSBTAG00000001353 | GO:0006508 (GO:0006508) | 0.0005713 | 0.9998572 | 38 | 232 | proteolysis |
| 230 | ENSBTAG00000003039 | GO:0006508 (GO:0006508) | 0.0005713 | 0.9998572 | 38 | 232 | proteolysis |
| 266 | ENSBTAG00000003395 | GO:0006508 (GO:0006508) | 0.0005713 | 0.9998572 | 38 | 232 | proteolysis |

#### Figure 1d

```
proteolysis_genes<-GO_BP_Cats_mRNA_AI_and_NB_and_NP[GO_BP_Cats_mRNA_AI_and_NB_and_NP$term == "proteolysis", 1]
pearson_correlation_b<-pearson_correlation_a[pearson_correlation_a$gene_1 %in% proteolysis_genes | pearson_correl
ation_a$gene_2 %in% proteolysis_genes ,]
pearson_correlation_b<-pearson_correlation_b[with(pearson_correlation_b, order(cor_NP)), ]

data_for_chart<-data.frame()
data_for_chart_a<-data.frame()
for(i in 1:6 ){
  expression_gene_1_AI<-rpkm_AIpreg[row.names(rpkm_AIpreg)==as.character(pearson_correlation_b$gene_1)[i],]
  expression_gene_2_AI<-rpkm_AIpreg[row.names(rpkm_AIpreg)==as.character(pearson_correlation_b$gene_2)[i],]
  expression_gene_1_NB<-rpkm_NBpreg[row.names(rpkm_NBpreg)==as.character(pearson_correlation_b$gene_1)[i],]
  expression_gene_2_NB<-rpkm_NBpreg[row.names(rpkm_NBpreg)==as.character(pearson_correlation_b$gene_2)[i],]
  expression_gene_1_NP<-rpkm_NOTpreg[row.names(rpkm_NOTpreg)==as.character(pearson_correlation_b$gene_1)[i],]
  expression_gene_2_NP<-rpkm_NOTpreg[row.names(rpkm_NOTpreg)==as.character(pearson_correlation_b$gene_2)[i],]

  symbol_gene_1<-annotation.ensembl.symbol[annotation.ensembl.symbol$ensembl_gene_id == as.character(pearson_corr
elation_b$gene_1)[i] , 2]
  symbol_gene_2<-annotation.ensembl.symbol[annotation.ensembl.symbol$ensembl_gene_id == as.character(pearson_corr
elation_b$gene_2)[i] , 2]
  chart<-i

  data_for_chart<-data.frame(gene_1=c(expression_gene_1_AI,expression_gene_1_NB,expression_gene_1_NP) ,
                             gene_2=c(expression_gene_2_AI,expression_gene_2_NB,expression_gene_2_NP),
                             symbol_gene_1,symbol_gene_2 ,chart,
                             group=rep(c("AI", "NB", "NP"), c(length(expression_gene_1_AI),length(expression_gene
_1_NB),length(expression_gene_1_NP))))

  data_for_chart_a<-rbind(data_for_chart_a,data_for_chart)
}

font_axis<-5
font_title<-8
ptsize<-0.05

plots<-list()
k<-1
for (j in c(1:6)){
  data_for_chart_b<-data_for_chart_a[data_for_chart_a$chart %in% j,]

plot<-ggplot(data_for_chart_b, aes(x=gene_1,y=gene_2))+
  geom_point()+
  geom_smooth(method=lm, size=ptsize, fill="lightgray")+
  scale_y_continuous(name=data_for_chart_b$symbol_gene_1[1])+
  scale_x_continuous(name=data_for_chart_b$symbol_gene_2[1])+
  facet_wrap(~group, ncol=3, scales="free")+
  theme(aspect.ratio = 1,
            panel.grid.major = element_blank(),
            panel.grid.minor = element_blank(),
            panel.background = element_blank(),
            plot.background  = element_blank(),
            axis.text = element_text( colour = 'black',size = font_axis),
            axis.title = element_text( colour = 'black' ,size = font_title, face="italic"),
            axis.ticks = element_line(size=0.1),
            panel.spacing.y = unit(0, "mm"),
            panel.spacing.x = unit(1.5, "mm"),
            legend.position="none",
            axis.line=element_line(size = 0.1, colour = "black"),
            strip.background=element_blank(),
            strip.text.x = element_blank())

 plots[[k]]<-plot
  k<-k+1
}

#png(filename =paste(result_path,"2019_11_14_plot_diff_cor_AI_NB_and_NP_gain_negative.png", sep="/"), width = 5.
2, height =3, units = 'in', res = 300,bg = "transparent")
#ggarrange(plotlist=plots, ncol=2, nrow=3)
#dev.off()
ggarrange(plotlist=plots, ncol=2, nrow=3)
```

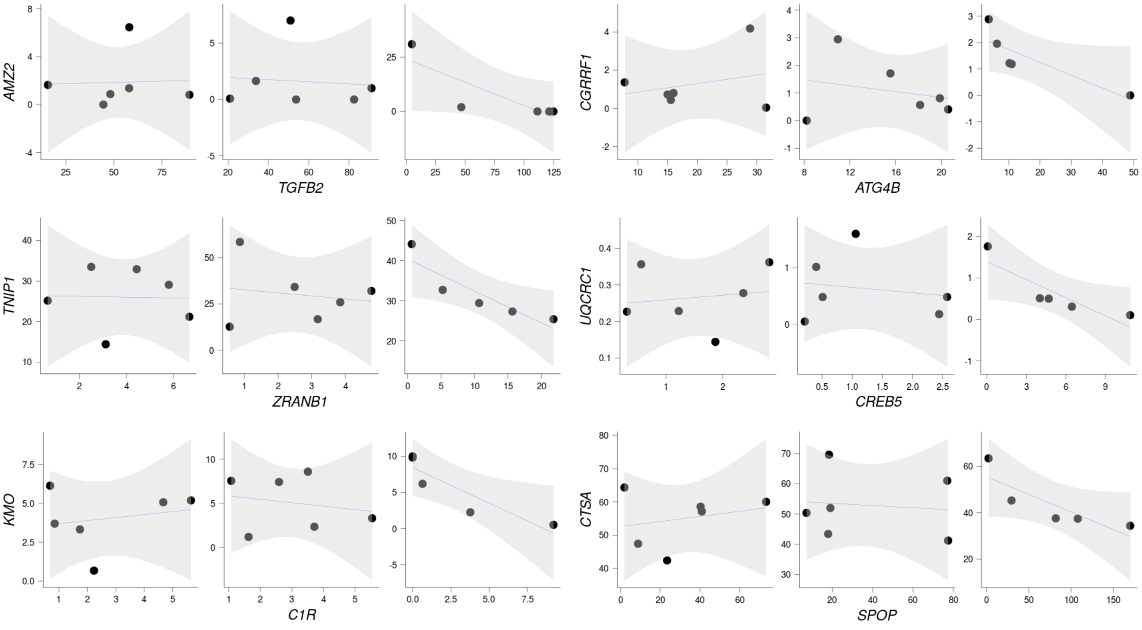


### Supplementary Table 5

```
pearson_correlation_a<-pearson_correlation[pearson_correlation$result_cor_AI_and_NB_cor_NP_a == "inverted",]
test.genes<-data.frame(a=unique(c(as.character(pearson_correlation_a$gene_1),as.character(pearson_correlation_a$g
ene_2))), stringsAsFactors=FALSE)
all_genes_numeric<-as.integer(all_genes$gene %in%test.genes$a)
names(all_genes_numeric)<-all_genes$gene

N_DEGs<-length(test.genes$a)

set.seed(9830)
pwf<-nullp(all_genes_numeric, bias.data=annotation.genelength.biomart_vector, plot.fit=FALSE )
GO_BP_Cats_mRNA_AI_and_NB_and_NP<-goseq(pwf,gene2cat=annotation.GO.BP.biomart, method ="Sampling", repcnt = 7000,
use_genes_without_cat=FALSE)
GO_BP_Cats_mRNA_AI_and_NB_and_NP<-GO_BP_Cats_mRNA_AI_and_NB_and_NP[GO_BP_Cats_mRNA_AI_and_NB_and_NP$numDEInCat>4
,]
GO_BP_Cats_mRNA_AI_and_NB_and_NP$BY_FDR<-p.adjust(GO_BP_Cats_mRNA_AI_and_NB_and_NP$over_represented_pvalue, metho
d ="fdr")
GO_BP_Cats_mRNA_AI_and_NB_and_NP<-GO_BP_Cats_mRNA_AI_and_NB_and_NP[with(GO_BP_Cats_mRNA_AI_and_NB_and_NP, order(B
Y_FDR,over_represented_pvalue, -numDEInCat)), ]
#head(GO_BP_Cats_mRNA_AI_and_NB_and_NP, n=20)

GO_BP_Cats_mRNA_AI_and_NB_and_NP$fold_enrichment<-(GO_BP_Cats_mRNA_AI_and_NB_and_NP$numDEInCat/N_DEGs)/(GO_BP_Cat
s_mRNA_AI_and_NB_and_NP$numInCat/N_expressed_genes)
annotation.GO.BP.biomart_testgenes<-annotation.GO.BP.biomart[annotation.GO.BP.biomart$ensembl_gene_id %in% test.g
enes$a, ]
GO_BP_Cats_mRNA_AI_and_NB_and_NP<-merge(GO_BP_Cats_mRNA_AI_and_NB_and_NP,annotation.GO.BP.biomart_testgenes, by.x
="category", by.y="go_id", all.x=TRUE, all.y=FALSE)
GO_BP_Cats_mRNA_AI_and_NB_and_NP<-merge(GO_BP_Cats_mRNA_AI_and_NB_and_NP, annotation.ensembl.symbol, by.x="ensemb
l_gene_id", by.y="ensembl_gene_id", all=FALSE, all.x=TRUE, all.y=FALSE)
GO_BP_Cats_mRNA_AI_and_NB_and_NP<-GO_BP_Cats_mRNA_AI_and_NB_and_NP[with(GO_BP_Cats_mRNA_AI_and_NB_and_NP, order(B
Y_FDR,term)), ]
GO_BP_Cats_mRNA_AI_and_NB_and_NP<-GO_BP_Cats_mRNA_AI_and_NB_and_NP[GO_BP_Cats_mRNA_AI_and_NB_and_NP$BY_FDR<0.2,]
#write.table(GO_BP_Cats_mRNA_AI_and_NB_and_NP, paste(result_path, "2019_12_06_GO_BP_Cats_mRNA_diff_coexpression_i
nverted_AI_and_NB_and_NP.txt", sep="/"), quote=FALSE, sep="\t", row.names=FALSE)
```

```
knitr::kable(head(GO_BP_Cats_mRNA_AI_and_NB_and_NP), caption = "First five rows of Supplementary Table 5",format
 = 'pandoc')
```

First five rows of Supplementary Table 5

| ensembl_gene_id | category | over_represented_pvalue | under_represented_pvalue | numDEInCat | numInCat | term | o |
|---|---|---|---|---|---|---|---|

| ensembl_gene_id | category | over_represented_pvalue | under_represented_pvalue | numDEInCat | numInCat | term | o |
|---|---|---|---|---|---|---|---|
| ENSBTAG00000000023 | GO:0000122 (GO:0000122) | 0.0204257 | 0.9950007 | 5 | 408 | negative regulation of transcription by RNA polymerase II | B |
| ENSBTAG00000001163 | GO:0000122 (GO:0000122) | 0.0204257 | 0.9950007 | 5 | 408 | negative regulation of transcription by RNA polymerase II | B |
| ENSBTAG00000005564 | GO:0000122 (GO:0000122) | 0.0204257 | 0.9950007 | 5 | 408 | negative regulation of transcription by RNA polymerase II | B |
| ENSBTAG00000012178 | GO:0000122 (GO:0000122) | 0.0204257 | 0.9950007 | 5 | 408 | negative regulation of transcription by RNA polymerase II | B |
| ENSBTAG00000021427 | GO:0000122 (GO:0000122) | 0.0204257 | 0.9950007 | 5 | 408 | negative regulation of transcription by RNA polymerase II | B |

#### Figure 1e

```
pearson_correlation_b<-pearson_correlation_a[pearson_correlation_a$gene_1 %in% GO_BP_Cats_mRNA_AI_and_NB_and_NP$e
nsembl_gene_id | pearson_correlation_a$gene_2 %in% GO_BP_Cats_mRNA_AI_and_NB_and_NP$ensembl_gene_id ,]

data_for_chart<-data.frame()
data_for_chart_a<-data.frame()
for(i in 1:dim(pearson_correlation_b)[1] ){
  expression_gene_1_AI<-rpkm_AIpreg[row.names(rpkm_AIpreg)==as.character(pearson_correlation_b$gene_1)[i],]
  expression_gene_2_AI<-rpkm_AIpreg[row.names(rpkm_AIpreg)==as.character(pearson_correlation_b$gene_2)[i],]
  expression_gene_1_NB<-rpkm_NBpreg[row.names(rpkm_NBpreg)==as.character(pearson_correlation_b$gene_1)[i],]
  expression_gene_2_NB<-rpkm_NBpreg[row.names(rpkm_NBpreg)==as.character(pearson_correlation_b$gene_2)[i],]
  expression_gene_1_NP<-rpkm_NOTpreg[row.names(rpkm_NOTpreg)==as.character(pearson_correlation_b$gene_1)[i],]
  expression_gene_2_NP<-rpkm_NOTpreg[row.names(rpkm_NOTpreg)==as.character(pearson_correlation_b$gene_2)[i],]

  symbol_gene_1<-annotation.ensembl.symbol[annotation.ensembl.symbol$ensembl_gene_id == as.character(pearson_corr
elation_b$gene_1)[i] , 2]
  symbol_gene_2<-annotation.ensembl.symbol[annotation.ensembl.symbol$ensembl_gene_id == as.character(pearson_corr
elation_b$gene_2)[i] , 2]
  chart<-i

  data_for_chart<-data.frame(gene_1=c(expression_gene_1_AI,expression_gene_1_NB,expression_gene_1_NP) ,
                             gene_2=c(expression_gene_2_AI,expression_gene_2_NB,expression_gene_2_NP),
                             symbol_gene_1,symbol_gene_2 ,chart,
                             group=rep(c("AI", "NB", "NP"), c(length(expression_gene_1_AI),length(expression_gene
_1_NB),length(expression_gene_1_NP))))

  data_for_chart_a<-rbind(data_for_chart_a,data_for_chart)
}

font_axis<-5
font_title<-8
ptsize<-0.05

plots<-list()
k<-1
for (j in c(1:4)){
  data_for_chart_b<-data_for_chart_a[data_for_chart_a$chart %in% j,]

plot<-ggplot(data_for_chart_b, aes(x=gene_1,y=gene_2))+
  geom_point()+
  geom_smooth(method=lm, size=ptsize, fill="lightgray")+
  scale_y_continuous(name=data_for_chart_b$symbol_gene_1[1])+
  scale_x_continuous(name=data_for_chart_b$symbol_gene_2[1])+
  facet_wrap(~group, ncol=3, scales="free")+
  theme(aspect.ratio = 1,
            panel.grid.major = element_blank(),
            panel.grid.minor = element_blank(),
            panel.background = element_blank(),
            plot.background  = element_blank(),
            axis.text = element_text( colour = 'black',size = font_axis),
            axis.title = element_text( colour = 'black' ,size = font_title, face="italic"),
            axis.ticks = element_line(size=0.1),
            panel.spacing.y = unit(0, "mm"),
            panel.spacing.x = unit(2, "mm"),
            legend.position="none",
            axis.line=element_line(size = 0.1, colour = "black"),
            strip.background=element_blank(),
            strip.text.x = element_blank())

 plots[[k]]<-plot
  k<-k+1
}

#png(filename =paste(result_path,"2019_11_14_plot_diff_cor_AI_NB_and_NP_inverted.png", sep="/"), width = 5.2, hei
ght =1.8, units = 'in', res = 300,bg = "transparent")
#ggarrange(plotlist=plots, ncol=2, nrow=2)
#dev.off()
ggarrange(plotlist=plots, ncol=2, nrow=2)
```

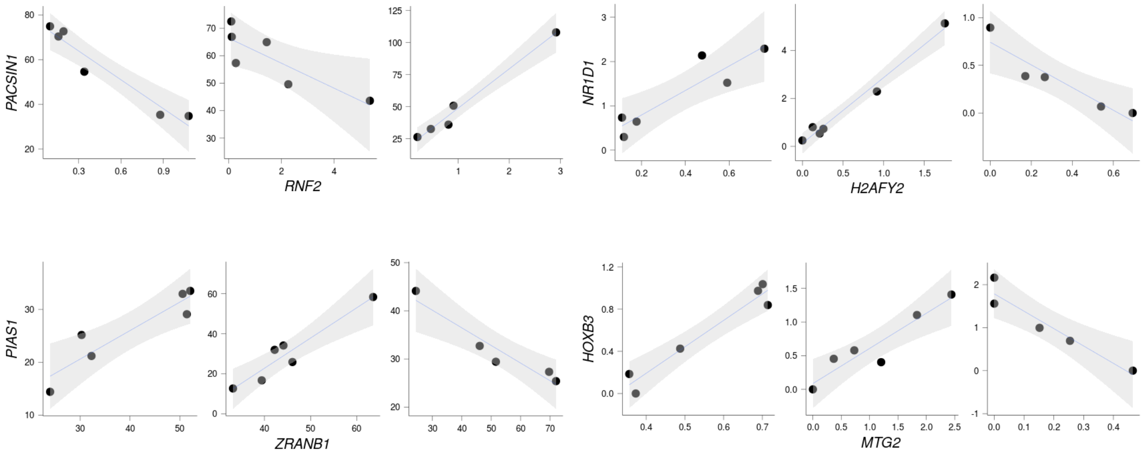


```
gc()
```

### Differential miRNA:mRNA coexpression associated with pregnancy outcome

#### Supplementary table 6

```
correlation_all_animals_miRNA<-corAndPvalue( t(log2(rpkm_pwbc +1)), t(log2(rpkm_miRNA +1)) ,method="pearson", use
="pairwise.complete.obs")
correlation_all_animals_miRNA_a<-melt(correlation_all_animals_miRNA$cor)
correlation_all_animals_miRNA_a$pvalue<-melt(correlation_all_animals_miRNA$p)$value

correlation_all_animals_miRNA_a<-correlation_all_animals_miRNA_a[!duplicated(t(apply(correlation_all_animals_miRN
A_a[,c(1:2)], 1, sort))),]

correlation_all_animals_miRNA_b<-correlation_all_animals_miRNA_a

correlation_all_animals_miRNA_b$gene_1<-annotation.ensembl.symbol$external_gene_name[match( correlation_all_anima
ls_miRNA_b$Var1, annotation.ensembl.symbol$ensembl_gene_id )]
correlation_all_animals_miRNA_b$gene_2<-annotation.ensembl.symbol$external_gene_name[match( correlation_all_anima
ls_miRNA_b$Var2, annotation.ensembl.symbol$ensembl_gene_id )]
correlation_all_animals_miRNA_b$Var1<-as.character(correlation_all_animals_miRNA_b$Var1)
correlation_all_animals_miRNA_b$Var2<-as.character(correlation_all_animals_miRNA_b$Var2)

correlation_all_animals_miRNA_b$gene_1<-ifelse(correlation_all_animals_miRNA_b$gene_1=="", correlation_all_animal
s_miRNA_b$Var1,correlation_all_animals_miRNA_b$gene_1)
correlation_all_animals_miRNA_b$gene_2<-ifelse(correlation_all_animals_miRNA_b$gene_2=="", correlation_all_animal
s_miRNA_b$Var2,correlation_all_animals_miRNA_b$gene_2)

#write.table(correlation_all_animals_miRNA_b, file= paste(result_path,"2019_12_20_mRNA_miNRA_correlation_all_anim
als_miRNA.txt", sep="/"), append = FALSE, quote = FALSE, sep = "\t" ,row.names = FALSE)
#system("bzip2 -9 /mnt/storage/auburn/heifer_pregnancy/proj_2018/analysis/results/2019_12_20_mRNA_miNRA_correlati
on_all_animals_miRNA.txt")


stri1 <- function(x){
  ind <- stri_endswith_fixed(x, "-1")
  x[ind] <- stri_sub(x[ind],1, -3)
  x
}

stri2 <- function(x){
  ind <- stri_endswith_fixed(x, "-2")
  x[ind] <- stri_sub(x[ind],1, -3)
  x
}

stri3 <- function(x){
  ind <- stri_endswith_fixed(x, "-3")
  x[ind] <- stri_sub(x[ind],1, -3)
  x
}

stri4 <- function(x){
  ind <- stri_endswith_fixed(x, "-4")
  x[ind] <- stri_sub(x[ind],1, -3)
  x
}

stri5 <- function(x){
  ind <- stri_endswith_fixed(x, "-5")
  x[ind] <- stri_sub(x[ind],1, -3)
  x
}

correlation_all_animals_miRNA_b$gene_2<-stri1(correlation_all_animals_miRNA_b$gene_2)
correlation_all_animals_miRNA_b$gene_2<-stri2(correlation_all_animals_miRNA_b$gene_2)
correlation_all_animals_miRNA_b$gene_2<-stri3(correlation_all_animals_miRNA_b$gene_2)
correlation_all_animals_miRNA_b$gene_2<-stri4(correlation_all_animals_miRNA_b$gene_2)
correlation_all_animals_miRNA_b$gene_2<-stri5(correlation_all_animals_miRNA_b$gene_2)


correlation_all_animals_miRNA_b<-correlation_all_animals_miRNA_b[abs(correlation_all_animals_miRNA_b$value) > 0.8
5,]
#write.table(correlation_all_animals_miRNA_b, file= paste(result_path,"2020_01_29_mRNA_miNRA_sig_correlation_all_
animals_miRNA.txt", sep="/"), append = FALSE, quote = FALSE, sep = "\t" ,row.names = FALSE)
```

```
knitr::kable(head(correlation_all_animals_miRNA_b), caption = "First five rows of Supplementary Table 6",format =
'pandoc')
```

First five rows of Supplementary Table 6

| | Var1 | Var2 | value | pvalue | gene_1 | gene_2 |
|---|---|---|---|---|---|---|
| 1695 | ENSBTAG00000004457 | ENSBTAG00000029760 | -0.8568286 | 1.12e-05 | ORAI1 | bta-mir-320a |

| | Var1 | Var2 | value | pvalue | gene_1 | gene_2 |
|---|---|---|---|---|---|---|
| 3125 | ENSBTAG00000008309 | ENSBTAG00000029760 | -0.8552897 | 1.21e-05 | ATP13A2 | bta-mir-320a |
| 4135 | ENSBTAG00000011064 | ENSBTAG00000029760 | -0.8519537 | 1.42e-05 | ADCK5 | bta-mir-320a |
| 4551 | ENSBTAG00000012172 | ENSBTAG00000029760 | -0.8553540 | 1.21e-05 | MIDN | bta-mir-320a |
| 10155 | ENSBTAG00000051917 | ENSBTAG00000029760 | -0.8514464 | 1.45e-05 | ZNF579 | bta-mir-320a |
| 12191 | ENSBTAG00000004457 | ENSBTAG00000029761 | -0.8749960 | 4.30e-06 | ORAI1 | bta-mir-320a |

### Supplementary Fig. 2

Calculate the empirical false discovery rate for Pearson’s co-expression and differential coexpression

```
permutation_matrix<-matrix(,nrow=200100,ncol=17)
#head(permutation_matrix)
#dim(permutation_matrix)

for (i in seq(1:200100)){
  sampling <- sample(1:17,17, replace = FALSE)
  if ( ! identical(sampling , c(1:17))){
    permutation_matrix[i,]<-sample(1:17,17, replace = FALSE)
  }}

permutation_matrix<-permutation_matrix[!duplicated(permutation_matrix),]
dim(permutation_matrix)
head(permutation_matrix)

permutation_matrix<-permutation_matrix[sample(1:dim(permutation_matrix)[1], 5000),]
rand<-dim(permutation_matrix)[1]

coexp<-seq(0.8, 1, 0.01)

results <- filebacked.big.matrix(length(coexp),rand, type="double", init=0, separated=FALSE,
                                 backingfile="incidence_matrix.bin",
                                 descriptor="incidence_matrix.desc")
mdesc_result<- describe(results)

cl <- makeCluster(20)
registerDoParallel(cl)

results[,]<-foreach(i = coexp, .combine='rbind',.packages=c("reshape2","bigmemory"), .inorder=TRUE, .verbose=TRUE
) %:%
            foreach(j = 1:rand, .combine='cbind',.packages=c("reshape2","bigmemory"), .inorder=FALSE, .verbose=TR
UE ) %dopar% {

   rpkm_pwbc_scramble<-log2(rpkm_pwbc[,permutation_matrix[j,]] +1)
   cor_1<-cor( t(rpkm_pwbc_scramble), t(log2(rpkm_miRNA +1)) ,method="pearson", use="pairwise.complete.obs")
   length(which(abs(cor_1) > i))
  }
stopCluster(cl)

total.rand <-  5000 * 10496 * 290

qvalue<-data.frame(correlation = coexp,
                   e.pvalue= (rowSums(results[,]+1))/(total.rand+1),
                   e.pvalue.round= round((rowSums(results[,]+1))/(total.rand+1) ,6))

qvalue


write.table(qvalue, file= paste(result_path,"2019_12_09_mRNA_miNRA_coexpression_empiricalFDR.txt", sep="/"), appe
nd = FALSE, quote = FALSE, sep = "\t" ,row.names = FALSE)


rm(results)
system("rm incidence_matrix.bin")
system("rm incidence_matrix.desc")
rm(permutation_matrix)
```

```
eFDR_coexpression<-read.delim(paste(result_path,"2019_12_09_mRNA_miNRA_coexpression_empiricalFDR.txt", sep="/"),
 row.names=NULL,header =TRUE, stringsAsFactors =FALSE)
```

```
font_size<-12 #font_size 12 for pdf  figure

scaleFUN <- function(x) sprintf("%.2f", x)

plot1<-ggplot()+
  geom_point(data=eFDR_coexpression, aes(x=correlation , y=e.pvalue.round),color="black", size=3, shape=16)+
  geom_line(data=eFDR_coexpression, aes(x=correlation , y=e.pvalue.round),color="black", size=0.1,linetype=3)+
  scale_y_continuous(name="empirical FDR", limits = c(0, 0.0002), breaks=seq(0,0.0002, 0.00001))+
  scale_x_continuous(name="correlation (r)", limits = c(0.8, 1), breaks=seq(0.8,1, 0.01), labels=scaleFUN)+
  ggtitle("mRNA and miRNA coexpression all heifers")+
  theme_bw()+
  theme(panel.grid= element_blank(),
        panel.background = element_blank(),
        panel.grid.minor = element_blank(),
        panel.grid.major = element_line(color="lightgray"),
        plot.background = element_blank(),
        axis.title=element_text(color="black", size=font_size),
        axis.text.y=element_text(color="black", size=font_size),
        axis.text.x=element_text(color="black", size=font_size, angle=90),
        panel.spacing = unit(c(0.4,0.4,0.4,0.4),"cm"),
        plot.margin = unit(c(0.5,0.5,0.5,0.5),"cm"),
        legend.position="none",
        plot.title = element_text(lineheight=.8, hjust=0.5, size= font_size))


#font_size 12 for figure
#pdf(paste(result_path,"2019_12_11_mRNA_miRNA_eFDR_plots.pdf", sep="/"), width=9, height=5)
#plot1
#dev.off()

plot1
```

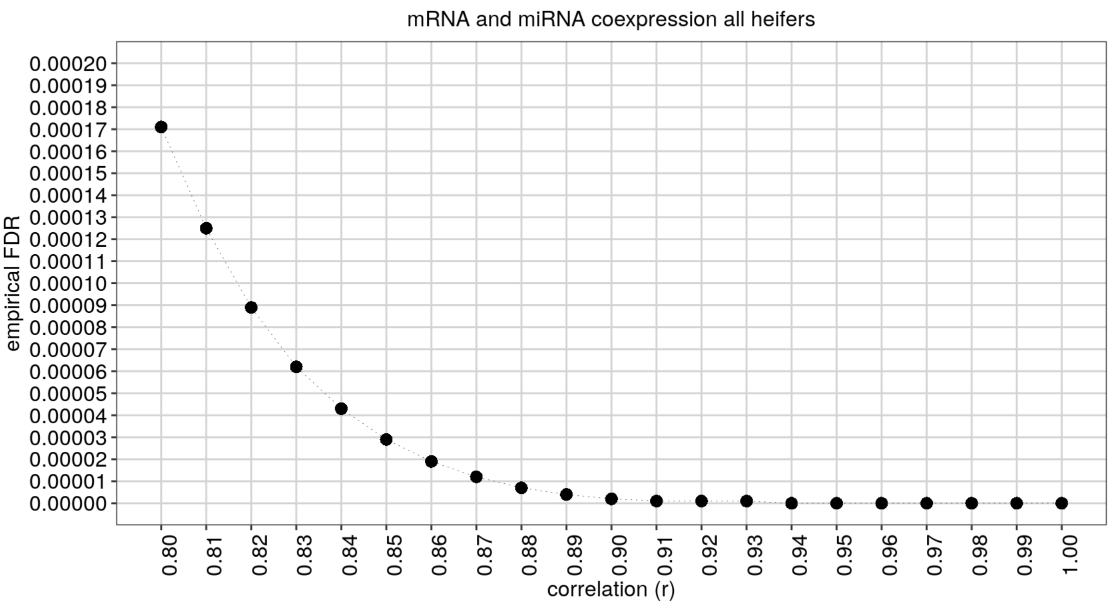


### Supplementary Table 7

```
test.genes<-data.frame(a=unique(as.character(correlation_all_animals_miRNA_b$Var1)), stringsAsFactors=FALSE)
all_genes_numeric<-as.integer(all_genes$gene %in%test.genes$a)
names(all_genes_numeric)<-all_genes$gene

N_DEGs<-length(test.genes$a)

set.seed(54345)
pwf<-nullp(all_genes_numeric, bias.data=annotation.genelength.biomart_vector, plot.fit=FALSE )
GO_BP_Cats_mRNA_miRNA_all_animals<-goseq(pwf,gene2cat=annotation.GO.BP.biomart, method ="Sampling", repcnt = 7000
, use_genes_without_cat=FALSE)
GO_BP_Cats_mRNA_miRNA_all_animals<-GO_BP_Cats_mRNA_miRNA_all_animals[GO_BP_Cats_mRNA_miRNA_all_animals$numDEInCat
>=4,]
GO_BP_Cats_mRNA_miRNA_all_animals$BY_FDR<-p.adjust(GO_BP_Cats_mRNA_miRNA_all_animals$over_represented_pvalue, met
hod ="fdr")
GO_BP_Cats_mRNA_miRNA_all_animals<-GO_BP_Cats_mRNA_miRNA_all_animals[with(GO_BP_Cats_mRNA_miRNA_all_animals, orde
r(BY_FDR,over_represented_pvalue, -numDEInCat)), ]
#head(GO_BP_Cats_mRNA_miRNA_all_animals, n=20)

GO_BP_Cats_mRNA_miRNA_all_animals$fold_enrichment<-(GO_BP_Cats_mRNA_miRNA_all_animals$numDEInCat/N_DEGs)/(GO_BP_C
ats_mRNA_miRNA_all_animals$numInCat/N_expressed_genes)
annotation.GO.BP.biomart_testgenes<-annotation.GO.BP.biomart[annotation.GO.BP.biomart$ensembl_gene_id %in% test.g
enes$a, ]
GO_BP_Cats_mRNA_miRNA_all_animals<-merge(GO_BP_Cats_mRNA_miRNA_all_animals,annotation.GO.BP.biomart_testgenes, b
y.x="category", by.y="go_id", all.x=TRUE, all.y=FALSE)
GO_BP_Cats_mRNA_miRNA_all_animals<-merge(GO_BP_Cats_mRNA_miRNA_all_animals, annotation.ensembl.symbol, by.x="ense
mbl_gene_id", by.y="ensembl_gene_id", all=FALSE, all.x=TRUE, all.y=FALSE)
GO_BP_Cats_mRNA_miRNA_all_animals<-GO_BP_Cats_mRNA_miRNA_all_animals[with(GO_BP_Cats_mRNA_miRNA_all_animals, orde
r(BY_FDR,term)), ]
GO_BP_Cats_mRNA_miRNA_all_animals<-GO_BP_Cats_mRNA_miRNA_all_animals[GO_BP_Cats_mRNA_miRNA_all_animals$BY_FDR<0.1
,]
#write.table(GO_BP_Cats_mRNA_miRNA_all_animals, paste(result_path, "2020_01_29_GO_BP_Cats_mRNA_miRNA_all_animals.
txt", sep="/"), quote=FALSE, sep="\t", row.names=FALSE)
```

```
knitr::kable(head(GO_BP_Cats_mRNA_miRNA_all_animals), caption = "First five rows of Supplementary Table 7",format
= 'pandoc')
```

First five rows of Supplementary Table 7

| | ensembl_gene_id | category | over_represented_pvalue | under_represented_pvalue | numDEInCat | numInCat | term |
|---|---|---|---|---|---|---|---|
| 17 | ENSBTAG00000005408 | GO:0018107 (GO:0018107) | 0.0004285 | 1.0000000 | 4 | 42 | peptidyl-threonine phosphoryl |
| 29 | ENSBTAG00000017401 | GO:0018107 (GO:0018107) | 0.0004285 | 1.0000000 | 4 | 42 | peptidyl-threonine phosphoryl |
| 48 | ENSBTAG00000033333 | GO:0018107 (GO:0018107) | 0.0004285 | 1.0000000 | 4 | 42 | peptidyl-threonine phosphoryl |
| 49 | ENSBTAG00000033801 | GO:0018107 (GO:0018107) | 0.0004285 | 1.0000000 | 4 | 42 | peptidyl-threonine phosphoryl |
| 35 | ENSBTAG00000019915 | GO:0060271 (GO:0060271) | 0.0164262 | 0.9972861 | 4 | 106 | cilium asse |
| 46 | ENSBTAG00000030175 | GO:0060271 (GO:0060271) | 0.0164262 | 0.9972861 | 4 | 106 | cilium asse |

### Figure 3a

```
correlation_all_animals_miRNA_c<-correlation_all_animals_miRNA_b[,c("gene_2", "gene_1", "value")]
correlation_all_animals_miRNA_c<-correlation_all_animals_miRNA_c[correlation_all_animals_miRNA_c$gene_1 %in% GO_B
P_Cats_mRNA_miRNA_all_animals$external_gene_name,]
colnames(correlation_all_animals_miRNA_c)<- c("source", "target", "value")

correlation_all_animals_miRNA_c<-rbind(correlation_all_animals_miRNA_c, data.frame(source=GO_BP_Cats_mRNA_miRNA_a
ll_animals[,c("external_gene_name")],target=GO_BP_Cats_mRNA_miRNA_all_animals[,c("term")], value=1))

ids <- unique(as.character(c(correlation_all_animals_miRNA_c$source, correlation_all_animals_miRNA_c$target)))
nodes <- data.frame(node = c(0:23),
                    name = ids)

results <- merge(correlation_all_animals_miRNA_c, nodes, by.x = "source", by.y = "name")
results <- merge(results, nodes, by.x = "target", by.y = "name")
links <- results[ , c("node.x", "node.y", "value")]
colnames(links) <- c("source", "target", "value")

links$value<-1

networkD3::sankeyNetwork(Links = links, Nodes = nodes, Source = 'source',
                         Target = 'target',  NodeID = 'name', Value = "value",
                         fontSize = 15,
                        nodeWidth = 5,
                        nodePadding = 5
                        )
```

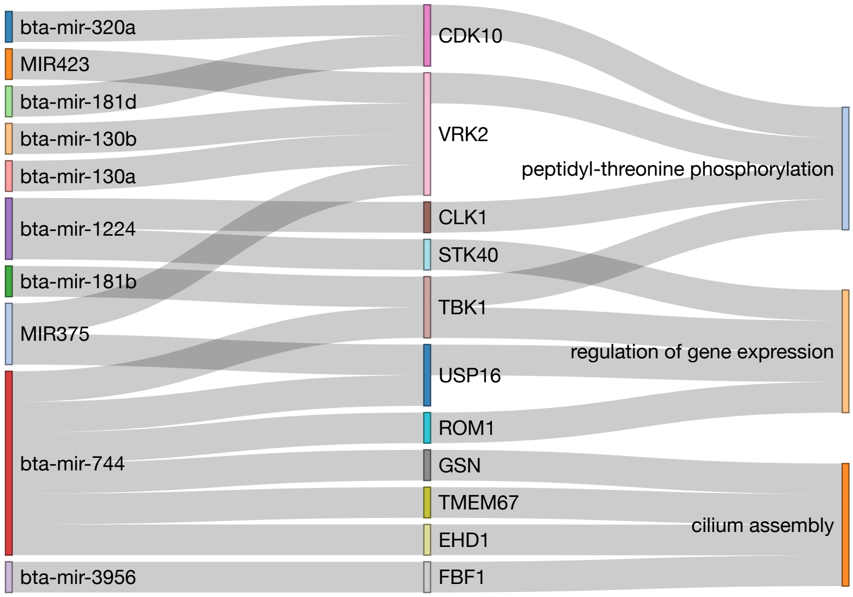

```
correlation_all_animals_miRNA_c<-correlation_all_animals_miRNA_b[correlation_all_animals_miRNA_b$Var1 %in% GO_BP_
Cats_mRNA_miRNA_all_animals$ensembl_gene_id[GO_BP_Cats_mRNA_miRNA_all_animals$term=='regulation of gene expressio
n'],]

data_for_chart<-data.frame()
data_for_chart_a<-data.frame()
for(i in 1:dim(correlation_all_animals_miRNA_c)[1]){
  expression_mRNA<-cpm_pwbc[as.character(correlation_all_animals_miRNA_c[i,1]),]
  expresson_miRNA<-cpm_miRNA[as.character(correlation_all_animals_miRNA_c[i,2]),]
  symbol_mRNA<-correlation_all_animals_miRNA_c[i,5]
  symbol_miRNA<-correlation_all_animals_miRNA_c[i,6]
  chart<-i
  data_for_chart<-data.frame(symbol_mRNA,symbol_miRNA,expression_mRNA,expresson_miRNA,chart)
  data_for_chart_a<-rbind(data_for_chart_a,data_for_chart)
}

data_for_chart_a[is.na(data_for_chart_a)]<-0

plots<-list()
k<-1
for (j in c(1:6)){
data_for_chart_b<-data_for_chart_a[data_for_chart_a$chart %in% j,]
plot<-ggplot(data_for_chart_b, aes(x=log(expresson_miRNA+1),y=log(expression_mRNA+1)))+
  geom_point()+
  geom_smooth(method=lm, size=0.1, fill="lightgray")+
  scale_y_continuous(name=data_for_chart_b$symbol_mRNA[1])+
  scale_x_continuous(name=data_for_chart_b$symbol_miRNA[1])+
  theme(aspect.ratio = 1,
            panel.grid.major = element_blank(),
            panel.grid.minor = element_blank(),
            panel.background = element_blank(),
            plot.background  = element_blank(),
            axis.text.x = element_text( colour = 'black' ,size = 7, angle=90),
            axis.text.y = element_text( colour = 'black',size = 7),
            axis.title= element_text( colour = 'black' ,size = 7, face="italic"),
            axis.ticks = element_line(size=0.1),
            panel.spacing = unit(0, "mm"),
            plot.margin = ggplot2::margin(1,1,1,1,unit= "mm"),
            legend.position="none",
            axis.line=element_line(size = 0.1, colour = "black"))

plots[[k]]<-plot
k<-k+1
}
#png(filename =paste(result_path,"2019_12_20_plot_mrna_mirna_all_samples_regulation_gene_expression.png", sep
="/"), width = 3, height = 2, units = 'in', res = 300)
#ggarrange(plotlist=plots, ncol=3, nrow=2)
#dev.off()
ggarrange(plotlist=plots, ncol=3, nrow=2)
```

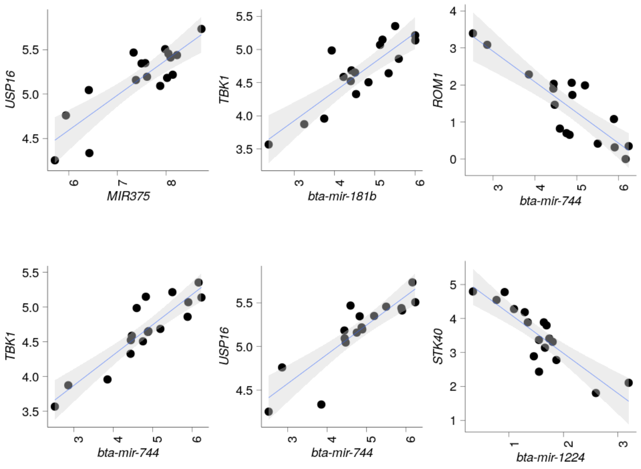

```
correlation_all_animals_miRNA_c<-correlation_all_animals_miRNA_b[correlation_all_animals_miRNA_b$Var1 %in% GO_BP_
Cats_mRNA_miRNA_all_animals$ensembl_gene_id[GO_BP_Cats_mRNA_miRNA_all_animals$term=='peptidyl-threonine phosphory
lation'],]

data_for_chart<-data.frame()
data_for_chart_a<-data.frame()
for(i in 1:dim(correlation_all_animals_miRNA_c)[1]){
  expression_mRNA<-cpm_pwbc[as.character(correlation_all_animals_miRNA_c[i,1]),]
  expresson_miRNA<-cpm_miRNA[as.character(correlation_all_animals_miRNA_c[i,2]),]
  symbol_mRNA<-correlation_all_animals_miRNA_c[i,5]
  symbol_miRNA<-correlation_all_animals_miRNA_c[i,6]
  chart<-i
  data_for_chart<-data.frame(symbol_mRNA,symbol_miRNA,expression_mRNA,expresson_miRNA,chart)
  data_for_chart_a<-rbind(data_for_chart_a,data_for_chart)
}

data_for_chart_a[is.na(data_for_chart_a)]<-0

plots<-list()
k<-1
for (j in c(1:9)){
data_for_chart_b<-data_for_chart_a[data_for_chart_a$chart %in% j,]
plot<-ggplot(data_for_chart_b, aes(x=log(expresson_miRNA+1),y=log(expression_mRNA+1)))+
  geom_point()+
  geom_smooth(method=lm, size=0.1, fill="lightgray")+
  scale_y_continuous(name=data_for_chart_b$symbol_mRNA[1])+
  scale_x_continuous(name=data_for_chart_b$symbol_miRNA[1])+
  theme(aspect.ratio = 1,
            panel.grid.major = element_blank(),
            panel.grid.minor = element_blank(),
            panel.background = element_blank(),
            plot.background  = element_blank(),
            axis.text.x = element_text( colour = 'black' ,size = 7, angle=90),
            axis.text.y = element_text( colour = 'black',size = 7),
            axis.title= element_text( colour = 'black' ,size = 7, face="italic"),
            axis.ticks = element_line(size=0.1),
            panel.spacing = unit(0, "mm"),
            plot.margin = ggplot2::margin(1,1,1,1,unit= "mm"),
            legend.position="none",
            axis.line=element_line(size = 0.1, colour = "black"))

plots[[k]]<-plot
k<-k+1
}
#png(filename =paste(result_path,"2019_12_20_plot_mrna_mirna_all_samples_peptidyl_threonine_phosphorylation.png",
sep="/"), width = 3, height = 3, units = 'in', res = 300)
#ggarrange(plotlist=plots, ncol=3, nrow=3)
#dev.off()
ggarrange(plotlist=plots, ncol=3, nrow=3)
```

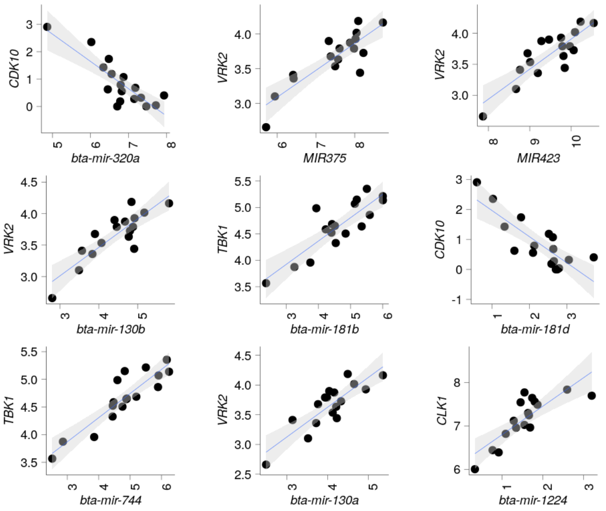


### Figure 3b

```
bta_miRWalk_3UTR$type<-"3UTR"
bta_miRWalk_5UTR$type<-"5UTR"
bta_miRWalk_CDS$type<-"CDS"

combined_bta_miRWalk<-rbind(bta_miRWalk_3UTR,rbind(bta_miRWalk_5UTR,bta_miRWalk_CDS))

combined_bta_miRWalk$miRNA_lower_case<-tolower(combined_bta_miRWalk$miRNA)
correlation_all_animals_miRNA_b$gene_2_lower_case<-tolower(correlation_all_animals_miRNA_b$gene_2)
correlation_all_animals_miRNA_c<-merge(correlation_all_animals_miRNA_b, combined_bta_miRWalk, by.x=c("gene_2_lowe
r_case", "gene_1" ), by.y=c("miRNA_lower_case", "Genesymbol" ), all=FALSE)
correlation_all_animals_miRNA_c<-correlation_all_animals_miRNA_c[!duplicated(correlation_all_animals_miRNA_c[,c(
"gene_2_lower_case", "gene_1")]),]
correlation_all_animals_miRNA_c<-correlation_all_animals_miRNA_c[correlation_all_animals_miRNA_c$value < -0.85,]
correlation_all_animals_miRNA_c<-correlation_all_animals_miRNA_c[order(correlation_all_animals_miRNA_c$gene_2_low
er_case, correlation_all_animals_miRNA_c$gene_1),]

#write.table(correlation_all_animals_miRNA_c, file= paste(result_path,"2019_12_24_mRNA_miNRA_correlation_all_anim
als_mirwalk.txt", sep="/"), append = FALSE, quote = FALSE, sep = "\t" ,row.names = FALSE)
#system("bzip2 -9 /mnt/storage/auburn/heifer_pregnancy/proj_2018/analysis/results/2019_12_24_mRNA_miNRA_correlati
on_all_animals_mirwalk.txt")


data_for_chart<-data.frame()
data_for_chart_a<-data.frame()
for(i in 1:dim(correlation_all_animals_miRNA_c)[1]){
  expression_mRNA<-cpm_pwbc[as.character(correlation_all_animals_miRNA_c[i,3]),]
  expresson_miRNA<-cpm_miRNA[as.character(correlation_all_animals_miRNA_c[i,4]),]
  symbol_mRNA<-correlation_all_animals_miRNA_c[i,2]
  symbol_miRNA<-correlation_all_animals_miRNA_c[i,1]
  chart<-i
  data_for_chart<-data.frame(symbol_mRNA,symbol_miRNA,expression_mRNA,expresson_miRNA,chart)
  data_for_chart_a<-rbind(data_for_chart_a,data_for_chart)
}

data_for_chart_a[is.na(data_for_chart_a)]<-0

plots<-list()
k<-1
for (j in c(1:30)){
data_for_chart_b<-data_for_chart_a[data_for_chart_a$chart %in% j,]
plot<-ggplot(data_for_chart_b, aes(x=log(expresson_miRNA+1),y=log(expression_mRNA+1)))+
  geom_point()+
  geom_smooth(method=lm, size=0.1, fill="lightgray")+
  scale_y_continuous(name=data_for_chart_b$symbol_mRNA[1])+
  scale_x_continuous(name=data_for_chart_b$symbol_miRNA[1])+
  theme(aspect.ratio = 1,
            panel.grid.major = element_blank(),
            panel.grid.minor = element_blank(),
            panel.background = element_blank(),
            plot.background  = element_blank(),
            axis.text.x = element_text( colour = 'black' ,size = 7, angle=90),
            axis.text.y = element_text( colour = 'black',size = 7),
            axis.title= element_text( colour = 'black' ,size = 7, face="italic"),
            axis.ticks = element_line(size=0.1),
            panel.spacing = unit(0, "mm"),
            plot.margin = ggplot2::margin(1,1,1,1,unit= "mm"),
            legend.position="none",
            axis.line=element_line(size = 0.1, colour = "black"))

plots[[k]]<-plot
k<-k+1
}
#png(filename =paste(result_path,"2019_12_21_plot_mrna_mirna_all_samples_binding_sites.png", sep="/"), width = 1
2, height = 3, units = 'in', res = 300)
#ggarrange(plotlist=plots, ncol=10, nrow=3)
#dev.off()
ggarrange(plotlist=plots, ncol=10, nrow=3)
```

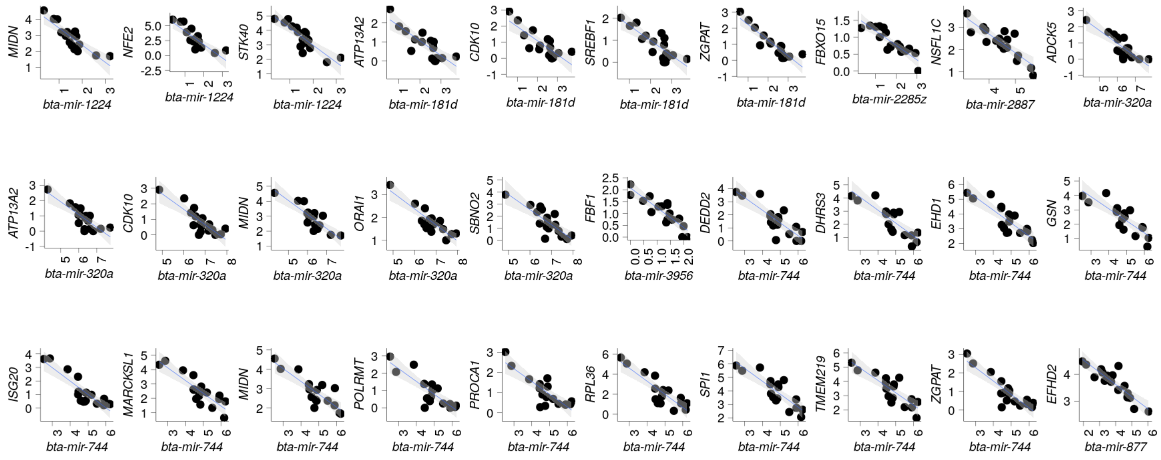


Create correlation matrices for AI pregnant NB pregnant and non pregnant animals.

```
rpkm_AIpreg_a<-t(log2(rpkm_AIpreg + 1))
rpkm_AIpreg_miRNA_a<-t(log2(rpkm_AIpreg_miRNA + 1))

rpkm_NBpreg_a<-t(log2(rpkm_NBpreg + 1))
rpkm_NBpreg_miRNA_a<-t(log2(rpkm_NBpreg_miRNA + 1))

rpkm_NOTpreg_a<-t(log2(rpkm_NOTpreg + 1))
rpkm_NOTpreg_miRNA_a<-t(log2(rpkm_NOTpreg_miRNA + 1))


correlation_AIpreg_mRNA_miRNA<-corAndPvalue(rpkm_AIpreg_a, rpkm_AIpreg_miRNA_a,method="pearson", use="pairwise.co
mplete.obs")
correlation_AIpreg_mRNA_miRNAa<-melt(correlation_AIpreg_mRNA_miRNA$cor)
correlation_AIpreg_mRNA_miRNAa$pvalue<-melt(correlation_AIpreg_mRNA_miRNA$p)$value
rm(correlation_AIpreg_mRNA_miRNA)
dim(correlation_AIpreg_mRNA_miRNAa)#3043840
```

```
## [1] 3043840       4
```

```
correlation_NBpreg_mRNA_miRNA<-corAndPvalue(rpkm_NBpreg_a, rpkm_NBpreg_miRNA_a,method="pearson", use="pairwise.co
mplete.obs")
correlation_NBpreg_mRNA_miRNAa<-melt(correlation_NBpreg_mRNA_miRNA$cor)
correlation_NBpreg_mRNA_miRNAa$pvalue<-melt(correlation_NBpreg_mRNA_miRNA$p)$value
rm(correlation_NBpreg_mRNA_miRNA)
dim(correlation_NBpreg_mRNA_miRNAa)#3043840
```

```
## [1] 3043840       4
```

```
correlation_NOTpreg_mRNA_miRNA<-corAndPvalue(rpkm_NOTpreg_a, rpkm_NOTpreg_miRNA_a,method="pearson", use="pairwis
e.complete.obs")
correlation_NOTpreg_mRNA_miRNAa<-melt(correlation_NOTpreg_mRNA_miRNA$cor)
correlation_NOTpreg_mRNA_miRNAa$pvalue<-melt(correlation_NOTpreg_mRNA_miRNA$p)$value
rm(correlation_NOTpreg_mRNA_miRNA)
dim(correlation_NOTpreg_mRNA_miRNAa) #3043840
```

```
## [1] 3043840       4
```

```
merged_correlation_mRNA_miRNA<-cbind(correlation_AIpreg_mRNA_miRNAa,cbind(correlation_NBpreg_mRNA_miRNAa,correlat
ion_NOTpreg_mRNA_miRNAa))
merged_correlation_mRNA_miRNA<-merged_correlation_mRNA_miRNA[,c(1,2,3,4,7,8,11,12)]
colnames(merged_correlation_mRNA_miRNA)<-c("mRNA_gene","miRNA_gene","cor_AI","pvalue_AI","cor_NB","pvalue_NB","co
r_NP","pvalue_NP")

merged_correlation_mRNA_miRNA$mRNA_symbol<-annotation.ensembl.symbol$external_gene_name[match( merged_correlation
_mRNA_miRNA$mRNA_gene, annotation.ensembl.symbol$ensembl_gene_id )]
merged_correlation_mRNA_miRNA$miRNA_symbol<-annotation.ensembl.symbol$external_gene_name[match( merged_correlatio
n_mRNA_miRNA$miRNA_gene, annotation.ensembl.symbol$ensembl_gene_id )]

merged_correlation_mRNA_miRNA$subtraction_cor_AI_cor_NP<-merged_correlation_mRNA_miRNA$cor_AI - merged_correlatio
n_mRNA_miRNA$cor_NP
merged_correlation_mRNA_miRNA$subtraction_cor_NB_cor_NP<-merged_correlation_mRNA_miRNA$cor_NB - merged_correlatio
n_mRNA_miRNA$cor_NP
merged_correlation_mRNA_miRNA$subtraction_cor_AI_cor_NB<-merged_correlation_mRNA_miRNA$cor_AI - merged_correlatio
n_mRNA_miRNA$cor_NB
```

#### Supplementary Fig. 3

```
setwd("/mnt/storage/auburn/heifer_pregnancy/proj_2018/analysis/results")

permutation_matrix<-matrix(,nrow=200100,ncol=17)
#head(permutation_matrix)
#dim(permutation_matrix)

for (i in seq(1:200100)){
  sampling <- sample(1:17,17, replace = FALSE)
  if ( ! identical(sampling , c(1:17))){
    permutation_matrix[i,]<-sample(1:17,17, replace = FALSE)
  }}

permutation_matrix<-permutation_matrix[!duplicated(permutation_matrix),]
dim(permutation_matrix)
head(permutation_matrix)

permutation_matrix<-permutation_matrix[sample(1:dim(permutation_matrix)[1], 5000),]
rand<-dim(permutation_matrix)[1]

sequence.pvalue<-seq(0.7, 1, 0.01)

results <- filebacked.big.matrix(length(sequence.pvalue),rand, type="double", init=0, separated=FALSE,
                                 backingfile="incidence_matrix.bin",
                                 descriptor="incidence_matrix.desc")  #this does not overwrite, create once and d
elete if want to create again
mdesc_result<- describe(results)

cl <- makeCluster(20)
registerDoParallel(cl)

results[,]<-foreach(i = sequence.pvalue, .combine='rbind', .inorder=TRUE,  .packages=c("bigmemory"), .verbose=FAL
SE) %:%
  foreach(j = 1:rand, .combine='cbind', .inorder=FALSE,.packages=c("reshape2","bigmemory"), .verbose=FALSE ) %dop
ar% {

   rpkm_pwbc_scramble<-log2(rpkm_pwbc[,permutation_matrix[j,]] +1)
   cor_1<-cor( t(rpkm_pwbc_scramble[,sample(17,6)]), t(log2(rpkm_miRNA +1)[,sample(17,6)]) ,method="pearson", use
="pairwise.complete.obs")
   length(which(abs(cor_1) > i))
  }

stopCluster(cl)

results[1:5,1:5]


total.rand <- rand * 10496 * 290

qvalue<-data.frame(correlation = sequence.pvalue,
                   e.pvalue= (rowSums(results[,]+1))/(total.rand+1),
                   e.pvalue.round= round((rowSums(results[,]+1))/(total.rand+1) ,6))

write.table(qvalue, file= paste(result_path,"/2019_12_23_mRNA_miRNA_within_group_coexpression_empiricalFDR.txt",
 sep=""), append = FALSE, quote = FALSE, sep = "\t" ,row.names = FALSE)

qvalue_two_tailled<-data.frame(correlation = sequence.pvalue,
                   e.pvalue= ( (rowSums(results[,]+1))/(total.rand+1) )/2  ,
                   e.pvalue.round= ( round((rowSums(results[,]+1))/(total.rand+1) ,6) )/2 )

write.table(qvalue, file= paste(result_path,"/2019_12_23_mRNA_miRNA_within_group_coexpression_empiricalFDR_twotai
l.txt", sep=""), append = FALSE, quote = FALSE, sep = "\t" ,row.names = FALSE)


rm(results)
system("rm incidence_matrix.bin")
system("rm incidence_matrix.desc")
```

```
eFDR_coexpression<-read.delim(paste(result_path,"2019_12_23_mRNA_miRNA_within_group_coexpression_empiricalFDR_two
tail.txt", sep="/"), row.names=NULL,header =TRUE, stringsAsFactors =FALSE)
```

```
font_size<-12 #font_size 12 for pdf  figure

scaleFUN <- function(x) sprintf("%.2f", x)

plot1<-ggplot()+
  geom_point(data=eFDR_coexpression, aes(x=correlation , y=e.pvalue.round),color="black", size=3, shape=16)+
  geom_line(data=eFDR_coexpression, aes(x=correlation , y=e.pvalue.round),color="black", size=0.1,linetype=3)+
  scale_y_continuous(name="empirical FDR", limits = c(0, 0.15), breaks=seq(0,0.2, 0.01))+
  scale_x_continuous(name="correlation (r)", limits = c(0.7, 1), breaks=seq(0.7,1, 0.01), labels=scaleFUN)+
  ggtitle("mRNA and miRNA coexpression within a group of heifers")+
  theme_bw()+
  theme(panel.grid= element_blank(),
        panel.background = element_blank(),
        panel.grid.minor = element_blank(),
        panel.grid.major = element_line(color="lightgray"),
        plot.background = element_blank(),
        axis.title=element_text(color="black", size=font_size),
        axis.text.y=element_text(color="black", size=font_size),
        axis.text.x=element_text(color="black", size=font_size, angle=90),
        panel.spacing = unit(c(0.4,0.4,0.4,0.4),"cm"),
        plot.margin = unit(c(0.5,0.5,0.5,0.5),"cm"),
        legend.position="none",
        plot.title = element_text(lineheight=.8, hjust=0.5, size= font_size))


#font_size 12 for figure
#pdf(paste(result_path,"2020_01_29_mRNA_miRNA_within_group_eFDR_plots.pdf", sep="/"), width=9, height=5)
#plot1
#dev.off()
plot1
```

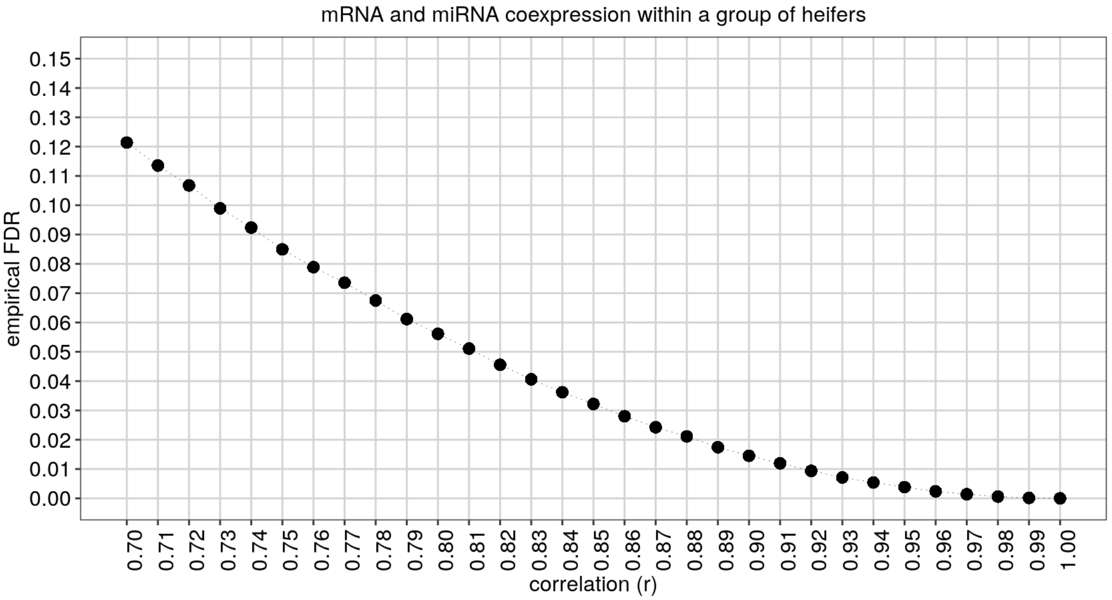


### Supplementary Table 8

```
merged_correlation_mRNA_miRNA$result_cor_AI_and_NB_cor_NP_a<-ifelse( (abs(merged_correlation_mRNA_miRNA$subtraction_cor_AI_cor_NP) + abs(merged_correlation_mRNA_miRNA$subtraction_cor_NB_cor_NP)) / 2 > 1.8 , "inverted",
                                                                      ifelse( (merged_correlation_mRNA_miRNA$cor_AI >0.90  & merged_correlation_mRNA_miRNA$cor_NB >0.90)  & (merged_correlation_mRNA_miRNA$cor_NP > -0.1 & merged_correlation_mRNA_miRNA$cor_NP < 0.1) , "loss_positive",
                                   ifelse( (merged_correlation_mRNA_miRNA$cor_AI < -0.90  & merged_correlation_mRNA_miRNA$cor_NB < -0.90)  & (merged_correlation_mRNA_miRNA$cor_NP > -0.1 & merged_correlation_mRNA_miRNA$cor_NP < 0.1) , "loss_negative",
                                   ifelse( (merged_correlation_mRNA_miRNA$cor_AI > -0.1 & merged_correlation_mRNA_miRNA$cor_AI < 0.1 & merged_correlation_mRNA_miRNA$cor_NB > -0.1 & merged_correlation_mRNA_miRNA$cor_NB < 0.1 ) &  merged_correlation_mRNA_miRNA$cor_NP > 0.90, "gain_positive" ,
                                   ifelse( (merged_correlation_mRNA_miRNA$cor_AI > -0.1 & merged_correlation_mRNA_miRNA$cor_AI < 0.1 & merged_correlation_mRNA_miRNA$cor_NB > -0.1 & merged_correlation_mRNA_miRNA$cor_NB < 0.1 ) &  merged_correlation_mRNA_miRNA$cor_NP < -0.90, "gain_negative" ,"nothing")))))

table(merged_correlation_mRNA_miRNA$result_cor_AI_and_NB_cor_NP_a)
```

```
##
## gain_negative gain_positive      inverted loss_negative loss_positive
##          2330          2738            51             2             3
##       nothing
##       3038716
```

```
#head(merged_correlation_mRNA_miRNA)

merged_correlation_mRNA_miRNA<-merged_correlation_mRNA_miRNA[!(merged_correlation_mRNA_miRNA$result_cor_AI_and_NB_cor_NP_a== "nothing"),]

#write.table(merged_correlation_mRNA_miRNA, file= paste(result_path,"/2020_01_29_mRNA_miRNA_sig_differential_coexpression.txt", sep=""), append = FALSE, quote = FALSE, sep = "\t" ,row.names = FALSE)


merged_correlation_mRNA_miRNA_gain_positive<-merged_correlation_mRNA_miRNA[(merged_correlation_mRNA_miRNA$result_cor_AI_and_NB_cor_NP_a== "gain_positive"),]
merged_correlation_mRNA_miRNA_gain_negative<-merged_correlation_mRNA_miRNA[(merged_correlation_mRNA_miRNA$result_cor_AI_and_NB_cor_NP_a== "gain_negative"),]
merged_correlation_mRNA_miRNA_inverted<-merged_correlation_mRNA_miRNA[(merged_correlation_mRNA_miRNA$result_cor_AI_and_NB_cor_NP_a== "inverted"),]
```

```
knitr::kable(head(merged_correlation_mRNA_miRNA), caption = "First five rows of Supplementary Table 8",format = 'pandoc')
```

First five rows of Supplementary Table 8

| | mRNA_gene | miRNA_gene | cor_AI | pvalue_AI | cor_NB | pvalue_NB | cor_NP | pvalue_NP | mRNA |
|---|---|---|---|---|---|---|---|---|---|
| 6583 | ENSBTAG00000017803 | ENSBTAG00000029760 | -0.0688122 | 0.8969446 | -0.0915422 | 0.8630703 | 0.9104692 | 0.0317229 | RAB1 |
| 10880 | ENSBTAG00000000993 | ENSBTAG00000029761 | -0.0444719 | 0.9333361 | -0.0853261 | 0.8723215 | 0.9038630 | 0.0352619 | N4BP |
| 14977 | ENSBTAG00000011994 | ENSBTAG00000029761 | 0.0547957 | 0.9178887 | 0.0724871 | 0.8914598 | -0.9244153 | 0.0246603 | NDEL |
| 15320 | ENSBTAG00000012972 | ENSBTAG00000029761 | 0.0031738 | 0.9952393 | 0.0484562 | 0.9273727 | 0.9230475 | 0.0253274 | CYP2 |
| 15513 | ENSBTAG00000013475 | ENSBTAG00000029761 | 0.0925901 | 0.8615118 | 0.0820110 | 0.8772593 | 0.9289427 | 0.0224938 | TRAF |
| 18569 | ENSBTAG00000021859 | ENSBTAG00000029761 | -0.0431445 | 0.9353233 | 0.0254350 | 0.9618558 | 0.9867504 | 0.0018271 | PDSS |

### Figure 3c

```
plot<-ggplot(merged_correlation_mRNA_miRNA, aes(y=(cor_AI+ cor_NB)/2, x=cor_NP))+
geom_point(alpha=0.2, size=1)+
scale_x_continuous("non pregnant")+
scale_y_continuous("Pregnant")+
coord_fixed(ratio = 1)+
#ggtitle("mRNA-miRNA differential coexpression")+
theme_bw()+
theme(
text = element_text(size=10, color="black"),
panel.background = element_rect(fill = "transparent"),
plot.background = element_rect(fill = "transparent", color = NA),
)
#png(filename =paste(result_path,"2019_12_23_plot_mrna_mirna_diff_coexpression.png", sep="/"), width = 2, height
 = 2, units = 'in', res = 300, bg = "transparent")
#plot
#dev.off()
plot
```

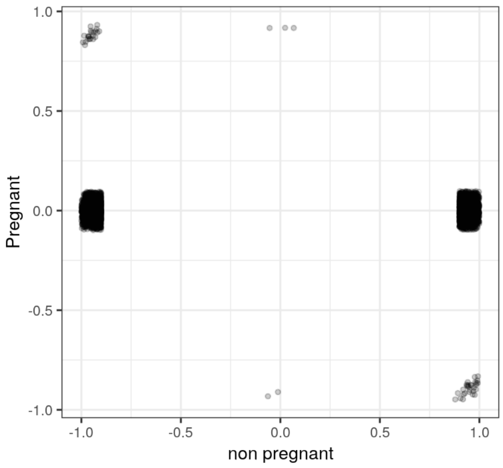

```
merged_correlation_mRNA_miRNA_gain_positive<-merged_correlation_mRNA_miRNA[(merged_correlation_mRNA_miRNA$result_
cor_AI_and_NB_cor_NP_a== "gain_positive"),]
merged_correlation_mRNA_miRNA_gain_positive<-merged_correlation_mRNA_miRNA_gain_positive[order(-merged_correlatio
n_mRNA_miRNA_gain_positive$cor_NP),]


data_for_chart<-data.frame()
i=1
  expression_gene_1_AI<-cpm_AIpreg[row.names(cpm_AIpreg)==as.character(merged_correlation_mRNA_miRNA_gain_positiv
e$mRNA_gene)[i],]
  expression_gene_2_AI<-cpm_AIpreg_miRNA[row.names(cpm_AIpreg_miRNA)==as.character(merged_correlation_mRNA_miRNA_
gain_positive$miRNA_gene)[i],]
  expression_gene_1_NB<-cpm_NBpreg[row.names(cpm_NBpreg)==as.character(merged_correlation_mRNA_miRNA_gain_positiv
e$mRNA_gene)[i],]
  expression_gene_2_NB<-cpm_NBpreg_miRNA[row.names(cpm_NBpreg_miRNA)==as.character(merged_correlation_mRNA_miRNA_
gain_positive$miRNA_gene)[i],]
  expression_gene_1_NP<-cpm_NOTpreg[row.names(cpm_NOTpreg)==as.character(merged_correlation_mRNA_miRNA_gain_posit
ive$mRNA_gene)[i],]
  expression_gene_2_NP<-cpm_NOTpreg_miRNA[row.names(cpm_NOTpreg_miRNA)==as.character(merged_correlation_mRNA_miRN
A_gain_positive$miRNA_gene)[i],]

  symbol_gene_1<-annotation.ensembl.symbol[annotation.ensembl.symbol$ensembl_gene_id == as.character(merged_corre
lation_mRNA_miRNA_gain_positive$mRNA_gene)[i] , 2]
  symbol_gene_2<-annotation.ensembl.symbol[annotation.ensembl.symbol$ensembl_gene_id == as.character(merged_corre
lation_mRNA_miRNA_gain_positive$miRNA_gene)[i] , 2]

  data_for_chart<-data.frame(gene_1=c(expression_gene_1_AI,expression_gene_1_NB,expression_gene_1_NP) ,
                             gene_2=c(expression_gene_2_AI,expression_gene_2_NB,expression_gene_2_NP),
                             symbol_gene_1,symbol_gene_2 ,chart,
                             group=rep(c("AI", "NB", "NP"), c(length(expression_gene_1_AI),length(expression_gene
_1_NB),length(expression_gene_1_NP))))

font_axis<-5
font_title<-8
ptsize<-0.05

plot<-ggplot(data_for_chart, aes(x=log(gene_1+1),y=log(gene_2+1)))+
  geom_point()+
  geom_smooth(method=lm, size=ptsize, fill="lightgray")+
  scale_y_continuous(name=data_for_chart$symbol_gene_1[1])+
  scale_x_continuous(name=data_for_chart$symbol_gene_2[1])+
  facet_wrap(~group, ncol=3, scales="free")+
  theme(aspect.ratio = 1,
            panel.grid.major = element_blank(),
            panel.grid.minor = element_blank(),
            panel.background = element_blank(),
            plot.background  = element_blank(),
            axis.text.y = element_text( colour = 'black',size = font_axis),
            axis.text.x = element_text( colour = 'black',size = font_axis, angle=90),
            axis.title = element_text( colour = 'black' ,size = font_title, face="italic"),
            axis.ticks = element_line(size=0.1),
            panel.spacing.y = unit(0, "mm"),
            panel.spacing.x = unit(2, "mm"),
            legend.position="none",
            axis.line=element_line(size = 0.1, colour = "black"),
            strip.background=element_blank(),
            strip.text.x = element_blank())

#png(filename =paste(result_path,"2019_12_23_plot_diff_cor_mRNA_miRNA_AI_NB_and_NP_gain_positive.png", sep="/"),
 width = 2.5, height =1, units = 'in', res = 300,bg = "transparent")
#plot
#dev.off()
plot
```

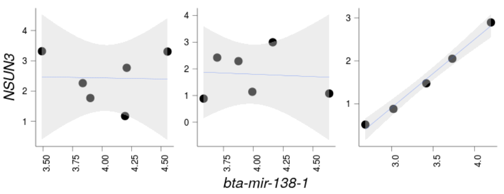

```
merged_correlation_mRNA_miRNA_gain_negative<-merged_correlation_mRNA_miRNA[(merged_correlation_mRNA_miRNA$result_
cor_AI_and_NB_cor_NP_a== "gain_negative"),]
merged_correlation_mRNA_miRNA_gain_negative<-merged_correlation_mRNA_miRNA_gain_negative[order(merged_correlation
_mRNA_miRNA_gain_negative$cor_NP),]


data_for_chart<-data.frame()
i=1
  expression_gene_1_AI<-cpm_AIpreg[row.names(cpm_AIpreg)==as.character(merged_correlation_mRNA_miRNA_gain_negativ
e$mRNA_gene)[i],]
  expression_gene_2_AI<-cpm_AIpreg_miRNA[row.names(cpm_AIpreg_miRNA)==as.character(merged_correlation_mRNA_miRNA_
gain_negative$miRNA_gene)[i],]
  expression_gene_1_NB<-cpm_NBpreg[row.names(cpm_NBpreg)==as.character(merged_correlation_mRNA_miRNA_gain_negativ
e$mRNA_gene)[i],]
  expression_gene_2_NB<-cpm_NBpreg_miRNA[row.names(cpm_NBpreg_miRNA)==as.character(merged_correlation_mRNA_miRNA_
gain_negative$miRNA_gene)[i],]
  expression_gene_1_NP<-cpm_NOTpreg[row.names(cpm_NOTpreg)==as.character(merged_correlation_mRNA_miRNA_gain_negat
ive$mRNA_gene)[i],]
  expression_gene_2_NP<-cpm_NOTpreg_miRNA[row.names(cpm_NOTpreg_miRNA)==as.character(merged_correlation_mRNA_miRN
A_gain_negative$miRNA_gene)[i],]

  symbol_gene_1<-annotation.ensembl.symbol[annotation.ensembl.symbol$ensembl_gene_id == as.character(merged_corre
lation_mRNA_miRNA_gain_negative$mRNA_gene)[i] , 2]
  symbol_gene_2<-annotation.ensembl.symbol[annotation.ensembl.symbol$ensembl_gene_id == as.character(merged_corre
lation_mRNA_miRNA_gain_negative$miRNA_gene)[i] , 2]

  data_for_chart<-data.frame(gene_1=c(expression_gene_1_AI,expression_gene_1_NB,expression_gene_1_NP) ,
                             gene_2=c(expression_gene_2_AI,expression_gene_2_NB,expression_gene_2_NP),
                             symbol_gene_1,symbol_gene_2 ,chart,
                             group=rep(c("AI", "NB", "NP"), c(length(expression_gene_1_AI),length(expression_gene
_1_NB),length(expression_gene_1_NP))))

font_axis<-5
font_title<-8
ptsize<-0.05

plot<-ggplot(data_for_chart, aes(x=log(gene_1+1),y=log(gene_2+1)))+
  geom_point()+
  geom_smooth(method=lm, size=ptsize, fill="lightgray")+
  scale_y_continuous(name=data_for_chart$symbol_gene_1[1])+
  scale_x_continuous(name=data_for_chart$symbol_gene_2[1])+
  facet_wrap(~group, ncol=3, scales="free")+
  theme(aspect.ratio = 1,
          panel.grid.major = element_blank(),
          panel.grid.minor = element_blank(),
          panel.background = element_blank(),
          plot.background  = element_blank(),
          axis.text.y = element_text( colour = 'black',size = font_axis),
          axis.text.x = element_text( colour = 'black',size = font_axis, angle=90),
          axis.title = element_text( colour = 'black' ,size = font_title, face="italic"),
          axis.ticks = element_line(size=0.1),
          panel.spacing.y = unit(0, "mm"),
          panel.spacing.x = unit(2, "mm"),
          legend.position="none",
          axis.line=element_line(size = 0.1, colour = "black"),
          strip.background=element_blank(),
          strip.text.x = element_blank())

#png(filename =paste(result_path,"2019_12_23_plot_diff_cor_mRNA_miRNA_AI_NB_and_NP_gain_negative.png", sep="/"),
 width = 2.5, height =1, units = 'in', res = 300,bg = "transparent")
#plot
#dev.off()
plot
```

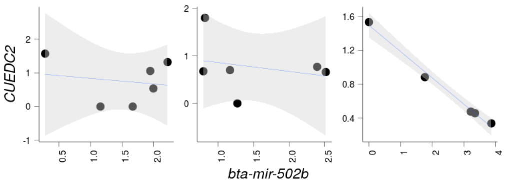

```
merged_correlation_mRNA_miRNA_inverted<-merged_correlation_mRNA_miRNA[(merged_correlation_mRNA_miRNA$result_cor_A
I_and_NB_cor_NP_a== "inverted"),]
merged_correlation_mRNA_miRNA_inverted<-merged_correlation_mRNA_miRNA_inverted[order(merged_correlation_mRNA_miRN
A_inverted$cor_NP),]


data_for_chart<-data.frame()
i=1
  expression_gene_1_AI<-cpm_AIpreg[row.names(cpm_AIpreg)==as.character(merged_correlation_mRNA_miRNA_inverted$mRN
A_gene)[i],]
  expression_gene_2_AI<-cpm_AIpreg_miRNA[row.names(cpm_AIpreg_miRNA)==as.character(merged_correlation_mRNA_miRNA_
inverted$miRNA_gene)[i],]
  expression_gene_1_NB<-cpm_NBpreg[row.names(cpm_NBpreg)==as.character(merged_correlation_mRNA_miRNA_inverted$mRN
A_gene)[i],]
  expression_gene_2_NB<-cpm_NBpreg_miRNA[row.names(cpm_NBpreg_miRNA)==as.character(merged_correlation_mRNA_miRNA_
inverted$miRNA_gene)[i],]
  expression_gene_1_NP<-cpm_NOTpreg[row.names(cpm_NOTpreg)==as.character(merged_correlation_mRNA_miRNA_inverted$m
RNA_gene)[i],]
  expression_gene_2_NP<-cpm_NOTpreg_miRNA[row.names(cpm_NOTpreg_miRNA)==as.character(merged_correlation_mRNA_miRN
A_inverted$miRNA_gene)[i],]

  symbol_gene_1<-annotation.ensembl.symbol[annotation.ensembl.symbol$ensembl_gene_id == as.character(merged_corre
lation_mRNA_miRNA_inverted$mRNA_gene)[i] , 2]
  symbol_gene_2<-annotation.ensembl.symbol[annotation.ensembl.symbol$ensembl_gene_id == as.character(merged_corre
lation_mRNA_miRNA_inverted$miRNA_gene)[i] , 2]

  data_for_chart<-data.frame(gene_1=c(expression_gene_1_AI,expression_gene_1_NB,expression_gene_1_NP) ,
                             gene_2=c(expression_gene_2_AI,expression_gene_2_NB,expression_gene_2_NP),
                             symbol_gene_1,symbol_gene_2 ,chart,
                             group=rep(c("AI", "NB", "NP"), c(length(expression_gene_1_AI),length(expression_gene
_1_NB),length(expression_gene_1_NP))))

font_axis<-5
font_title<-8
ptsize<-0.05

plot<-ggplot(data_for_chart, aes(x=log(gene_1+1),y=log(gene_2+1)))+
  geom_point()+
  geom_smooth(method=lm, size=ptsize, fill="lightgray")+
  scale_y_continuous(name=data_for_chart$symbol_gene_1[1])+
  scale_x_continuous(name=data_for_chart$symbol_gene_2[1])+
  facet_wrap(~group, ncol=3, scales="free")+
  theme(aspect.ratio = 1,
          panel.grid.major = element_blank(),
          panel.grid.minor = element_blank(),
          panel.background = element_blank(),
          plot.background  = element_blank(),
          axis.text.y = element_text( colour = 'black',size = font_axis),
          axis.text.x = element_text( colour = 'black',size = font_axis, angle=90),
          axis.title = element_text( colour = 'black' ,size = font_title, face="italic"),
          axis.ticks = element_line(size=0.1),
          panel.spacing.y = unit(0, "mm"),
          panel.spacing.x = unit(2, "mm"),
          legend.position="none",
          axis.line=element_line(size = 0.1, colour = "black"),
          strip.background=element_blank(),
          strip.text.x = element_blank())

#png(filename =paste(result_path,"2019_12_23_plot_diff_cor_mRNA_miRNA_AI_NB_and_NP_inverted_negative.png", sep
="/"), width = 2.5, height =1, units = 'in', res = 300,bg = "transparent")
#plot
#dev.off()
plot
```

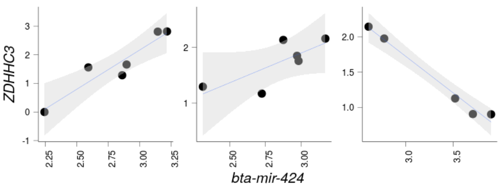

```
merged_correlation_mRNA_miRNA_inverted<-merged_correlation_mRNA_miRNA_inverted[order(-merged_correlation_mRNA_miRNA_inverted$cor_NP),]

data_for_chart<-data.frame()
i=5
  expression_gene_1_AI<-cpm_AIpreg[row.names(cpm_AIpreg)==as.character(merged_correlation_mRNA_miRNA_inverted$mRNA_gene)[i],]
  expression_gene_2_AI<-cpm_AIpreg_miRNA[row.names(cpm_AIpreg_miRNA)==as.character(merged_correlation_mRNA_miRNA_inverted$miRNA_gene)[i],]
  expression_gene_1_NB<-cpm_NBpreg[row.names(cpm_NBpreg)==as.character(merged_correlation_mRNA_miRNA_inverted$mRNA_gene)[i],]
  expression_gene_2_NB<-cpm_NBpreg_miRNA[row.names(cpm_NBpreg_miRNA)==as.character(merged_correlation_mRNA_miRNA_inverted$miRNA_gene)[i],]
  expression_gene_1_NP<-cpm_NOTpreg[row.names(cpm_NOTpreg)==as.character(merged_correlation_mRNA_miRNA_inverted$mRNA_gene)[i],]
  expression_gene_2_NP<-cpm_NOTpreg_miRNA[row.names(cpm_NOTpreg_miRNA)==as.character(merged_correlation_mRNA_miRNA_inverted$miRNA_gene)[i],]

  symbol_gene_1<-annotation.ensembl.symbol[annotation.ensembl.symbol$ensembl_gene_id == as.character(merged_correlation_mRNA_miRNA_inverted$mRNA_gene)[i] , 2]
  symbol_gene_2<-annotation.ensembl.symbol[annotation.ensembl.symbol$ensembl_gene_id == as.character(merged_correlation_mRNA_miRNA_inverted$miRNA_gene)[i] , 2]

  data_for_chart<-data.frame(gene_1=c(expression_gene_1_AI,expression_gene_1_NB,expression_gene_1_NP) ,
                             gene_2=c(expression_gene_2_AI,expression_gene_2_NB,expression_gene_2_NP),
                             symbol_gene_1,symbol_gene_2 ,chart,
                             group=rep(c("AI", "NB", "NP"), c(length(expression_gene_1_AI),length(expression_gene_1_NB),length(expression_gene_1_NP))))

font_axis<-5
font_title<-8
ptsize<-0.05

plot<-ggplot(data_for_chart, aes(x=log(gene_1+1),y=log(gene_2+1)))+
  geom_point()+
  geom_smooth(method=lm, size=ptsize, fill="lightgray")+
  scale_y_continuous(name=data_for_chart$symbol_gene_1[1])+
  scale_x_continuous(name=data_for_chart$symbol_gene_2[1])+
  facet_wrap(~group, ncol=3, scales="free")+
  theme(aspect.ratio = 1,
            panel.grid.major = element_blank(),
            panel.grid.minor = element_blank(),
            panel.background = element_blank(),
            plot.background  = element_blank(),
            axis.text.y = element_text( colour = 'black',size = font_axis),
            axis.text.x = element_text( colour = 'black',size = font_axis, angle=90),
            axis.title = element_text( colour = 'black' ,size = font_title, face="italic"),
            axis.ticks = element_line(size=0.1),
            panel.spacing.y = unit(0, "mm"),
            panel.spacing.x = unit(2, "mm"),
            legend.position="none",
            axis.line=element_line(size = 0.1, colour = "black"),
            strip.background=element_blank(),
            strip.text.x = element_blank())

#png(filename =paste(result_path,"2019_12_23_plot_diff_cor_mRNA_miRNA_AI_NB_and_NP_inverted_positive.png", sep="/"), width = 2.5, height =1, units = 'in', res = 300,bg = "transparent")
#plot
#dev.off()
plot
```

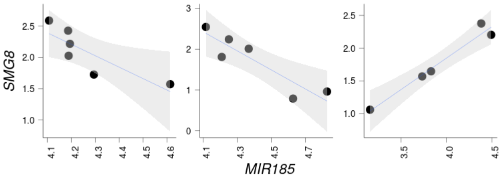


### Supplementary Table 9

```
test.genes<-data.frame(a=unique(c(as.character(merged_correlation_mRNA_miRNA_gain_negative$mRNA_gene))), stringsA
sFactors=FALSE)
all_genes_numeric<-as.integer(all_genes$gene %in%test.genes$a)
names(all_genes_numeric)<-all_genes$gene

N_DEGs<-length(test.genes$a)

set.seed(9830)
pwf<-nullp(all_genes_numeric, bias.data=annotation.genelength.biomart_vector, plot.fit=FALSE )
GO_BP_Cats_mRNA_AI_NB_Preg_NP<-goseq(pwf,gene2cat=annotation.GO.BP.biomart, method ="Sampling", repcnt = 7000, us
e_genes_without_cat=FALSE)
GO_BP_Cats_mRNA_AI_NB_Preg_NP<-GO_BP_Cats_mRNA_AI_NB_Preg_NP[GO_BP_Cats_mRNA_AI_NB_Preg_NP$numDEInCat>4,]
GO_BP_Cats_mRNA_AI_NB_Preg_NP$BY_FDR<-p.adjust(GO_BP_Cats_mRNA_AI_NB_Preg_NP$over_represented_pvalue, method ="fd
r")
GO_BP_Cats_mRNA_AI_NB_Preg_NP<-GO_BP_Cats_mRNA_AI_NB_Preg_NP[with(GO_BP_Cats_mRNA_AI_NB_Preg_NP, order(BY_FDR,ove
r_represented_pvalue, -numDEInCat)), ]
#head(GO_BP_Cats_mRNA_AI_NB_Preg_NP, n=20)

GO_BP_Cats_mRNA_AI_NB_Preg_NP$fold_enrichment<-(GO_BP_Cats_mRNA_AI_NB_Preg_NP$numDEInCat/N_DEGs)/(GO_BP_Cats_mRNA
_AI_NB_Preg_NP$numInCat/N_expressed_genes)
annotation.GO.BP.biomart_testgenes<-annotation.GO.BP.biomart[annotation.GO.BP.biomart$ensembl_gene_id %in% test.g
enes$a, ]
GO_BP_Cats_mRNA_AI_NB_Preg_NP<-merge(GO_BP_Cats_mRNA_AI_NB_Preg_NP,annotation.GO.BP.biomart_testgenes, by.x="cate
gory", by.y="go_id", all.x=TRUE, all.y=FALSE)
GO_BP_Cats_mRNA_AI_NB_Preg_NP<-merge(GO_BP_Cats_mRNA_AI_NB_Preg_NP, annotation.ensembl.symbol, by.x="ensembl_gene
_id", by.y="ensembl_gene_id", all=FALSE, all.x=TRUE, all.y=FALSE)
GO_BP_Cats_mRNA_AI_NB_Preg_NP<-GO_BP_Cats_mRNA_AI_NB_Preg_NP[with(GO_BP_Cats_mRNA_AI_NB_Preg_NP, order(BY_FDR,ter
m)), ]
GO_BP_Cats_mRNA_AI_NB_Preg_NP<-GO_BP_Cats_mRNA_AI_NB_Preg_NP[GO_BP_Cats_mRNA_AI_NB_Preg_NP$BY_FDR < 0.1,]

#write.table(GO_BP_Cats_mRNA_AI_NB_Preg_NP, paste(result_path, "2019_12_23_GO_BP_Cats_mRNA_miRNA_gain_neg_coexpre
ssion_AI_NB_Preg_NP.txt", sep="/"), quote=FALSE, sep="\t", row.names=FALSE)
```

```
knitr::kable(head(GO_BP_Cats_mRNA_AI_NB_Preg_NP), caption = "First five rows of Supplementary Table 9",format =
'pandoc')
```

First five rows of Supplementary Table 9

| | ensembl_gene_id | category | over_represented_pvalue | under_represented_pvalue | numDEInCat | numInCat | term | ontology |
|---|---|---|---|---|---|---|---|---|
| 55 | ENSBTAG00000000563 | GO:0016064 (GO:0016064) | 0.0005713 | 1.0000000 | 5 | 8 | immunoglobulin mediated immune response | BP |
| 1343 | ENSBTAG00000015228 | GO:0016064 (GO:0016064) | 0.0005713 | 1.0000000 | 5 | 8 | immunoglobulin mediated immune response | BP |
| 1979 | ENSBTAG00000024503 | GO:0016064 (GO:0016064) | 0.0005713 | 1.0000000 | 5 | 8 | immunoglobulin mediated immune response | BP |
| 2126 | ENSBTAG00000032122 | GO:0016064 (GO:0016064) | 0.0005713 | 1.0000000 | 5 | 8 | immunoglobulin mediated immune response | BP |
| 2289 | ENSBTAG00000047680 | GO:0016064 (GO:0016064) | 0.0005713 | 1.0000000 | 5 | 8 | immunoglobulin mediated immune response | BP |
| 37 | ENSBTAG00000000396 | GO:0043433 (GO:0043433) | 0.0004285 | 0.9998572 | 11 | 34 | negative regulation of DNA-binding transcription factor activity | BP |

### Supplementary Table 10

```
test.genes<-data.frame(a=unique(c(as.character(merged_correlation_mRNA_miRNA_gain_positive$mRNA_gene))), stringsA
sFactors=FALSE)
all_genes_numeric<-as.integer(all_genes$gene %in%test.genes$a)
names(all_genes_numeric)<-all_genes$gene

N_DEGs<-length(test.genes$a)

set.seed(9830)
pwf<-nullp(all_genes_numeric, bias.data=annotation.genelength.biomart_vector, plot.fit=FALSE )
```

```
## Warning in pcls(G): initial point very close to some inequality constraints
```

```
GO_BP_Cats_mRNA_AI_NB_Preg_NP<-goseq(pwf,gene2cat=annotation.GO.BP.biomart, method ="Sampling", repcnt = 7000, us
e_genes_without_cat=FALSE)
GO_BP_Cats_mRNA_AI_NB_Preg_NP<-GO_BP_Cats_mRNA_AI_NB_Preg_NP[GO_BP_Cats_mRNA_AI_NB_Preg_NP$numDEInCat>4,]
GO_BP_Cats_mRNA_AI_NB_Preg_NP$BY_FDR<-p.adjust(GO_BP_Cats_mRNA_AI_NB_Preg_NP$over_represented_pvalue, method ="fd
r")
GO_BP_Cats_mRNA_AI_NB_Preg_NP<-GO_BP_Cats_mRNA_AI_NB_Preg_NP[with(GO_BP_Cats_mRNA_AI_NB_Preg_NP, order(BY_FDR,ove
r_represented_pvalue, -numDEInCat)), ]
#head(GO_BP_Cats_mRNA_AI_NB_Preg_NP, n=20)

GO_BP_Cats_mRNA_AI_NB_Preg_NP$fold_enrichment<-(GO_BP_Cats_mRNA_AI_NB_Preg_NP$numDEInCat/N_DEGs)/(GO_BP_Cats_mRNA
_AI_NB_Preg_NP$numInCat/N_expressed_genes)
annotation.GO.BP.biomart_testgenes<-annotation.GO.BP.biomart[annotation.GO.BP.biomart$ensembl_gene_id %in% test.g
enes$a, ]
GO_BP_Cats_mRNA_AI_NB_Preg_NP<-merge(GO_BP_Cats_mRNA_AI_NB_Preg_NP,annotation.GO.BP.biomart_testgenes, by.x="cate
gory", by.y="go_id", all.x=TRUE, all.y=FALSE)
GO_BP_Cats_mRNA_AI_NB_Preg_NP<-merge(GO_BP_Cats_mRNA_AI_NB_Preg_NP, annotation.ensembl.symbol, by.x="ensembl_gene
_id", by.y="ensembl_gene_id", all=FALSE, all.x=TRUE, all.y=FALSE)
GO_BP_Cats_mRNA_AI_NB_Preg_NP<-GO_BP_Cats_mRNA_AI_NB_Preg_NP[with(GO_BP_Cats_mRNA_AI_NB_Preg_NP, order(BY_FDR,ter
m)), ]
GO_BP_Cats_mRNA_AI_NB_Preg_NP<-GO_BP_Cats_mRNA_AI_NB_Preg_NP[GO_BP_Cats_mRNA_AI_NB_Preg_NP$BY_FDR < 0.1,]
#write.table(GO_BP_Cats_mRNA_AI_NB_Preg_NP, paste(result_path, "2019_12_23_GO_BP_Cats_mRNA_miRNA_gain_positive_co
expression_AI_NB_Preg_NP.txt", sep="/"), quote=FALSE, sep="\t", row.names=FALSE)
```

```
knitr::kable(head(GO_BP_Cats_mRNA_AI_NB_Preg_NP), caption = "First five rows of Supplementary Table 10",format =
'pandoc')
```

First five rows of Supplementary Table 10

| | ensembl_gene_id | category | over_represented_pvalue | under_represented_pvalue | numDEInCat | numInCat | term | ontology | BY |
|---|---|---|---|---|---|---|---|---|---|
| 683 | ENSBTAG00000006065 | GO:0006275 (GO:0006275) | 0.0001428 | 1 | 8 | 15 | regulation of DNA replication | BP | 0.04 |
| 755 | ENSBTAG00000006551 | GO:0006275 (GO:0006275) | 0.0001428 | 1 | 8 | 15 | regulation of DNA replication | BP | 0.04 |
| 1550 | ENSBTAG00000013905 | GO:0006275 (GO:0006275) | 0.0001428 | 1 | 8 | 15 | regulation of DNA replication | BP | 0.04 |
| 1729 | ENSBTAG00000015338 | GO:0006275 (GO:0006275) | 0.0001428 | 1 | 8 | 15 | regulation of DNA replication | BP | 0.04 |
| 1878 | ENSBTAG00000016572 | GO:0006275 (GO:0006275) | 0.0001428 | 1 | 8 | 15 | regulation of DNA replication | BP | 0.04 |
| 1984 | ENSBTAG00000017329 | GO:0006275 (GO:0006275) | 0.0001428 | 1 | 8 | 15 | regulation of DNA replication | BP | 0.04 |

```
gc()
```

### Differential gene expression associated with pregnancy outcome

#### AI Pregnant vs non pregnant

##### Results from edgeR

```
#colnames(count_pwbc)
group<-factor(c("Preg_AI","Preg_AI","Preg_AI","Preg_AI","Preg_AI","Preg_AI","Preg_NB","Preg_NB","Preg_NB","Preg_N
B","Preg_NB","Preg_NB","Not_Preg","Not_Preg","Not_Preg","Not_Preg","Not_Preg"), levels=c("Preg_AI","Preg_NB", "No
t_Preg"))
design<-model.matrix(~group)
dds<-DGEList(count=count_pwbc, group=group)
dds<-estimateDisp(dds, design, robust=TRUE)
dds<-glmFit(dds, design)
dds<-glmLRT(dds,coef="groupNot_Preg")
res_not_preg_preg_AI_edgeR<-topTags(dds,n=Inf)$table
#head(res_not_preg_preg_AI_edgeR,n=30)
#write.table(res_not_preg_preg_AI_edgeR, paste(result_path, "2019_04_30_edgeR_AI_NP_mRNA.txt"), quote=FALSE, sep
="\t", row.names=FALSE)
```

##### Results from edgeR both years

```
#colnames(merged_datasets)
group<-factor(c("Preg_AI","Preg_AI","Preg_AI","Preg_AI","Preg_AI","Preg_AI","Preg_NB","Preg_NB","Preg_NB","Preg_N
B","Preg_NB","Preg_NB","Not_Preg","Not_Preg","Not_Preg","Not_Preg","Not_Preg",
                "Preg_AI","Not_Preg", "Not_Preg", "Not_Preg", "Preg_AI", "Preg_AI","Preg_AI", "Preg_AI", "Not_Pre
g", "Preg_AI",
                "Not_Preg", "Not_Preg"), levels=c("Preg_AI","Preg_NB", "Not_Preg"))
year<-factor(rep(c("A","B"), c(17,12)))
design<-model.matrix(~0+group+year)
dds<-DGEList(count=merged_datasets, group=group)
dds<-estimateDisp(dds, design, robust=TRUE)
contrast_not_preg_preg_AI<-makeContrasts( groupNot_Preg-groupPreg_AI, levels=design)
dds<-glmFit(dds, design)
dds<-glmLRT(dds,contrast=contrast_not_preg_preg_AI)
res_not_preg_preg_AI_edgeR_merged<-topTags(dds,n=Inf)$table
#head(res_not_preg_preg_AI_edgeR_merged,n=30)
#write.table(res_not_preg_preg_AI_edgeR, paste(result_path, "2019_04_16_edgeR_AI_NP_mRNA.txt"), quote=FALSE, sep
="\t", row.names=FALSE)
```

##### Results from DESeq

```
group<-factor(c("Preg_AI","Preg_AI","Preg_AI","Preg_AI","Preg_AI","Preg_AI","Preg_NB","Preg_NB","Preg_NB","Preg_N
B","Preg_NB","Preg_NB","Not_Preg","Not_Preg","Not_Preg","Not_Preg","Not_Preg"), levels=c("Preg_AI","Preg_NB", "No
t_Preg"))
design<-model.matrix(~group)
colData<-data.frame("group"=group)
rownames(colData)<-colnames(count_pwbc)
dds<-DESeqDataSetFromMatrix(countData=count_pwbc, colData=colData, design= ~group)
dds<-DESeq(dds)
res_not_preg_preg_AI_DeSeq<-results(dds, contrast=c("group","Not_Preg" ,"Preg_AI"), pAdjustMethod="fdr", tidy=TRU
E)
res_not_preg_preg_AI_DeSeq<-res_not_preg_preg_AI_DeSeq[with(res_not_preg_preg_AI_DeSeq, order(pvalue)), ]
#head(res_not_preg_preg_AI_DeSeq,n=30)
#write.table(res_not_preg_preg_AI_DeSeq, paste(result_path, "2019_04_30_DESeq_AI_NP_mRNA.txt"), quote=FALSE, sep
="\t", row.names=FALSE)
```

#### Results from DESeq from both years

```
group<-factor(c("Preg_AI","Preg_AI","Preg_AI","Preg_AI","Preg_AI","Preg_AI","Preg_NB","Preg_NB","Preg_NB","Preg_N
B","Preg_NB","Preg_NB","Not_Preg","Not_Preg","Not_Preg","Not_Preg","Not_Preg",
                "Preg_AI","Not_Preg", "Not_Preg", "Not_Preg", "Preg_AI", "Preg_AI","Preg_AI", "Preg_AI", "Not_Pre
g", "Preg_AI",
                "Not_Preg", "Not_Preg"), levels=c("Preg_AI","Preg_NB", "Not_Preg"))
design<-model.matrix(~group+year)
colData<-data.frame("group"=group, "year"=year)
rownames(colData)<-colnames(merged_datasets)
dds<-DESeqDataSetFromMatrix(countData=merged_datasets, colData=colData, design= ~group+year)
dds<-DESeq(dds)
res_not_preg_preg_AI_DeSeq_merged<-results(dds, contrast=c("group", "Not_Preg", "Preg_AI") , pAdjustMethod="fdr",
tidy=TRUE)
res_not_preg_preg_AI_DeSeq_merged<-res_not_preg_preg_AI_DeSeq_merged[with(res_not_preg_preg_AI_DeSeq_merged, orde
r(pvalue)), ]
#head(res_not_preg_preg_AI_DeSeq_merged,n=30)
#write.table(res_not_preg_preg_AI_DeSeq, paste(result_path, "2019_04_22_DESeq_AI_NP_mRNA.txt"), quote=FALSE, sep
="\t", row.names=FALSE)
```

### Supplementary Fig. 4

```
setwd("/mnt/storage/auburn/heifer_pregnancy/proj_2018/analysis/results/")
#Create the permutations matrix
perm_without_replacement <- function(n, r){
  return(factorial(n)/factorial(n - r))
}

permutation_matrix<-matrix(,nrow=200100,ncol=17)
head(permutation_matrix)
dim(permutation_matrix)

for (i in seq(1:200100)){
  sampling <- sample(1:17,17, replace = FALSE)
  if ( ! identical(sampling , c(1:17))){
    permutation_matrix[i,]<-sample(1:17,17, replace = FALSE)
  }}

permutation_matrix<-permutation_matrix[!duplicated(permutation_matrix),]
dim(permutation_matrix)
head(permutation_matrix)

permutation_matrix<-permutation_matrix[sample(1:dim(permutation_matrix)[1], 10000),]
rand<-dim(permutation_matrix)[1]

sequence.pvalue<-seq(0.005, 0.05, 0.005)

#edgeR
results <- filebacked.big.matrix(length(sequence.pvalue),rand, type="double", init=0, separated=FALSE,
                                 backingfile="incidence_matrix.bin",
                                 descriptor="incidence_matrix.desc")
mdesc_result<- describe(results)

#edgeR
cl <- makeCluster(30)
registerDoParallel(cl)

results[,]<-foreach(i = sequence.pvalue, .combine='rbind', .inorder=TRUE,  .packages=c("edgeR","bigmemory"), .ver
bose=FALSE) %:%

  foreach(j = 1:rand, .combine='cbind', .inorder=FALSE,.packages=c("edgeR","bigmemory"), .verbose=FALSE ) %dopar%
{

    group<-factor(c("Preg_AI","Preg_AI","Preg_AI","Preg_AI","Preg_AI","Preg_AI","Preg_NB","Preg_NB","Preg_NB","Pr
eg_NB","Preg_NB","Preg_NB","Not_Preg","Not_Preg","Not_Preg","Not_Preg","Not_Preg"), levels=c("Preg_AI","Preg_NB",
"Not_Preg"))
    group<-group[permutation_matrix[j,]]
    group<-factor(group, levels=c("Preg_AI","Not_Preg", "Preg_NB"))
    design <- model.matrix(~group)
    dds<-DGEList(count=count_pwbc, group=group)
    dds<-estimateDisp(dds, design, robust=TRUE)
    dds <- glmFit(dds, design)
    dds <- glmLRT(dds, coef="groupNot_Preg")
    length(which(topTags(dds,n=Inf)$table$PValue < i))

  }
stopCluster(cl)
results[1:5,1:5]
total.rand <- rand * 10496

qvalue<-data.frame(raw.pvalue = sequence.pvalue,
                   e.pvalue.edgeR= (rowSums(results[,]+1))/(total.rand+1),
                   e.pvalue.edgeR.round= round((rowSums(results[,]+1))/(total.rand+1) ,4))

#qvalue
rm(results)
system("rm incidence_matrix.bin")
system("rm incidence_matrix.desc")

#DESEQ2

results.rand <- filebacked.big.matrix(length(sequence.pvalue),rand, type="double", init=0, separated=FALSE,
                                      backingfile="incidence_matrix.bin",
                                      descriptor="incidence_matrix.desc")
mdesc_result<- describe(results.rand)

cl <- makeCluster(30)
registerDoParallel(cl)

results.rand[,]<-foreach(i = sequence.pvalue, .combine='rbind', .inorder=TRUE,  .packages=c("DESeq2","bigmemory"
), .verbose=TRUE) %:%

  foreach(j = 1:rand, .combine='cbind', .inorder=FALSE,.packages=c("DESeq2","bigmemory"), .verbose=TRUE ) %dopar%
```

```
{

    group<- c("Preg_AI","Preg_AI","Preg_AI","Preg_AI","Preg_AI","Preg_AI","Preg_NB","Preg_NB","Preg_NB","Preg_NB"
,"Preg_NB","Preg_NB","Not_Preg","Not_Preg","Not_Preg","Not_Preg","Not_Preg")
    group<-group[permutation_matrix[j,]]
    group<-factor(group, levels=c("Preg_AI","Not_Preg", "Preg_NB"))
    design <- model.matrix(~group)

    colData<-data.frame("group"=group)
    rownames(colData)<-colnames(count_pwbc)
    dds<-DESeqDataSetFromMatrix(countData=count_pwbc, colData=colData, design= ~group)
    dds <- DESeq(dds)
    DeSeq_AI_NP <- results(dds, contrast=c("group","Not_Preg" ,"Preg_AI"), pAdjustMethod="fdr", tidy=TRUE)

    length(which(DeSeq_AI_NP$pvalue < i))

  }

stopCluster(cl)

results.rand[1:5,1:5]

total.rand <- rand * 10496


qvalue2<-data.frame(raw.pvalue = sequence.pvalue,
                    e.pvalue.DESEQ2= (rowSums(results.rand[,]+1))/(total.rand+1),
                    e.pvalue.DESEQ2.round= round((rowSums(results.rand[,]+1))/(total.rand+1) ,4))

qvalue2


qvalue3<-cbind(qvalue,qvalue2)

qvalue3

rm(results.rand)
system("rm incidence_matrix.bin")
system("rm incidence_matrix.desc")

#write.table(qvalue3,paste(result_path,"2019_12_26_empirical_FDR_for_DEG.txt",sep="/"), quote=FALSE, sep="\t", ro
w.names=FALSE)
```

```
eFDR_differential_expression<-read.delim(paste(result_path,"2019_12_26_empirical_FDR_for_DEG.txt",sep="/"), row.n
ames=NULL,header =TRUE, stringsAsFactors =FALSE)
```

```
font_size<-9

plot1<-ggplot()+
  geom_point(data=eFDR_differential_expression, aes(x=e.pvalue.edgeR.round , y=raw.pvalue),color="black", size=3,
shape=16)+
  geom_line(data=eFDR_differential_expression, aes(x=e.pvalue.edgeR.round , y=raw.pvalue),color="black", size=0.1
,linetype=3)+
  scale_x_continuous(name="empirical FDR", limits = c(0, 0.08), breaks=seq(0, 0.08, 0.005))+
  scale_y_continuous(name="nominal P value", limits = c(0, 0.05), breaks=seq(0, 0.06, 0.005))+
  ggtitle("Differential gene expression with edgeR")+
  theme_bw()+
  theme(panel.grid= element_blank(),
        panel.background = element_blank(),
        panel.grid.minor = element_blank(),
        panel.grid.major = element_line(color="lightgray"),
        plot.background = element_blank(),
        axis.text.y=element_text(color="black", size=font_size),
        axis.text.x=element_text(color="black", size=font_size, angle=90),
        panel.spacing = unit(c(0.4,0.4,0.4,0.4),"cm"),
        plot.margin = unit(c(0.5,0.5,0.5,0.5),"cm"),
        legend.position="none",
        plot.title = element_text(lineheight=.8, hjust=0.5, size= font_size))

plot2<-ggplot()+
  geom_point(data=eFDR_differential_expression, aes(x=e.pvalue.DESEQ2.round , y=raw.pvalue),color="black", size=3
, shape=16)+
  geom_line(data=eFDR_differential_expression, aes(x=e.pvalue.DESEQ2.round , y=raw.pvalue),color="black", size=0.
1,linetype=3)+
  scale_x_continuous(name="empirical FDR", limits = c(0, 0.08), breaks=seq(0, 0.08, 0.005))+
  scale_y_continuous(name="nominal P value", limits = c(0, 0.05), breaks=seq(0, 0.05, 0.005))+
  ggtitle("Differential gene expression with DESEQ2")+
  theme_bw()+
  theme(panel.grid= element_blank(),
        panel.background = element_blank(),
        panel.grid.minor = element_blank(),
        panel.grid.major = element_line(color="lightgray"),
        plot.background = element_blank(),
        axis.title=element_text(color="black", size=font_size),
        axis.text.y=element_text(color="black", size=font_size),
        axis.text.x=element_text(color="black", size=font_size, angle=90),
        panel.spacing = unit(c(0.4,0.4,0.4,0.4),"cm"),
        plot.margin = unit(c(0.5,0.5,0.5,0.5),"cm"),
        legend.position="none",
        plot.title = element_text(lineheight=.8, hjust=0.5, size= font_size))

#font_size 12 for figure
#pdf(paste(result_path,"2019_12_27_eFDR_plots_differential_gene_expression.pdf", sep="/"), width=10, height=4)
#grid.arrange(plot1,plot2, ncol = 2)
#dev.off()
grid.arrange(plot1,plot2, ncol = 2)
```

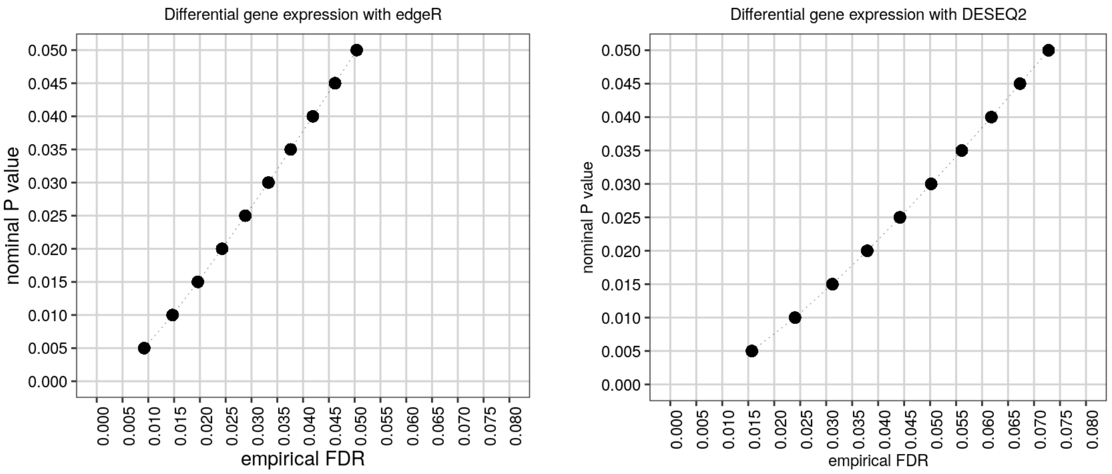

```
merged_res_not_preg_preg_AI_mRNA<-merge(res_not_preg_preg_AI_edgeR, res_not_preg_preg_AI_DeSeq, by.x="row.names", by.y="row")
merged_res_not_preg_preg_AI_mRNA<-merged_res_not_preg_preg_AI_mRNA[merged_res_not_preg_preg_AI_mRNA$PValue<=0.03 & merged_res_not_preg_preg_AI_mRNA$pvalue<=0.03,]
merged_res_not_preg_preg_AI_mRNA<-merged_res_not_preg_preg_AI_mRNA[with(merged_res_not_preg_preg_AI_mRNA, order(logFC)), ]
#dim(merged_res_not_preg_preg_AI_mRNA) #353
#head(merged_res_not_preg_preg_AI_mRNA)
#write.table(merged_res_not_preg_preg_AI_mRNA, paste(result_path, "2019_04_30_merged_AI_NP_mRNA_before_LOOCV.txt"), quote=FALSE, sep="\t", row.names=FALSE)
DEG_AI_NP_mRNA<-merge(merged_res_not_preg_preg_AI_mRNA,annotation.ensembl.symbol, by.x="Row.names", by.y="ensembl_gene_id", all=FALSE)
DEG_AI_NP_mRNA<-DEG_AI_NP_mRNA[with(DEG_AI_NP_mRNA, order(logFC)), ]
is.num <- sapply(DEG_AI_NP_mRNA, is.numeric)
DEG_AI_NP_mRNA[is.num] <- lapply(DEG_AI_NP_mRNA[is.num], round, 4)
#write.table(DEG_AI_NP_mRNA, paste(result_path, "2019_12_26_merged_AI_NP_mRNA_gene_names_before_LOOCV.txt", sep="/"), quote=FALSE, sep="\t", row.names=FALSE)
```

```
merged_res_not_preg_preg_AI_mRNA_merged<-merge(res_not_preg_preg_AI_edgeR_merged, res_not_preg_preg_AI_DeSeq_merged, by.x="row.names", by.y="row")
merged_res_not_preg_preg_AI_mRNA_merged_a<-merged_res_not_preg_preg_AI_mRNA_merged[merged_res_not_preg_preg_AI_mRNA_merged$PValue<=0.03 & merged_res_not_preg_preg_AI_mRNA_merged$pvalue<=0.03,]
#write.table(merged_res_not_preg_preg_AI_mRNA, paste(result_path, "2019_04_22_merged_AI_NP_mRNA.txt"), quote=FALSE, sep="\t", row.names=FALSE)
DEG_AI_NP_mRNA_merged<-merge(merged_res_not_preg_preg_AI_mRNA_merged_a,annotation.ensembl.symbol, by.x="Row.names", by.y="ensembl_gene_id", all=FALSE)
DEG_AI_NP_mRNA_merged<-DEG_AI_NP_mRNA_merged[with(DEG_AI_NP_mRNA_merged, order(logFC)), ]
is.num <- sapply(DEG_AI_NP_mRNA_merged, is.numeric)
DEG_AI_NP_mRNA_merged[is.num] <- lapply(DEG_AI_NP_mRNA_merged[is.num], round, 4)
#write.table(DEG_AI_NP_mRNA_merged, paste(result_path, "2019_12_27_merged_AI_NP_mRNA_gene_names_both_years.txt", sep="/"), quote=FALSE, sep="\t", row.names=FALSE)
```

Plot the charts for all genes inferred as differentially expressed between AI and non pregnant. Not added as a supplementary material due to space.

```
cpm_AIpreg<-cpm_pwbc[,c(1,2,3,4,5,6)]
cpm_NOTpreg<-cpm_pwbc[,c(13,14,15,16,17)]
cpm_NBpreg<-cpm_pwbc[,c(7,8,9,10,11,12)]

data_chart_1<-data.frame( stringsAsFactors=FALSE)
data_chart_2<-data.frame( stringsAsFactors=FALSE)

for (i in  seq(dim(DEG_AI_NP_mRNA)[1])){

  gene_id<-DEG_AI_NP_mRNA[i,1]
  gene_symbol<-DEG_AI_NP_mRNA$external_gene_name[i]
  if(gene_symbol==""){gene_symbol<-gene_id}

  cpm_AI<-cpm_AIpreg[rownames(cpm_AIpreg)==gene_id,]
  cpm_NP<-cpm_NOTpreg[rownames(cpm_NOTpreg)==gene_id,]

  data_chart_1<-rbind(data.frame(cpm=cpm_AI), data.frame(cpm=cpm_NP))
  data_chart_1$group<-rep(c("Preg_AI","Not_Preg"),c(6,5))
  data_chart_1$shape<-as.factor(c(0:10))
  data_chart_1$gene<-gene_symbol
  data_chart_1$chart<-i
  data_chart_1$fold_change<-DEG_AI_NP_mRNA[DEG_AI_NP_mRNA$Row.names==gene_id,]$logFC
  data_chart_1$pvalue<-DEG_AI_NP_mRNA[DEG_AI_NP_mRNA$Row.names==gene_id,]$PValue

  data_chart_2<-rbind(data_chart_2,data_chart_1)
}

pdf(file=paste(result_path,"2019_12_27_plot_all_DEGs_preg_AI_not_preg_ordered_by_FC_before_LOOCV.pdf",sep="/"), w
idth= 11, paper="USr", bg="transparent", onefile=TRUE)

for (i in seq(1,dim(DEG_AI_NP_mRNA)[1], 24)) {
  plots <- list()
  k<-1
  for (j in c(i:(i+23))){
    data_chart_3<-data_chart_2[data_chart_2$chart %in% j , ]
    plot<-ggplot(data=data_chart_3, aes(x=group , y=cpm, colour=group))+
      geom_jitter(position = position_jitter(width = .1, height=0), aes( shape=shape), size=2)+
      scale_shape_manual(values=c(0:11))+
      scale_x_discrete(data_chart_3$gene[1],labels=c("Preg_AI"="Preg_AI", "Not_Preg"="Not_Preg"))+
      scale_colour_manual(name=NULL, values = c("#D55E00", "#0072B2"))+
      stat_summary(fun.y = median, fun.ymin = median, fun.ymax = median,colour = "gray", size = 0.2,  geom = "cro
ssbar", alpha=0.5)+
      scale_y_continuous("CPM")+
      ggtitle(paste("LogFC","=", round(data_chart_3$fold_change[1],1),"  ","P","=",round(data_chart_3$pvalue[1],4
), sep=c("")))+
      theme_bw()+
      theme(panel.grid= element_blank(),
            panel.background = element_blank(),
            panel.grid.minor = element_blank(),
            panel.grid.major = element_blank(),
            plot.background = element_blank(),
            strip.background = element_rect(fill = "white"),
            #strip.text.x = element_text(colour = 'black', face="italic",size = 9),
            #legend.text = element_text( colour = 'black', size = 10 ),
            axis.text.x = element_blank(),
            axis.text.y = element_text( colour = 'black', size = 8 ),
            axis.title = element_text( colour = 'black', size = 9 ),
            legend.key.size = unit(0.9,"cm"),
            legend.position="none",
            axis.ticks.x = element_blank(),
            plot.title = element_text(size = 7)
      )

    plots[[k]] <- plot
    k<-k+1
  }
  multiplot(plotlist = plots, cols = 6,layout = matrix(1:24, nrow=4, byrow = TRUE))
}
dev.off()
```

Plot the charts for all genes inferred as differentially expressed between AI and non pregnant for merged dataset. Not added as a supplementary material due to space.

```
cpm_AIpreg<-cpm_merged_datasets[,c(1,2,3,4,5,6,18,22,23,24,25,27)]
cpm_NOTpreg<-cpm_merged_datasets[,c(13,14,15,16,17,19,20,21,26,28,29)]


data_chart_1<-data.frame( stringsAsFactors=FALSE)
data_chart_2<-data.frame( stringsAsFactors=FALSE)

for (i in  seq(dim(DEG_AI_NP_mRNA_merged)[1])){

  gene_id<-DEG_AI_NP_mRNA_merged[i,1]
  gene_symbol<-DEG_AI_NP_mRNA_merged$external_gene_name[i]
  if(gene_symbol==""){gene_symbol<-gene_id}

  cpm_AI<-cpm_AIpreg[rownames(cpm_AIpreg)==gene_id,]
  cpm_NP<-cpm_NOTpreg[rownames(cpm_NOTpreg)==gene_id,]

  data_chart_1<-rbind(data.frame(cpm=cpm_AI), data.frame(cpm=cpm_NP))
  data_chart_1$group<-rep(c("Preg_AI","Not_Preg"),c(12,11))
  data_chart_1$shape<-as.factor(c(0:22))
  data_chart_1$gene<-gene_symbol
  data_chart_1$chart<-i
  data_chart_1$fold_change<-DEG_AI_NP_mRNA_merged[DEG_AI_NP_mRNA_merged$Row.names==gene_id,]$logFC
  data_chart_1$pvalue<-DEG_AI_NP_mRNA_merged[DEG_AI_NP_mRNA_merged$Row.names==gene_id,]$PValue


  data_chart_2<-rbind(data_chart_2,data_chart_1)
}

pdf(file=paste(result_path,"2019_12_27_plot_all_DEGs_preg_AI_not_preg_merged.pdf",sep="/"), width= 11, paper="US
r", bg="transparent", onefile=TRUE)

for (i in seq(1,dim(DEG_AI_NP_mRNA_merged)[1], 24)) {
  plots <- list()
  k<-1
  for (j in c(i:(i+23))){
    data_chart_3<-data_chart_2[data_chart_2$chart %in% j , ]
    plot<-ggplot(data=data_chart_3, aes(x=group , y=cpm, colour=group))+
      geom_jitter(position = position_jitter(width = .1, height=0), aes( shape=shape), size=2)+
      scale_shape_manual(values=c(0:22))+
      scale_x_discrete(data_chart_3$gene[1],labels=c("Preg_AI"="Preg_AI", "Not_Preg"="Not_Preg"))+
      scale_colour_manual(name=NULL, values = c("#D55E00", "#0072B2"))+
      stat_summary(fun.y = median, fun.ymin = median, fun.ymax = median,colour = "gray", size = 0.2,  geom = "cro
ssbar", alpha=0.5)+
      scale_y_continuous("CPM")+
      ggtitle(paste("LogFC","=", round(data_chart_3$fold_change[1],1),"  ","P","=",round(data_chart_3$pvalue[1],4
), sep=c("")))+
      theme_bw()+
      theme(panel.grid= element_blank(),
            panel.background = element_blank(),
            panel.grid.minor = element_blank(),
            panel.grid.major = element_blank(),
            plot.background = element_blank(),
            strip.background = element_rect(fill = "white"),
            #strip.text.x = element_text(colour = 'black', face="italic",size = 9),
            #legend.text = element_text( colour = 'black', size = 10 ),
            axis.text.x = element_blank(),
            axis.text.y = element_text( colour = 'black', size = 8 ),
            axis.title = element_text( colour = 'black', size = 9 ),
            legend.key.size = unit(0.9,"cm"),
            legend.position="none",
            axis.ticks.x = element_blank(),
            plot.title = element_text(size = 7)
      )

    plots[[k]] <- plot
    k<-k+1
  }
  multiplot(plotlist = plots, cols = 6,layout = matrix(1:24, nrow=4, byrow = TRUE))
}
dev.off()
```

### Supplementary table 11

Leave-one-out cross validation.

```
cl <- makeCluster(12)
registerDoParallel(cl)

group<-factor(c("Preg_AI","Preg_AI","Preg_AI","Preg_AI","Preg_AI","Preg_AI","Preg_NB","Preg_NB","Preg_NB","Preg_N
B","Preg_NB","Preg_NB","Not_Preg","Not_Preg","Not_Preg","Not_Preg","Not_Preg"), levels=c("Preg_AI","Preg_NB", "No
t_Preg"))

out_list_IDs_LOOCV<-foreach(i = c(1:6,13:17), .inorder=FALSE,.packages=c("edgeR","DESeq2"), .verbose=FALSE ) %dop
ar% {

group_a<-group[-i]
count_pwbc_a<-count_pwbc[,-i]

design<-model.matrix(~group_a)
dds<-DGEList(count=count_pwbc_a, group=group_a)
dds<-estimateDisp(dds, design, robust=TRUE)
dds<-glmFit(dds, design)
dds<-glmLRT(dds,coef="group_aNot_Preg")
res_not_preg_preg_AI_edgeR<-topTags(dds,n=Inf)$table

#DESeq
colData<-data.frame("group"=group_a)
rownames(colData)<-colnames(count_pwbc_a)
dds<-DESeqDataSetFromMatrix(countData=count_pwbc_a, colData=colData, design= ~group)
dds<-DESeq(dds)
res_not_preg_preg_AI_DeSeq<-results(dds, contrast=c("group","Not_Preg" ,"Preg_AI"), pAdjustMethod="fdr", tidy=TRU
E)


merged_res_not_preg_preg_AI_mRNA_a<-merge(res_not_preg_preg_AI_edgeR, res_not_preg_preg_AI_DeSeq, by.x="row.name
s", by.y="row")
merged_res_not_preg_preg_AI_mRNA_a<-merged_res_not_preg_preg_AI_mRNA_a[merged_res_not_preg_preg_AI_mRNA_a$PValue<
=0.03 & merged_res_not_preg_preg_AI_mRNA_a$pvalue<=0.03,]

rm(group_a,count_pwbc_a)

merged_res_not_preg_preg_AI_mRNA_a$Row.names
}

stopCluster(cl)

overlapping_genes_LOOCV<-Reduce(intersect, out_list_IDs_LOOCV)

merged_res_not_preg_preg_AI_mRNA<-merge(res_not_preg_preg_AI_edgeR, res_not_preg_preg_AI_DeSeq, by.x="row.names",
by.y="row")
merged_res_not_preg_preg_AI_mRNA_a<-merged_res_not_preg_preg_AI_mRNA[merged_res_not_preg_preg_AI_mRNA$Row.names %
in% overlapping_genes_LOOCV,]
DEG_AI_NP_mRNA<-merge(merged_res_not_preg_preg_AI_mRNA_a,annotation.ensembl.symbol, by.x="Row.names", by.y="ensem
bl_gene_id", all=FALSE)
DEG_AI_NP_mRNA<-DEG_AI_NP_mRNA[with(DEG_AI_NP_mRNA, order(logFC)), ]
is.num <- sapply(DEG_AI_NP_mRNA, is.numeric)
DEG_AI_NP_mRNA[is.num] <- lapply(DEG_AI_NP_mRNA[is.num], round, 4)
#dim(DEG_AI_NP_mRNA)
#head(DEG_AI_NP_mRNA)
#write.table(DEG_AI_NP_mRNA, paste(result_path, "2019_12_27_merged_AI_NP_mRNA_gene_names_after_LOOCV.txt",sep
="/"), quote=FALSE, sep="\t", row.names=FALSE)
rm(out_list_IDs_LOOCV)
```

```
knitr::kable(head(DEG_AI_NP_mRNA), caption = "First five rows of Supplementary Table 11",format = 'pandoc')
```

First five rows of Supplementary Table 11

| | Row.names | logFC | logCPM | LR | PValue | FDR | baseMean | log2FoldChange | lfcSE | stat | pvalue | padj | external_gene_n |
|---|---|---|---|---|---|---|---|---|---|---|---|---|---|
| 64 | ENSBTAG00000052118 | -4.4422 | 0.1925 | 8.1231 | 0.0044 | 0.6588 | 17.1086 | -4.5092 | 1.5625 | -2.8860 | 0.0039 | 0.2636 | |
| 49 | ENSBTAG00000034281 | -3.0239 | 1.0672 | 7.2748 | 0.0070 | 0.6588 | 36.8808 | -2.8357 | 1.0866 | -2.6098 | 0.0091 | 0.3176 | MGAT4D |
| 66 | ENSBTAG00000052658 | -2.2694 | 4.0733 | 11.1769 | 0.0008 | 0.6467 | 291.2199 | -1.7625 | 0.6162 | -2.8603 | 0.0042 | 0.2636 | NKG2A |
| 62 | ENSBTAG00000051183 | -2.1635 | 4.1301 | 13.4758 | 0.0002 | 0.3624 | 308.3543 | -1.9229 | 0.5047 | -3.8097 | 0.0001 | 0.1073 | |
| 14 | ENSBTAG00000007247 | -2.1530 | 4.7636 | 8.3500 | 0.0039 | 0.6588 | 471.4847 | -2.0381 | 0.6571 | -3.1018 | 0.0019 | 0.2122 | NUF2 |

| | Row.names | logFC | logCPM | LR | PValue | FDR | baseMean | log2FoldChange | lfcSE | stat | pvalue | padj | external_gene_n |
|---|---|---|---|---|---|---|---|---|---|---|---|---|---|
| 63 | ENSBTAG00000051500 | -2.1186 | 2.2460 | 6.7875 | 0.0092 | 0.6588 | 81.1065 | -2.0127 | 0.7156 | -2.8127 | 0.0049 | 0.2676 | ZNF389 |

Plot the charts for all genes inferred as differentially expressed between AI and non pregnant. Not added to the supplementary material due do space.

```
cpm_AIpreg<-cpm_pwbc[,c(1,2,3,4,5,6)]
cpm_NOTpreg<-cpm_pwbc[,c(13,14,15,16,17)]
cpm_NBpreg<-cpm_pwbc[,c(7,8,9,10,11,12)]

data_chart_1<-data.frame( stringsAsFactors=FALSE)
data_chart_2<-data.frame( stringsAsFactors=FALSE)

for (i in  seq(dim(DEG_AI_NP_mRNA)[1])){

  gene_id<-DEG_AI_NP_mRNA[i,1]
  gene_symbol<-DEG_AI_NP_mRNA$external_gene_name[i]
  if(gene_symbol==""){gene_symbol<-gene_id}

  cpm_AI<-cpm_AIpreg[rownames(cpm_AIpreg)==gene_id,]
  cpm_NP<-cpm_NOTpreg[rownames(cpm_NOTpreg)==gene_id,]

  data_chart_1<-rbind(data.frame(cpm=cpm_AI), data.frame(cpm=cpm_NP))
  data_chart_1$group<-rep(c("Preg_AI","Not_Preg"),c(6,5))
  data_chart_1$shape<-as.factor(c(0:10))
  data_chart_1$gene<-gene_symbol
  data_chart_1$chart<-i
  data_chart_1$fold_change<-DEG_AI_NP_mRNA[DEG_AI_NP_mRNA$Row.names==gene_id,]$logFC
  data_chart_1$pvalue<-DEG_AI_NP_mRNA[DEG_AI_NP_mRNA$Row.names==gene_id,]$PValue


  data_chart_2<-rbind(data_chart_2,data_chart_1)
}

pdf(file=paste(result_path,"2019_12_27_plot_all_DEGs_preg_AI_not_preg_ordered_by_FC_after_LOOCV.pdf",sep="/"), wi
dth= 11, paper="USr", bg="transparent", onefile=TRUE)


for (i in seq(1,dim(DEG_AI_NP_mRNA)[1], 24)) {
  plots <- list()
  k<-1
  for (j in c(i:(i+23))){
    data_chart_3<-data_chart_2[data_chart_2$chart %in% j , ]
    plot<-ggplot(data=data_chart_3, aes(x=group , y=cpm, colour=group))+
      geom_jitter(position = position_jitter(width = .1, height=0), aes( shape=shape), size=2)+
      scale_shape_manual(values=c(0:11))+
      scale_x_discrete(data_chart_3$gene[1],labels=c("Preg_AI"="Preg_AI", "Not_Preg"="Not_Preg"))+
      scale_colour_manual(name=NULL, values = c("#D55E00", "#0072B2"))+
      stat_summary(fun.y = median, fun.ymin = median, fun.ymax = median,colour = "gray", size = 0.2,  geom = "cro
ssbar", alpha=0.5)+
      scale_y_continuous("CPM")+
      ggtitle(paste("LogFC","=", round(data_chart_3$fold_change[1],1),"  ","P","=",round(data_chart_3$pvalue[1],4
), sep=c("")))+
      theme_bw()+
      theme(panel.grid= element_blank(),
            panel.background = element_blank(),
            panel.grid.minor = element_blank(),
            panel.grid.major = element_blank(),
            plot.background = element_blank(),
            strip.background = element_rect(fill = "white"),
            #strip.text.x = element_text(colour = 'black', face="italic",size = 9),
            #legend.text = element_text( colour = 'black', size = 10 ),
            axis.text.x = element_blank(),
            axis.text.y = element_text( colour = 'black', size = 8 ),
            axis.title = element_text( colour = 'black', size = 9 ),
            legend.key.size = unit(0.9,"cm"),
            legend.position="none",
            axis.ticks.x = element_blank(),
            plot.title = element_text(size = 7)
      )

    plots[[k]] <- plot
    k<-k+1
  }
  multiplot(plotlist = plots, cols = 6,layout = matrix(1:24, nrow=4, byrow = TRUE))
}
dev.off()
```

Create the background gene lists for gene ontology.

```
all_genes<-data.frame( gene=rownames(count_pwbc), stringsAsFactors=FALSE )
rownames(all_genes)<-all_genes$gene

N_expressed_genes<-length(all_genes$gene)

annotation.genelength.biomart_vector<-gene_length[,2]
names(annotation.genelength.biomart_vector)<-gene_length[,1]

annotation.GO.BP.biomart<-annotation.GO.biomart[annotation.GO.biomart$namespace_1003=="biological_process", c(1,3
)]
annotation.GO.BP.biomart<-annotation.GO.BP.biomart[annotation.GO.BP.biomart$ensembl_gene_id %in% rownames(all_gen
es),]
annotation.GO.MF.biomart<-annotation.GO.biomart[annotation.GO.biomart$namespace_1003=="molecular_function", c(1,3
)]
annotation.GO.MF.biomart<-annotation.GO.MF.biomart[annotation.GO.MF.biomart$ensembl_gene_id %in% rownames(all_gen
es),]
```

Test biological processes for enrichment with DEGs and calculate fold change enrichment. (AI Pregnant vs non pregnant)

```
test.genes<-data.frame(a=unique(c(as.character(merged_res_not_preg_preg_AI_mRNA_a$Row.names))), stringsAsFactors=
FALSE)
all_genes_numeric<-as.integer(all_genes$gene %in%test.genes$a)
names(all_genes_numeric)<-all_genes$gene

N_DEGs<-length(test.genes$a)

set.seed(9830)
pwf<-nullp(all_genes_numeric, bias.data=annotation.genelength.biomart_vector, plot.fit=FALSE )
GO_BP_Cats_mRNA_AI_NP<-goseq(pwf,gene2cat=annotation.GO.BP.biomart, method ="Sampling", repcnt = 10000, use_genes
_without_cat=FALSE)
GO_BP_Cats_mRNA_AI_NP<-GO_BP_Cats_mRNA_AI_NP[GO_BP_Cats_mRNA_AI_NP$numDEInCat>4,]
GO_BP_Cats_mRNA_AI_NP$BY_FDR<-p.adjust(GO_BP_Cats_mRNA_AI_NP$over_represented_pvalue, method ="fdr")
GO_BP_Cats_mRNA_AI_NP<-GO_BP_Cats_mRNA_AI_NP[with(GO_BP_Cats_mRNA_AI_NP, order(BY_FDR,over_represented_pvalue, -n
umDEInCat)), ]
#head(GO_BP_Cats_mRNA_AI_NP, n=20)

GO_BP_Cats_mRNA_AI_NP$fold_enrichment<-(GO_BP_Cats_mRNA_AI_NP$numDEInCat/N_DEGs)/(GO_BP_Cats_mRNA_AI_NP$numInCat/
N_expressed_genes)
annotation.GO.BP.biomart_testgenes<-annotation.GO.BP.biomart[annotation.GO.BP.biomart$ensembl_gene_id %in% test.g
enes$a, ]
GO_BP_Cats_mRNA_AI_NP<-merge(GO_BP_Cats_mRNA_AI_NP,annotation.GO.BP.biomart_testgenes, by.x="category", by.y="go_
id", all.x=TRUE, all.y=FALSE)
GO_BP_Cats_mRNA_AI_NP<-merge(GO_BP_Cats_mRNA_AI_NP, annotation.ensembl.symbol, by.x="ensembl_gene_id", by.y="ense
mbl_gene_id", all=FALSE, all.x=TRUE, all.y=FALSE)
GO_BP_Cats_mRNA_AI_NP<-GO_BP_Cats_mRNA_AI_NP[GO_BP_Cats_mRNA_AI_NP$BY_FDR<0.1,]
GO_BP_Cats_mRNA_AI_NP<-GO_BP_Cats_mRNA_AI_NP[with(GO_BP_Cats_mRNA_AI_NP, order(BY_FDR,term)), ]
#write.table(GO_BP_Cats_mRNA_AI_NP, paste(result_path, "2019_12_27_GO_BP_Cats_DEG_mRNA_AI_NP.txt",sep="/"), quote
=FALSE, sep="\t", row.names=FALSE)
```

```
knitr::kable(head(GO_BP_Cats_mRNA_AI_NP), caption = "First five rows GO-BP analysis DEGs",format = 'pandoc')
```

First five rows GO-BP analysis DEGs

| ensembl_gene_id | category | over_represented_pvalue | under_represented_pvalue | numDEInCat | numInCat | term | ontology | BY_FD |
|---|---|---|---|---|---|---|---|---|
| ENSBTAG00000005018 | GO:0007165 (GO:0007165) | 0.0457954 | 0.9853015 | 5 | 348 | signal transduction | BP | 0.045795 |
| ENSBTAG00000010447 | GO:0007165 (GO:0007165) | 0.0457954 | 0.9853015 | 5 | 348 | signal transduction | BP | 0.045795 |
| ENSBTAG00000012500 | GO:0007165 (GO:0007165) | 0.0457954 | 0.9853015 | 5 | 348 | signal transduction | BP | 0.045795 |
| ENSBTAG00000015938 | GO:0007165 (GO:0007165) | 0.0457954 | 0.9853015 | 5 | 348 | signal transduction | BP | 0.045795 |

| ensembl_gene_id | category | over_represented_pvalue | under_represented_pvalue | numDEInCat | numInCat | term | ontology | BY_FD |
|---|---|---|---|---|---|---|---|---|
| ENSBTAG00000040188 | GO:0007165 (GO:0007165) | 0.0457954 | 0.9853015 | 5 | 348 | signal transduction | BP | 0.045795 |

Test molecular functions for enrichment with DEGs and calculate fold change enrichment. (AI Pregnant vs non pregnant)

```
set.seed(9830)
pwf<-nullp(all_genes_numeric, bias.data=annotation.genelength.biomart_vector, plot.fit=FALSE )
GO_MF_Cats_mRNA_AI_NP<-goseq(pwf,gene2cat=annotation.GO.MF.biomart, method ="Sampling", repcnt = 10000, use_genes
_without_cat=FALSE)
GO_MF_Cats_mRNA_AI_NP<-GO_MF_Cats_mRNA_AI_NP[GO_MF_Cats_mRNA_AI_NP$numDEInCat>4,]
GO_MF_Cats_mRNA_AI_NP$BY_FDR<-p.adjust(GO_MF_Cats_mRNA_AI_NP$over_represented_pvalue, method ="fdr")
GO_MF_Cats_mRNA_AI_NP<-GO_MF_Cats_mRNA_AI_NP[with(GO_MF_Cats_mRNA_AI_NP, order(BY_FDR,over_represented_pvalue, -n
umDEInCat)), ]
#head(GO_MF_Cats_mRNA_AI_NP, n=20)

GO_MF_Cats_mRNA_AI_NP$fold_enrichment<-(GO_MF_Cats_mRNA_AI_NP$numDEInCat/N_DEGs)/(GO_MF_Cats_mRNA_AI_NP$numInCat/
N_expressed_genes)
annotation.GO.MF.biomart_testgenes<-annotation.GO.MF.biomart[annotation.GO.MF.biomart$ensembl_gene_id %in% test.g
enes$a, ]
GO_MF_Cats_mRNA_AI_NP<-merge(GO_MF_Cats_mRNA_AI_NP,annotation.GO.MF.biomart_testgenes, by.x="category", by.y="go_
id", all.x=TRUE, all.y=FALSE)
GO_MF_Cats_mRNA_AI_NP<-merge(GO_MF_Cats_mRNA_AI_NP, annotation.ensembl.symbol, by.x="ensembl_gene_id", by.y="ense
mbl_gene_id", all=FALSE, all.x=TRUE, all.y=FALSE)
GO_MF_Cats_mRNA_AI_NP<-GO_MF_Cats_mRNA_AI_NP[GO_MF_Cats_mRNA_AI_NP$BY_FDR<0.1,]
GO_MF_Cats_mRNA_AI_NP<-GO_MF_Cats_mRNA_AI_NP[with(GO_MF_Cats_mRNA_AI_NP, order(BY_FDR,term)), ]
#write.table(GO_MF_Cats_mRNA_AI_NP, paste(result_path, "2019_10_10_GO_MF_Cats_mRNA_AI_NP.txt",sep="/"), quote=FAL
SE, sep="\t", row.names=FALSE)
```

Pathway analysis through KEGG Pathways. (AI Pregnant vs non pregnant)

```
all_genes_entrez<- stack(mget(all_genes$gene, org.Bt.egENSEMBL2EG, ifnotfound = NA))
all_genes_entrez<-all_genes_entrez[complete.cases(all_genes_entrez),]
all_genes_entrez<-all_genes_entrez[!duplicated(all_genes_entrez$values),]

test.genes_entrez <- stack(mget(test.genes$a, org.Bt.egENSEMBL2EG, ifnotfound = NA))
test.genes_entrez<-test.genes_entrez[complete.cases(test.genes_entrez),]

all_genes_entrez_numeric<-as.integer(all_genes_entrez$values %in% test.genes_entrez$values)
names(all_genes_entrez_numeric)<-all_genes_entrez$values

entrez_kegg<- stack(mget(all_genes_entrez$values, org.Bt.egPATH, ifnotfound = NA))
entrez_kegg<-entrez_kegg[complete.cases(entrez_kegg),]
entrez_kegg<-entrez_kegg[,c(2,1)]
entrez_kegg_test_genes<-entrez_kegg[entrez_kegg$ind %in% test.genes_entrez$values,]


pwf<-nullp(all_genes_entrez_numeric, 'bosTau4','refGene', plot.fit=FALSE )

set.seed(8503)
kegg_AI_NP_subset<-goseq(pwf, gene2cat=entrez_kegg,method ="Sampling", repcnt = 10000, use_genes_without_cat=FALS
E)
kegg_AI_NP_subset<-kegg_AI_NP_subset[kegg_AI_NP_subset$numDEInCat>4,]
kegg_AI_NP_subset$BY_FDR<-p.adjust(kegg_AI_NP_subset$over_represented_pvalue, method ="fdr")
kegg_AI_NP_subset<-merge(kegg_AI_NP_subset,entrez_kegg_test_genes, by.x="category", by.y="values", all.x=TRUE, al
l.y=FALSE)
kegg_AI_NP_subset<-merge(kegg_AI_NP_subset,all_genes_entrez, by.x="ind", by.y="values", all.x=TRUE, all.y=FALSE,
 suffixes =c('.entrez','.ensembl'))
kegg_AI_NP_subset<-kegg_AI_NP_subset[with(kegg_AI_NP_subset, order(BY_FDR, -numDEInCat)), ]
kegg_AI_NP_subset<-kegg_AI_NP_subset[,c(2:8)]
kegg_AI_NP_subset<-merge(kegg_AI_NP_subset, annotation.ensembl.symbol, by.x="ind.ensembl", by.y="ensembl_gene_id"
, all.x=TRUE, all.y=FALSE)
kegg_AI_NP_subset<-kegg_AI_NP_subset[with(kegg_AI_NP_subset, order(BY_FDR, -numDEInCat)), ]
kegg_AI_NP_subset<-kegg_AI_NP_subset[kegg_AI_NP_subset$BY_FDR<0.1,]
```

### DEG Analysis for mRNA

### AI Pregnant vs Pregnant to Natural Service

#### Results from edgeR

```
#colnames(count_pwbc)
group<-factor(c("Preg_AI","Preg_AI","Preg_AI","Preg_AI","Preg_AI","Preg_AI","Preg_NB","Preg_NB","Preg_NB","Preg_N
B","Preg_NB","Preg_NB","Not_Preg","Not_Preg","Not_Preg","Not_Preg","Not_Preg"), levels=c("Preg_AI","Preg_NB", "No
t_Preg"))
design<-model.matrix(~group)
dds<-DGEList(count=count_pwbc, group=group)
dds<-estimateDisp(dds, design, robust=TRUE)
dds<-glmFit(dds, design)
dds<-glmLRT(dds,coef="groupPreg_NB")
res_preg_AI_preg_NB_mRNA_edgeR<-topTags(dds,n=Inf)$table
#head(res_preg_AI_preg_NB_mRNA_edgeR,n=30)
#write.table(res_preg_AI_preg_NB_mRNA_edgeR, paste(result_path, "2019_04_30_res_preg_AI_preg_NB_mRNA_edgeR.txt"),
quote=FALSE, sep="\t", row.names=FALSE)
```

#### Results from DESeq

```
group<-factor(c("Preg_AI","Preg_AI","Preg_AI","Preg_AI","Preg_AI","Preg_AI","Preg_NB","Preg_NB","Preg_NB","Preg_N
B","Preg_NB","Preg_NB","Not_Preg","Not_Preg","Not_Preg","Not_Preg","Not_Preg"), levels=c("Preg_AI","Preg_NB", "No
t_Preg"))
design<-model.matrix(~group)
colData<-data.frame("group"=group)
rownames(colData)<-colnames(count_pwbc)
dds<-DESeqDataSetFromMatrix(countData=count_pwbc, colData=colData, design= ~group)
dds<-DESeq(dds)
res_preg_AI_preg_NB_mRNA_DeSeq<-results(dds, contrast=c("group","Preg_NB" ,"Preg_AI"), pAdjustMethod="fdr", tidy=
TRUE)
res_preg_AI_preg_NB_mRNA_DeSeq<-res_preg_AI_preg_NB_mRNA_DeSeq[with(res_preg_AI_preg_NB_mRNA_DeSeq, order(pvalu
e)), ]
#head(res_preg_AI_preg_NB_mRNA_DeSeq,n=30)
#write.table(res_preg_AI_preg_NB_mRNA_DeSeq, paste(result_path, "2019_04_30_res_preg_AI_preg_NB_mRNA_DeSeq.txt",
 sep="/"), quote=FALSE, sep="\t", row.names=FALSE)
```

Calculation of the eFDR for mRNA comparisons

```
#Create the permutations matrix
perm_without_replacement <- function(n, r){
  return(factorial(n)/factorial(n - r))
}

permutation_matrix<-matrix(,nrow=200100,ncol=17)
head(permutation_matrix)
dim(permutation_matrix)

for (i in seq(1:200100)){
  sampling <- sample(1:17,17, replace = FALSE)
  if ( ! identical(sampling , c(1:17))){
    permutation_matrix[i,]<-sample(1:17,17, replace = FALSE)
  }}

permutation_matrix<-permutation_matrix[!duplicated(permutation_matrix),]
dim(permutation_matrix) #number changes each time
head(permutation_matrix)

permutation_matrix<-permutation_matrix[sample(1:dim(permutation_matrix)[1], 10000),]   #should this number for th
e dim of the earlier
rand<-dim(permutation_matrix)[1]

sequence.pvalue<-seq(0.005, 0.05, 0.005)

#edgeR
results <- filebacked.big.matrix(length(sequence.pvalue),rand, type="double", init=0, separated=FALSE,
                                 backingfile="incidence_matrix.bin",
                                 descriptor="incidence_matrix.desc")  #this does not overwrite, create once and d
elete if want to create again
mdesc_result<- describe(results)

#edgeR
cl <- makeCluster(10)
registerDoParallel(cl)

results[,]<-foreach(i = sequence.pvalue, .combine='rbind', .inorder=TRUE,  .packages=c("edgeR","bigmemory"), .ver
bose=FALSE) %:%

  foreach(j = 1:rand, .combine='cbind', .inorder=FALSE,.packages=c("edgeR","bigmemory"), .verbose=FALSE ) %dopar%
{

    group<-factor(c("Preg_AI","Preg_AI","Preg_AI","Preg_AI","Preg_AI","Preg_AI","Preg_NB","Preg_NB","Preg_NB","Pr
eg_NB","Preg_NB","Preg_NB","Not_Preg","Not_Preg","Not_Preg","Not_Preg","Not_Preg"), levels=c("Preg_AI","Preg_NB",
"Not_Preg"))
    group<-group[permutation_matrix[j,]]
    group<-factor(group, levels=c("Preg_AI","Not_Preg", "Preg_NB"))
    design <- model.matrix(~group)
    dds<-DGEList(count=count_pwbc, group=group)
    dds<-estimateDisp(dds, design, robust=TRUE)
    dds <- glmFit(dds, design)
    dds <- glmLRT(dds, coef="groupPreg_NB")
    length(which(topTags(dds,n=Inf)$table$PValue < i))

  }

stopCluster(cl)

results[1:5,1:5]

total.rand <- rand * 10496

qvalue<-data.frame(raw.pvalue = sequence.pvalue,
                   e.pvalue.edgeR= (rowSums(results[,]+1))/(total.rand+1),
                   e.pvalue.edgeR.round= round((rowSums(results[,]+1))/(total.rand+1) ,4))

qvalue

rm(results)
system("rm incidence_matrix.bin")
system("rm incidence_matrix.desc")

#DESEQ2

results.rand <- filebacked.big.matrix(length(sequence.pvalue),rand, type="double", init=0, separated=FALSE,
                                      backingfile="incidence_matrix.bin",
                                      descriptor="incidence_matrix.desc")
mdesc_result<- describe(results.rand)

cl <- makeCluster(10)
registerDoParallel(cl)
```

```
results.rand[,]<-foreach(i = sequence.pvalue, .combine='rbind', .inorder=TRUE,  .packages=c("DESeq2","bigmemory"
), .verbose=TRUE) %:%

  foreach(j = 1:rand, .combine='cbind', .inorder=FALSE,.packages=c("DESeq2","bigmemory"), .verbose=TRUE ) %dopar%
{

    group<- c("Preg_AI","Preg_AI","Preg_AI","Preg_AI","Preg_AI","Preg_AI","Preg_NB","Preg_NB","Preg_NB","Preg_NB"
,"Preg_NB","Preg_NB","Not_Preg","Not_Preg","Not_Preg","Not_Preg","Not_Preg")
    group<-group[permutation_matrix[j,]]
    group<-factor(group, levels=c("Preg_AI","Not_Preg", "Preg_NB"))
    design <- model.matrix(~group)

    colData<-data.frame("group"=group)
    rownames(colData)<-colnames(count_pwbc)
    dds<-DESeqDataSetFromMatrix(countData=count_pwbc, colData=colData, design= ~group)
    dds <- DESeq(dds)
    DeSeq_AI_NP <- results(dds, contrast=c("group","Preg_NB" ,"Preg_AI"), pAdjustMethod="fdr", tidy=TRUE)

    length(which(DeSeq_AI_NP$pvalue < i))

  }

stopCluster(cl)

results.rand[1:5,1:5]

total.rand <- rand * 10496


qvalue2<-data.frame(raw.pvalue = sequence.pvalue,
                    e.pvalue.DESEQ2= (rowSums(results.rand[,]+1))/(total.rand+1),
                    e.pvalue.DESEQ2.round= round((rowSums(results.rand[,]+1))/(total.rand+1) ,4))

qvalue2

qvalue3<-cbind(qvalue,qvalue2)

qvalue3

rm(results.rand)
system("rm incidence_matrix.bin")
system("rm incidence_matrix.desc")

write.table(qvalue3,paste(result_path,"2019_12_27_empirical_FDR_AI_NB.txt",sep="/"), quote=FALSE, sep="\t", row.n
ames=FALSE)
```

Merge the results for edgeR and DESeq. (AI Pregnant vs Pregnant to Natural Service)

```
merged_res_preg_AI_preg_NB_mRNA<-merge(res_preg_AI_preg_NB_mRNA_edgeR, res_preg_AI_preg_NB_mRNA_DeSeq, by.x="row.
names", by.y="row")
merged_res_preg_AI_preg_NB_mRNA<-merged_res_preg_AI_preg_NB_mRNA[merged_res_preg_AI_preg_NB_mRNA$PValue<=0.03 & m
erged_res_preg_AI_preg_NB_mRNA$pvalue<=0.03,]
#dim(merged_res_preg_AI_preg_NB_mRNA) #12
merged_res_preg_AI_preg_NB_mRNA<-merged_res_preg_AI_preg_NB_mRNA[with(merged_res_preg_AI_preg_NB_mRNA, order(logF
C)), ]
#head(merged_res_preg_AI_preg_NB_mRNA)
DEG_AI_NB_mRNA<-merge(merged_res_preg_AI_preg_NB_mRNA,annotation.ensembl.symbol, by.x="Row.names", by.y="ensembl_
gene_id", all=FALSE)
DEG_AI_NB_mRNA<-DEG_AI_NB_mRNA[with(DEG_AI_NB_mRNA, order(logFC)), ]
#write.table(merged_res_preg_AI_preg_NB_mRNA, paste(result_path, "2019_04_30_merged_res_preg_AI_preg_NB_mRNA_befo
re_LOOCV.txt"), quote=FALSE, sep="\t", row.names=FALSE)
#write.table(DEG_AI_NB_mRNA, paste(result_path, "2019_12_27_merged_AI_NB_mRNA_gene_names_before_LOOCV.txt", sep
="/"), quote=FALSE, sep="\t", row.names=FALSE)
```

Plot the charts for all genes inferred as differentially expressed between AI and NB pregnant

```
cpm_AIpreg<-cpm_pwbc[,c(1,2,3,4,5,6)]
cpm_NOTpreg<-cpm_pwbc[,c(13,14,15,16,17)]
cpm_NBpreg<-cpm_pwbc[,c(7,8,9,10,11,12)]

data_chart_1<-data.frame( stringsAsFactors=FALSE)
data_chart_2<-data.frame( stringsAsFactors=FALSE)

for (i in  seq(dim(DEG_AI_NB_mRNA)[1])){

  gene_id<-DEG_AI_NB_mRNA[i,1]
  gene_symbol<-DEG_AI_NB_mRNA$external_gene_name[i]
  if(gene_symbol==""){gene_symbol<-gene_id}

  cpm_AI<-cpm_AIpreg[rownames(cpm_AIpreg)==gene_id,]
  cpm_NB<-cpm_NBpreg[rownames(cpm_NBpreg)==gene_id,]

  data_chart_1<-rbind(data.frame(cpm=cpm_AI), data.frame(cpm=cpm_NB))
  data_chart_1$group<-rep(c("Preg_AI","Preg_NB"),c(6,6))
  data_chart_1$shape<-as.factor(c(0:11))
  data_chart_1$gene<-gene_symbol
  data_chart_1$chart<-i
  data_chart_1$fold_change<-DEG_AI_NB_mRNA[DEG_AI_NB_mRNA$Row.names==gene_id,]$logFC
  data_chart_1$pvalue<-DEG_AI_NB_mRNA[DEG_AI_NB_mRNA$Row.names==gene_id,]$PValue


  data_chart_2<-rbind(data_chart_2,data_chart_1)
}

pdf(file=paste(result_path,"2019_12_27_plot_all_DEGs_preg_AI_preg_NB_ordered_by_FC_before_LOOCV.pdf",sep="/"), wi
dth= 11, height=4, bg="transparent", onefile=TRUE)

for (i in seq(1,dim(DEG_AI_NB_mRNA)[1], 24)) {
  plots <- list()
  k<-1
  for (j in c(i:(i+11))){
    data_chart_3<-data_chart_2[data_chart_2$chart %in% j , ]
    plot<-ggplot(data=data_chart_3, aes(x=group , y=cpm, colour=group))+
      geom_jitter(position = position_jitter(width = .1, height=0), aes( shape=shape), size=2)+
      scale_shape_manual(values=c(0:11))+
      scale_x_discrete(data_chart_3$gene[1],labels=c("Preg_AI"="Preg_AI", "Preg_NB"="Preg_NB"))+
      scale_colour_manual(name=NULL, values = c("#0072B2", "#000000"))+
      stat_summary(fun.y = median, fun.ymin = median, fun.ymax = median,colour = "gray", size = 0.2,  geom = "cro
ssbar", alpha=0.5)+
      scale_y_continuous("CPM")+
      ggtitle(paste("LogFC","=", round(data_chart_3$fold_change[1],1),"  ","P","=",round(data_chart_3$pvalue[1],4
), sep=c("")))+
      theme_bw()+
      theme(panel.grid= element_blank(),
            panel.background = element_blank(),
            panel.grid.minor = element_blank(),
            panel.grid.major = element_blank(),
            plot.background = element_blank(),
            strip.background = element_rect(fill = "white"),
            #strip.text.x = element_text(colour = 'black', face="italic",size = 9),
            #legend.text = element_text( colour = 'black', size = 10 ),
            axis.text.x = element_blank(),
            axis.text.y = element_text( colour = 'black', size = 8 ),
            axis.title = element_text( colour = 'black', size = 9 ),
            legend.key.size = unit(0.9,"cm"),
            legend.position="none",
            axis.ticks.x = element_blank(),
            plot.title = element_text(size = 7)
      )

    plots[[k]] <- plot
    k<-k+1
  }
  multiplot(plotlist = plots, cols = 6,layout = matrix(1:12, nrow=2, byrow = TRUE))
}
dev.off()
```

Leave-one-out cross validation

```
group<-factor(c("Preg_AI","Preg_AI","Preg_AI","Preg_AI","Preg_AI","Preg_AI","Preg_NB","Preg_NB","Preg_NB","Preg_N
B","Preg_NB","Preg_NB","Not_Preg","Not_Preg","Not_Preg","Not_Preg","Not_Preg"), levels=c("Preg_AI","Preg_NB", "No
t_Preg"))
rm(out_list_IDs_LOOCV,count_pwbc_a,group_a)

cl <- makeCluster(12)
registerDoParallel(cl)

out_list_IDs_LOOCV<-foreach(i = c(1:12), .inorder=FALSE,.packages=c("edgeR","DESeq2"), .verbose=FALSE ) %dopar% {

group_a<-group[-i]
count_pwbc_a<-count_pwbc[,-i]

design<-model.matrix(~group_a)
dds<-DGEList(count=count_pwbc_a, group=group_a)
dds<-estimateDisp(dds, design, robust=TRUE)
dds<-glmFit(dds, design)
dds<-glmLRT(dds,coef="group_aPreg_NB")
res_preg_AI_preg_NB_edgeR<-topTags(dds,n=Inf)$table

#DESeq
colData<-data.frame("group"=group_a)
rownames(colData)<-colnames(count_pwbc_a)
dds<-DESeqDataSetFromMatrix(countData=count_pwbc_a, colData=colData, design= ~group)
dds<-DESeq(dds)
res_preg_AI_preg_NB_DeSeq<-results(dds, contrast=c("group","Preg_NB" ,"Preg_AI"), pAdjustMethod="fdr", tidy=TRUE)


merged_res_preg_AI_preg_NB_mRNA_a<-merge(res_preg_AI_preg_NB_edgeR, res_preg_AI_preg_NB_DeSeq, by.x="row.names",
 by.y="row")
merged_res_preg_AI_preg_NB_mRNA_a<-merged_res_preg_AI_preg_NB_mRNA_a[merged_res_preg_AI_preg_NB_mRNA_a$PValue<=0.
03 & merged_res_preg_AI_preg_NB_mRNA_a$pvalue<=0.03,]

rm(group_a,count_pwbc_a)

merged_res_preg_AI_preg_NB_mRNA_a$Row.names
}
stopCluster(cl)

overlapping_genes_LOOCV<-Reduce(intersect, out_list_IDs_LOOCV)

#merged_res_preg_AI_preg_NB_mRNA<-merge(res_preg_AI_preg_NB_edgeR, res_preg_AI_preg_NB_DeSeq, by.x="row.names", b
y.y="row")
merged_res_preg_AI_preg_NB_mRNA_a<-merged_res_preg_AI_preg_NB_mRNA[merged_res_preg_AI_preg_NB_mRNA$Row.names %in%
overlapping_genes_LOOCV,]
dim(merged_res_preg_AI_preg_NB_mRNA_a) #2
```

```
## [1]  2 12
```

```
merged_res_preg_AI_preg_NB_mRNA_a<-merged_res_preg_AI_preg_NB_mRNA_a[with(merged_res_preg_AI_preg_NB_mRNA_a, orde
r(logFC)), ]
DEG_AI_NB_mRNA<-merge(merged_res_preg_AI_preg_NB_mRNA_a,annotation.ensembl.symbol, by.x="Row.names", by.y="ensemb
l_gene_id", all=FALSE)
DEG_AI_NB_mRNA<-DEG_AI_NB_mRNA[with(DEG_AI_NB_mRNA, order(logFC)), ]
dim(DEG_AI_NB_mRNA)
```

```
## [1]  2 17
```

```
#head(DEG_AI_NB_mRNA)
#write.table(DEG_AI_NB_mRNA, paste(result_path, "2019_12_27_merged_AI_NB_mRNA_gene_names_after_LOOCV.txt", sep
="/"), quote=FALSE, sep="\t", row.names=FALSE)
rm(out_list_IDs_LOOCV)
```

```
knitr::kable(head(DEG_AI_NB_mRNA), caption = "First five rows AI pregnant vs NB pregnant",format = 'pandoc')
```

First five rows AI pregnant vs NB pregnant

| | Row.names | logFC | logCPM | LR | PValue | FDR | baseMean | log2FoldChange | lfcSE | stat | pvalue | |
|---|---|---|---|---|---|---|---|---|---|---|---|---|
| 2 | ENSBTAG00000049416 | 2.888986 | 0.5520443 | 7.151642 | 0.0074895 | 0.9994483 | 31.42871 | 3.232006 | 1.130232 | 2.859596 | 0.0042418 | 0.999 |

| | Row.names | logFC | logCPM | LR | PValue | FDR | baseMean | log2FoldChange | lfcSE | stat | pvalue | |
|---|---|---|---|---|---|---|---|---|---|---|---|---|
| 1 | ENSBTAG00000040188 | 6.887927 | 0.8187581 | 12.657093 | 0.0003741 | 0.9994483 | 37.68545 | 7.300996 | 1.748430 | 4.175745 | 0.0000297 | 0.310[illegible] |

Plot the charts for all genes inferred as differentially expressed between AI and NB pregnant

```
cpm_AIpreg<-cpm_pwbc[,c(1,2,3,4,5,6)]
cpm_NOTpreg<-cpm_pwbc[,c(13,14,15,16,17)]
cpm_NBpreg<-cpm_pwbc[,c(7,8,9,10,11,12)]

data_chart_1<-data.frame( stringsAsFactors=FALSE)
data_chart_2<-data.frame( stringsAsFactors=FALSE)

for (i in  seq(dim(DEG_AI_NB_mRNA)[1])){

  gene_id<-DEG_AI_NB_mRNA[i,1]
  gene_symbol<-DEG_AI_NB_mRNA$external_gene_name[i]
  if(gene_symbol==""){gene_symbol<-gene_id}

  cpm_AI<-cpm_AIpreg[rownames(cpm_AIpreg)==gene_id,]
  cpm_NB<-cpm_NBpreg[rownames(cpm_NBpreg)==gene_id,]

  data_chart_1<-rbind(data.frame(cpm=cpm_AI), data.frame(cpm=cpm_NB))
  data_chart_1$group<-rep(c("Preg_AI","Preg_NB"),c(6,6))
  data_chart_1$shape<-as.factor(c(0:11))
  data_chart_1$gene<-gene_symbol
  data_chart_1$chart<-i
  data_chart_1$fold_change<-DEG_AI_NB_mRNA[DEG_AI_NB_mRNA$Row.names==gene_id,]$logFC
  data_chart_1$pvalue<-DEG_AI_NB_mRNA[DEG_AI_NB_mRNA$Row.names==gene_id,]$PValue


  data_chart_2<-rbind(data_chart_2,data_chart_1)
}

pdf(file=paste(result_path,"2019_12_27_plot_all_DEGs_preg_AI_preg_NB_ordered_by_FC_after_LOOCV.pdf",sep="/"), wid
th= 3,height=2, bg="transparent", onefile=TRUE)

# for (i in seq(1,dim(DEG_AI_NB_mRNA)[1], 24)) {
  plots <- list()
  k<-1
  for (j in c(i:2)){
    data_chart_3<-data_chart_2[data_chart_2$chart %in% j , ]
    plot<-ggplot(data=data_chart_3, aes(x=group , y=cpm, colour=group))+
      geom_jitter(position = position_jitter(width = .1, height=0), aes( shape=shape), size=2)+
      scale_shape_manual(values=c(0:11))+
      scale_x_discrete(data_chart_3$gene[1],labels=c("Preg_AI"="Preg_AI", "Preg_NB"="Preg_NB"))+
      scale_colour_manual(name=NULL, values = c("blue", "black"))+
      stat_summary(fun.y = median, fun.ymin = median, fun.ymax = median,colour = "gray", size = 0.2,  geom = "cro
ssbar", alpha=0.5)+
      scale_y_continuous("CPM")+
      ggtitle(paste("LogFC","=", round(data_chart_3$fold_change[1],1),"  ","P","=",round(data_chart_3$pvalue[1],4
), sep=c("")))+
      theme_bw()+
      theme(panel.grid= element_blank(),
            panel.background = element_blank(),
            panel.grid.minor = element_blank(),
            panel.grid.major = element_blank(),
            plot.background = element_blank(),
            strip.background = element_rect(fill = "white"),
            #strip.text.x = element_text(colour = 'black', face="italic",size = 9),
            #legend.text = element_text( colour = 'black', size = 10 ),
            axis.text.x = element_blank(),
            axis.text.y = element_text( colour = 'black', size = 8 ),
            axis.title = element_text( colour = 'black', size = 9 ),
            legend.key.size = unit(0.9,"cm"),
            legend.position="none",
            axis.ticks.x = element_blank(),
            plot.title = element_text(size = 7)
      )

    plots[[k]] <- plot
    k<-k+1
  }
  multiplot(plotlist = plots, cols = 2,layout = matrix(1:2, nrow=1, byrow = TRUE))
#}
dev.off()
```

### DEG Analysis for mRNA

### Pregnant to Natural Service vs non pregnant

#### Results from edgeR

```
#colnames(count_pwbc)
group<-factor(c("Preg_AI","Preg_AI","Preg_AI","Preg_AI","Preg_AI","Preg_AI","Preg_NB","Preg_NB","Preg_NB","Preg_N
B","Preg_NB","Preg_NB","Not_Preg","Not_Preg","Not_Preg","Not_Preg","Not_Preg"), levels=c("Preg_NB","Preg_AI", "No
t_Preg"))
design<-model.matrix(~group)
dds<-DGEList(count=count_pwbc, group=group)
dds<-estimateDisp(dds, design, robust=TRUE)
dds<-glmFit(dds, design)
dds<-glmLRT(dds,coef="groupNot_Preg")
res_not_preg_preg_NB_edgeR<-topTags(dds,n=Inf)$table
#head(res_not_preg_preg_NB_edgeR,n=30)
#write.table(res_not_preg_preg_NB_edgeR, paste(result_path, "2019_04_30_res_not_preg_preg_NB_edgeR.txt"), quote=F
ALSE, sep="\t", row.names=FALSE)
```

#### Results from DESeq

```
design<-model.matrix(~group)
colData<-data.frame("group"=group)
rownames(colData)<-colnames(count_pwbc)
dds<-DESeqDataSetFromMatrix(countData=count_pwbc, colData=colData, design= ~group)
dds<-DESeq(dds)
res_not_preg_preg_NB_DeSeq<-results(dds, contrast=c("group","Not_Preg" ,"Preg_NB"), pAdjustMethod="fdr", tidy=TRU
E)
res_not_preg_preg_NB_DeSeq<-res_not_preg_preg_NB_DeSeq[with(res_not_preg_preg_NB_DeSeq, order(pvalue)), ]
#head(res_not_preg_preg_NB_DeSeq,n=30)
#write.table(res_not_preg_preg_NB_DeSeq, paste(result_path, "2019_04_30_res_not_preg_preg_NB_DeSeq.txt"), quote=F
ALSE, sep="\t", row.names=FALSE)
```

Merge the results for edgeR and DESeq. (Pregnant to Natural Service vs non pregnant)

```
merged_res_not_preg_preg_NB_mRNA<-merge(res_not_preg_preg_NB_edgeR, res_not_preg_preg_NB_DeSeq, by.x="row.names",
by.y="row")
merged_res_not_preg_preg_NB_mRNA<-merged_res_not_preg_preg_NB_mRNA[merged_res_not_preg_preg_NB_mRNA$PValue<=0.03
 & merged_res_not_preg_preg_NB_mRNA$pvalue<=0.03,]
dim(merged_res_not_preg_preg_NB_mRNA) #414
```

```
## [1] 414  12
```

```
#head(merged_res_not_preg_preg_NB_mRNA)
DEG_NB_NP_mRNA<-merge(merged_res_not_preg_preg_NB_mRNA,annotation.ensembl.symbol, by.x="Row.names", by.y="ensembl
_gene_id", all=FALSE)
DEG_NB_NP_mRNA<-DEG_NB_NP_mRNA[with(DEG_NB_NP_mRNA, order(logFC)), ]
#write.table(merged_res_not_preg_preg_NB_mRNA, paste(result_path, "2019_04_30_merged_res_not_preg_preg_NB_mRNA_be
fore_LOOCV.txt"), quote=FALSE, sep="\t", row.names=FALSE)
#write.table(DEG_NB_NP_mRNA, paste(result_path, "2019_12_27_merged_NB_NP_mRNA_gene_names_before_LOOCV.txt", sep
="/"), quote=FALSE, sep="\t", row.names=FALSE)
```

Plot the charts for all genes inferred as differentially expressed between NB pregnant and non pregnant

```
cpm_AIpreg<-cpm_pwbc[,c(1,2,3,4,5,6)]
cpm_NOTpreg<-cpm_pwbc[,c(13,14,15,16,17)]
cpm_NBpreg<-cpm_pwbc[,c(7,8,9,10,11,12)]

data_chart_1<-data.frame( stringsAsFactors=FALSE)
data_chart_2<-data.frame( stringsAsFactors=FALSE)

for (i in  seq(dim(DEG_NB_NP_mRNA)[1])){

  gene_id<-DEG_NB_NP_mRNA[i,1]
  gene_symbol<-DEG_NB_NP_mRNA$external_gene_name[i]
  if(gene_symbol==""){gene_symbol<-gene_id}

  cpm_NB<-cpm_NBpreg[rownames(cpm_NBpreg)==gene_id,]
  cpm_NP<-cpm_NOTpreg[rownames(cpm_NOTpreg)==gene_id,]

  data_chart_1<-rbind(data.frame(cpm=cpm_NB), data.frame(cpm=cpm_NP))
  data_chart_1$group<-rep(c("Preg_NB","Not_Preg"),c(6,5))
  data_chart_1$shape<-as.factor(c(0:10))
  data_chart_1$gene<-gene_symbol
  data_chart_1$chart<-i
  data_chart_1$fold_change<-DEG_NB_NP_mRNA[DEG_NB_NP_mRNA$Row.names==gene_id,]$logFC
  data_chart_1$pvalue<-DEG_NB_NP_mRNA[DEG_NB_NP_mRNA$Row.names==gene_id,]$PValue


  data_chart_2<-rbind(data_chart_2,data_chart_1)
}

pdf(file=paste(result_path,"2019_12_27_plot_all_DEGs_preg_NB_not_preg_ordered_by_FC_before_LOOCV.pdf",sep="/"), w
idth= 11, paper="USr", bg="transparent", onefile=TRUE)

for (i in seq(1,dim(DEG_NB_NP_mRNA)[1], 24)) {
  plots <- list()
  k<-1
  for (j in c(i:(i+23))){
    data_chart_3<-data_chart_2[data_chart_2$chart %in% j , ]
    plot<-ggplot(data=data_chart_3, aes(x=group , y=cpm, colour=group))+
      geom_jitter(position = position_jitter(width = .1, height=0), aes( shape=shape), size=2)+
      scale_shape_manual(values=c(0:10))+
      scale_x_discrete(data_chart_3$gene[1],labels=c("Preg_NB"="Preg_NB", "Not_Preg"="Not_Preg"))+
      scale_colour_manual(name=NULL, values = c("#D55E00", "#000000"))+
      stat_summary(fun.y = median, fun.ymin = median, fun.ymax = median,colour = "gray", size = 0.2,  geom = "cro
ssbar", alpha=0.5)+
      scale_y_continuous("CPM")+
      ggtitle(paste("LogFC","=", round(data_chart_3$fold_change[1],1),"  ","P","=",round(data_chart_3$pvalue[1],4
), sep=c("")))+
      theme_bw()+
      theme(panel.grid= element_blank(),
            panel.background = element_blank(),
            panel.grid.minor = element_blank(),
            panel.grid.major = element_blank(),
            plot.background = element_blank(),
            strip.background = element_rect(fill = "white"),
            #strip.text.x = element_text(colour = 'black', face="italic",size = 9),
            #legend.text = element_text( colour = 'black', size = 10 ),
            axis.text.x = element_blank(),
            axis.text.y = element_text( colour = 'black', size = 8 ),
            axis.title = element_text( colour = 'black', size = 9 ),
            legend.key.size = unit(0.9,"cm"),
            legend.position="none",
            axis.ticks.x = element_blank(),
            plot.title = element_text(size = 7)
      )

    plots[[k]] <- plot
    k<-k+1
  }
  multiplot(plotlist = plots, cols = 6,layout = matrix(1:24, nrow=4, byrow = TRUE))
}
dev.off()
```

### Supplementary table 12

Leave-one-out cross validation

```
group<-factor(c("Preg_AI","Preg_AI","Preg_AI","Preg_AI","Preg_AI","Preg_AI","Preg_NB","Preg_NB","Preg_NB","Preg_N
B","Preg_NB","Preg_NB","Not_Preg","Not_Preg","Not_Preg","Not_Preg","Not_Preg"), levels=c("Preg_NB", "Preg_AI","No
t_Preg"))
rm(out_list_IDs_LOOCV,count_pwbc_a,group_a)

cl <- makeCluster(12)
registerDoParallel(cl)

out_list_IDs_LOOCV<-foreach(i = c(7:17), .inorder=FALSE,.packages=c("edgeR","DESeq2"), .verbose=FALSE ) %dopar% {

group_a<-group[-i]
count_pwbc_a<-count_pwbc[,-i]

design<-model.matrix(~group_a)
dds<-DGEList(count=count_pwbc_a, group=group_a)
dds<-estimateDisp(dds, design, robust=TRUE)
dds<-glmFit(dds, design)
dds<-glmLRT(dds,coef="group_aNot_Preg")
res_not_preg_preg_NB_edgeR<-topTags(dds,n=Inf)$table

#DESeq
colData<-data.frame("group"=group_a)
rownames(colData)<-colnames(count_pwbc_a)
dds<-DESeqDataSetFromMatrix(countData=count_pwbc_a, colData=colData, design= ~group)
dds<-DESeq(dds)
res_not_preg_preg_NB_DeSeq<-results(dds, contrast=c("group","Preg_NB" ,"Not_Preg"), pAdjustMethod="fdr", tidy=TRU
E)


merged_res_not_preg_preg_NB_mRNA_a<-merge(res_not_preg_preg_NB_edgeR, res_not_preg_preg_NB_DeSeq, by.x="row.name
s", by.y="row")
merged_res_not_preg_preg_NB_mRNA_a<-merged_res_not_preg_preg_NB_mRNA_a[merged_res_not_preg_preg_NB_mRNA_a$PValue<
=0.03 & merged_res_not_preg_preg_NB_mRNA_a$pvalue<=0.03,]

rm(group_a,count_pwbc_a)

merged_res_not_preg_preg_NB_mRNA_a$Row.names
}
stopCluster(cl)
overlapping_genes_LOOCV<-Reduce(intersect, out_list_IDs_LOOCV)

merged_res_not_preg_preg_NB_mRNA<-merge(res_not_preg_preg_NB_edgeR, res_not_preg_preg_NB_DeSeq, by.x="row.names",
by.y="row")
merged_res_not_preg_preg_NB_mRNA_a<-merged_res_not_preg_preg_NB_mRNA[merged_res_not_preg_preg_NB_mRNA$Row.names %
in% overlapping_genes_LOOCV,]
dim(merged_res_not_preg_preg_NB_mRNA_a) #81
```

```
## [1] 81 12
```

```
DEG_NB_NP_mRNA<-merge(merged_res_not_preg_preg_NB_mRNA_a,annotation.ensembl.symbol, by.x="Row.names", by.y="ensem
bl_gene_id", all=FALSE)
DEG_NB_NP_mRNA<-DEG_NB_NP_mRNA[with(DEG_NB_NP_mRNA, order(logFC)), ]
#write.table(DEG_NB_NP_mRNA, paste(result_path, "2019_12_27_merged_NB_NP_mRNA_gene_names_after_LOOCV.txt", sep
="/"), quote=FALSE, sep="\t", row.names=FALSE)
rm(out_list_IDs_LOOCV)
```

```
knitr::kable(head(DEG_NB_NP_mRNA), caption = "First five rows Supplementary table 12",format = 'pandoc')
```

First five rows Supplementary table 12

| | Row.names | logFC | logCPM | LR | PValue | FDR | baseMean | log2FoldChange | lfcSE | stat | pvalue | |
|---|---|---|---|---|---|---|---|---|---|---|---|---|
| 79 | ENSBTAG00000052118 | -4.492594 | 0.1925364 | 8.266168 | 0.0040391 | 0.5315921 | 17.10857 | -4.674455 | 1.5654939 | -2.985930 | 0.0028272 | 0. |
| 3 | ENSBTAG00000000517 | -4.008047 | 1.6725720 | 13.410263 | 0.0002503 | 0.3283297 | 56.86040 | -3.734326 | 0.9896552 | -3.773360 | 0.0001611 | 0. |
| 59 | ENSBTAG00000034281 | -3.406753 | 1.0671814 | 8.971424 | 0.0027423 | 0.5315921 | 36.88080 | -3.330845 | 1.0879922 | -3.061460 | 0.0022026 | 0. |
| 41 | ENSBTAG00000014791 | -2.644484 | 1.6574465 | 7.079575 | 0.0077967 | 0.5464846 | 59.61641 | -2.916556 | 0.9095127 | -3.206724 | 0.0013426 | 0. |

| | Row.names | logFC | logCPM | LR | PValue | FDR | baseMean | log2FoldChange | lfcSE | stat | pvalue | |
|---|---|---|---|---|---|---|---|---|---|---|---|---|
| 23 | ENSBTAG00000008227 | -2.366188 | 2.2527690 | 11.986830 | 0.0005358 | 0.3763416 | 88.74001 | -2.406923 | 0.6201177 | -3.881398 | 0.0001039 | 0. |
| 75 | ENSBTAG00000049285 | -2.111184 | 3.1991884 | 9.014732 | 0.0026781 | 0.5315921 | 177.45930 | -2.320851 | 0.6485780 | -3.578368 | 0.0003457 | 0. |

Plot the charts for all genes inferred as differentially expressed between NB pregnant and non pregnant

```
cpm_AIpreg<-cpm_pwbc[,c(1,2,3,4,5,6)]
cpm_NOTpreg<-cpm_pwbc[,c(13,14,15,16,17)]
cpm_NBpreg<-cpm_pwbc[,c(7,8,9,10,11,12)]

data_chart_1<-data.frame( stringsAsFactors=FALSE)
data_chart_2<-data.frame( stringsAsFactors=FALSE)

for (i in  seq(dim(DEG_NB_NP_mRNA)[1])){

  gene_id<-DEG_NB_NP_mRNA[i,1]
  gene_symbol<-DEG_NB_NP_mRNA$external_gene_name[i]
  if(gene_symbol==""){gene_symbol<-gene_id}

  cpm_NB<-cpm_NBpreg[rownames(cpm_NBpreg)==gene_id,]
  cpm_NP<-cpm_NOTpreg[rownames(cpm_NOTpreg)==gene_id,]

  data_chart_1<-rbind(data.frame(cpm=cpm_NB), data.frame(cpm=cpm_NP))
  data_chart_1$group<-rep(c("Preg_NB","Not_Preg"),c(6,5))
  data_chart_1$shape<-as.factor(c(0:10))
  data_chart_1$gene<-gene_symbol
  data_chart_1$chart<-i
  data_chart_1$fold_change<-DEG_NB_NP_mRNA[DEG_NB_NP_mRNA$Row.names==gene_id,]$logFC
  data_chart_1$pvalue<-DEG_NB_NP_mRNA[DEG_NB_NP_mRNA$Row.names==gene_id,]$PValue


  data_chart_2<-rbind(data_chart_2,data_chart_1)
}

pdf(file=paste(result_path,"2019_12_27_plot_all_DEGs_preg_NB_not_preg_ordered_by_FC_after_LOOCV.pdf",sep="/"), wi
dth= 11, paper="USr", bg="transparent", onefile=TRUE)

for (i in seq(1,dim(DEG_NB_NP_mRNA)[1], 24)) {
  plots <- list()
  k<-1
  for (j in c(i:(i+23))){
    data_chart_3<-data_chart_2[data_chart_2$chart %in% j , ]
    plot<-ggplot(data=data_chart_3, aes(x=group , y=cpm, colour=group))+
      geom_jitter(position = position_jitter(width = .1, height=0), aes( shape=shape), size=2)+
      scale_shape_manual(values=c(0:10))+
      scale_x_discrete(data_chart_3$gene[1],labels=c("Preg_NB"="Preg_NB", "Not_Preg"="Not_Preg"))+
      scale_colour_manual(name=NULL, values = c("#D55E00", "#000000"))+
      stat_summary(fun.y = median, fun.ymin = median, fun.ymax = median,colour = "gray", size = 0.2,  geom = "cro
ssbar", alpha=0.5)+
      scale_y_continuous("CPM")+
      ggtitle(paste("LogFC","=", round(data_chart_3$fold_change[1],1),"  ","P","=",round(data_chart_3$pvalue[1],4
), sep=c("")))+
      theme_bw()+
      theme(panel.grid= element_blank(),
            panel.background = element_blank(),
            panel.grid.minor = element_blank(),
            panel.grid.major = element_blank(),
            plot.background = element_blank(),
            strip.background = element_rect(fill = "white"),
            #strip.text.x = element_text(colour = 'black', face="italic",size = 9),
            #legend.text = element_text( colour = 'black', size = 10 ),
            axis.text.x = element_blank(),
            axis.text.y = element_text( colour = 'black', size = 8 ),
            axis.title = element_text( colour = 'black', size = 9 ),
            legend.key.size = unit(0.9,"cm"),
            legend.position="none",
            axis.ticks.x = element_blank(),
            plot.title = element_text(size = 7)
      )

    plots[[k]] <- plot
    k<-k+1
  }
  multiplot(plotlist = plots, cols = 6,layout = matrix(1:24, nrow=4, byrow = TRUE))
}
dev.off()
```

Test biological processes for enrichment with DEGs and calculate fold change enrichment. (Pregnant to Natural Service vs non pregnant)

```
test.genes<-data.frame(a=unique(c(as.character(merged_res_not_preg_preg_NB_mRNA_a$Row.names))), stringsAsFactors=
FALSE)
#(test.genes$a %in% all_genes$gene)
all_genes_numeric<-as.integer(all_genes$gene %in%test.genes$a)
names(all_genes_numeric)<-all_genes$gene
N_DEGs<-length(test.genes$a)

set.seed(9830)
pwf<-nullp(all_genes_numeric, bias.data=annotation.genelength.biomart_vector, plot.fit=FALSE )
GO_BP_Cats_mRNA_NB_NP<-goseq(pwf,gene2cat=annotation.GO.BP.biomart, method ="Sampling", repcnt = 7000, use_genes_
without_cat=FALSE)
GO_BP_Cats_mRNA_NB_NP<-GO_BP_Cats_mRNA_NB_NP[GO_BP_Cats_mRNA_NB_NP$numDEInCat>4,]
GO_BP_Cats_mRNA_NB_NP$BY_FDR<-p.adjust(GO_BP_Cats_mRNA_NB_NP$over_represented_pvalue, method ="fdr")
GO_BP_Cats_mRNA_NB_NP<-GO_BP_Cats_mRNA_NB_NP[with(GO_BP_Cats_mRNA_NB_NP, order(BY_FDR,over_represented_pvalue, -n
umDEInCat)), ]
#head(GO_BP_Cats_mRNA_NB_NP, n=20)

GO_BP_Cats_mRNA_NB_NP$fold_enrichment<-(GO_BP_Cats_mRNA_NB_NP$numDEInCat/N_DEGs)/(GO_BP_Cats_mRNA_NB_NP$numInCat/
N_expressed_genes)
annotation.GO.BP.biomart_testgenes<-annotation.GO.BP.biomart[annotation.GO.BP.biomart$ensembl_gene_id %in% test.g
enes$a, ]
GO_BP_Cats_mRNA_NB_NP<-merge(GO_BP_Cats_mRNA_NB_NP,annotation.GO.BP.biomart_testgenes, by.x="category", by.y="go_
id", all.x=TRUE, all.y=FALSE)
GO_BP_Cats_mRNA_NB_NP<-merge(GO_BP_Cats_mRNA_NB_NP, annotation.ensembl.symbol, by.x="ensembl_gene_id", by.y="ense
mbl_gene_id", all=FALSE, all.x=TRUE, all.y=FALSE)
GO_BP_Cats_mRNA_NB_NP<-GO_BP_Cats_mRNA_NB_NP[with(GO_BP_Cats_mRNA_NB_NP, order(BY_FDR,term)), ]
GO_BP_Cats_mRNA_NB_NP<-GO_BP_Cats_mRNA_NB_NP[GO_BP_Cats_mRNA_NB_NP$BY_FDR <0.1, ]
#write.table(GO_BP_Cats_mRNA_NB_NP, paste(result_path, "2019_05_07_GO_BP_Cats_mRNA_NB_NP.txt", sep="/"), quote=FA
LSE, sep="\t", row.names=FALSE)
```

```
knitr::kable(head(GO_BP_Cats_mRNA_NB_NP), caption = "First five rows",format = 'pandoc')
```

First five rows

| | ensembl_gene_id | category | over_represented_pvalue | under_represented_pvalue | numDEInCat | numInCat | term | ontology | BY[illegible] |
|---|---|---|---|---|---|---|---|---|---|
| 2 | ENSBTAG00000004211 | GO:0007165 (GO:0007165) | 0.0309956 | 0.9901443 | 6 | 348 | signal transduction | BP | 0.06[illegible] |
| 3 | ENSBTAG00000010047 | GO:0007165 (GO:0007165) | 0.0309956 | 0.9901443 | 6 | 348 | signal transduction | BP | 0.06[illegible] |
| 4 | ENSBTAG00000010447 | GO:0007165 (GO:0007165) | 0.0309956 | 0.9901443 | 6 | 348 | signal transduction | BP | 0.06[illegible] |
| 5 | ENSBTAG00000015938 | GO:0007165 (GO:0007165) | 0.0309956 | 0.9901443 | 6 | 348 | signal transduction | BP | 0.06[illegible] |
| 10 | ENSBTAG00000044071 | GO:0007165 (GO:0007165) | 0.0309956 | 0.9901443 | 6 | 348 | signal transduction | BP | 0.06[illegible] |
| 11 | ENSBTAG00000049285 | GO:0007165 (GO:0007165) | 0.0309956 | 0.9901443 | 6 | 348 | signal transduction | BP | 0.06[illegible] |

Test molecular functions for enrichment with DEGs and calculate fold change enrichment. (Pregnant to Natural Service vs non pregnant)

```
set.seed(9830)
pwf<-nullp(all_genes_numeric, bias.data=annotation.genelength.biomart_vector, plot.fit=FALSE )
GO_MF_Cats_mRNA_NB_NP<-goseq(pwf,gene2cat=annotation.GO.MF.biomart, method ="Sampling", repcnt = 7000, use_genes_
without_cat=FALSE)
GO_MF_Cats_mRNA_NB_NP<-GO_MF_Cats_mRNA_NB_NP[GO_MF_Cats_mRNA_NB_NP$numDEInCat>4,]
GO_MF_Cats_mRNA_NB_NP$BY_FDR<-p.adjust(GO_MF_Cats_mRNA_NB_NP$over_represented_pvalue, method ="fdr")
GO_MF_Cats_mRNA_NB_NP<-GO_MF_Cats_mRNA_NB_NP[with(GO_MF_Cats_mRNA_NB_NP, order(BY_FDR,over_represented_pvalue, -n
umDEInCat)), ]
#head(GO_MF_Cats_mRNA_NB_NP, n=20)

GO_MF_Cats_mRNA_NB_NP$fold_enrichment<-(GO_MF_Cats_mRNA_NB_NP$numDEInCat/N_DEGs)/(GO_MF_Cats_mRNA_NB_NP$numInCat/
N_expressed_genes)
annotation.GO.MF.biomart_testgenes<-annotation.GO.MF.biomart[annotation.GO.MF.biomart$ensembl_gene_id %in% test.g
enes$a, ]
GO_MF_Cats_mRNA_NB_NP<-merge(GO_MF_Cats_mRNA_NB_NP,annotation.GO.MF.biomart_testgenes, by.x="category", by.y="go_
id", all.x=TRUE, all.y=FALSE)
GO_MF_Cats_mRNA_NB_NP<-merge(GO_MF_Cats_mRNA_NB_NP, annotation.ensembl.symbol, by.x="ensembl_gene_id", by.y="ense
mbl_gene_id", all=FALSE, all.x=TRUE, all.y=FALSE)
GO_MF_Cats_mRNA_NB_NP<-GO_MF_Cats_mRNA_NB_NP[with(GO_MF_Cats_mRNA_NB_NP, order(BY_FDR,term)), ]
GO_MF_Cats_mRNA_NB_NP<-GO_MF_Cats_mRNA_NB_NP[GO_MF_Cats_mRNA_NB_NP$BY_FDR<0.1, ]
#write.table(GO_MF_Cats_mRNA_NB_NP, paste(result_path, "2019_05_07_GO_MF_Cats_mRNA_NB_NP.txt"), quote=FALSE, sep
="\t", row.names=FALSE)
```

Pathway analysis through KEGG Pathways. (Pregnant to Natural Service vs non pregnant)

```
all_genes_entrez<- stack(mget(all_genes$gene, org.Bt.egENSEMBL2EG, ifnotfound = NA))
all_genes_entrez<-all_genes_entrez[complete.cases(all_genes_entrez),]
all_genes_entrez<-all_genes_entrez[!duplicated(all_genes_entrez$values),]

test.genes_entrez <- stack(mget(test.genes$a, org.Bt.egENSEMBL2EG, ifnotfound = NA))
test.genes_entrez<-test.genes_entrez[complete.cases(test.genes_entrez),]

all_genes_entrez_numeric<-as.integer(all_genes_entrez$values %in% test.genes_entrez$values)
names(all_genes_entrez_numeric)<-all_genes_entrez$values

entrez_kegg<- stack(mget(all_genes_entrez$values, org.Bt.egPATH, ifnotfound = NA))
entrez_kegg<-entrez_kegg[complete.cases(entrez_kegg),]
entrez_kegg<-entrez_kegg[,c(2,1)]
entrez_kegg_test_genes<-entrez_kegg[entrez_kegg$ind %in% test.genes_entrez$values,]

pwf<-nullp(all_genes_entrez_numeric, 'bosTau4','refGene', plot.fit=FALSE )

set.seed(8503)
kegg_NB_NP_subset<-goseq(pwf, gene2cat=entrez_kegg,method ="Sampling", repcnt = 7000, use_genes_without_cat=FALSE
)
kegg_NB_NP_subset<-kegg_NB_NP_subset[kegg_NB_NP_subset$numDEInCat>4,]
kegg_NB_NP_subset$BY_FDR<-p.adjust(kegg_NB_NP_subset$over_represented_pvalue, method ="fdr")
kegg_NB_NP_subset<-merge(kegg_NB_NP_subset,entrez_kegg_test_genes, by.x="category", by.y="values", all.x=TRUE, al
l.y=FALSE)
kegg_NB_NP_subset<-merge(kegg_NB_NP_subset,all_genes_entrez, by.x="ind", by.y="values", all.x=TRUE, all.y=FALSE,
 suffixes =c('.entrez','.ensembl'))
kegg_NB_NP_subset<-kegg_NB_NP_subset[with(kegg_NB_NP_subset, order(BY_FDR, -numDEInCat)), ]
kegg_NB_NP_subset<-kegg_NB_NP_subset[,c(2:8)]
kegg_NB_NP_subset<-merge(kegg_NB_NP_subset, annotation.ensembl.symbol, by.x="ind.ensembl", by.y="ensembl_gene_id"
, all.x=TRUE, all.y=FALSE)
kegg_NB_NP_subset<-kegg_NB_NP_subset[with(kegg_NB_NP_subset, order(BY_FDR, -numDEInCat)), ]
kegg_NB_NP_subset<-kegg_NB_NP_subset[kegg_NB_NP_subset$BY_FDR<0.1,]
#write.table(kegg_NB_NP_subset, paste(result_path, "2019_12_27_kegg_NB_NP_subset.txt", sep="/"), quote=FALSE, sep
="\t", row.names=FALSE)
```

```
knitr::kable(head(kegg_NB_NP_subset), caption = "First five rows KEEG pathways NB vs NP",format = 'pandoc')
```

First five rows KEEG pathways NB vs NP

| | ind.ensembl | category | over_represented_pvalue | under_represented_pvalue | numDEInCat | numInCat | BY_FDR | external_gene_name |
|---|---|---|---|---|---|---|---|---|
| 1 | ENSBTAG00000000273 | 04060 | 0.0022854 | 0.9997143 | 5 | 100 | 0.0045708 | IL5RA |
| 4 | ENSBTAG00000004211 | 04060 | 0.0022854 | 0.9997143 | 5 | 100 | 0.0045708 | TNFRSF1A |

| | ind.ensembl | category | over_represented_pvalue | under_represented_pvalue | numDEInCat | numInCat | BY_FDR | external_gene_name |
|---|---|---|---|---|---|---|---|---|
| 8 | ENSBTAG00000012899 | 04060 | 0.0022854 | 0.9997143 | 5 | 100 | 0.0045708 | IFNGR2 |
| 11 | ENSBTAG00000020674 | 04060 | 0.0022854 | 0.9997143 | 5 | 100 | 0.0045708 | LTB |
| 14 | ENSBTAG00000049285 | 04060 | 0.0022854 | 0.9997143 | 5 | 100 | 0.0045708 | TNFRSF17 |

### Figure 4a

```
DEG_AI_NB_only<-unique(DEG_AI_NB_mRNA$Row.names[! DEG_AI_NB_mRNA$Row.names %in% c(DEG_AI_NP_mRNA$Row.names, DEG_N
B_NP_mRNA$Row.names)]  )
DEG_AI_NB_AI_NP<-unique(DEG_AI_NB_mRNA$Row.names[DEG_AI_NB_mRNA$Row.names %in% c(DEG_AI_NP_mRNA$Row.names, DEG_NB
_NP_mRNA$Row.names)]  )
DEG_AI_NP_only<-unique(DEG_AI_NP_mRNA$Row.names[! (DEG_AI_NP_mRNA$Row.names %in% c(DEG_AI_NB_mRNA$Row.names, DEG_
NB_NP_mRNA$Row.names))]  )
DEG_AI_NP_NB_NP<-unique(DEG_NB_NP_mRNA$Row.names[ (DEG_NB_NP_mRNA$Row.names %in% DEG_AI_NP_mRNA$Row.names)]  )
DEG_NB_NP_only<-unique(DEG_NB_NP_mRNA$Row.names[! (DEG_NB_NP_mRNA$Row.names %in% c(DEG_AI_NB_mRNA$Row.names, DEG_
AI_NP_mRNA$Row.names))]  )

DEG_AI_NP_only<-DEG_AI_NP_only[flashClust(dist(log2(rpkm_pwbc[c(DEG_AI_NP_only),]+1)), method = "complete")$orde
r]
DEG_AI_NP_NB_NP<-DEG_AI_NP_NB_NP[flashClust(dist(log2(rpkm_pwbc[c(DEG_AI_NP_NB_NP),]+1)), method = "complete")$or
der]
DEG_NB_NP_only<-DEG_NB_NP_only[flashClust(dist(log2(rpkm_pwbc[c(DEG_NB_NP_only),]+1)), method = "complete")$orde
r]

rpkm_AIpreg_heatmap<-log2(rpkm_AIpreg[rownames(rpkm_AIpreg)%in%c(DEG_AI_NB_only,DEG_AI_NB_AI_NP, DEG_AI_NP_only,
 DEG_AI_NP_NB_NP, DEG_NB_NP_only),]+1)
rpkm_AIpreg_heatmap <- rpkm_AIpreg_heatmap[c(DEG_AI_NB_only,DEG_AI_NB_AI_NP, DEG_AI_NP_only, DEG_AI_NP_NB_NP, DEG
_NB_NP_only),]

rpkm_NBpreg_heatmap<-log2(rpkm_NBpreg[rownames(rpkm_NBpreg)%in%c(DEG_AI_NB_only,DEG_AI_NB_AI_NP, DEG_AI_NP_only,
 DEG_AI_NP_NB_NP, DEG_NB_NP_only),]+1)
rpkm_NBpreg_heatmap <- rpkm_NBpreg_heatmap[c(DEG_AI_NB_only,DEG_AI_NB_AI_NP, DEG_AI_NP_only, DEG_AI_NP_NB_NP, DEG
_NB_NP_only),]

rpkm_NOTpreg_heatmap<-log2(rpkm_NOTpreg[rownames(rpkm_NOTpreg)%in%c(DEG_AI_NB_only,DEG_AI_NB_AI_NP, DEG_AI_NP_onl
y, DEG_AI_NP_NB_NP, DEG_NB_NP_only),]+1)
rpkm_NOTpreg_heatmap <- rpkm_NOTpreg_heatmap[c(DEG_AI_NB_only,DEG_AI_NB_AI_NP, DEG_AI_NP_only, DEG_AI_NP_NB_NP, D
EG_NB_NP_only),]

sample_Tree_AI <- flashClust(as.dist((1-cor(rpkm_AIpreg_heatmap))/2), method = "complete")
rpkm_AIpreg_heatmap<-rpkm_AIpreg_heatmap[,sample_Tree_AI$order ]
sample_Tree_NB <- flashClust(as.dist((1-cor(rpkm_NBpreg_heatmap))/2), method = "complete")
rpkm_NBpreg_heatmap<-rpkm_NBpreg_heatmap[,sample_Tree_NB$order ]
sample_Tree_NP<- flashClust(as.dist((1-cor(rpkm_NOTpreg_heatmap))/2), method = "complete")
rpkm_NOTpreg_heatmap<-data.frame(rpkm_NOTpreg_heatmap[,sample_Tree_NP$order ])

rpkm_NOTpreg_heatmap$gene_id<-rownames(rpkm_NOTpreg_heatmap)
rpkm_NOTpreg_heatmap$gene_symbol<-annotation.ensembl.symbol$external_gene_name[match( rpkm_NOTpreg_heatmap$gene_i
d, annotation.ensembl.symbol$ensembl_gene_id )]
rpkm_NOTpreg_heatmap$gene_symbol<-ifelse(rpkm_NOTpreg_heatmap$gene_symbol=="", rpkm_NOTpreg_heatmap$gene_id, rpkm
_NOTpreg_heatmap$gene_symbol)

rownames(rpkm_NOTpreg_heatmap)<-rpkm_NOTpreg_heatmap$gene_symbol
rpkm_NOTpreg_heatmap<-rpkm_NOTpreg_heatmap[,c(1:5)]

annot_matrix<-cbind(rep(c(1,0,0,0,0),c(1,1,40,26,55)) , rep(c(0,1,0,0,0),c(1,1,40,26,55)), rep(c(0,0,1,0,0),c(1,1
,40,26,55)),rep(c(0,0,0,1,0),c(1,1,40,26,55)), rep(c(0,0,0,0,1),c(1,1,40,26,55)))

annotation<-Heatmap(annot_matrix,
        cluster_rows= FALSE,
        cluster_columns = FALSE,
        show_row_names = FALSE,
        show_column_names = FALSE,
        show_heatmap_legend = FALSE,
        col = colorRamp2(c(0,1), c("white","black"), space = "LAB"),
        split= rep(c(1,2,3,4,5), c(length(DEG_AI_NB_only),length(DEG_AI_NB_AI_NP),length(DEG_AI_NP_only),length(D
EG_AI_NP_NB_NP),length(DEG_NB_NP_only))),
        width = unit(5, "mm"))

ht_AIpreg<-Heatmap(rpkm_AIpreg_heatmap,
        column_title = "AI pregnant",
        column_title_gp = gpar(fill = "#0072B2", col = "white", border = "#FFFFFF", fontsize=9),
        cluster_rows= FALSE,
        cluster_columns = FALSE,
        show_row_names = FALSE,
        show_column_names = FALSE,
        show_heatmap_legend = FALSE,
        col = colorRamp2(c(0,15), c("#eef5fb","#3182bd"), space = "LAB"),
        split= rep(c(1,2,3,4,5), c(length(DEG_AI_NB_only),length(DEG_AI_NB_AI_NP),length(DEG_AI_NP_only),length(D
EG_AI_NP_NB_NP),length(DEG_NB_NP_only))),
        width = unit(2, "cm"))

ht_NBpreg<-Heatmap(rpkm_NBpreg_heatmap,
        column_title = "NB pregnant",
        column_title_gp = gpar(fill = "#000000", col = "white", border = "#FFFFFF", fontsize=9),
        cluster_rows= FALSE,
        cluster_columns = FALSE,
        show_row_names = FALSE,
```

```
        show_column_names = FALSE,
        show_heatmap_legend = FALSE,
        col = colorRamp2(c(0,15), c("#eef5fb","#3182bd"), space = "LAB"),
        split= rep(c(1,2,3,4,5), c(length(DEG_AI_NB_only),length(DEG_AI_NB_AI_NP),length(DEG_AI_NP_only),length(D
EG_AI_NP_NB_NP),length(DEG_NB_NP_only))),
        width = unit(2, "cm"))

ht_NOTpreg<-Heatmap(rpkm_NOTpreg_heatmap,
        column_title = "non pregnant",
        column_title_gp = gpar(fill = "#D55E00", col = "white", border = "#FFFFFF", fontsize=9),
        cluster_rows= FALSE,
        cluster_columns = FALSE,
        show_row_names = TRUE,
        row_names_gp = gpar(fontsize=4),
        show_column_names = FALSE,
        show_heatmap_legend = TRUE,
        heatmap_legend_param = list(title = "Log2(FPKM+1)", direction = "horizontal"),
        col = colorRamp2(c(0,15), c("#eef5fb","#3182bd"), space = "LAB"),
        split= rep(c(1,2,3,4,5), c(length(DEG_AI_NB_only),length(DEG_AI_NB_AI_NP),length(DEG_AI_NP_only),length(D
EG_AI_NP_NB_NP),length(DEG_NB_NP_only))),
        width = unit(2, "cm"))

#png(paste(result_path,"2020_01_03_DEG_heatmap.png", sep="/"),width = 1600, height = 2000,res = 300)
#draw(annotation + ht_AIpreg + ht_NBpreg + ht_NOTpreg)
#dev.off()

draw(annotation + ht_AIpreg + ht_NBpreg + ht_NOTpreg)
```

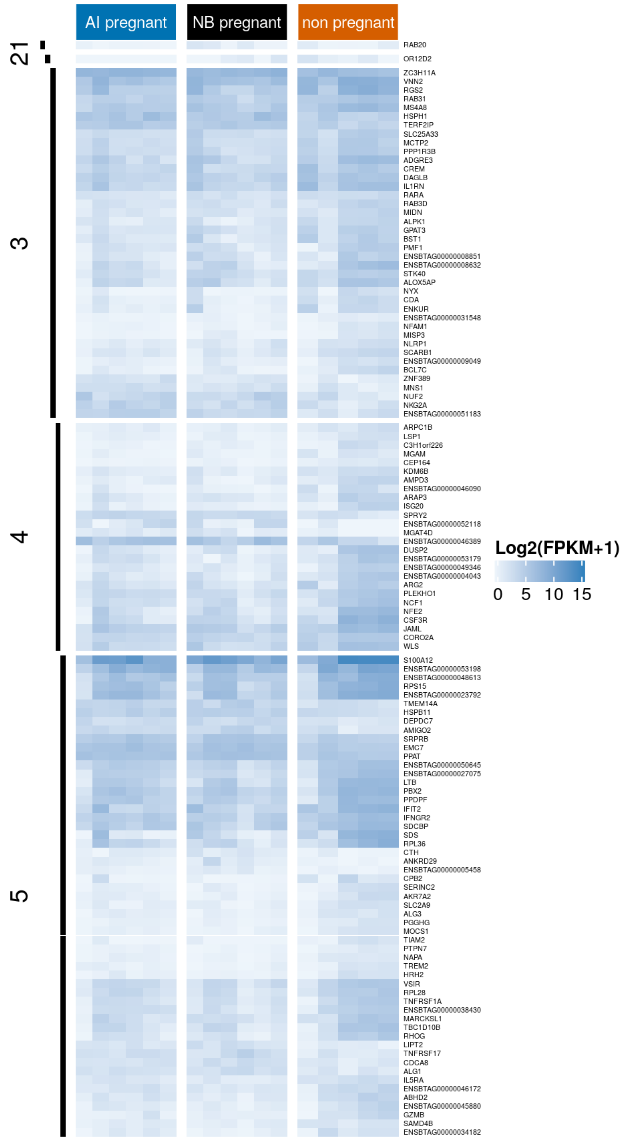

```
merged_correlation_mRNA_miRNA_DEG_AI_NB_only<- merged_correlation_mRNA_miRNA[merged_correlation_mRNA_miRNA$mRNA_g
ene %in% c(DEG_AI_NB_only),]
merged_correlation_mRNA_miRNA_DEG_AI_NB_AI_NP<- merged_correlation_mRNA_miRNA[merged_correlation_mRNA_miRNA$mRNA_
gene %in% c(DEG_AI_NB_AI_NP),]
merged_correlation_mRNA_miRNA_DEG_AI_NP_only<- merged_correlation_mRNA_miRNA[merged_correlation_mRNA_miRNA$mRNA_g
ene %in% c(DEG_AI_NP_only),]
merged_correlation_mRNA_miRNA_DEG_AI_NP_NB_NP<- merged_correlation_mRNA_miRNA[merged_correlation_mRNA_miRNA$mRNA_
gene %in% c( DEG_AI_NP_NB_NP),]
merged_correlation_mRNA_miRNA_DEG_NB_NP_only<- merged_correlation_mRNA_miRNA[merged_correlation_mRNA_miRNA$mRNA_g
ene %in% c( DEG_NB_NP_only),]


#head(DEG_AI_NP_mRNA)
#head(DEG_NB_NP_mRNA)

merged_DEG_AI_NP_mRNA_DEG_NB_NP_mRNA<-merge(DEG_AI_NP_mRNA, DEG_NB_NP_mRNA, by="Row.names", all=TRUE)

#head(merged_DEG_AI_NP_mRNA_DEG_NB_NP_mRNA)

merged_DEG_AI_NP_mRNA_DEG_NB_NP_mRNA_subset<-merged_DEG_AI_NP_mRNA_DEG_NB_NP_mRNA[ (merged_DEG_AI_NP_mRNA_DEG_NB_
NP_mRNA$logFC.x < 0 & merged_DEG_AI_NP_mRNA_DEG_NB_NP_mRNA$logFC.y < 0) |
(merged_DEG_AI_NP_mRNA_DEG_NB_NP_mRNA$logFC.x > 0 & merged_DEG_AI_NP_mRNA_DEG_NB_NP_mRNA$logFC.y > 0),]

merged_DEG_AI_NP_mRNA_DEG_NB_NP_mRNA_subset<-merged_DEG_AI_NP_mRNA_DEG_NB_NP_mRNA_subset[complete.cases(merged_DE
G_AI_NP_mRNA_DEG_NB_NP_mRNA_subset),]


merged_DEG_AI_NP_mRNA_DEG_NB_NP_mRNA_subset_miRNA<-merge(merged_DEG_AI_NP_mRNA_DEG_NB_NP_mRNA_subset,merged_corre
lation_mRNA_miRNA, by.x="Row.names", by.y="mRNA_gene", all=FALSE)

#merged_DEG_AI_NP_mRNA_DEG_NB_NP_mRNA_subset_miRNA
#write.table(merged_DEG_AI_NP_mRNA_DEG_NB_NP_mRNA_subset_miRNA, paste(result_path, "2020_01_04_merged_DEG_AI_NP_m
RNA_DEG_NB_NP_mRNA_subset_miRNA.txt", sep="/"), quote=FALSE, sep="\t", row.names=FALSE)
```

#### Figure 4b

```
cpm_AIpreg<-cpm_pwbc[,c(1,2,3,4,5,6)]
cpm_NOTpreg<-cpm_pwbc[,c(13,14,15,16,17)]
cpm_NBpreg<-cpm_pwbc[,c(7,8,9,10,11,12)]

data_chart_1<-data.frame( stringsAsFactors=FALSE)
data_chart_2<-data.frame( stringsAsFactors=FALSE)

for (i in  seq(dim(merged_DEG_AI_NP_mRNA_DEG_NB_NP_mRNA_subset)[1])){
  gene_id<-merged_DEG_AI_NP_mRNA_DEG_NB_NP_mRNA_subset[i,1]
  gene_symbol<-merged_DEG_AI_NP_mRNA_DEG_NB_NP_mRNA_subset$external_gene_name.x[i]
  if(gene_symbol==""){gene_symbol<-gene_id}

  cpm_AI<-cpm_AIpreg[rownames(cpm_AIpreg)==gene_id,]
  cpm_NB<-cpm_NBpreg[rownames(cpm_NBpreg)==gene_id,]
  cpm_NP<-cpm_NOTpreg[rownames(cpm_NOTpreg)==gene_id,]

  data_chart_1<-rbind(data.frame(cpm=cpm_AI),data.frame(cpm=cpm_NB), data.frame(cpm=cpm_NP))
  data_chart_1$group<-rep(c("Preg_AI","Preg_NB","Not_Preg"),c(6,6,5))
  data_chart_1$gene<-gene_symbol
  data_chart_2<-rbind(data_chart_2,data_chart_1)
}

data_chart_2$group<-factor(data_chart_2$group,levels =c( "Preg_AI" , "Preg_NB" , "Not_Preg"))
plot<-ggplot(data=data_chart_2, aes(x=group , y=cpm, colour=group))+
      geom_jitter(position = position_jitter(width = .1, height=0),  size=1)+
      scale_shape_manual(values=c(0:10))+
      scale_x_discrete(labels=c("Preg_AI"="Preg_AI" , "Preg_NB"="Preg_NB", "Not_Preg"="Not_Preg"))+
      scale_colour_manual(name=NULL, values = c("#0072B2",  "#000000", "#D55E00" ))+
      stat_summary(fun.y = median, fun.ymin = median, fun.ymax = median,colour = "gray", size = 0.2,  geom = "cro
ssbar", alpha=0.5)+
      scale_y_continuous("CPM")+
      facet_wrap(~gene, scales = "free", ncol=4)+
      theme_bw()+
      theme(panel.grid= element_blank(),
            panel.background = element_blank(),
            panel.grid.minor = element_blank(),
            panel.grid.major = element_blank(),
            plot.background = element_blank(),
            strip.background = element_rect(fill = "white"),
            strip.text.x = element_text(colour = 'black', size = 4),
            #legend.text = element_text( colour = 'black', size = 10 ),
            axis.text.x = element_blank(),
            axis.text.y = element_text( colour = 'black', size = 4 ),
            axis.title.y = element_text( colour = 'black', size = 4 ),
            axis.title.x = element_blank(),
            legend.text = element_text( colour = 'black', size = 4 ),
            legend.key.size = unit(0.4,"cm"),
            legend.position=c(0.85, 0.05),
            axis.ticks.x = element_blank(),
            panel.spacing.x=unit(0.5, "mm"),
            panel.spacing.y=unit(0.5, "mm")

      )
#png(file=paste(result_path,"2020_01_05_plot_DEGs_preg_AI_not_preg_NB_not_preg_after_LOOCV.png",sep="/"), width=
 1000, height=2000, bg="transparent",res=300 )
    #plot
#dev.off()
plot
```

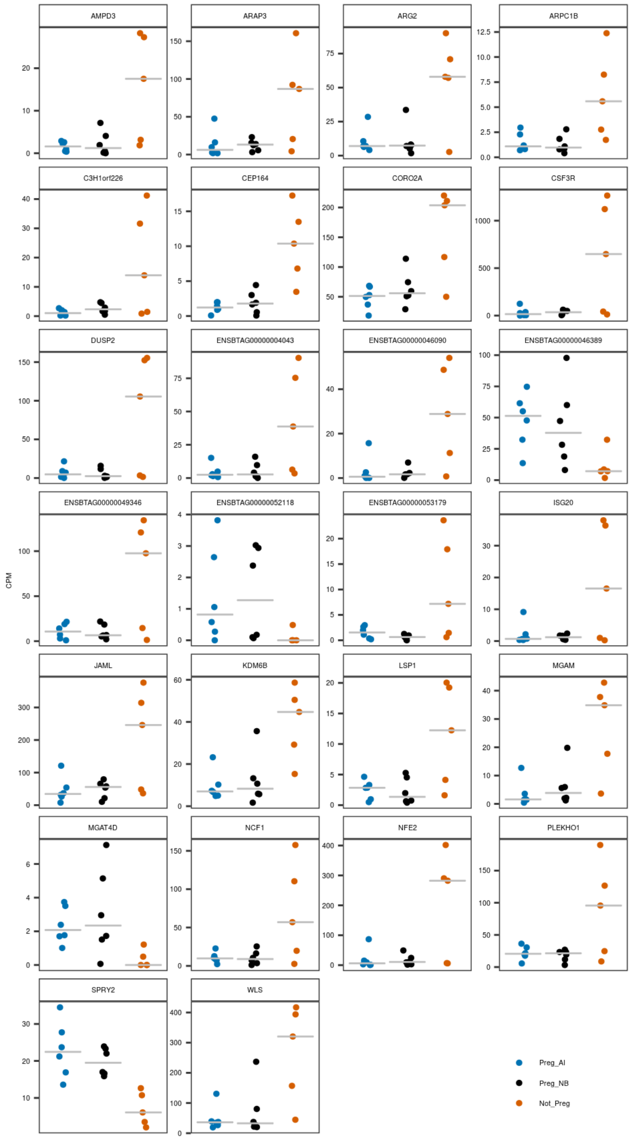
AMPD3
ARAP3
ARG2
ARPC1B
C3H1orf226
CEP164
CORO2A
CSF3R
DUSP2
ENSBTAG00000004043
ENSBTAG00000046090
ENSBTAG00000046389
ENSBTAG00000049346
ENSBTAG00000052118
ENSBTAG00000053179
ISG20
JAML
KDM6B
LSP1
MGAM
MGAT4D
NCF1
NFE2
PLEKHO1
SPRY2
WLS
CPM
Preg_AI
Preg_NB
Not_Preg

### Figure 4c

```
merged_DEG_AI_NP_mRNA_DEG_NB_NP_mRNA_subset_miRNA_a<-merged_DEG_AI_NP_mRNA_DEG_NB_NP_mRNA_subset_miRNA[!(merged_D
EG_AI_NP_mRNA_DEG_NB_NP_mRNA_subset_miRNA$description.x==""),]
merged_DEG_AI_NP_mRNA_DEG_NB_NP_mRNA_subset_miRNA_a[10,41]<-"FCGR2"
merged_DEG_AI_NP_mRNA_DEG_NB_NP_mRNA_subset_miRNA_a[20,41]<-"CEACAM4"

data_for_chart<-data.frame()
data_for_chart_a<-data.frame()
for(i in 1:dim(merged_DEG_AI_NP_mRNA_DEG_NB_NP_mRNA_subset_miRNA_a)[1] ){
  expression_gene_1_AI<-cpm_AIpreg[row.names(cpm_AIpreg)==as.character(merged_DEG_AI_NP_mRNA_DEG_NB_NP_mRNA_subse
t_miRNA_a$Row.names)[i],]
  expression_gene_2_AI<-cpm_AIpreg_miRNA[row.names(cpm_AIpreg_miRNA)==as.character(merged_DEG_AI_NP_mRNA_DEG_NB_N
P_mRNA_subset_miRNA_a$miRNA_gene)[i],]
  expression_gene_1_NB<-cpm_NBpreg[row.names(cpm_NBpreg)==as.character(merged_DEG_AI_NP_mRNA_DEG_NB_NP_mRNA_subse
t_miRNA_a$Row.names)[i],]
  expression_gene_2_NB<-cpm_NBpreg_miRNA[row.names(cpm_NBpreg_miRNA)==as.character(merged_DEG_AI_NP_mRNA_DEG_NB_N
P_mRNA_subset_miRNA_a$miRNA_gene)[i],]
  expression_gene_1_NP<-cpm_NOTpreg[row.names(cpm_NOTpreg)==as.character(merged_DEG_AI_NP_mRNA_DEG_NB_NP_mRNA_sub
set_miRNA_a$Row.names)[i],]
  expression_gene_2_NP<-cpm_NOTpreg_miRNA[row.names(cpm_NOTpreg_miRNA)==as.character(merged_DEG_AI_NP_mRNA_DEG_NB
_NP_mRNA_subset_miRNA_a$miRNA_gene)[i],]

  symbol_gene_1<-as.character(merged_DEG_AI_NP_mRNA_DEG_NB_NP_mRNA_subset_miRNA_a$mRNA_symbol)[i]
  symbol_gene_2<-as.character(merged_DEG_AI_NP_mRNA_DEG_NB_NP_mRNA_subset_miRNA_a$miRNA_symbol)[i]

  chart<-i

  data_for_chart<-data.frame(gene_1=c(expression_gene_1_AI,expression_gene_1_NB,expression_gene_1_NP) ,
                             gene_2=c(expression_gene_2_AI,expression_gene_2_NB,expression_gene_2_NP),
                             symbol_gene_1,symbol_gene_2 ,chart,
                             group=rep(c("AI", "NB", "NP"), c(length(expression_gene_1_AI),length(expression_gene
_1_NB),length(expression_gene_1_NP))))

  data_for_chart_a<-rbind(data_for_chart_a,data_for_chart)
}

font_axis<-5
font_title<-5
ptsize<-0.05

plots<-list()
k<-1
for (j in c(1:20)){
  data_for_chart_b<-data_for_chart_a[data_for_chart_a$chart %in% j,]

plot<-ggplot(data_for_chart_b, aes(y=gene_1,x=gene_2))+
  geom_point(size=0.5)+
  geom_smooth(method=lm, size=ptsize, fill="lightgray")+
  scale_y_continuous(name=data_for_chart_b$symbol_gene_1[1])+
  scale_x_continuous(name=data_for_chart_b$symbol_gene_2[1])+
  facet_wrap(~group, ncol=3, scales="free")+
  theme(aspect.ratio = 1,
            panel.grid.major = element_blank(),
            panel.grid.minor = element_blank(),
            panel.background = element_blank(),
            plot.background  = element_blank(),
            axis.text.x = element_text( colour = 'black',size = font_axis, angle=90),
            axis.text.y = element_text( colour = 'black',size = font_axis),
            axis.title = element_text( colour = 'black' ,size = font_title, face="italic"),
            axis.ticks = element_line(size=0.1),
            panel.spacing.y = unit(0, "mm"),
            panel.spacing.x = unit(1.5, "mm"),
            legend.position="none",
            axis.line=element_line(size = 0.1, colour = "black"),
            strip.background=element_blank(),
            strip.text.x = element_blank())

 plots[[k]]<-plot
  k<-k+1
}

#png(filename =paste(result_path,"2020_02_19_plot_DE_diff_coexpression.png", sep="/"), width = 9, height =3, unit
s = 'in', res = 300,bg = "transparent")
#ggarrange(plotlist=plots, ncol=5, nrow=4)
#dev.off()
ggarrange(plotlist=plots, ncol=5, nrow=4)
```

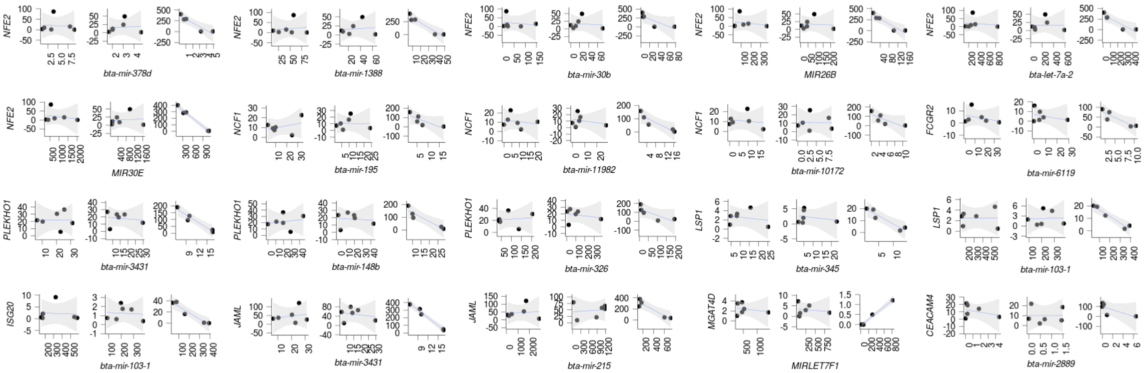

### DEG Analysis for miRNA

#### Pregnant to Artificial Insemination vs non pregnant

##### Results from edgeR

```
colnames(count_miRNA_plasma)
```

```
##  [1] "AI_92"  "AI_99"  "AI_114" "AI_117" "AI_119" "AI_120" "NB_93"  "NB_103"
##  [9] "NB_108" "NB_111" "NB_116" "NB_121" "NP_94"  "NP_101" "NP_107" "NP_113"
## [17] "NP_115"
```

```
group<-factor(c("Preg_AI","Preg_AI","Preg_AI","Preg_AI","Preg_AI","Preg_AI","Preg_NB","Preg_NB","Preg_NB","Preg_N
B","Preg_NB","Preg_NB","Not_Preg","Not_Preg","Not_Preg","Not_Preg","Not_Preg"), levels=c("Preg_AI","Preg_NB", "No
t_Preg"))
design<-model.matrix(~group)
dds<-DGEList(count=count_miRNA_plasma, group=group)
dds<-estimateDisp(dds, design, robust=TRUE)
dds<-glmFit(dds, design)
dds<-glmLRT(dds,coef="groupNot_Preg")
res_not_preg_preg_AI_miRNA_edgeR<-topTags(dds,n=Inf)$table
#head(res_not_preg_preg_AI_miRNA_edgeR,n=30)
#write.table(res_not_preg_preg_AI_miRNA_edgeR, paste(result_path, "2019_04_30_res_not_preg_preg_AI_miRNA_edgeR.tx
t"), quote=FALSE, sep="\t", row.names=FALSE)
```

##### Results from DESeq

```
design<-model.matrix(~group)
colData<-data.frame("group"=group)
rownames(colData)<-colnames(count_miRNA_plasma)
dds<-DESeqDataSetFromMatrix(countData=count_miRNA_plasma, colData=colData, design= ~group)
dds<-DESeq(dds)
res_not_preg_preg_AI_miRNA_DeSeq<-results(dds, contrast=c("group","Not_Preg" ,"Preg_AI"), pAdjustMethod="fdr", ti
dy=TRUE)
res_not_preg_preg_AI_miRNA_DeSeq<-res_not_preg_preg_AI_miRNA_DeSeq[with(res_not_preg_preg_AI_miRNA_DeSeq, order(p
value)), ]
#head(res_not_preg_preg_AI_miRNA_DeSeq,n=30)
#write.table(res_not_preg_preg_AI_miRNA_DeSeq, paste(result_path, "2019_04_30_res_not_preg_preg_AI_miRNA_DeSeq.tx
t"), quote=FALSE, sep="\t", row.names=FALSE)
```

Merge and annotate the results from edgeR and DESeq. (AI Pregnant vs non pregnant)

```
merged_res_not_preg_preg_AI_miRNA<-merge(res_not_preg_preg_AI_miRNA_edgeR, res_not_preg_preg_AI_miRNA_DeSeq, by.x
="row.names", by.y="row")
merged_res_not_preg_preg_AI_miRNA<-merged_res_not_preg_preg_AI_miRNA[merged_res_not_preg_preg_AI_miRNA$PValue<=0.
03 & merged_res_not_preg_preg_AI_miRNA$pvalue<=0.03,]
dim(merged_res_not_preg_preg_AI_miRNA) #8
```

```
## [1]  8 12
```

```
merged_res_not_preg_preg_AI_miRNA<-merged_res_not_preg_preg_AI_miRNA[with(merged_res_not_preg_preg_AI_miRNA, orde
r(logFC)), ]
#head(merged_res_not_preg_preg_AI_miRNA)
#write.table(res_not_preg_preg_AI_miRNA_DeSeq, paste(result_path, "2019_04_30_res_not_preg_preg_AI_miRNA_DeSeq.tx
t"), quote=FALSE, sep="\t", row.names=FALSE)

annotated_merged_res_not_preg_preg_AI_miRNA<-merge(merged_res_not_preg_preg_AI_miRNA,annotation.ensembl.symbol, b
y.x="Row.names", by.y="ensembl_gene_id", all=FALSE)
#head(annotated_merged_res_not_preg_preg_AI_miRNA)
annotated_merged_res_not_preg_preg_AI_miRNA<-annotated_merged_res_not_preg_preg_AI_miRNA[with(annotated_merged_re
s_not_preg_preg_AI_miRNA, order(logFC)), ]
dim(annotated_merged_res_not_preg_preg_AI_miRNA) #7
```

```
## [1]  7 17
```

```
#write.table(annotated_merged_res_not_preg_preg_AI_miRNA, paste(result_path, "2019_04_30_annotated_merged_res_not
_preg_preg_AI_miRNA.txt"), quote=FALSE, sep="\t", row.names=FALSE)
```

Plot the charts for all genes inferred as differentially expressed between AI and non pregnant

```
cpm_AIpreg<-cpm_miRNA[,c(1,2,3,4,5,6)]
cpm_NOTpreg<-cpm_miRNA[,c(13,14,15,16,17)]
cpm_NBpreg<-cpm_miRNA[,c(7,8,9,10,11,12)]

data_chart_1<-data.frame( stringsAsFactors=FALSE)
data_chart_2<-data.frame( stringsAsFactors=FALSE)

for (i in  seq(dim(annotated_merged_res_not_preg_preg_AI_miRNA)[1])){

  gene_id<-annotated_merged_res_not_preg_preg_AI_miRNA[i,1]
  gene_symbol<-annotated_merged_res_not_preg_preg_AI_miRNA$external_gene_name[i]
  if(gene_symbol==""){gene_symbol<-gene_id}

  cpm_AI<-cpm_AIpreg[rownames(cpm_AIpreg)==gene_id,]
  cpm_NP<-cpm_NOTpreg[rownames(cpm_NOTpreg)==gene_id,]

  data_chart_1<-rbind(data.frame(cpm=cpm_AI), data.frame(cpm=cpm_NP))
  data_chart_1$group<-rep(c("Preg_AI","Not_Preg"),c(6,5))
  data_chart_1$shape<-as.factor(c(0:10))
  data_chart_1$gene<-gene_symbol
  data_chart_1$chart<-i
  data_chart_1$fold_change<-annotated_merged_res_not_preg_preg_AI_miRNA[annotated_merged_res_not_preg_preg_AI_miR
NA$Row.names==gene_id,]$logFC
  data_chart_1$pvalue<-annotated_merged_res_not_preg_preg_AI_miRNA[annotated_merged_res_not_preg_preg_AI_miRNA$Ro
w.names==gene_id,]$PValue


  data_chart_2<-rbind(data_chart_2,data_chart_1)
}

pdf(file=paste(result_path,"04_30_plot_all_DEGs_preg_AI_not_preg_miRNA_before_LOOCV.pdf",sep="/"), width= 11, pap
er="USr", bg="transparent", onefile=TRUE)


for (i in seq(1,dim(annotated_merged_res_not_preg_preg_AI_miRNA)[1], 24)) {
  plots <- list()
  k<-1
  for (j in c(i:(i+23))){
    data_chart_3<-data_chart_2[data_chart_2$chart %in% j , ]
    plot<-ggplot(data=data_chart_3, aes(x=group , y=cpm, colour=group))+
      geom_jitter(position = position_jitter(width = .1, height=0), aes( shape=shape), size=2)+
      scale_shape_manual(values=c(0:11))+
      scale_x_discrete(data_chart_3$gene[1],labels=c("Preg_AI"="Preg_AI", "Not_Preg"="Not_Preg"))+
      scale_colour_manual(name=NULL, values = c("red", "blue"))+
      stat_summary(fun.y = median, fun.ymin = median, fun.ymax = median,colour = "gray", size = 0.2,  geom = "cro
ssbar", alpha=0.5)+
      scale_y_continuous("CPM")+
      ggtitle(paste("LogFC","=", round(data_chart_3$fold_change[1],1),"  ","P","=",round(data_chart_3$pvalue[1],2
), sep=c("")))+
      theme_bw()+
      theme(panel.grid= element_blank(),
            panel.background = element_blank(),
            panel.grid.minor = element_blank(),
            panel.grid.major = element_blank(),
            plot.background = element_blank(),
            strip.background = element_rect(fill = "white"),
            #strip.text.x = element_text(colour = 'black', face="italic",size = 9),
            #legend.text = element_text( colour = 'black', size = 10 ),
            axis.text.x = element_blank(),
            axis.text.y = element_text( colour = 'black', size = 8 ),
            axis.title = element_text( colour = 'black', size = 9 ),
            legend.key.size = unit(0.9,"cm"),
            legend.position="none",
            axis.ticks.x = element_blank(),
            plot.title = element_text(size = 7)
      )

    plots[[k]] <- plot
    k<-k+1
  }
  multiplot(plotlist = plots, cols = 6,layout = matrix(1:24, nrow=4, byrow = TRUE))
}
dev.off()
```

Leave-one-out cross validation

```
group<-factor(c("Preg_AI","Preg_AI","Preg_AI","Preg_AI","Preg_AI","Preg_AI","Preg_NB","Preg_NB","Preg_NB","Preg_N
B","Preg_NB","Preg_NB","Not_Preg","Not_Preg","Not_Preg","Not_Preg","Not_Preg"), levels=c("Preg_AI","Preg_NB", "No
t_Preg"))

out_list_IDs_LOOCV<-foreach(i = c(1:6,13:17), .inorder=FALSE,.packages=c("edgeR","DESeq2"), .verbose=FALSE ) %do%
{

group_a<-group[-i]
count_miRNA_plasma_a<-count_miRNA_plasma[,-i]

design<-model.matrix(~group_a)
dds<-DGEList(count=count_miRNA_plasma_a, group=group_a)
dds<-estimateDisp(dds, design, robust=TRUE)
dds<-glmFit(dds, design)
dds<-glmLRT(dds,coef="group_aNot_Preg")
res_not_preg_preg_AI_edgeR<-topTags(dds,n=Inf)$table

#DESeq
colData<-data.frame("group"=group_a)
rownames(colData)<-colnames(count_miRNA_plasma_a)
dds<-DESeqDataSetFromMatrix(countData=count_miRNA_plasma_a, colData=colData, design= ~group)
dds<-DESeq(dds)
res_not_preg_preg_AI_DeSeq<-results(dds, contrast=c("group","Not_Preg" ,"Preg_AI"), pAdjustMethod="fdr", tidy=TRU
E)


merged_res_not_preg_preg_AI_miRNA_a<-merge(res_not_preg_preg_AI_edgeR, res_not_preg_preg_AI_DeSeq, by.x="row.name
s", by.y="row")
merged_res_not_preg_preg_AI_miRNA_a<-merged_res_not_preg_preg_AI_miRNA_a[merged_res_not_preg_preg_AI_miRNA_a$PVal
ue<=0.03 & merged_res_not_preg_preg_AI_miRNA_a$pvalue<=0.03,]

rm(group_a,count_miRNA_plasma_a)

merged_res_not_preg_preg_AI_miRNA_a$Row.names
}

overlapping_genes_LOOCV<-Reduce(intersect, out_list_IDs_LOOCV)

merged_res_not_preg_preg_AI_miRNA<-merge(res_not_preg_preg_AI_miRNA_edgeR, res_not_preg_preg_AI_miRNA_DeSeq, by.x
="row.names", by.y="row")
merged_res_not_preg_preg_AI_miRNA_a<-merged_res_not_preg_preg_AI_miRNA[merged_res_not_preg_preg_AI_miRNA$Row.name
s %in% overlapping_genes_LOOCV,]
annotated_merged_res_not_preg_preg_AI_miRNA<-merge(merged_res_not_preg_preg_AI_miRNA_a,annotation.ensembl.symbol,
by.x="Row.names", by.y="ensembl_gene_id", all=FALSE)
annotated_merged_res_not_preg_preg_AI_miRNA<-annotated_merged_res_not_preg_preg_AI_miRNA[with(annotated_merged_re
s_not_preg_preg_AI_miRNA, order(logFC)), ]
dim(annotated_merged_res_not_preg_preg_AI_miRNA)#1
```

```
## [1]  1 17
```

```
#head(annotated_merged_res_not_preg_preg_AI_miRNA)
#write.table(annotated_merged_res_not_preg_preg_AI_miRNA, paste(result_path, "2020_01_04_annotated_merged_res_not
_preg_preg_AI_miRNA_after_LOOCV.txt"), quote=FALSE, sep="\t", row.names=FALSE)
rm(out_list_IDs_LOOCV)
```

```
knitr::kable(annotated_merged_res_not_preg_preg_AI_miRNA,format = 'pandoc')
```

| Row.names | logFC | logCPM | LR | PValue | FDR | baseMean | log2FoldChange | lfcSE | stat | pvalue | padj |
|---|---|---|---|---|---|---|---|---|---|---|---|
| ENSBTAG00000052419 | -5.096042 | 0.5841013 | 12.22747 | 0.0004709 | 0.068282 | 7.394997 | -5.113215 | 1.951268 | -2.620458 | 0.0087812 | 0.362537 |

Plot the charts for all genes inferred as differentially expressed between AI and non pregnant

```
cpm_AIpreg<-cpm_miRNA[,c(1,2,3,4,5,6)]
cpm_NOTpreg<-cpm_miRNA[,c(13,14,15,16,17)]
cpm_NBpreg<-cpm_miRNA[,c(7,8,9,10,11,12)]

data_chart_1<-data.frame( stringsAsFactors=FALSE)
data_chart_2<-data.frame( stringsAsFactors=FALSE)

for (i in  seq(dim(annotated_merged_res_not_preg_preg_AI_miRNA)[1])){

  gene_id<-annotated_merged_res_not_preg_preg_AI_miRNA[i,1]
  gene_symbol<-annotated_merged_res_not_preg_preg_AI_miRNA$external_gene_name[i]
  if(gene_symbol==""){gene_symbol<-gene_id}

  cpm_AI<-cpm_AIpreg[rownames(cpm_AIpreg)==gene_id,]
  cpm_NP<-cpm_NOTpreg[rownames(cpm_NOTpreg)==gene_id,]

  data_chart_1<-rbind(data.frame(cpm=cpm_AI), data.frame(cpm=cpm_NP))
  data_chart_1$group<-rep(c("Preg_AI","Not_Preg"),c(6,5))
  data_chart_1$shape<-as.factor(c(0:10))
  data_chart_1$gene<-gene_symbol
  data_chart_1$chart<-i
  data_chart_1$fold_change<-annotated_merged_res_not_preg_preg_AI_miRNA[annotated_merged_res_not_preg_preg_AI_miR
NA$Row.names==gene_id,]$logFC
  data_chart_1$pvalue<-annotated_merged_res_not_preg_preg_AI_miRNA[annotated_merged_res_not_preg_preg_AI_miRNA$Ro
w.names==gene_id,]$PValue


  data_chart_2<-rbind(data_chart_2,data_chart_1)
}

pdf(file=paste(result_path,"04_30_plot_all_DEGs_preg_AI_not_preg_miRNA_after_LOOCV.pdf",sep="/"), width= 11, pape
r="USr", bg="transparent", onefile=TRUE)


for (i in seq(1,dim(annotated_merged_res_not_preg_preg_AI_miRNA)[1], 24)) {
  plots <- list()
  k<-1
  for (j in c(i:(i+23))){
    data_chart_3<-data_chart_2[data_chart_2$chart %in% j , ]
    plot<-ggplot(data=data_chart_3, aes(x=group , y=cpm, colour=group))+
      geom_jitter(position = position_jitter(width = .1, height=0), aes( shape=shape), size=2)+
      scale_shape_manual(values=c(0:11))+
      scale_x_discrete(data_chart_3$gene[1],labels=c("Preg_AI"="Preg_AI", "Not_Preg"="Not_Preg"))+
      scale_colour_manual(name=NULL, values = c("red", "blue"))+
      stat_summary(fun.y = median, fun.ymin = median, fun.ymax = median,colour = "gray", size = 0.2,  geom = "cro
ssbar", alpha=0.5)+
      scale_y_continuous("CPM")+
      ggtitle(paste("LogFC","=", round(data_chart_3$fold_change[1],1),"  ","P","=",round(data_chart_3$pvalue[1],2
), sep=c("")))+
      theme_bw()+
      theme(panel.grid= element_blank(),
            panel.background = element_blank(),
            panel.grid.minor = element_blank(),
            panel.grid.major = element_blank(),
            plot.background = element_blank(),
            strip.background = element_rect(fill = "white"),
            #strip.text.x = element_text(colour = 'black', face="italic",size = 9),
            #legend.text = element_text( colour = 'black', size = 10 ),
            axis.text.x = element_blank(),
            axis.text.y = element_text( colour = 'black', size = 8 ),
            axis.title = element_text( colour = 'black', size = 9 ),
            legend.key.size = unit(0.9,"cm"),
            legend.position="none",
            axis.ticks.x = element_blank(),
            plot.title = element_text(size = 7)
      )

    plots[[k]] <- plot
    k<-k+1
  }
  multiplot(plotlist = plots, cols = 6,layout = matrix(1:24, nrow=4, byrow = TRUE))
}
dev.off()
```

### AI Pregnant vs Pregnant to Natural Service

#### Results from edgeR

```
#colnames(count_miRNA_plasma)
group<-factor(c("Preg_AI","Preg_AI","Preg_AI","Preg_AI","Preg_AI","Preg_AI","Preg_NB","Preg_NB","Preg_NB","Preg_N
B","Preg_NB","Preg_NB","Not_Preg","Not_Preg","Not_Preg","Not_Preg","Not_Preg"), levels=c("Preg_AI","Preg_NB", "No
t_Preg"))
design<-model.matrix(~group)
dds<-DGEList(count=count_miRNA_plasma, group=group)
dds<-estimateDisp(dds, design, robust=TRUE)
dds<-glmFit(dds, design)
dds<-glmLRT(dds,coef="groupPreg_NB")
res_preg_AI_preg_NB_miRNA_edgeR<-topTags(dds,n=Inf)$table
#head(res_preg_AI_preg_NB_miRNA_edgeR,n=30)
#write.table(res_preg_AI_preg_NB_miRNA_edgeR, paste(result_path, "2019_04_30_res_preg_AI_preg_NB_miRNA_edgeR.tx
t"), quote=FALSE, sep="\t", row.names=FALSE)
```

#### Results from DESeq

```
design<-model.matrix(~group)
colData<-data.frame("group"=group)
rownames(colData)<-colnames(count_miRNA_plasma)
dds<-DESeqDataSetFromMatrix(countData=count_miRNA_plasma, colData=colData, design= ~group)
dds<-DESeq(dds)
res_preg_AI_preg_NB_miRNA_DeSeq<-results(dds, contrast=c("group","Preg_NB" ,"Preg_AI"), pAdjustMethod="fdr", tidy
=TRUE)
res_preg_AI_preg_NB_miRNA_DeSeq<-res_preg_AI_preg_NB_miRNA_DeSeq[with(res_preg_AI_preg_NB_miRNA_DeSeq, order(pval
ue)), ]
#head(res_preg_AI_preg_NB_miRNA_DeSeq,n=30)
#write.table(res_preg_AI_preg_NB_miRNA_DeSeq, paste(result_path, "2019_04_30_res_preg_AI_preg_NB_miRNA_DeSeq.tx
t"), quote=FALSE, sep="\t", row.names=FALSE)
```

Merge the results from edgeR and DESeq. (AI Pregnant vs Pregnant to Natural Service)

```
merged_res_preg_AI_preg_NB_miRNA<-merge(res_preg_AI_preg_NB_miRNA_edgeR, res_preg_AI_preg_NB_miRNA_DeSeq, by.x="r
ow.names", by.y="row")
merged_res_preg_AI_preg_NB_miRNA<-merged_res_preg_AI_preg_NB_miRNA[merged_res_preg_AI_preg_NB_miRNA$PValue<=0.03
 & merged_res_preg_AI_preg_NB_miRNA$pvalue<=0.03,]
dim(merged_res_preg_AI_preg_NB_miRNA) #0
```

```
## [1]  1 12
```

```
#head(merged_res_preg_AI_preg_NB_miRNA)
```

### Pregnant to Natural Service vs non pregnant

#### Results from edgeR

```
#colnames(count_miRNA_plasma)
group<-factor(c("Preg_AI","Preg_AI","Preg_AI","Preg_AI","Preg_AI","Preg_AI","Preg_NB","Preg_NB","Preg_NB","Preg_N
B","Preg_NB","Preg_NB","Not_Preg","Not_Preg","Not_Preg","Not_Preg","Not_Preg"), levels=c("Preg_NB","Preg_AI","Not
_Preg"))
design<-model.matrix(~group)
dds<-DGEList(count=count_miRNA_plasma, group=group)
dds<-estimateDisp(dds, design, robust=TRUE)
dds<-glmFit(dds, design)
dds<-glmLRT(dds,coef="groupNot_Preg")
res_not_preg_preg_NB_miRNA_edgeR<-topTags(dds,n=Inf)$table
#head(res_not_preg_preg_NB_miRNA_edgeR,n=30)
#write.table(res_not_preg_preg_NB_miRNA_edgeR, paste(result_path, "2019_04_30_res_not_preg_preg_NB_miRNA_edgeR.tx
t"), quote=FALSE, sep="\t", row.names=FALSE)
```

#### Results from DESeq

```
design<-model.matrix(~group)
colData<-data.frame("group"=group)
rownames(colData)<-colnames(count_miRNA_plasma)
dds<-DESeqDataSetFromMatrix(countData=count_miRNA_plasma, colData=colData, design= ~group)
dds<-DESeq(dds)
res_not_preg_preg_NB_miRNA_DeSeq<-results(dds, contrast=c("group","Not_Preg" ,"Preg_NB"), pAdjustMethod="fdr", ti
dy=TRUE)
res_not_preg_preg_NB_miRNA_DeSeq<-res_not_preg_preg_NB_miRNA_DeSeq[with(res_not_preg_preg_NB_miRNA_DeSeq, order(p
value)), ]
#head(res_not_preg_preg_NB_miRNA_DeSeq,n=30)
#write.table(res_not_preg_preg_NB_miRNA_DeSeq, paste(result_path, "2019_04_30_res_not_preg_preg_NB_miRNA_DeSeq.tx
t"), quote=FALSE, sep="\t", row.names=FALSE)
```

Merge the results from edgeR and DESeq. (Pregnant to Natural Service vs non pregnant)

```
merged_res_not_preg_preg_NB_miRNA<-merge(res_not_preg_preg_NB_miRNA_edgeR, res_not_preg_preg_NB_miRNA_DeSeq, by.x
="row.names", by.y="row")
merged_res_not_preg_preg_NB_miRNA<-merged_res_not_preg_preg_NB_miRNA[merged_res_not_preg_preg_NB_miRNA$PValue<=0.
03 & merged_res_not_preg_preg_NB_miRNA$pvalue<=0.03,]
dim(merged_res_not_preg_preg_NB_miRNA) #1
```

```
## [1]  2 12
```

```
#head(merged_res_not_preg_preg_NB_miRNA)

annotated_merged_res_not_preg_preg_NB_miRNA<-merge(merged_res_not_preg_preg_NB_miRNA,annotation.ensembl.symbol, b
y.x="Row.names", by.y="ensembl_gene_id", all=FALSE)
#head(annotated_merged_res_not_preg_preg_NB_miRNA)
#write.table(annotated_merged_res_not_preg_preg_NB_miRNA, paste(result_path, "2019_04_30_annotated_merged_res_not
_preg_preg_NB_miRNA.txt"), quote=FALSE, sep="\t", row.names=FALSE)
```

Plot the charts for all genes inferred as differentially expressed between Pregnant to Natural Service and non pregnant

```
cpm_AIpreg<-cpm_miRNA[,c(1,2,3,4,5,6)]
cpm_NOTpreg<-cpm_miRNA[,c(13,14,15,16,17)]
cpm_NBpreg<-cpm_miRNA[,c(7,8,9,10,11,12)]

data_chart_1<-data.frame( stringsAsFactors=FALSE)
data_chart_2<-data.frame( stringsAsFactors=FALSE)

for (i in  seq(dim(annotated_merged_res_not_preg_preg_NB_miRNA)[1])){

  gene_id<-annotated_merged_res_not_preg_preg_NB_miRNA[i,1]
  gene_symbol<-annotated_merged_res_not_preg_preg_NB_miRNA$external_gene_name[i]
  if(gene_symbol==""){gene_symbol<-gene_id}

  cpm_NB<-cpm_NBpreg[rownames(cpm_NBpreg)==gene_id,]
  cpm_NP<-cpm_NOTpreg[rownames(cpm_NOTpreg)==gene_id,]

  data_chart_1<-rbind(data.frame(cpm=cpm_NB), data.frame(cpm=cpm_NP))
  data_chart_1$group<-rep(c("Preg_NB","Not_Preg"),c(6,5))
  data_chart_1$shape<-as.factor(c(0:10))
  data_chart_1$gene<-gene_symbol
  data_chart_1$chart<-i
  data_chart_1$fold_change<-annotated_merged_res_not_preg_preg_NB_miRNA[annotated_merged_res_not_preg_preg_NB_miR
NA$Row.names==gene_id,]$logFC
  data_chart_1$pvalue<-annotated_merged_res_not_preg_preg_NB_miRNA[annotated_merged_res_not_preg_preg_NB_miRNA$Ro
w.names==gene_id,]$PValue


  data_chart_2<-rbind(data_chart_2,data_chart_1)
}

pdf(file=paste(result_path,"04_30_plot_all_DEGs_preg_NB_not_preg_miRNA_before_LOOCV.pdf",sep="/"), width= 11, pap
er="USr", bg="transparent", onefile=TRUE)


for (i in seq(1,dim(annotated_merged_res_not_preg_preg_NB_miRNA)[1], 24)) {
  plots <- list()
  k<-1
  for (j in c(i:(i+23))){
    data_chart_3<-data_chart_2[data_chart_2$chart %in% j , ]
    plot<-ggplot(data=data_chart_3, aes(x=group , y=cpm, colour=group))+
      geom_jitter(position = position_jitter(width = .1, height=0), aes( shape=shape), size=2)+
      scale_shape_manual(values=c(0:11))+
      scale_x_discrete(data_chart_3$gene[1],labels=c("Preg_NB"="Preg_NB", "Not_Preg"="Not_Preg"))+
      scale_colour_manual(name=NULL, values = c("red", "black"))+
      stat_summary(fun.y = median, fun.ymin = median, fun.ymax = median,colour = "gray", size = 0.2,  geom = "cro
ssbar", alpha=0.5)+
      scale_y_continuous("CPM")+
      ggtitle(paste("LogFC","=", round(data_chart_3$fold_change[1],1),"  ","P","=",round(data_chart_3$pvalue[1],2
), sep=c("")))+
      theme_bw()+
      theme(panel.grid= element_blank(),
            panel.background = element_blank(),
            panel.grid.minor = element_blank(),
            panel.grid.major = element_blank(),
            plot.background = element_blank(),
            strip.background = element_rect(fill = "white"),
            #strip.text.x = element_text(colour = 'black', face="italic",size = 9),
            #legend.text = element_text( colour = 'black', size = 10 ),
            axis.text.x = element_blank(),
            axis.text.y = element_text( colour = 'black', size = 8 ),
            axis.title = element_text( colour = 'black', size = 9 ),
            legend.key.size = unit(0.9,"cm"),
            legend.position="none",
            axis.ticks.x = element_blank(),
            plot.title = element_text(size = 7)
      )

    plots[[k]] <- plot
    k<-k+1
  }
  multiplot(plotlist = plots, cols = 6,layout = matrix(1:24, nrow=4, byrow = TRUE))
}
dev.off()
```

Leave-one-out cross validation

```
group<-factor(c("Preg_AI","Preg_AI","Preg_AI","Preg_AI","Preg_AI","Preg_AI","Preg_NB","Preg_NB","Preg_NB","Preg_N
B","Preg_NB","Preg_NB","Not_Preg","Not_Preg","Not_Preg","Not_Preg","Not_Preg"), levels=c("Preg_NB", "Preg_AI","No
t_Preg"))

out_list_IDs_LOOCV<-foreach(i = c(7:17), .inorder=FALSE,.packages=c("edgeR","DESeq2"), .verbose=FALSE ) %do% {

group_a<-group[-i]
count_miRNA_plasma_a<-count_miRNA_plasma[,-i]

design<-model.matrix(~group_a)
dds<-DGEList(count=count_miRNA_plasma_a, group=group_a)
dds<-estimateDisp(dds, design, robust=TRUE)
dds<-glmFit(dds, design)
dds<-glmLRT(dds,coef="group_aNot_Preg")
res_not_preg_preg_NB_edgeR<-topTags(dds,n=Inf)$table

#DESeq
colData<-data.frame("group"=group_a)
rownames(colData)<-colnames(count_miRNA_plasma_a)
dds<-DESeqDataSetFromMatrix(countData=count_miRNA_plasma_a, colData=colData, design= ~group)
dds<-DESeq(dds)
res_not_preg_preg_NB_DeSeq<-results(dds, contrast=c("group","Not_Preg" ,"Preg_NB"), pAdjustMethod="fdr", tidy=TRU
E)


merged_res_not_preg_preg_NB_miRNA_a<-merge(res_not_preg_preg_NB_edgeR, res_not_preg_preg_NB_DeSeq, by.x="row.name
s", by.y="row")
merged_res_not_preg_preg_NB_miRNA_a<-merged_res_not_preg_preg_NB_miRNA_a[merged_res_not_preg_preg_NB_miRNA_a$PVal
ue<=0.03 & merged_res_not_preg_preg_NB_miRNA_a$pvalue<=0.03,]

rm(group_a,count_miRNA_plasma_a)

merged_res_not_preg_preg_NB_miRNA_a$Row.names
}

overlapping_genes_LOOCV<-Reduce(intersect, out_list_IDs_LOOCV)

merged_res_not_preg_preg_NB_miRNA<-merge(res_not_preg_preg_NB_miRNA_edgeR, res_not_preg_preg_NB_miRNA_DeSeq, by.x
="row.names", by.y="row")
merged_res_not_preg_preg_NB_miRNA_a<-merged_res_not_preg_preg_NB_miRNA[merged_res_not_preg_preg_NB_miRNA$Row.name
s %in% overlapping_genes_LOOCV,]
annotated_merged_res_not_preg_preg_NB_miRNA<-merge(merged_res_not_preg_preg_NB_miRNA_a,annotation.ensembl.symbol,
by.x="Row.names", by.y="ensembl_gene_id", all=FALSE)
annotated_merged_res_not_preg_preg_NB_miRNA<-annotated_merged_res_not_preg_preg_NB_miRNA[with(annotated_merged_re
s_not_preg_preg_NB_miRNA, order(logFC)), ]
dim(annotated_merged_res_not_preg_preg_NB_miRNA)#0
```

```
## [1]  0 17
```

```
#head(annotated_merged_res_not_preg_preg_NB_miRNA)
```

### Assessment of transcript levels as predictors of pregnancy outcome

#### Supplementary Fig. 5

```
group<-factor(c("Preg_AI","Preg_AI","Preg_AI","Preg_AI","Preg_AI","Preg_AI",
                "Not_Preg","Not_Preg","Not_Preg","Not_Preg","Not_Preg",
                "Preg_AI","Not_Preg", "Not_Preg", "Not_Preg", "Preg_AI",
                "Preg_AI","Preg_AI", "Preg_AI", "Not_Preg", "Preg_AI",
                "Not_Preg", "Not_Preg"), levels=c("Preg_AI", "Not_Preg"))

class<-data.frame(group=group)

merged_datasets_a<-merged_datasets[,c(1:6,13:17,18:29)]

#variance stablize the data

design<-model.matrix(~group)
colData<-data.frame("group"=group)
rownames(colData)<-colnames(data)
dds<-DESeqDataSetFromMatrix(countData=merged_datasets_a, colData=colData, design= ~group)
dds<-DESeq(dds)
vsd <- varianceStabilizingTransformation(dds, fitType='local')
data_vsd <- assay(vsd)

data_vsd<-data_vsd[rownames(data_vsd) %in% merged_res_not_preg_preg_AI_mRNA_merged_a$Row.names,]

data.train <- data.frame(t(data_vsd[ ,c(12:23)]))
data.test  <- data.frame(t(data_vsd[ ,c(1:11)]))
classtr <-  as.factor(group[c(12:23)])
classts <-  as.factor(group[c(1:11)])

set.seed(1)
number <-4
repeats <- 5

control <- trainControl(method = "repeatedcv",
                        number = number ,
                        repeats = repeats,
                        classProbs = TRUE,
                        savePredictions = "final",
                        index = createResample(classtr, repeats*number),
                        #summaryFunction = twoClassSummary,
                        #returnResamp = "all",
                        allowParallel = TRUE)


my_models<-c( "ada","AdaBag","adaboost","avNNet","bagEarth","bagEarthGCV","bagFDAGCV","bayesglm","blackboost","Bs
tLm","bstSm",
 "bstTree","C5.0","C5.0Cost","C5.0Rules","C5.0Tree","cforest","CSimca","ctree","ctree2","deepboost","dnn",
 "dwdLinear","dwdRadial","earth","FRBCS.CHI","FRBCS.W","gamboost","gamLoess","gamSpline","gaussprLinear","gausspr
Poly","gaussprRadial",
 "gcvEarth","glm","glmboost","glmnet","glmStepAIC","hdda","hdrda","kknn","knn","lda","lda2",
 "LogitBoost","Mlda","mlp","mlpML","mlpWeightDecay","mlpWeightDecayML", "monmlp","msaenet","multinom","naive_baye
s","nb",
 "nnet","null","ordinalNet","ORFlog","ORFpls","ORFridge","ORFsvm","ownn","pam","parRF","partDSA",
 "pcaNNet","pda","pda2","plr","pls","PRIM","protoclass","ranger","rbfDDA","Rborist","regLogistic",
 "rf","RFlda","rfRules","rlda","rocc" )


#initial testing of the models not run here due to time constrain

cl <- makeCluster(32)
registerDoParallel(cl)
rand_integer<-sample.int(10000, 50, replace = FALSE)
data_runs<-data.frame()
data_runs<-foreach(model = my_models,.combine='rbind', .packages=c('caret'), .inorder=FALSE, .verbose=TRUE , .err
orhandling='remove') %:%
foreach(i = rand_integer, .combine='rbind', .packages=c('caret'), .inorder=FALSE, .verbose=TRUE , .errorhandling=
'remove') %dopar% {
  rm(model_train)
  set.seed(i)
  model_train<-train(data.train, classtr,  method = model, trControl=control)
  data.frame(model,t(caret::confusionMatrix(predict(model_train, data.test),classts)$overall), seed=i)
}
stopCluster(cl)

#write.table(data_runs, file = paste(result_path,"2020_01_25_prediction_metrics_all_models.txt",sep="/"), append
 = FALSE, quote = FALSE, sep = "\t",  eol = "\n", na = "NA", dec = ".", row.names = FALSE, col.names = TRUE)

#system("bzip2 -9 /mnt/storage/auburn/heifer_pregnancy/proj_2018/analysis/results/2020_01_25_prediction_metrics_a
ll_models.txt")
```

```
data_runs<-read.delim(paste(result_path,"2020_01_25_prediction_metrics_all_models.txt.bz2",sep="/"), stringsAsFac
tors=FALSE)

sumary_models<-aggregate( Accuracy ~ model, data_runs,  FUN=function(x) c(mn = mean(x), median = median(x), std=s
d(x), n=length(x)))
sumary_models<-sumary_models[order(-sumary_models$Accuracy[,1]),]
#head(sumary_models, 10)

sumary_models<-sumary_models[1:15,]
sumary_models_1<-sumary_models$Accuracy
sumary_models_1<-data.frame(sumary_models_1)
sumary_models_1$model<-factor(sumary_models$model, levels=rev(sumary_models$model))


ggplot(sumary_models_1,aes(y=mn, x=model)) +
geom_pointrange(aes(ymin = mn-std, ymax = mn+std),colour="black",size=0.1)+
geom_point(size=2 )+
scale_y_continuous(name="Accuracy")+
coord_flip()+
theme_minimal()+
theme(
text = element_text(size=12),
axis.title.x = element_text(color="black"),
axis.text.x  = element_text(color="black"),
axis.text.y  = element_text(color="black"),
axis.title.y  = element_blank()
)
```

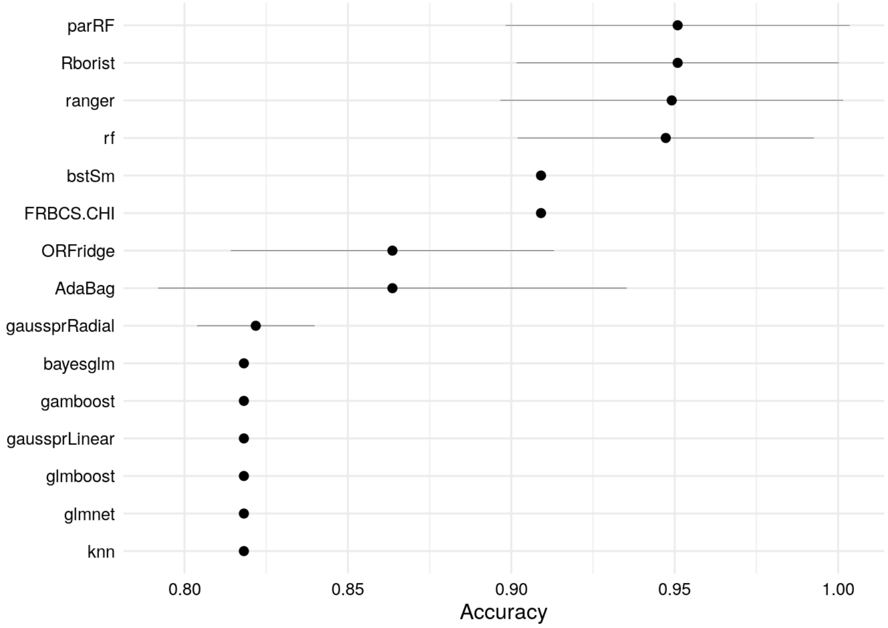

```
cl <- makeCluster(32)
registerDoParallel(cl)

rand_integer<-sample.int(10000, 2000, replace = FALSE)
data_runs<-data.frame()
data_runs<-foreach(i = rand_integer, .combine='rbind', .packages=c('caret'), .inorder=FALSE, .verbose=TRUE , .err
orhandling='remove') %dopar% {
  rm(model_train)
  set.seed(i)
  model_train<-train(data.train, classtr,  method = "parRF", trControl=control)
  data.frame(t(caret::confusionMatrix(predict(model_train, data.test),classts)$overall), seed=i)
}
stopCluster(cl)

attach(data_runs)
data_runs_b <- data_runs[order(-Accuracy),]
detach(data_runs)

write.table(data_runs_b, file= paste(result_path,"/2020_01_25_data_runs_b_2000_randomizations_parRF.txt", sep=""
), append = FALSE, quote = FALSE, sep = "\t" ,row.names = FALSE)
system("bzip2 -9 /mnt/storage/auburn/heifer_pregnancy/proj_2018/analysis/results/2020_01_25_data_runs_b_2000_rand
omizations_parRF.txt")


rand_integer<-sample.int(10000, 2000, replace = FALSE)
data_runs<-data.frame()
data_runs<-foreach(i = rand_integer, .combine='rbind', .packages=c('caret'), .inorder=FALSE, .verbose=TRUE , .err
orhandling='remove') %do% {
  rm(model_train)
  set.seed(i)
  model_train<-train(data.train, classtr,  method = "Rborist", trControl=control)
  data.frame(t(caret::confusionMatrix(predict(model_train, data.test),classts)$overall), seed=i)
}

attach(data_runs)
data_runs_b <- data_runs[order(-Accuracy),]
detach(data_runs)

write.table(data_runs_b, file= paste(result_path,"/2020_02_05_data_runs_b_2000_randomizations_Rborist.txt", sep=
""), append = FALSE, quote = FALSE, sep = "\t" ,row.names = FALSE)
system("bzip2 -9 /mnt/storage/auburn/heifer_pregnancy/proj_2018/analysis/results/2020_02_05_data_runs_b_2000_rand
omizations_Rborist.txt")


cl <- makeCluster(32)
registerDoParallel(cl)

rand_integer<-sample.int(10000, 2000, replace = FALSE)
data_runs<-data.frame()
data_runs<-foreach(i = rand_integer, .combine='rbind', .packages=c('caret'), .inorder=FALSE, .verbose=TRUE , .err
orhandling='remove') %dopar% {
  rm(model_train)
  set.seed(i)
  model_train<-train(data.train, classtr,  method = "ranger", trControl=control)
  data.frame(t(caret::confusionMatrix(predict(model_train, data.test),classts)$overall), seed=i)
}
stopCluster(cl)

attach(data_runs)
data_runs_b <- data_runs[order(-Accuracy),]
detach(data_runs)

write.table(data_runs_b, file= paste(result_path,"/2020_02_05_data_runs_b_2000_randomizations_ranger.txt", sep=""
), append = FALSE, quote = FALSE, sep = "\t" ,row.names = FALSE)
system("bzip2 -9 /mnt/storage/auburn/heifer_pregnancy/proj_2018/analysis/results/2020_02_05_data_runs_b_2000_rand
omizations_ranger.txt")


cl <- makeCluster(32)
registerDoParallel(cl)

rand_integer<-sample.int(10000, 2000, replace = FALSE)
data_runs<-data.frame()
data_runs<-foreach(i = rand_integer, .combine='rbind', .packages=c('caret'), .inorder=FALSE, .verbose=TRUE , .err
orhandling='remove') %dopar% {
  rm(model_train)
  set.seed(i)
  model_train<-train(data.train, classtr,  method = "rf", trControl=control)
  data.frame(t(caret::confusionMatrix(predict(model_train, data.test),classts)$overall), seed=i)
}
stopCluster(cl)
```

```
attach(data_runs)
data_runs_b <- data_runs[order(-Accuracy),]
detach(data_runs)

write.table(data_runs_b, file= paste(result_path,"/2020_02_05_data_runs_b_2000_randomizations_rf.txt", sep=""), a
ppend = FALSE, quote = FALSE, sep = "\t" ,row.names = FALSE)
system("bzip2 -9 /mnt/storage/auburn/heifer_pregnancy/proj_2018/analysis/results/2020_02_05_data_runs_b_2000_rand
omizations_rf.txt")



cl <- makeCluster(32)
registerDoParallel(cl)

rand_integer<-sample.int(10000, 2000, replace = FALSE)
data_runs<-data.frame()
data_runs<-foreach(i = rand_integer, .combine='rbind', .packages=c('caret'), .inorder=FALSE, .verbose=TRUE , .err
orhandling='remove') %dopar% {
  rm(model_train)
  set.seed(i)
  model_train<-train(data.train, classtr,  method = "bstSm", trControl=control)
  data.frame(t(caret::confusionMatrix(predict(model_train, data.test),classts)$overall), seed=i)
}
stopCluster(cl)

attach(data_runs)
data_runs_b <- data_runs[order(-Accuracy),]
detach(data_runs)

write.table(data_runs_b, file= paste(result_path,"/2020_01_25_data_runs_b_2000_randomizations_bstSm.txt", sep=""
), append = FALSE, quote = FALSE, sep = "\t" ,row.names = FALSE)
system("bzip2 -9 /mnt/storage/auburn/heifer_pregnancy/proj_2018/analysis/results/2020_01_25_data_runs_b_2000_rand
omizations_bstSm.txt")



cl <- makeCluster(32)
registerDoParallel(cl)

rand_integer<-sample.int(10000, 2000, replace = FALSE)
data_runs<-data.frame()
data_runs<-foreach(i = rand_integer, .combine='rbind', .packages=c('caret'), .inorder=FALSE, .verbose=TRUE , .err
orhandling='remove') %dopar% {
  rm(model_train)
  set.seed(i)
  model_train<-train(data.train, classtr,  method = "FRBCS.CHI", trControl=control)
  data.frame(t(caret::confusionMatrix(predict(model_train, data.test),classts)$overall), seed=i)
}
stopCluster(cl)

attach(data_runs)
data_runs_b <- data_runs[order(-Accuracy),]
detach(data_runs)

write.table(data_runs_b, file= paste(result_path,"/2020_01_25_data_runs_b_2000_randomizations_FRBCS.CHI.txt", sep
=""), append = FALSE, quote = FALSE, sep = "\t" ,row.names = FALSE)
system("bzip2 -9 /mnt/storage/auburn/heifer_pregnancy/proj_2018/analysis/results/2020_01_25_data_runs_b_2000_rand
omizations_FRBCS.CHI.txt")
```

### Figure 5d

```
acuracy_parRF<-read.delim("/mnt/storage/auburn/heifer_pregnancy/proj_2018/analysis/results/2020_01_25_data_runs_b
_2000_randomizations_parRF.txt.bz2")
table(acuracy_parRF$Accuracy)
```

```
## 
## 0.818181818181818 0.909090909090909                 1 
##                11              1063               926
```

```
acuracy_parRF$Accuracy<-round(acuracy_parRF$Accuracy,2)

acuracy_parRF_point<-round(unique(acuracy_parRF[,-c(2,8)]),2)

plot_point<-ggplot(acuracy_parRF_point, aes(y=Accuracy, x=as.factor(Accuracy)))+
geom_errorbar(aes(ymin = AccuracyLower, ymax = AccuracyUpper), width = 0.1)+
geom_point(size=2, color="red")+
scale_y_continuous(name=paste("Accuracy ", " CI", sep = "\u00B1"))+
theme_minimal()+
theme(
text = element_text(size=11),
axis.title.x = element_blank(),
axis.text.x  = element_blank(),
axis.text.y  = element_text(color="black"),
axis.title.y  = element_text(color="black"),
plot.margin = unit(c(0,0,0,0.3), "cm")
)

plot_histogram<-ggplot(acuracy_parRF, aes(as.factor(Accuracy)))+
geom_bar(aes(y = (..count..)/sum(..count..)),width=0.4)+
scale_x_discrete(name="Accuracy")+
scale_y_continuous(labels =percent_format(accuracy = 2) )+
coord_flip()+
theme_minimal()+
theme(
text = element_text(size=11),
axis.text.y  = element_text(color="black"),
axis.text.x  = element_text(color="black", angle=90),
axis.title.x  = element_blank(),
plot.margin = unit(c(0,0,0,0), "cm")
)

#png(filename=paste(result_path,"2020_02_10_accuracy_prediction_parRF.png",sep="/"), width=750, height=450, res=3
00, bg="transparent")
#grid.arrange( plot_histogram,plot_point, nrow = 1)
#dev.off()

grid.arrange( plot_histogram,plot_point, nrow = 1)
```

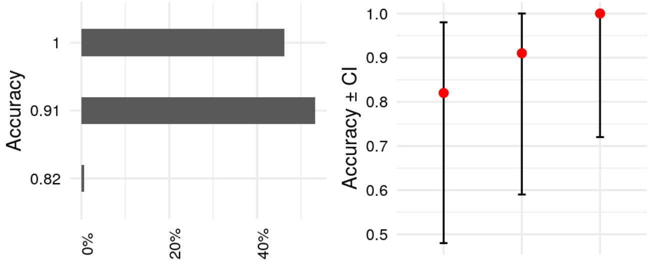


### Figure 5e

```
acuracy_parRF<-read.delim("/mnt/storage/auburn/heifer_pregnancy/proj_2018/analysis/results/2020_01_25_data_runs_b
_2000_randomizations_parRF.txt.bz2")
table(acuracy_parRF$Accuracy)
acuracy_parRF$Accuracy<-round(acuracy_parRF$Accuracy,2)

acuracy_parRF_point<-round(unique(acuracy_parRF[,-c(2,8)]),2)

acuracy_parRF_09<-acuracy_parRF[acuracy_parRF$Accuracy>=0.9 ,]

cl <- makeCluster(32)
registerDoParallel(cl)
variable_importance_09<-data.frame()
variable_importance_09<-foreach(i = acuracy_parRF_09$seed, .combine='cbind', .packages=c('caret'), .inorder=FALSE
, .verbose=TRUE , .errorhandling='remove') %dopar% {
  rm(model_train)
  set.seed(i)
  model_train<-train(data.train, classtr,  method = "parRF", trControl=control)
  importance<-varImp(model_train, scale=F)$importance
  importance$gene<-(rownames(importance))
  importance<-importance[with(importance, order(gene)),]$Overall
}
stopCluster(cl)

rownames(variable_importance_09)<-colnames(data.train)

write.table(variable_importance_09, file= paste(result_path,"/2020_03_11_variable_importance_parRF.txt", sep=""),
append = FALSE, quote = FALSE, sep = "\t" ,row.names = TRUE)
system("bzip2 -9 /mnt/storage/auburn/heifer_pregnancy/proj_2018/analysis/results/2020_03_11_variable_importance_p
arRF.txt")
```

```
variable_importance_09<-as.matrix(read.delim(paste(result_path,"/2020_03_11_variable_importance_parRF.txt.bz2", s
ep=""),row.names=1))
variable_importance_09<-cbind(variable_importance_09,average=rowMeans2(variable_importance_09), std=rowVars(varia
ble_importance_09,std = TRUE))

variable_importance_09<-variable_importance_09[order(-variable_importance_09[,1990]),]

variable_importance_09_plot<-as.data.frame(variable_importance_09[1:5,1990:1991 ])
variable_importance_09_plot$genes<-factor(rownames(variable_importance_09_plot),levels=rev(c("ENSBTAG00000049161"
,"ENSBTAG00000051249","ENSBTAG00000002615","ENSBTAG00000017243","ENSBTAG00000001412")))

plot_point<-ggplot(variable_importance_09_plot, aes(y=average, x=genes))+
geom_pointrange(aes(ymin = average-std, ymax = average+std),colour="blue",size=0.1)+
geom_point(size=0.4, colour="blue" )+
scale_y_continuous(name="Importance")+
scale_x_discrete(labels=c("N6AMT1", "GATA3", "LONRF3", "SMIM26", "RPL39"))+
coord_flip()+
theme_minimal()+
theme(
text = element_text(size=12),
axis.title.x = element_text(color="black"),
axis.text.x  = element_text(color="black"),
axis.text.y  = element_text(color="black", face="italic"),
axis.title.y  = element_blank()
)

#png(filename=paste(result_path,"2020_02_10_gene_importance.png",sep="/"), width=600, height=450, res=310, bg="tr
ansparent")
#plot_point
#dev.off()
plot_point
```

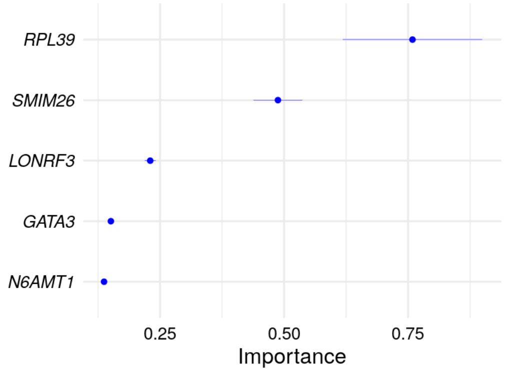

### Figure 5f

```
variable_importance_09_plot$gene_symbols<-c("RPL39","SMIM26" ,"LONRF3", "GATA3" , "N6AMT1")

cpm_AIpreg<-cpm_merged_datasets[,c(1,2,3,4,5,6,18,22,23,24,25,27)]
cpm_NOTpreg<-cpm_merged_datasets[,c(13,14,15,16,17,19,20,21,26,28,29)]


data_chart_1<-data.frame( stringsAsFactors=FALSE)
data_chart_2<-data.frame( stringsAsFactors=FALSE)

for (i in  seq(dim(variable_importance_09_plot)[1])){

  gene_id<-variable_importance_09_plot$genes[i]
  gene_symbol<-variable_importance_09_plot$gene_symbols[i]
  if(gene_symbol==""){gene_symbol<-gene_id}

  cpm_AI<-cpm_AIpreg[rownames(cpm_AIpreg)==gene_id,]
  cpm_NP<-cpm_NOTpreg[rownames(cpm_NOTpreg)==gene_id,]

  data_chart_1<-rbind(data.frame(cpm=cpm_AI), data.frame(cpm=cpm_NP))
  data_chart_1$group<-as.factor(rep(c("Preg_AI","Not_Preg"),c(12,11)))
  data_chart_1$shape<-as.factor(rep(c(2,1,2,1),c(6,6,5,6)) )
  data_chart_1$gene<-gene_symbol
  data_chart_1$chart<-i
  data_chart_1$fold_change<-DEG_AI_NP_mRNA_merged[DEG_AI_NP_mRNA_merged$Row.names==gene_id,]$logFC
  data_chart_1$pvalue<-DEG_AI_NP_mRNA_merged[DEG_AI_NP_mRNA_merged$Row.names==gene_id,]$PValue

  data_chart_2<-rbind(data_chart_2,data_chart_1)
}

data_chart_2$gene<-factor(data_chart_2$gene,levels=c("RPL39","SMIM26" ,"LONRF3", "GATA3" , "N6AMT1"))


plot<-ggplot(data=data_chart_2, aes(x=group , y=cpm, colour=group, shape=shape) )+
      geom_jitter(position = position_jitterdodge(dodge.width = 1), size=2)+
      scale_shape_manual(values=c(15,16), labels=c("year 1", "year 2"))+
      scale_colour_manual(name=NULL, values = c("#D55E00", "#0072B2"),labels=c("non preg", "AI preg"))+
      #stat_summary(fun.y = median, fun.ymin = median, fun.ymax = median,colour = "gray", size = 0.2,  geom = "cr
ossbar", alpha=0.5)+
      scale_y_continuous("CPM")+
      facet_wrap(~gene, scales="free", nrow=1)+
      theme_bw()+
      theme(panel.grid= element_blank(),
            panel.background = element_blank(),
            panel.grid.minor = element_blank(),
            panel.grid.major = element_blank(),
            plot.background = element_blank(),
            strip.background = element_rect(fill = "white"),
            strip.text.x = element_text(colour = 'black', face="italic",size = 10),
            legend.text = element_text( colour = 'black', size = 10 ),
            axis.text.x = element_blank(),
            axis.text.y = element_text( colour = 'black', size = 10 ),
            axis.title.y = element_text( colour = 'black', size = 10 ),
            axis.title.x = element_blank(),
            #legend.key.size = unit(0.9,"cm"),
            legend.position="bottom",
            legend.margin=margin(0,0,0,0),
            legend.box.margin=margin(-10,-10,-10,-10),
            legend.title = element_blank(),
            axis.ticks.x = element_blank()
      )

#png(file=paste(result_path,"2020_02_10_plot_DEGs_predictors_random_forest.png",sep="/"), width= 1100, height= 49
0,  bg="transparent", res=300)
#plot
#dev.off()
plot
```

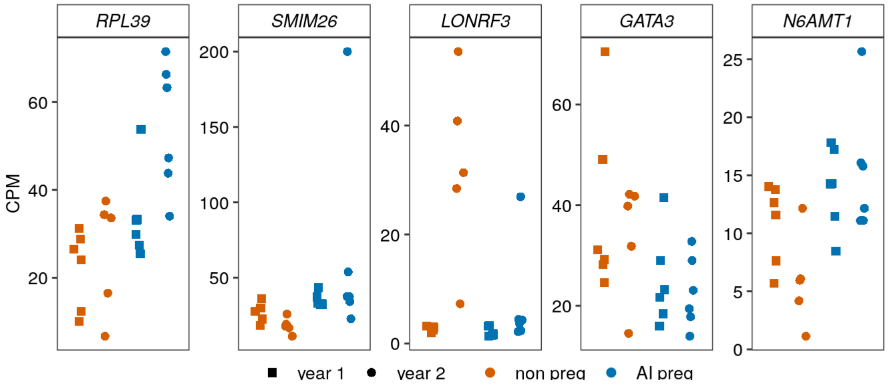


```
sessionInfo()
```

```
## R version 4.0.1 (2020-06-06)
## Platform: x86_64-pc-linux-gnu (64-bit)
## Running under: Ubuntu 18.04.4 LTS
## 
## Matrix products: default
## BLAS:   /usr/lib/x86_64-linux-gnu/blas/libblas.so.3.7.1
## LAPACK: /usr/lib/x86_64-linux-gnu/lapack/liblapack.so.3.7.1
## 
## locale:
##  [1] LC_CTYPE=en_US.UTF-8       LC_NUMERIC=C              
##  [3] LC_TIME=en_US.UTF-8        LC_COLLATE=en_US.UTF-8    
##  [5] LC_MONETARY=en_US.UTF-8    LC_MESSAGES=en_US.UTF-8   
##  [7] LC_PAPER=en_US.UTF-8       LC_NAME=C                 
##  [9] LC_ADDRESS=C               LC_TELEPHONE=C            
## [11] LC_MEASUREMENT=en_US.UTF-8 LC_IDENTIFICATION=C       
## 
## attached base packages:
##  [1] grid      parallel  stats4    stats     graphics  grDevices utils    
##  [8] datasets  methods   base     
## 
## other attached packages:
##  [1] networkD3_0.4               stringi_1.4.6              
##  [3] scales_1.1.1                TestCor_0.0.1.1            
##  [5] caretEnsemble_2.0.1         caret_6.0-86               
##  [7] lattice_0.20-41             circlize_0.4.10            
##  [9] ComplexHeatmap_2.4.2        parallelDist_0.2.4         
## [11] flashClust_1.01-2           gridExtra_2.3              
## [13] ggpubr_0.3.0                Rtsne_0.15                 
## [15] DT_0.13                     org.Bt.eg.db_3.11.4        
## [17] AnnotationDbi_1.50.0        gtools_3.8.2               
## [19] doParallel_1.0.15           iterators_1.0.12           
## [21] foreach_1.5.0               bigmemory_4.5.36           
## [23] reshape2_1.4.4              WGCNA_1.69                 
## [25] fastcluster_1.1.25          dynamicTreeCut_1.63-1      
## [27] goseq_1.40.0                geneLenDataBase_1.24.0     
## [29] BiasedUrn_1.07              biomaRt_2.44.1             
## [31] ggrepel_0.8.2               ggplot2_3.3.2              
## [33] statmod_1.4.34              edgeR_3.30.3               
## [35] limma_3.44.3                DESeq2_1.28.1              
## [37] SummarizedExperiment_1.18.1 DelayedArray_0.14.0        
## [39] matrixStats_0.56.0          Biobase_2.48.0             
## [41] GenomicRanges_1.40.0        GenomeInfoDb_1.24.2        
## [43] IRanges_2.22.2              S4Vectors_0.26.1           
## [45] BiocGenerics_0.34.0        
## 
## loaded via a namespace (and not attached):
##   [1] tidyselect_1.1.0         RSQLite_2.2.0            htmlwidgets_1.5.1       
##   [4] BiocParallel_1.22.0      pROC_1.16.2              munsell_0.5.0           
##   [7] codetools_0.2-16         preprocessCore_1.50.0    withr_2.2.0             
##  [10] colorspace_1.4-1         highr_0.8                knitr_1.28              
##  [13] rstudioapi_0.11          ggsignif_0.6.0           labeling_0.3            
##  [16] GenomeInfoDbData_1.2.3   farver_2.0.3             bit64_0.9-7             
##  [19] vctrs_0.3.1              generics_0.0.2           ipred_0.9-9             
##  [22] xfun_0.15                BiocFileCache_1.12.0     R6_2.4.1                
##  [25] markdown_1.1             clue_0.3-57              locfit_1.5-9.4          
##  [28] bitops_1.0-6             assertthat_0.2.1         nnet_7.3-14             
##  [31] gtable_0.3.0             timeDate_3043.102        rlang_0.4.6             
##  [34] genefilter_1.70.0        GlobalOptions_0.1.2      splines_4.0.1           
##  [37] rtracklayer_1.48.0       rstatix_0.6.0            ModelMetrics_1.2.2.2    
##  [40] acepack_1.4.1            impute_1.62.0            broom_0.5.6             
##  [43] checkmate_2.0.0          yaml_2.2.1               abind_1.4-5             
##  [46] GenomicFeatures_1.40.0   backports_1.1.8          Hmisc_4.4-0             
##  [49] tools_4.0.1              lava_1.6.7               ellipsis_0.3.1          
##  [52] RColorBrewer_1.1-2       Rcpp_1.0.4.6             plyr_1.8.6              
##  [55] base64enc_0.1-3          progress_1.2.2           zlibbioc_1.34.0         
##  [58] purrr_0.3.4              RCurl_1.98-1.2           prettyunits_1.1.1       
##  [61] rpart_4.1-15             openssl_1.4.1            pbapply_1.4-2           
##  [64] GetoptLong_1.0.0         cowplot_1.0.0            haven_2.3.1             
##  [67] cluster_2.1.0            magrittr_1.5             data.table_1.12.8       
##  [70] openxlsx_4.1.5           hms_0.5.3                mime_0.9                
##  [73] evaluate_0.14            xtable_1.8-4             XML_3.99-0.3            
##  [76] rio_0.5.16               jpeg_0.1-8.1             readxl_1.3.1            
##  [79] shape_1.4.4              compiler_4.0.1           tibble_3.0.1            
##  [82] crayon_1.3.4             htmltools_0.5.0          mgcv_1.8-31             
##  [85] Formula_1.2-3            tidyr_1.1.0              geneplotter_1.66.0      
##  [88] RcppParallel_5.0.1       lubridate_1.7.9          DBI_1.1.0               
##  [91] dbplyr_1.4.4             MASS_7.3-51.6            rappdirs_0.3.1          
##  [94] Matrix_1.2-18            car_3.0-8                igraph_1.2.5            
##  [97] gower_0.2.1              forcats_0.5.0            pkgconfig_2.0.3         
## [100] bigmemory.sri_0.1.3      GenomicAlignments_1.24.0 foreign_0.8-80          
## [103] recipes_0.1.12           annotate_1.66.0          XVector_0.28.0          
```

```
## [106] prodlim_2019.11.13       stringr_1.4.0            digest_0.6.25
## [109] Biostrings_2.56.0        rmarkdown_2.3            cellranger_1.1.0
## [112] htmlTable_2.0.0          curl_4.3                 Rsamtools_2.4.0
## [115] rjson_0.2.20             jsonlite_1.6.1           lifecycle_0.2.0
## [118] nlme_3.1-148             carData_3.0-4            askpass_1.1
## [121] pillar_1.4.4             httr_1.4.1               survival_3.2-3
## [124] GO.db_3.11.4             glue_1.4.1               zip_2.0.4
## [127] png_0.1-7                bit_1.1-15.2             class_7.3-17
## [130] blob_1.2.1               latticeExtra_0.6-29      memoise_1.1.0
## [133] dplyr_1.0.0
```
