## Supplementary figures for "Rewiring of gene expression in circulating white blood cells is associated with pregnancy outcome in heifers (Bos taurus)"

**This PDF file includes:**

Supplementary Fig. S1 2
Supplementary Fig. S2 3
Supplementary Fig. S3 4
Supplementary Fig. S4 5
Supplementary Fig. S5 6

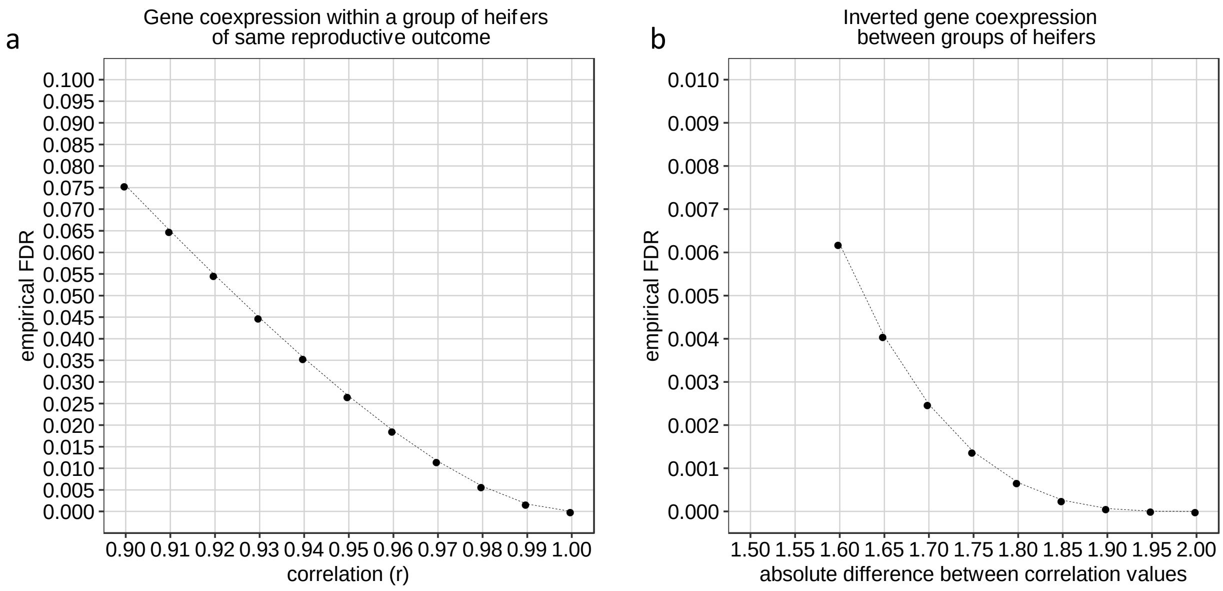


Supplementary Fig. S1. Empirical false discovery rate for within group mRNA:mRNA gene co-expression (a) and (b) inverted differential co-expression.

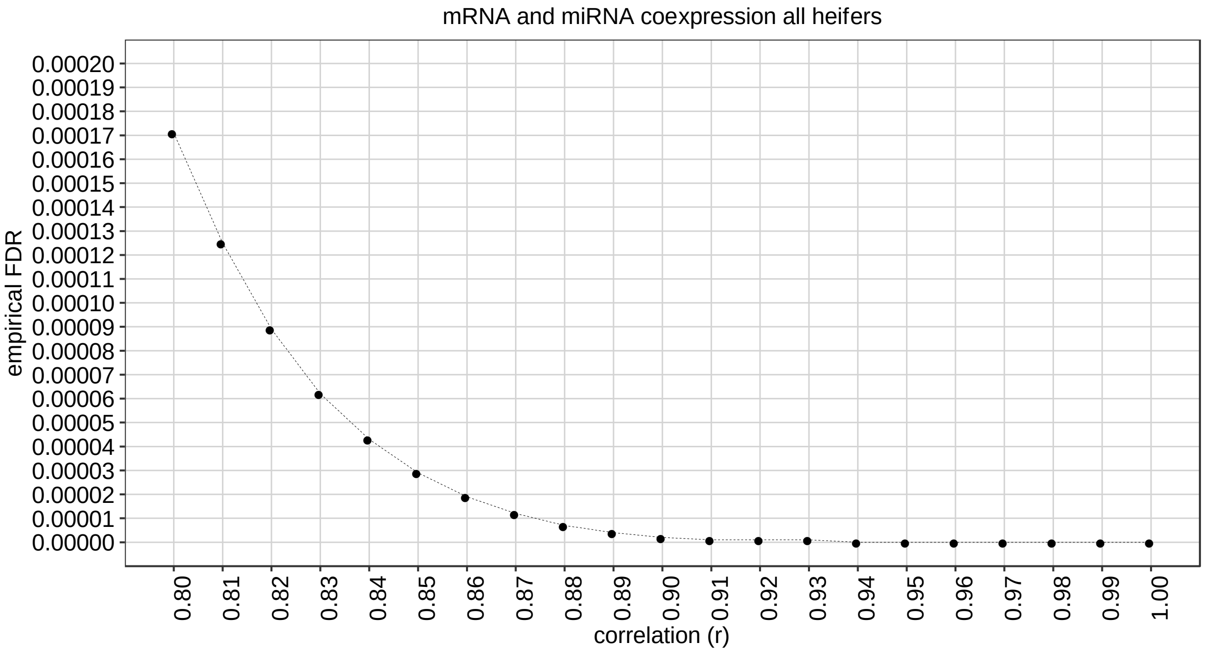


Supplementary Fig. S2. Empirical false discovery rate for correlation between mRNA and miRNA.

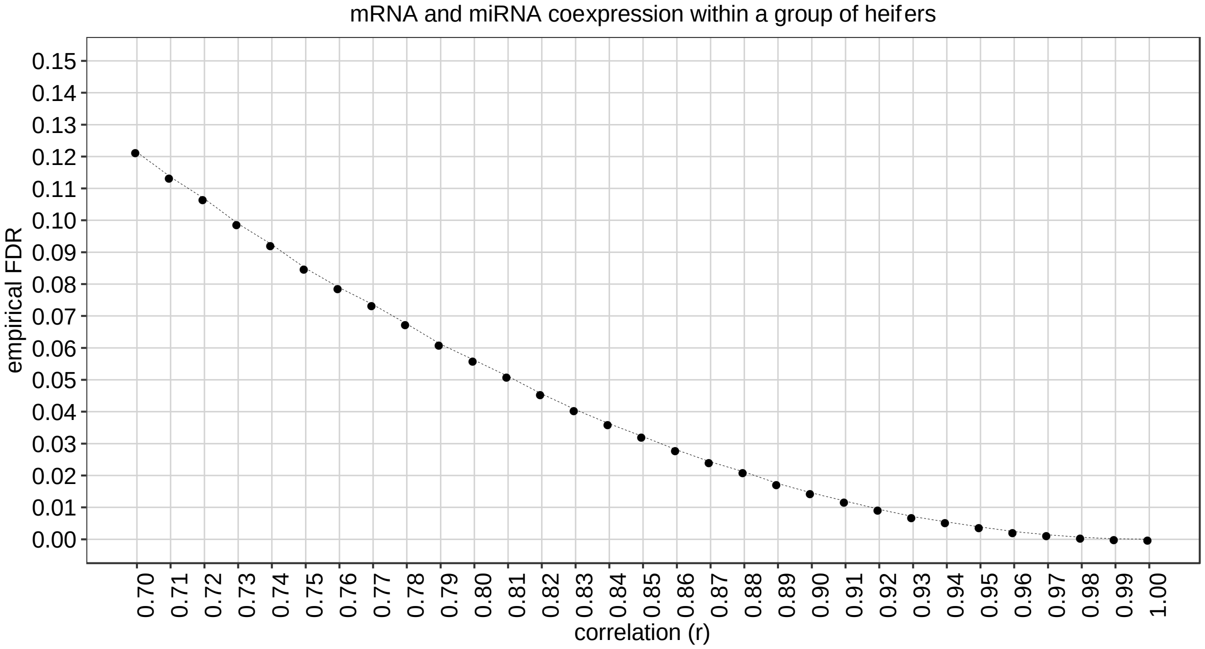


Supplementary Fig. S3. Empirical false discovery rate for correlation between mRNA and miRNA within a group of heifers (n=6).

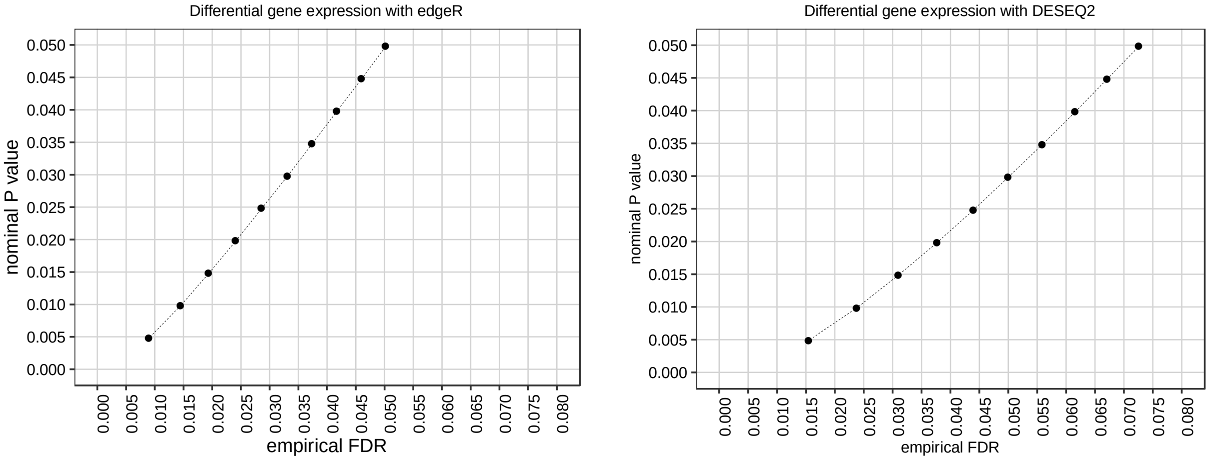


Supplementary Fig. S4. Empirical false discovery rate for differential gene expression considering both edgeR and DESEQ2 Bioconductor packages estimated after 10,000 randomizations.

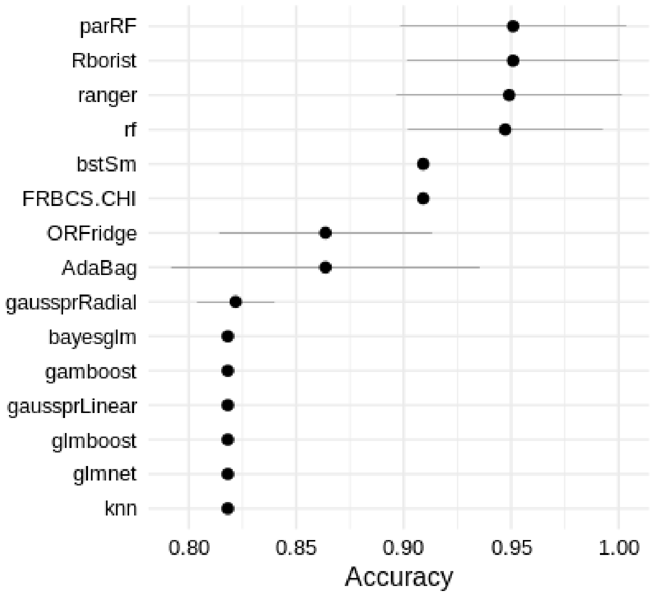

Supplementary Fig. S5. Average and standard deviation of the accuracies calculated for 50 randomizations on different machine learning models. Only 15 models that generated the 15 highest accuracy averages are shown.
